# Whole-Brain Millisecond-Scale Effective Connectivity Atlas of Auditory and Visual Naming

**DOI:** 10.1101/2025.11.18.689014

**Authors:** Ryuzaburo Kochi, Aya Kanno, Hiroshi Uda, Keisuke Hatano, Masaki Sonoda, Hidenori Endo, Michael Cools, Robert Rothermel, Aimee F. Luat, Eishi Asano

## Abstract

Neurobiological models suggest that speech relies on interactions among distant cortical regions interconnected by white matter pathways. Prior studies have shown that speech-related functional coactivation—simultaneous high-gamma augmentation across distributed regions—reflects underlying neural interactions, as validated by electrical stimulation mapping. However, it remains unclear *at the whole-brain level* when, for how long, and in which directions cortical regions transmit information, and how dynamic information flows contribute to speech. Here, we investigated the causal roles of directional neural information flow during auditory and visual naming using intracranial EEG from 9,526 artifact-free nonepileptic sites across 127 patients. Information flow, estimated by transfer entropy–based effective connectivity, was classified as *excitatory* when high-gamma acceleration in one region predicted subsequent acceleration in another, and *inhibitory* when deceleration predicted downstream deceleration. Following stimulus onset, excitatory flows emerged from modality-specific sensory cortices and propagated to higher-order regions, later becoming bidirectional as functional coactivation developed. Each excitatory flow event was transient (<500 ms), typically followed by inhibitory flows and decreased coactivation, during which excitatory flows often emerged along new pathways. During auditory naming, faster response times were associated with stronger excitatory flows in left perisylvian regions; during visual naming, faster responses were linked to stronger flows in bilateral basal temporal cortices. Stronger excitatory flows at specific time points predicted a higher probability of stimulation-induced symptoms (Spearman’s rho: 0.54–0.81; p<0.00001), whereas associations with inhibitory flows peaked later. Excitatory flows near visual naming response onset were particularly associated with stimulation-induced speech arrest and face sensorimotor symptoms, whereas functional coactivation alone failed to reveal comparable associations. These findings demonstrate that transient acceleration of directional neural interactions through white matter supports successive stages of speech processing. As activity within one pathway decelerates, excitatory flow accelerates along another, enabling functional transitions critical for naming. By establishing the causal contribution of temporally precise, anatomically specific white matter pathways, this study substantiates and extends existing neurobiological models of speech. To facilitate replication and dynamic whole-brain visualization, we provide open access to the full dataset (62.15 GB) and analysis code.

## Introduction

Speech arises from rapid, large-scale neural interactions distributed across cortical regions and coordinated through long-range white matter pathways [1–7]. These interactions unfold on a millisecond timescale and depend on the dynamic exchange of information across structurally interconnected, functionally specialized circuits. Capturing their spatiotemporal organization at the whole-brain level remains a major challenge, requiring integrative approaches that simultaneously resolve directional flow dynamics, structural connectivity pathways, and underlying functional information [8]. Although printed schematic models outline key cortical hubs and tracts, they often fail to capture the temporal dynamics of neural interactions across distributed pathways. Dynamic visualizations can reveal how transient, time-locked activity spreads across functionally and anatomically coupled regions [9–10]. The causal necessity of these regions has long been inferred from direct electrical stimulation mapping, where transient behavioral impairments mark sites essential for speech processing [11–13]. Such stimulation responses serve as a ground truth for evaluating the functional relevance of network-level measurements [14]. Building on this framework, our prior work using intracranial EEG (iEEG) showed that structurally connected cortical regions coactivate during speech at high temporal resolution (5-ms steps), and that the timing and intensity of coactivation predict the emergence of specific stimulation-induced symptoms [9–10]. These findings support the view that dynamic interactions mediated by white matter are not merely correlated with speech—they are causally involved. Yet a fundamental question remains unanswered: whether and when directional information flows occur between cortical regions around functional coactivation, and how these flows contribute to speech generation.

To address this question, we analyzed iEEG data from a large patient cohort to resolve the timing, directionality, and white matter pathways underlying neural information flows during auditory and visual naming. By linking these temporally resolved flows to electrical stimulation outcomes and behavioral response times, we identified distinct phases of modality-specific speech processing—including perception, semantic access, lexical retrieval, response planning, and articulatory execution [10]. Because perceived speech stimuli must be relayed from primary sensory cortices to downstream conceptual and motor systems [15–17], we hypothesized that unidirectional information flow from sensory to higher-order areas would precede functional coactivation, and that bidirectional interactions would emerge later. Consistent with prior work showing peri-response interactions in perisylvian frontotemporal and occipitotemporal circuits during auditory and visual language tasks, respectively [14, 18–19], we also expected white matter pathways linking these regions to mediate critical functional transitions. Given that lesions disrupting the arcuate fasciculus and inferior longitudinal fasciculi lead to persistent speech impairments [7,20–21], we predicted that stronger information flows along these tracts would be associated with both faster naming and a higher probability of stimulation-induced speech symptoms. While previous studies have examined individual elements—such as stimulation mapping or functional coactivation—this is, to our knowledge, the first large-cohort study to integrate *whole-brain* visualization of directionally specific neural information flows through white matter with direct measures of behavior and stimulation-based causality. We studied patients undergoing iEEG as part of presurgical evaluation for focal epilepsy, using auditory and visual naming tasks to elicit modality-specific speech processes. Neural information flow was quantified using transfer entropy–based effective connectivity analysis [22–26], which detects directed statistical dependence between time series. Flows were classified as *excitatory* when acceleration of high-gamma activity (70–110 Hz) in one region predicted subsequent acceleration in another connected region, and *inhibitory* when deceleration predicted downstream deceleration. In parallel, we measured functional coactivation and co-deactivation, defined as sustained, simultaneous high-gamma augmentation or attenuation across structurally connected cortical regions [9–10]. We focused on high-gamma amplitude because it tightly correlates with population-level neuronal firing [27–28], metabolic demand [29], and hemodynamic activation [30], while offering superior temporal resolution compared to lower-frequency oscillations. Recent work shows that task-related high-gamma modulations outperform slower bands in predicting postoperative cognitive outcomes, further reinforcing its functional relevance [31–32].

## Materials and methods

### Participants

From January 2007 through December 2024, we recruited individuals with focal seizures who fulfilled the eligibility criteria [10]. Eligible participants were native English speakers, four years of age or older, who completed auditory or visual naming tasks during extraoperative iEEG at the Detroit Medical Center (**eFigure 1**). We excluded those with sensory deficits, a history of epilepsy surgery, major malformations, or evidence of right-hemisphere language dominance, determined by Wada testing or by the combination of left-handedness and congenital left-hemispheric cortical lesions. In the present study, with these criteria, the left hemisphere was assumed to contain essential language functions [31,33].

### Intracranial electrodes

Intracranial implantation of platinum disk or depth electrodes was performed [10,31]. A cortical surface reconstruction was generated, and electrode locations were determined by registering the post-implant CT scan to the preoperative MRI using FreeSurfer (http://surfer.nmr.mgh.harvard.edu) [34]. Electrode coordinates were transformed into the FreeSurfer standard brain space and mapped to predefined regions of interest (ROIs) for group-level analysis (**eFigure 2**) [35].

### Structural connectivity

We assessed structural connectivity between ROIs using tractography data from the Human Connectome Project (http://brain.labsolver.org/diffusion-mri-templates/hcp-842-hcp-1021) [36], following procedures described in our previous studies [10,35]. This approach is appropriate, as the goal of the present study was to model neural interactions that are representative of broader populations rather than tailored to a single patient. Our prior analysis demonstrated that white-matter trajectories in children with drug-resistant epilepsy are highly consistent with those observed in healthy individuals (r: 0.80) [37]. White-matter connections linking cortical ROI pairs were visualized in Montreal Neurological Institute space using DSI Studio (http://dsi-studio.labsolver.org/). Specific parameters for tract reconstruction are detailed previously [10], and the corresponding white-matter imaging dataset is publicly available, as indicated below.

### iEEG

After implantation, patients were monitored with video-iEEG using a Nihon Kohden system (Foothill Ranch, CA), with signals digitized at 1,000 Hz. Analyses were restricted to artifact-free electrodes positioned at nonepileptic regions, excluding those related to seizure onset, interictal epileptiform discharges, or MRI-visible lesions.

### Tasks

Participants completed both auditory and visual naming paradigms [10,31]. For the auditory condition, up to 100 spoken questions (e.g., “What flies in the sky?”) required brief noun responses. Stimulus and response onsets were tracked with a photosensor and microphone, and response latency was calculated from the end of the auditory cue to the response onset. In the visual condition, up to 60 grayscale pictures (e.g., “dog”) were presented on a screen for overt naming, with response latency defined from stimulus onset to response onset. Trials lacking valid responses were excluded from analysis.

### Time-frequency analysis

We adopted the same time–frequency analysis as described in our prior report [10]. High-gamma amplitude (70–110 Hz) was extracted using a bipolar montage, and a Gabor transform with 5-ms windows and 10-Hz frequency bins was performed in BESA Software (BESA, Gräfelfing, Germany) [38]. Amplitude values, defined as the square root of power, were converted into percent changes relative to a 400-ms baseline spanning 200–600 ms before stimulus onset. iEEG recordings were aligned to stimulus onset, stimulus offset, and response onset during auditory naming, and to stimulus and response onsets during visual naming. Percent changes in high-gamma amplitude were computed within −200 to +500 ms of stimulus onset, −500 to +500 ms of stimulus offset, and −500 to +500 ms of response onset. The spatial distribution of high-gamma modulation was rendered on a FreeSurfer cortical surface with 10-mm interpolation in MATLAB R2023 (MathWorks, Natick, MA).

### Group-level assessment of regional high-gamma amplitudes

We quantified group-level high-gamma amplitudes at each of the 66 ROIs, with every ROI informed by data from a minimum of seven patients [39]. This group-level approach offers a valid framework for modeling population-level interactions [40–41], analogous to cross-sectional designs or meta-analyses where longitudinal data are lacking or a single cohort dataset is insufficient. Because comprehensive bilateral sampling of nonepileptic regions is neither ethical nor feasible, analysis of individual iEEG data alone cannot capture neural interactions at the whole-brain level. Group-level mean high-gamma measures from sufficiently large samples are expected to be robust and reproducible across populations, as detailed in our previous studies [9–10]. To confirm robustness of group-level high-gamma amplitudes at each ROI, we displayed temporal dynamics with confidence intervals and a leave-one-out approach (**eFigures 3–6**); Spearman’s tests demonstrated consistency in mean high-gamma amplitudes across datasets each excluding one patient (rho values: ≥0.95; p-values: <0.00001).

We assessed the timing and regional distribution of high-gamma acceleration, deceleration, augmentation, and attenuation at each ROI. **eFigures 3-4** illustrate the distinctions between acceleration and augmentation, as well as between deceleration and attenuation. The timing of high-gamma acceleration and deceleration was used to define neural information flow between ROIs, whereas the timing of augmentation and attenuation independently defined functional co-activation and co-deactivation [10]. Precisely, high-gamma acceleration (deceleration) was considered significant when five consecutive 5-ms bins showed amplitudes significantly greater (smaller) than the mean of the preceding 50-ms time window (i.e., ten 5-ms bins), as determined by t-tests. High-gamma amplitude was classified as augmented (attenuated) when it exceeded (fell below) the 400-ms baseline before stimulus presentation. Augmentation was considered significant when the lower bound of the 99.99% confidence interval exceeded zero for at least five consecutive 5-ms bins, whereas attenuation was defined when the upper bound remained below zero for the same duration [10]. Accordingly, for example, if amplitude remained above baseline but declined relative to the immediately preceding period, high-gamma activity was classified as augmented with deceleration.

### Excitatory and inhibitory neural information flows

We quantified the strength and direction of *excitatory* and *inhibitory* neural information flow along white matter pathways using a transfer entropy–based effective connectivity analysis [22–26]. Transfer entropy was computed within 100-ms windows sliding in 5-ms steps, with each 5-ms bin coded as 1 when high-gamma acceleration was significant and 0 otherwise. This binary time series was then used as input to the transfer entropy algorithm [24] to estimate information flows between ROI pairs. Excitatory flow from ROI_1_ to ROI_2_ was considered present when two conditions were met: (i) ROI_1_ and ROI_2_ were structurally connected by a white matter tract, and (ii) transfer entropy was greater than zero, such that significant high-gamma acceleration at ROI_1_ predicted subsequent acceleration at ROI_2_. Under this definition, the transfer entropy value was interpreted as efferent flow at ROI_1_ and afferent flow at ROI_2_. For visualization, the strength of information flow at each 5-ms bin was estimated by computing a Gaussian-weighted summation of transfer entropy values across the corresponding 100-ms window. Inhibitory information flow was defined analogously, when significant high-gamma deceleration at ROI_1_ predicted subsequent deceleration at ROI_2_.

We tested the hypothesis that, during the auditory naming task, excitatory information flow would begin as unidirectional from the left posterior superior temporal gyrus (pSTG) to the posterior middle temporal gyrus (pMTG) and subsequently become bidirectional. During the visual naming task, we tested the hypothesis that excitatory flow would likewise begin as unidirectional from the lateral occipital gyrus to the posterior fusiform gyrus and later evolve into bidirectional interactions within each hemisphere.

### Functional coactivation and co-deactivation

We analyzed temporal windows and white matter pathways exhibiting functional coactivation or co-deactivation [9–10]. ROI pairs were classified as coactivated or co-deactivated when (i) both regions showed significant high-gamma modulations (as compared to the baseline before stimulus onset) in the same direction for at least five consecutive 5-ms bins, and (ii) the regions were directly connected by a streamline on tractography. These interactions were evaluated in 5-ms steps. The strength of functional coactivation or co-deactivation was quantified as the square root of the product of the high-gamma percent changes of the paired ROIs.

### Effect of patient demographics and clinical factors on behavioral and neural measures

In our prior investigation involving 125 patients, we found that older age correlated with faster median response times, whereas both left-handedness and the presence of a left-hemispheric epileptogenic focus were linked to reduced left-hemispheric dominance of high-gamma augmentation [10]. In the present cohort, we used mixed-effects modeling to test whether these associations could be replicated (**eResults**).

### Stimulation

Electrical stimulation mapping was used to validate excitatory flows and functional co-activation. Full methodological descriptions are provided elsewhere [9–10,31]. During 50-Hz stimulation of neighboring electrode pairs, patients were asked to answer *wh*-questions or name objects, and blinded examiners confirmed reproducible responses. Analyzed manifestations included auditory hallucinations, receptive and expressive aphasia, speech arrest, facial sensorimotor phenomena, phosphenes, visual distortions, and visual-naming errors.

### Association between neural dynamics and stimulation-induced manifestations

We examined whether the intensity of excitatory flow at specific time bins, quantified by transfer entropy, was associated with the probability of given stimulation-induced symptoms [9–10]. For each cortical ROI, efferent and afferent flow intensities were computed every 5 ms by averaging transfer entropy values across all ROI pairs. Spearman’s rank correlation was then used to test whether higher excitatory flow intensity at an ROI was associated with a greater probability (%) of specific symptoms. Statistical significance was defined as a Bonferroni-corrected, two-sided p < 0.05 across 500 time bins. We likewise examined the association between inhibitory flows and stimulation-induced symptoms.

### Neural correlates of within-individual variability in response times

We examined the relationship between excitatory flow and response times, while controlling for inter-patient variability by stratifying each patient’s trials into six response time categories. For each category, we computed the mean flow intensity across all ROI pairs and assessed its association with response times using Spearman’s rank correlation [10]. We hypothesized that stronger excitatory flow at the cortices along the left arcuate fasciculus and bilateral inferior longitudinal fasciculi would be associated with shorter response times during auditory and visual naming tasks, respectively. We likewise examined the association between inhibitory flows and response times.

## Results

### Impact of patient profiles on response time and neural measures

We found 127 eligible patients (age 5–49 years; 61 females; **eTable 1**); 121 completed the auditory task, 110 the visual task, and 104 completed both. We analyzed 9,526 artifact-free nonepileptic sites (9,178 for auditory, 8,585 for visual naming; **eTable 2**).

The median response times, averaged across patients, were 1.46 and 1.38 seconds for auditory and visual naming, respectively (**eTable 1**). This study replicated our previous findings from 125 patients [10], showing that older age was associated with faster median response times, whereas left-handedness and the presence of a left-hemispheric epileptogenic focus were linked to reduced left-hemispheric dominance of high-gamma augmentation (**eResult**).

### Overview of network dynamics

Given the extensive spatiotemporal information on neural interactions provided in this study, we recommend first examining the videos for a comprehensive overview of the whole-brain findings. Neural information flows during auditory naming and visual naming tasks are presented with functional coactivation/co-deactivation (**eVideos 1–2**) and cortical high-gamma modulation (**eVideos 3–4**). In general, excitatory flow initially propagated from modality-specific sensory cortices to higher-order regions, later becoming bidirectional and coinciding with sustained functional coactivation. Excitatory flows were transient, lasting up to 500 ms and followed by inhibitory flows; in contrast, functional coactivation persisted and slowly subsided.

During auditory naming, bidirectional excitatory flows were most prominent along the left arcuate fasciculus, whereas during visual naming, they were distributed more evenly across multiple tracts, including the bilateral inferior longitudinal, left arcuate, and left inferior fronto-occipital fasciculi. Additional fasciculi also exhibited excitatory flows concurrently. **eFigures 7–10** depict the proportion of each fasciculus demonstrating excitatory and inhibitory flows across time bins, and **eFigures 11–14** depict the proportion exhibiting functional coactivation and co-deactivation. These **eFigures** assist readers in appreciating that excitatory flows are transient, whereas functional coactivation persists over a longer period.

### Neural information flows during auditory naming

Following stimulus onset, excitatory flow initially propagated from the pSTG to adjacent gyri before becoming bidirectional and coinciding with sustained functional coactivation (**eVideo 1; Figure 1A**). For example, unidirectional intrahemispheric excitatory flows emerged from the left pSTG to the pMTG via U-fibers, peaking at 45 ms after stimulus onset (**Figure 2A**). Subsequently, bidirectional excitatory flows between the left pMTG and pSTG were observed around 200 ms after stimulus onset. Concurrently, bidirectional excitatory flows were observed between the right and left pSTG via callosal fibers.

**Figure 1.**
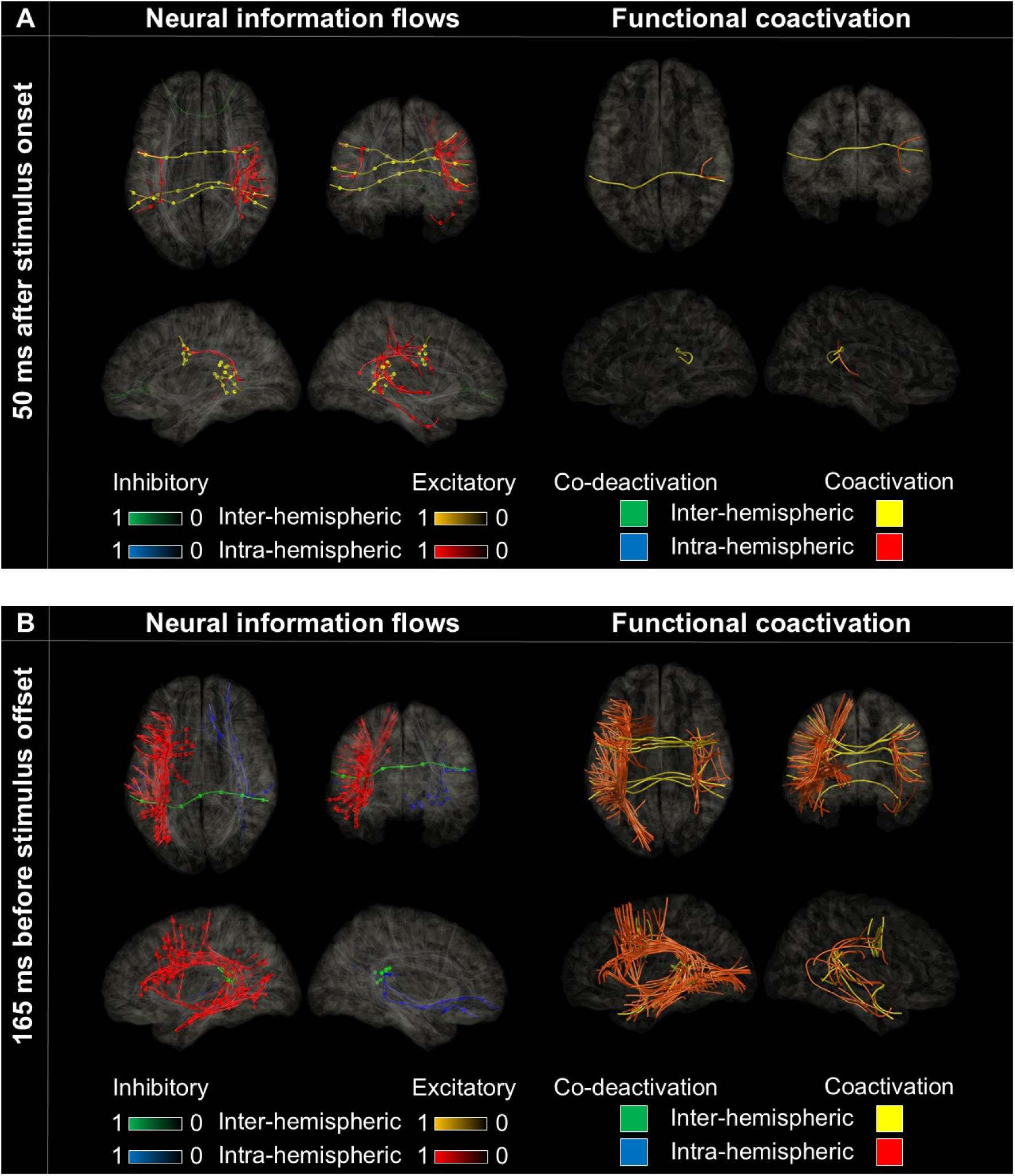
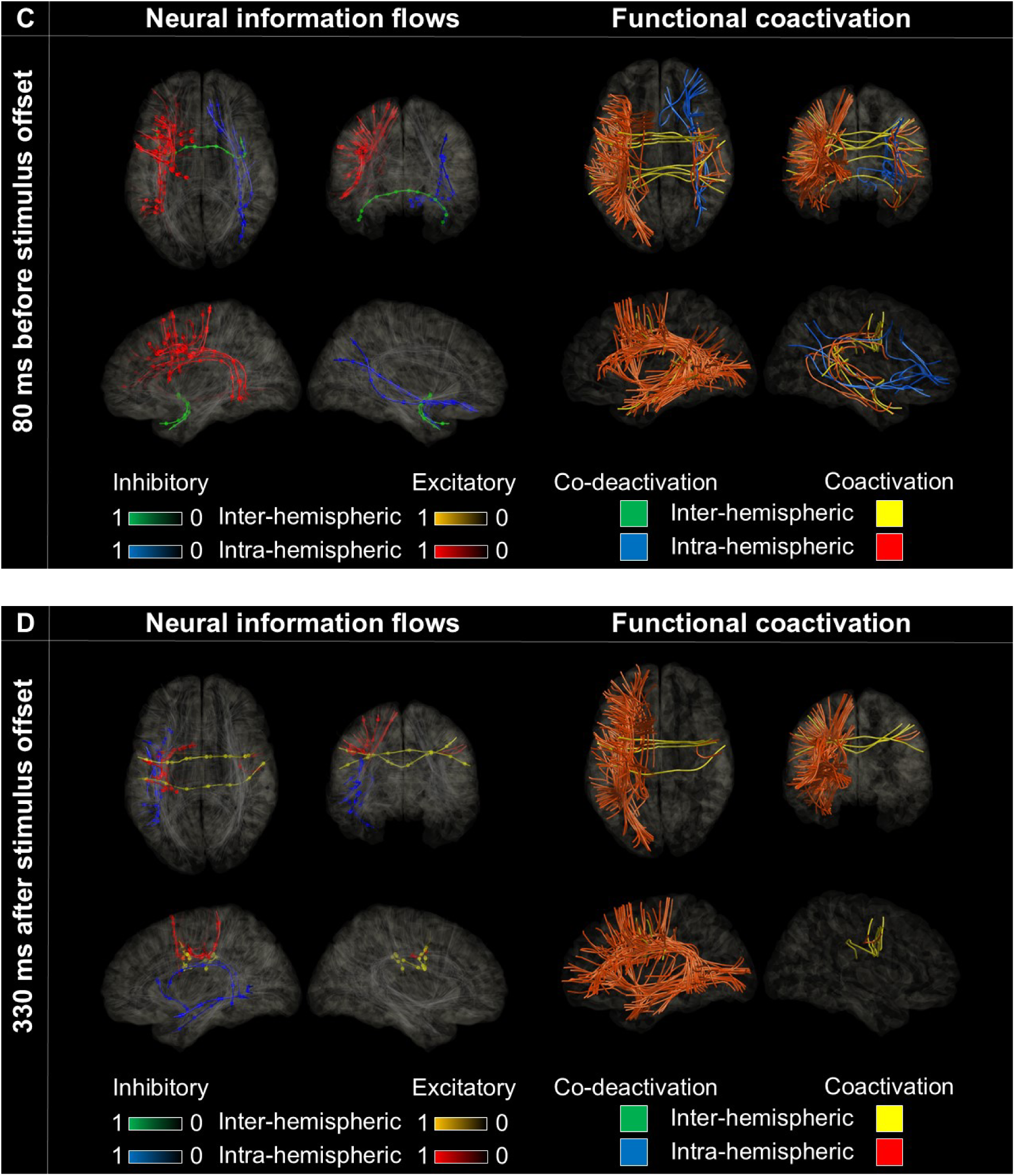
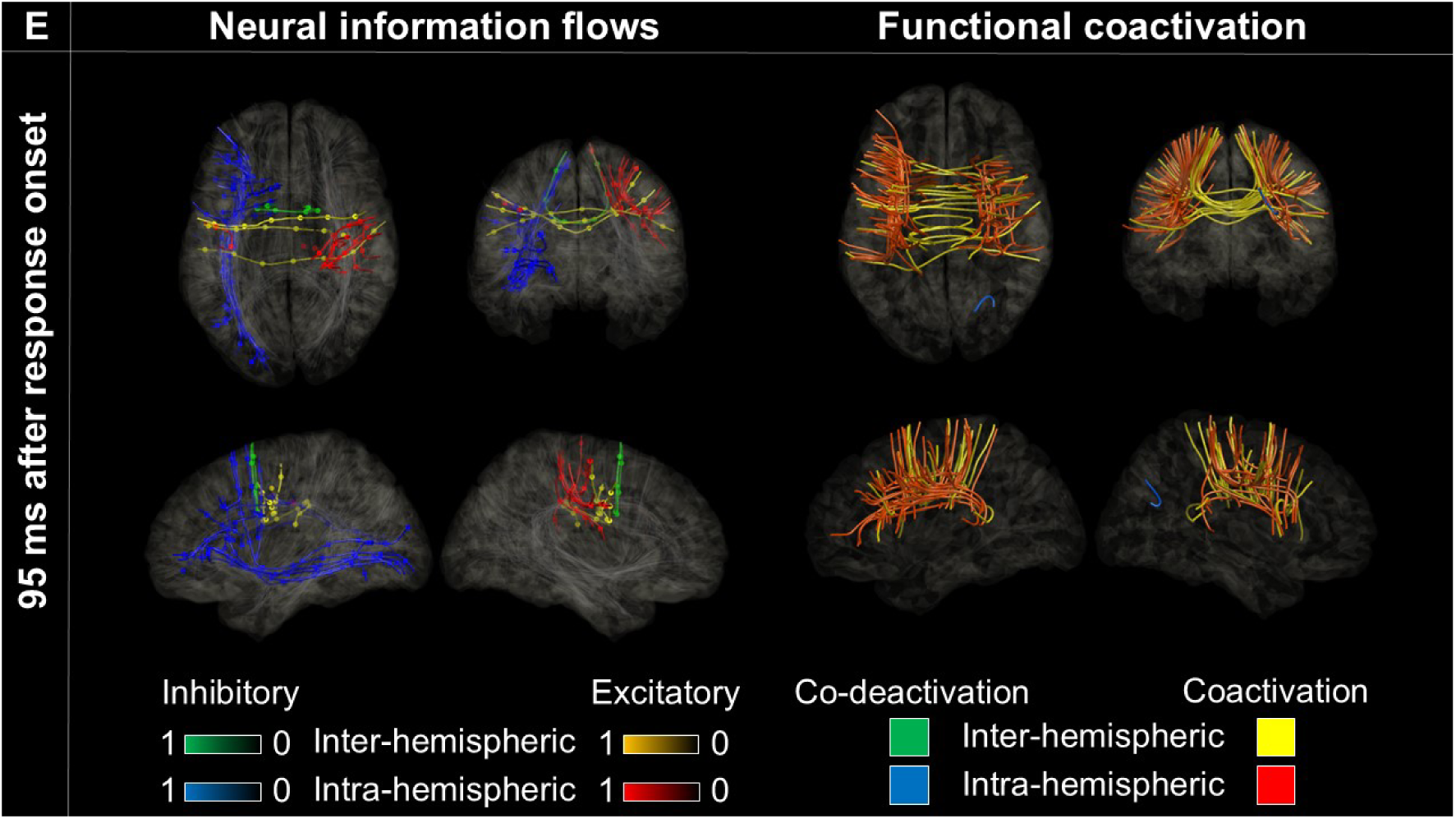
Neural information flows and functional coactivation/co-deactivation during auditory naming. Left: Excitatory (red/yellow) and inhibitory (blue/green) flows through white matter tracts. Right: Functional coactivation (red/yellow) and co-deactivation (blue/green). **A** 50 ms after stimulus onset, efferent excitatory flow intensity at a given ROI correlated with probability of stimulation-induced auditory hallucinations. **B** 165 ms before stimulus offset, both efferent and afferent excitatory flows correlated with receptive aphasia. **C** 80 ms before stimulus offset, both efferent and afferent excitatory flows correlated with expressive aphasia. **D** 330 ms after stimulus offset, both efferent and afferent excitatory flows correlated with speech arrest. **E** 95 ms after response onset, both efferent and afferent excitatory flows correlated with face sensorimotor symptoms.

Peaking at 55 ms after stimulus onset, unidirectional intrahemispheric excitatory flows likewise emerged from the pSTG to the inferior precentral gyrus via the left arcuate fasciculus, (**Figure 2B**). Subsequently, excitatory flows from the left inferior precentral gyrus to the pSTG were observed around 180 ms after stimulus onset.

**Figure 2.**
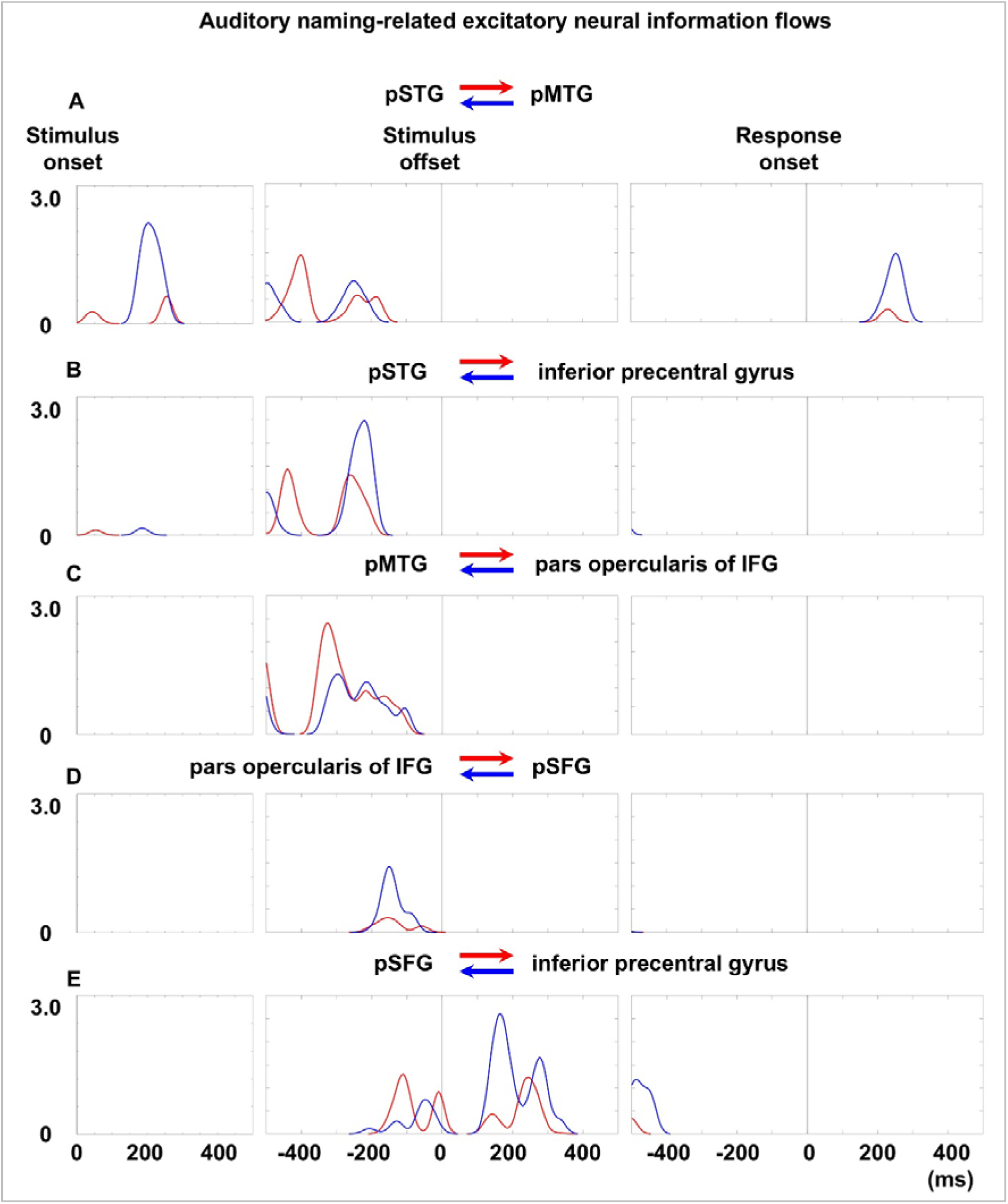
Auditory naming–related excitatory neural information flows within the left hemisphere. X-axis: time; Y-axis: flow intensity (transfer entropy). Plot colors denote the direction of excitatory flows between the following regions of interest. **A** posterior superior temporal gyrus (pSTG) and posterior middle temporal gyrus (pMTG). **B** pSTG and inferior precentral gyrus. **C** pMTG and pars opercularis of the inferior frontal gyrus (IFG). **D** pars opercularis of the IFG and posterior superior frontal gyrus (pSFG). **E** pSFG and inferior precentral gyrus.

Peaking 100–300 ms before stimulus offset, bidirectional excitatory flows were evident between the lateral temporal and perisylvian frontal regions via the left arcuate fasciculus (**Figures 1B–1C**). For example, **Figure 2C** illustrates the dynamics of bidirectional flows between the left pMTG and the pars opercularis of the inferior frontal gyrus (IFG) during this period.

Around stimulus offset, bidirectional excitatory flows through the left arcuate fasciculus subsided (**eVideo 1**), whereas bidirectional excitatory flows across the left frontal lobe became prominent via the frontal aslant fasciculus and U-fibers. For example, **Figure 2D** shows the dynamics of bidirectional excitatory flows between the pars opercularis of the left IFG and the posterior superior frontal gyrus (pSFG). Peaking 200–300 ms after stimulus offset, bidirectional excitatory flows became evident between the left pSFG and the inferior precentral gyrus (**Figure 2E**). As time elapsed after stimulus offset, bidirectional inhibitory information flows through the left arcuate fasciculus became increasingly prominent, while the extent of functional coactivation along this pathway gradually diminished (**Figure 1D**).

Around response onset, bidirectional excitatory flows were confined to intra- and interhemispheric pathways involving the Rolandic cortices and pSTG (**Figure 1E**).

### Auditory naming-related neural information flows and stimulation-induced manifestations

**eFigure 15** shows probability maps of stimulation-induced symptoms, while **Figure 3** illustrates how correlations between efferent excitatory flow intensity and symptom probability evolve over time. Peak correlations occurred at different time points:

- Auditory hallucination: 5 ms post-stimulus onset (rho: +0.58; p-value: 4.7×10^-7^)
- Receptive aphasia: 165 ms pre-stimulus offset (rho: +0.60; p-value: 1.5×10^-7^)
- Expressive aphasia: 80 ms pre-stimulus offset (rho: +0.81; p-value: 2.1×10^-16^)
- Speech arrest: 330 ms post-stimulus offset (rho: +0.69; p-value: 1.4×10^-10^)
- Facial sensorimotor symptom: 95 ms post-response onset (rho: +0.60; p-value: 1.1×10^-7^).

**Figure 3.**
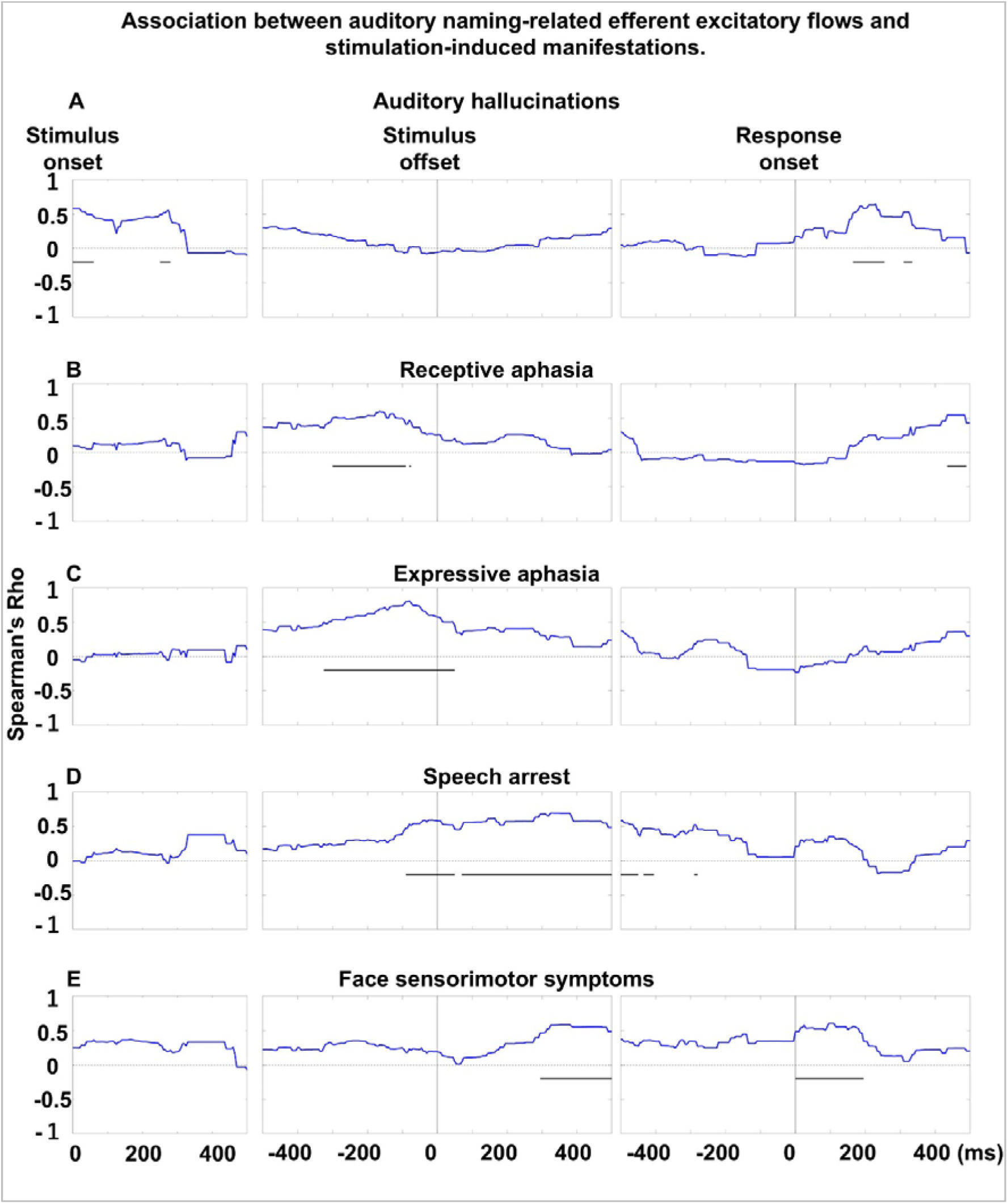
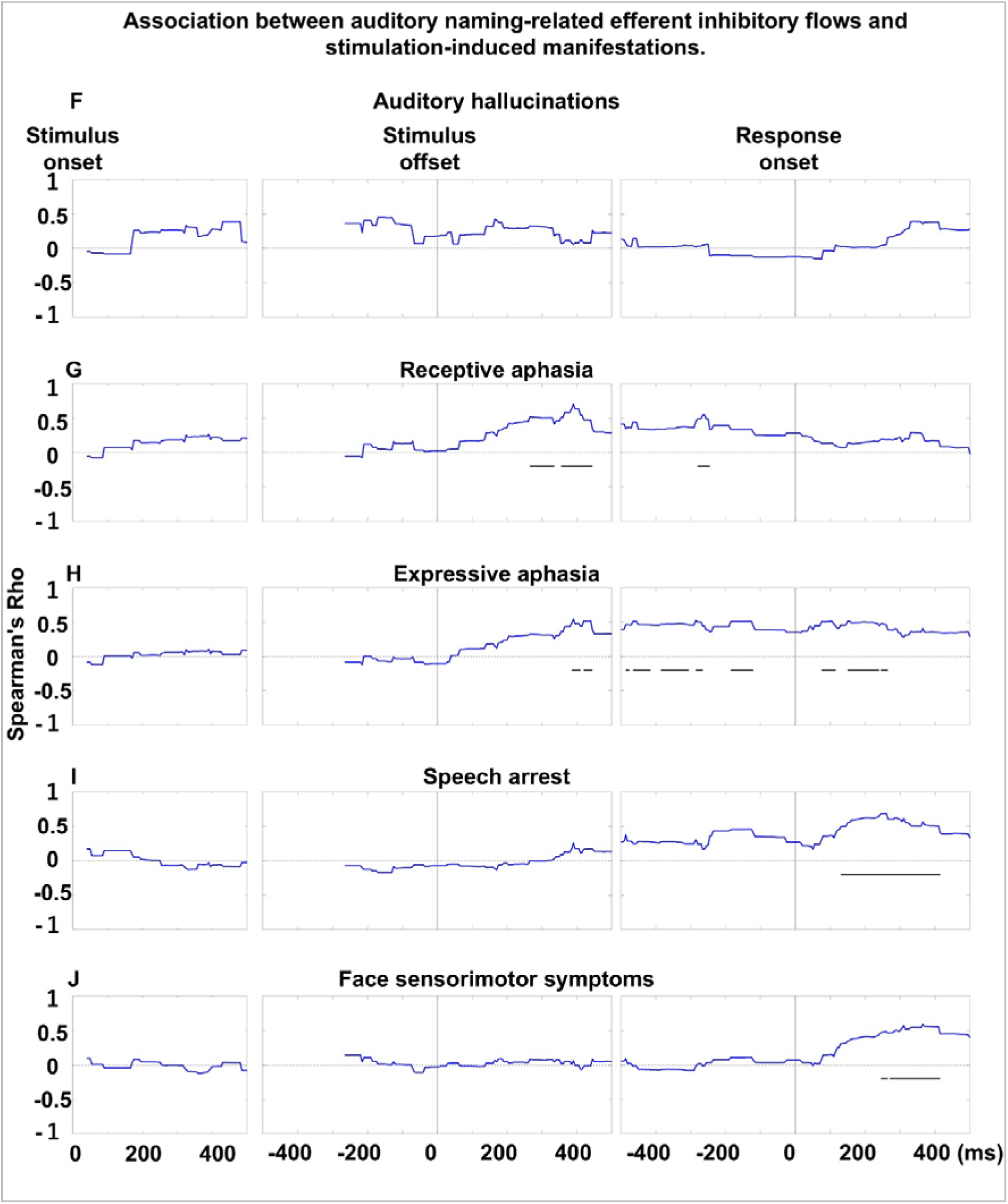
Association between auditory naming–related excitatory and inhibitory efferent flows and stimulation-induced manifestations. Each plot shows Spearman’s rho, reflecting the correlation between mean flow intensity and the likelihood of stimulation-induced symptoms across time bins. **A-E** Correlations between efferent excitatory flows and stimulation-induced manifestations. Rho values > 0 indicate that, at a given moment, the probability of a specific manifestation was higher in regions emitting excitatory information flows to other regions via white matter tracts, as determined by transfer entropy-based effective connectivity analysis. Horizontal bars mark intervals of significant correlation. **F-J** Correlations between efferent inhibitory flows and stimulation-induced manifestations. Rho values >0 indicate that the probability of a given manifestation was higher in regions emitting inhibitory flows to other regions. Spearman’s rho was not computed for time bins lacking neural information flows.

The temporal correlation patterns of afferent and efferent excitatory flows with symptom probability were similar (**eFigure 16**).

Additionally, the intensity of efferent inhibitory flow was correlated with symptom probability as follows:

- Auditory hallucination: no significant correlation
- Receptive aphasia: 390 ms post-stimulus offset (rho: +0.70; p-value: 5.4×10^-11^)
- Expressive aphasia: 390 ms post-stimulus offset (rho: +0.55; p-value: 2.3×10^-6^)
- Speech arrest: 260 ms post-response onset (rho: +0.69; p-value: 2.2 ×10^-10^)
- Facial sensorimotor symptom: 365 ms post-response onset (rho: +0.59; p-value: 2.3×10^-7^).

### Neural correlates of variability in response times in auditory naming

**eFigure 18** presents response time–sorted neural information flow across the 66 ROIs, whereas **Figure 4** illustrates the timing and location of significant correlations between neural information flow intensity and response time. Around 400 ms after stimulus onset, stronger afferent inhibitory flow to the right pSTG was associated with slower responses (maximum rho: +1.00; p-value: 0.003; **Figure 4A**). Around 250 ms before stimulus offset, stronger efferent and afferent excitatory flows at the left pMTG and inferior precentral gyrus were associated with faster responses (minimum rho: –0.94; p-value: 0.016; **Figure 4B**); these cortical regions are connected through the arcuate fasciculus.

**Figure 4.**
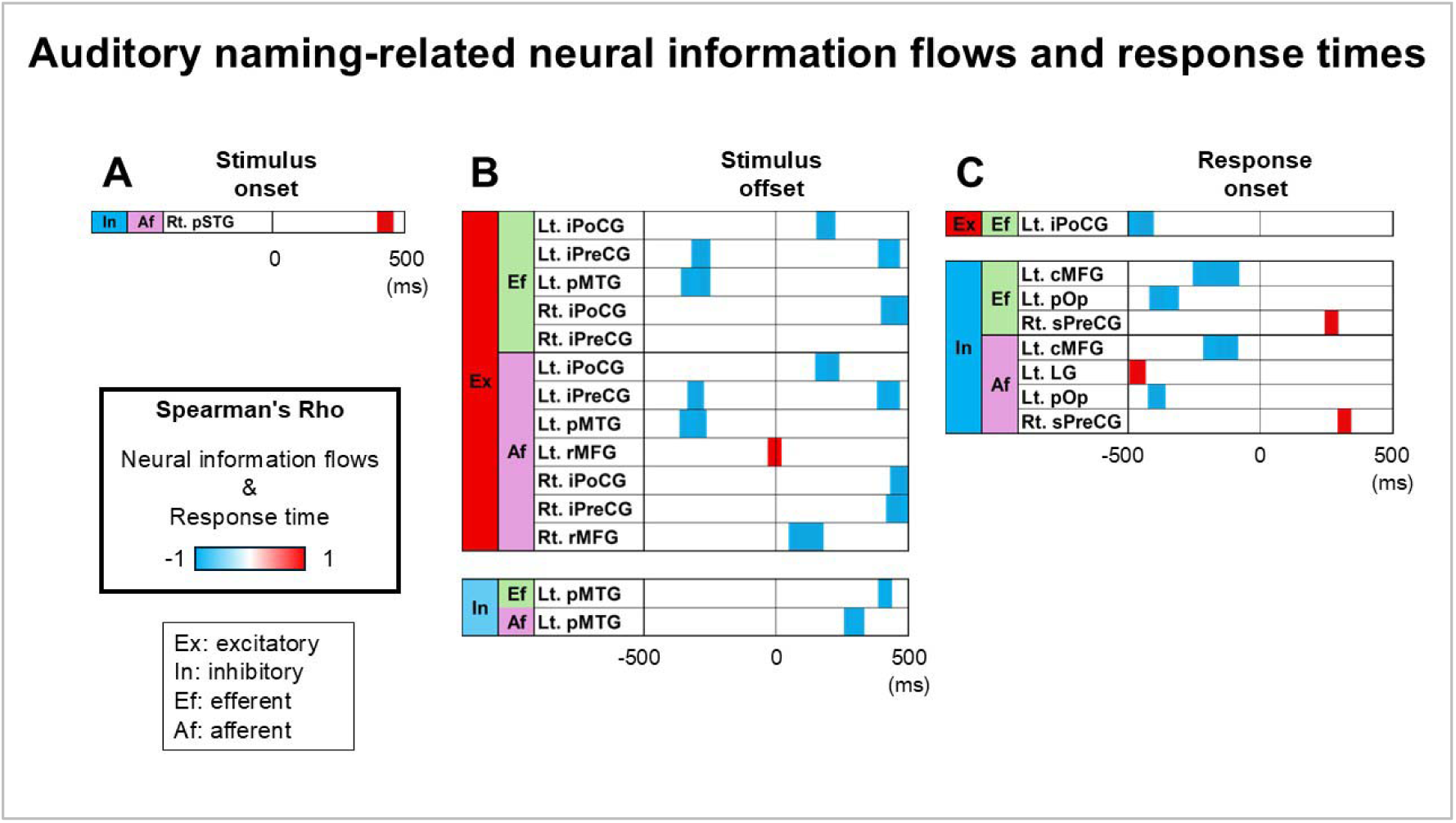
Auditory naming-related neural information flows and response times. Spearman’s rho over time illustrating correlations between neural information flow intensity and naming response times. Blue indicates increased flows associated with faster responses; red indicates slower responses. Only regions of interest (ROIs) with significant correlations are shown. Significance was defined as p-value < 0.05 sustained across a 50-ms period. **A** following stimulus onset. **B** around stimulus offset. **C** around response onset. Full names of all regions of interest are provided in **eTable 2**.

### Neural information flows during visual naming

After stimulus onset, excitatory flow propagated from the lateral occipital gyrus to the posterior fusiform gyrus in each hemisphere before becoming bidirectional and coinciding with sustained functional coactivation (**eVideo 2; Figures 5A-5B**). For instance, **Figure 6A** illustrates unidirectional intra-hemispheric excitatory flows emerging from the left lateral occipital to posterior fusiform gyrus via the inferior longitudinal fasciculus, peaking at 65 ms post-stimulus onset. Subsequently, excitatory flows from the left posterior fusiform to lateral occipital gyrus appeared, with peak latency of 400 ms (**Figure 6A**). Concurrently, bidirectional excitatory flows were observed between these posterior cortical regions via callosal fibers.

**Figure 5.**
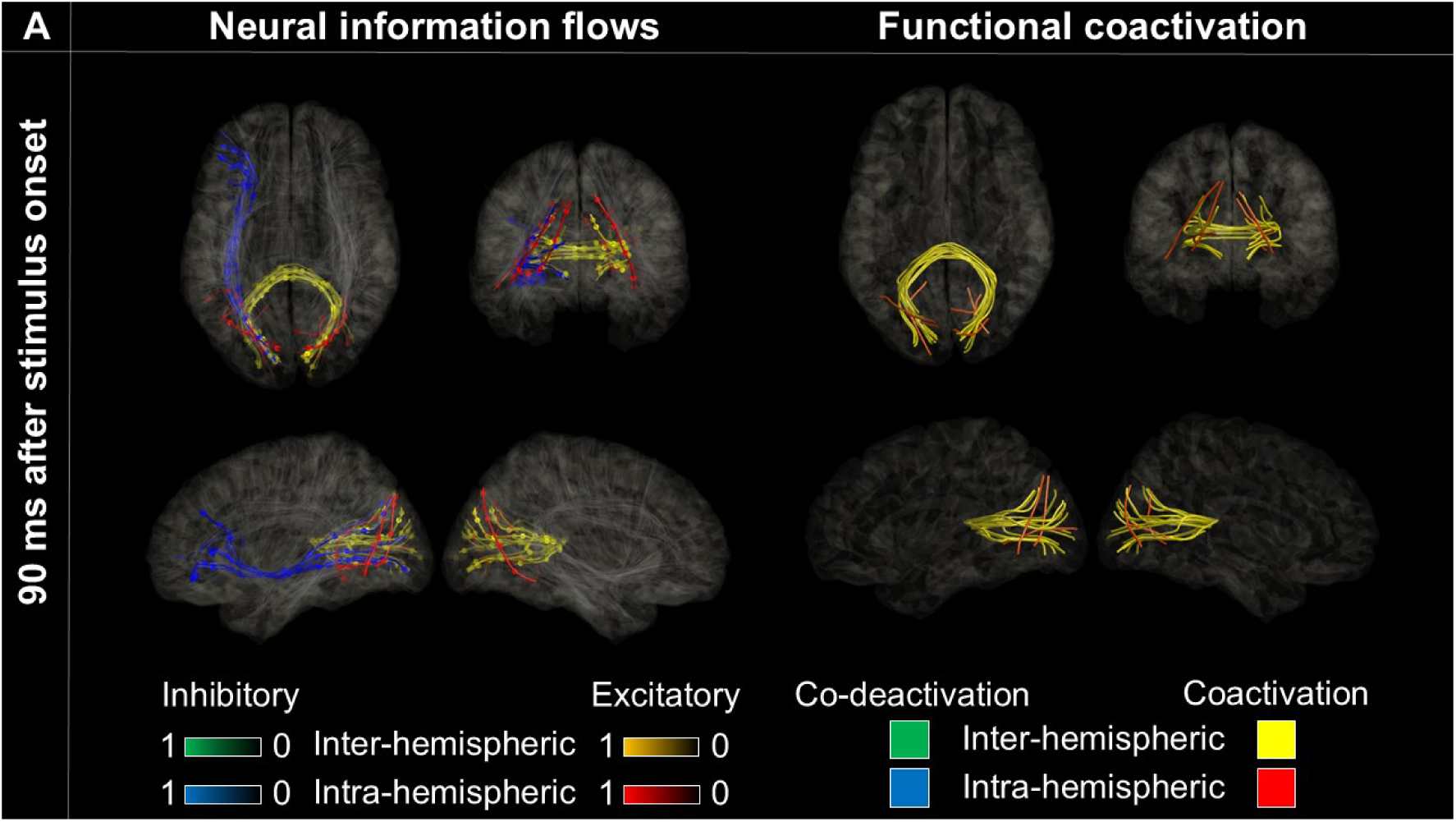
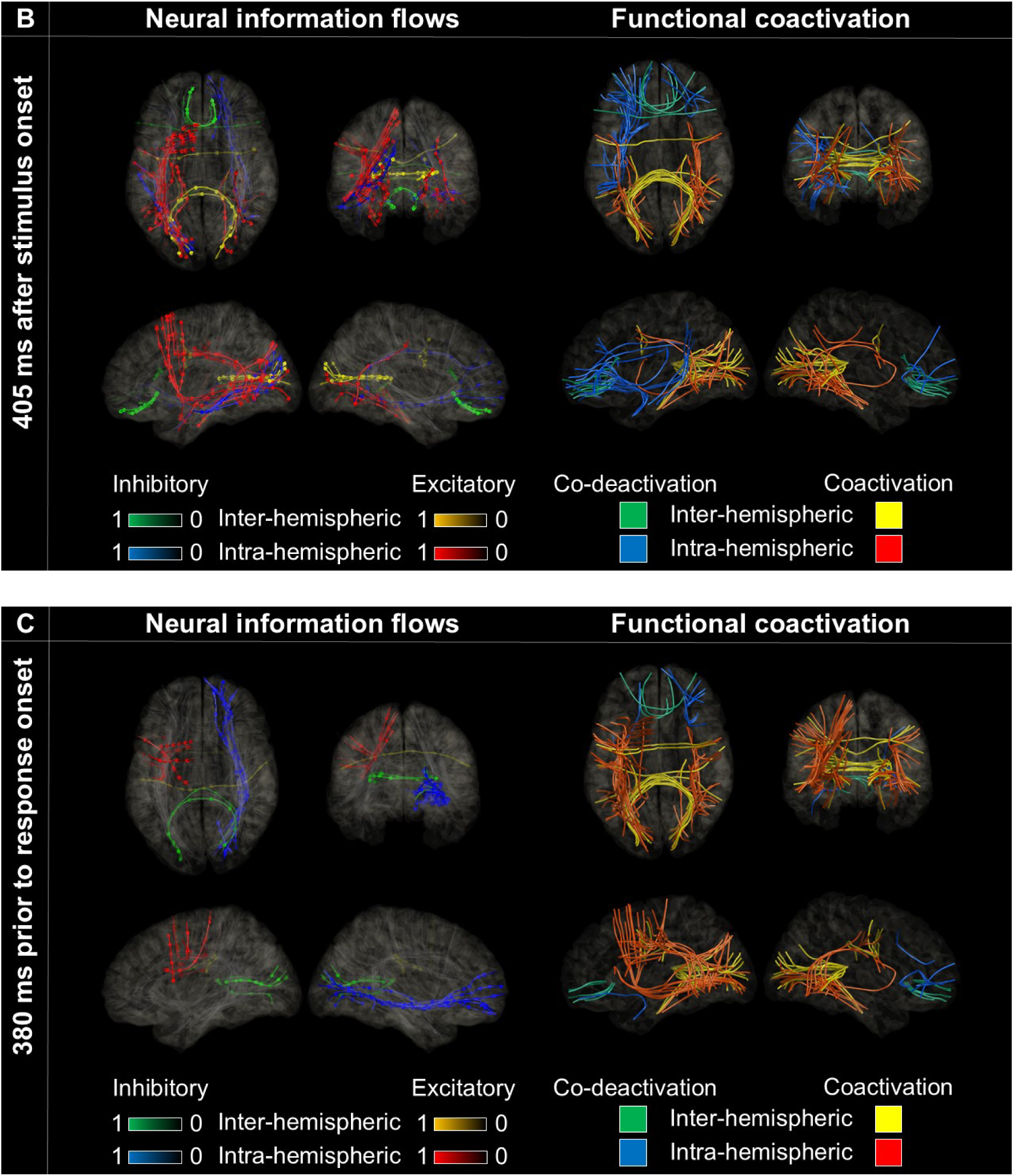
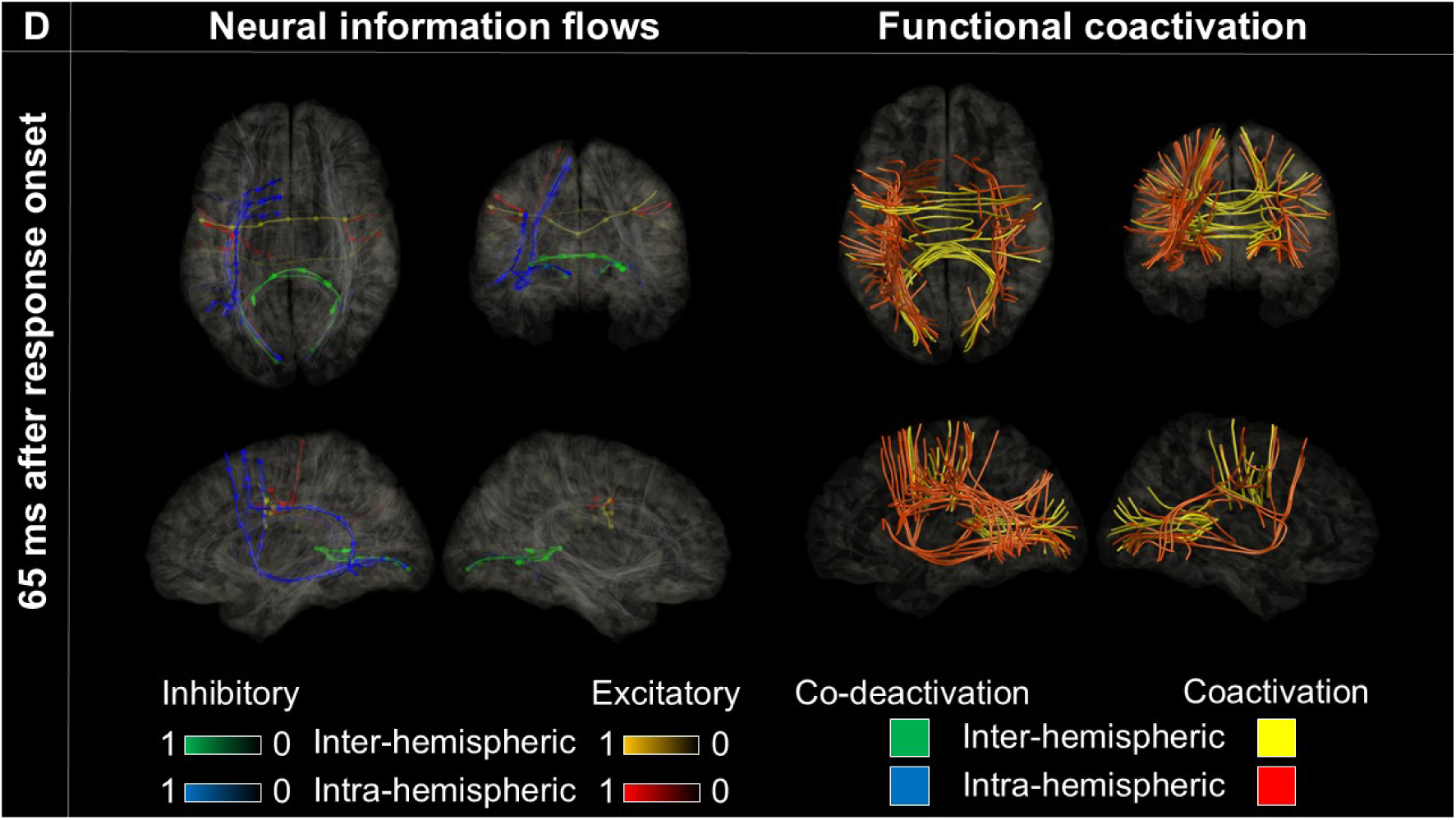
Neural information flows and functional coactivation/co-deactivation during visual naming. **A** 90 ms after stimulus onset, both efferent and afferent excitatory flow intensity correlated with stimulation-induced phosphene and visual distortion. **B** 405 ms after stimulus onset, both efferent and afferent excitatory flow intensity correlated with stimulation-induced visual distortion. **C** 380 ms prior to response onset, both efferent and afferent excitatory flow intensity correlated with speech arrest. **D** 65 ms after response onset, both efferent and afferent excitatory flow intensity correlated with face sensorimotor symptoms.

Between 350 and 450 ms after stimulus onset, bidirectional excitatory flows also took place between the lateral occipital and posterior inferior temporal gyrus (pITG), between the left lateral occipital gyrus and inferior precentral gyrus via the superior longitudinal fasciculus, and between the left pITG and inferior precentral gyrus via the arcuate fasciculus (**Figures 5B, 6B, and 6C**).

Between 350 and 450 ms prior to response onset, bidirectional excitatory flows were observed across the inferior precentral gyrus, pars opercularis of the IFG, and pSFG in the left hemisphere (**Figures 5C, 6D, and 6E**).

During responses, intra- and inter-hemispheric excitatory flows were confined to the Rolandic areas, whereas functional coactivation continued to encompass posterior head regions (**Figure 5D**).

### Visual naming-related neural information flows and stimulation-induced manifestations

Correlations between efferent excitatory flow and stimulation-induced manifestations peaked at distinct times (**Figure 7**):

- Phosphenes at 90 ms post-stimulus (rho: +0.75; p-value: 1.8×10D¹³)
- Visual distortions at 405 ms post-stimulus (rho: +0.63; p-value: 1.1×10D^8^)
- Visual naming error: no significant correlation
- Speech arrest: 380 ms pre-response onset (rho: +0.56; p-value: 1.0×10^-6^)
- Facial sensorimotor: 65 ms post-response onset (rho: +0.54; p-value: 2.0×10^-6^).

Afferent and efferent excitatory flow intensities demonstrated similar patterns of correlation with symptom probability (**eFigure 17**).

Correlations between efferent inhibitory flow and stimulation-induced manifestations peaked as follows (**Figure 7**):

- Phosphenes at 240 ms pre-response onset (rho: +0.68; p-value: 2.3×10D^10^)
- Visual distortions at 80 ms pre-response onset (rho: +0.70; p-value: 3.8×10D^11^)
- No correlation with visual naming error, speech arrest, or facial sensorimotor symptoms.

### Neural correlates of within-individual variability in response times in visual naming

**eFigure 19** presents response time–sorted neural information flow at given ROIs. During a period between 200 and 500 ms following stimulus onset, stronger efferent or afferent excitatory flows in the basal posterior temporal regions, including the bilateral pITG and the right parahippocampal gyrus were associated with faster responses (minimum rho: –1.0; p-value: 0.003; **Figure 8A**).

## Discussion

### Summary

This study provides the causal evidence that transient neural information flows through white matter pathways—captured with millisecond-level resolution—underlie distinct stages of speech processing during auditory and visual naming. By correlating the strength and timing of excitatory and inhibitory flows with stimulation-induced symptoms, we identified dynamic, directionally specific interactions that unfold along structurally constrained circuits. Excitatory flows first propagate from sensory cortices to higher-order regions, become bidirectional with recurrent coactivation, and are later followed by inhibitory flows. Each excitatory event is brief (<500 ms) and task-specific. During auditory naming, faster responses were associated with stronger excitatory flows across left perisylvian regions; during visual naming, faster responses aligned with stronger flows in bilateral posterior basal temporal cortices. Excitatory flows at key time points predicted stimulation-induced speech symptoms, while inhibitory flows showed later, distinct symptom associations. These findings suggest that rapid, transient accelerations of neural activity through white-matter pathways support sequential stages of speech—from perceptual encoding to lexical retrieval and motor planning—and are followed by inhibitory deceleration phases that likely mediate processing reallocation. Dynamic flow trajectories, their structural substrates, and corresponding clinical correlations are visualized in **eVideos 1–4** and expanded upon in **eFigures 7–17**.

### Neural information flows underlying rapid auditory naming

The present study revealed a spatiotemporal sequence of excitatory flows consistent with the dual-stream model of speech processing [1–4], which proposes reciprocal interactions among auditory, temporal, and frontal cortical regions. Peaking at 45 ms after auditory stimulus onset, unidirectional excitatory flows emerged from the left pSTG to the pMTG via U-fibers (**Figure 2A**). By 55 ms, additional unidirectional excitatory flows arose from the left pSTG to the inferior precentral gyrus through the arcuate fasciculus (**Figure 2B**). Cortical regions sending stronger excitatory flows were associated with a higher probability of stimulation-induced auditory hallucinations (**Figure 3A**). The initial pSTG-originating flows likely reflect feed-forward transmissions from regions mediating early acoustic analysis to two parallel cortical streams: (i) ventral-stream lateral temporal regions that perform phonetic-to-semantic transformations, linking acoustic–phonological representations to word-level meaning [21,42–43]; and (ii) dorsal-stream frontal regions that support auditory–motor mapping, transforming perceptual features of speech into articulatory motor commands via the arcuate fasciculus [44–45]. Approximately 200 ms after stimulus onset, these feed-forward flows were followed by reverse-direction excitatory flows toward the pSTG. Such feedback is likely generated by perisylvian structures, including the pMTG and inferior precentral gyrus, which send predictive signals and corollary discharges back to auditory cortex to refine the perceptual representation of incoming speech input [46–49]. Given their early timing, these recurrent interactions probably operate at a sub-lexical phonological level, dynamically optimizing speech perception.

Variations in neural information flows during auditory stimulus presentation may underlie within-individual fluctuations in response times. An increase in afferent inhibitory flow to the right pSTG around 400 ms after stimulus onset was associated with slower auditory naming responses, whereas faster responses were linked to reduced inhibitory inflow to this region (**Figure 4A**). Consistently, our previous study showed that faster responses coincided with greater high-gamma augmentation in the right pSTG during the same time window [10]. Taken together, these findings indicate that delayed responses may result from diminished attention to auditory stimuli, leading to reduced neural gain and attenuated processing of auditory inputs [50–52].

We also found that acceleration and sustained engagement of left perisylvian interactions facilitated the transformation of semantic representations into articulatory plans for naming. Specifically, stronger excitatory flows involving the left pMTG and inferior precentral gyrus approximately 250 ms before stimulus offset were associated with faster auditory naming responses (**Figure 4B**). This temporal window likely reflects the beginning of a convergence phase in which ventral-stream regions supporting semantic processing interact with dorsal frontal regions responsible for lexical retrieval and motor preparation. Moreover, cortical regions exhibiting stronger excitatory flows showed a higher probability of stimulation-induced receptive and expressive aphasia (**Figures 3B–3C**), suggesting that these excitatory interactions are necessary for both lexical access and fluent speech production. Consistently, our previous study demonstrated that stronger coactivation across the left perisylvian network—including the pMTG, pSTG, IFG, and inferior precentral gyrus—during the 500-ms period centered on the stimulus offset was associated with faster auditory naming responses [10]. These temporal and frontal regions are directly interconnected via the arcuate fasciculus, which mediates bidirectional information exchange between semantic-lexical and motor systems [37,53–54]; damage to this tract produces severe naming deficits [7]. Together, these findings provide converging evidence for posterior temporal–inferior frontal (i.e., Wernicke–Broca) coordination via a direct dorsal stream pathway supporting auditory naming, addressing an outstanding question highlighted by Dick and Tremblay [2].

Our observations directly support the model proposed by Binder [16], which holds that the left pMTG functions as a central hub that constructs word meaning through reciprocal exchanges with adjacent temporal lobe cortices. The pMTG synthesizes phonological and syntactic inputs into unified semantic representations by dynamically coordinating activity across distributed perisylvian regions. Our findings refine this model by specifying the precise timing of accelerated bidirectional interactions involving the left pMTG (**Figure 2**) and by demonstrating that its coupling with frontal perisylvian regions is reciprocal—mediated predominantly through the arcuate fasciculus—rather than unidirectional from temporal to frontal areas. This bidirectional architecture may reflect continuous feedback between semantic integration and response selection processes that supports real-time updating of lexical meaning during speech comprehension [55–56].

We also found that inhibitory flows emerging after excitatory flows form part of a temporally coordinated physiological mechanism that reallocates neural resources across successive stages of speech processing. Before stimulus offset, excitatory flows propagated along the left arcuate fasciculus and were associated with stimulation-induced receptive and expressive aphasia (**Figures 3B–3C**), implicating this tract in lexical-semantic retrieval. Following stimulus offset, inhibitory flows emerged along the same tract, while excitatory flows arose via U-fibers linking left frontal and Rolandic regions—corresponding with stimulation-induced speech arrest (**Figure 3D**). This sequential excitatory–inhibitory transition likely reflects a transient gating mechanism, suppressing earlier semantic processes to enable the recruitment of motor circuits for speech initiation. Consistent with this interpretation, prior iEEG studies have reported reciprocal activation patterns between task-positive regions and the default mode network [57–59]. Our findings uniquely demonstrate that excitatory and inhibitory flows operate along the same eloquent pathways in a temporally ordered manner, reflecting an intrinsic acceleration–deceleration cycle of information transfer. This dynamic sequencing aligns with the multiple-network model [6,60], proposes that sentence processing and speech production emerge from coordinated transitions across distributed functional systems, spanning conceptual access to articulatory planning.

### Neural information flows underlying rapid visual naming

Visual naming–related information flows likewise engaged both ventral and dorsal streams, with ventral engagement preceding dorsal involvement. Peaking at 65 ms after picture stimulus onset, unidirectional excitatory flows emerged from the left lateral occipital gyrus to the posterior fusiform gyrus via the inferior longitudinal fasciculus (**Figure 6A**). Cortical regions exhibiting stronger efferent excitatory flows during this early window were associated with a higher probability of stimulation-induced phosphene and visual distortion (**Figures 7A–7B**). These initial flows from the left lateral occipital gyrus likely reflect feed-forward transmissions from cortical regions mediating lower-order visual feature extraction to ventral-stream posterior temporal regions engaged in hierarchical object analysis, including higher-order visual processing and object recognition [15,42,61]. Around 400 ms after stimulus onset, bidirectional excitatory flows emerged along both ventral and dorsal streams, marking the transition from visual object recognition to lexical–semantic processing and early articulatory planning. In the ventral stream, these flows propagated bilaterally through the inferior longitudinal fasciculus between the lateral occipital gyrus and both the posterior fusiform gyrus and pITG (**Figures 6A–6B**). Greater efferent flows from these basal posterior temporal regions between 200–500 ms post-stimulus onset were associated with faster visual naming responses (**Figure 8A**), consistent with their role in transforming visual object representations into lexical–semantic codes. Concurrently, within the dorsal stream, bidirectional excitatory flows appeared through the left superior longitudinal fasciculus, linking the <u>lateral</u> occipital gyrus with the inferior precentral gyrus and pSFG (**Figure 5B**). In parallel, excitatory flows emerged through the arcuate fasciculus between the left pITG and inferior precentral gyrus (**Figure 6C**), notably bypassing the left pSTG and pMTG—regions prominently engaged during auditory naming [62]. These findings suggest that visual and auditory naming rely on distinct dorsal stream configurations, thereby addressing an outstanding question raised by Dick and Tremblay [2]. The engagement of these frontal lobe structures likely reflects the initiation of motor preparatory processes, integrating lexical–semantic retrieval with articulatory planning by mapping retrieved lexical representations onto motor command frameworks [44–45]. Cortical regions exhibiting stronger efferent excitatory flows around 400 ms after stimulus onset were associated with a higher probability of stimulation-induced visual distortion (**Figure 7B**), and stimulation of the left fusiform gyrus and pITG frequently elicited picture-naming errors (**eFigure 15**) [4,42]. Studies in healthy individuals further demonstrate that scalp-recorded N400 potentials are modulated by the semantic demands of picture stimuli [63–64].

**Figure 6.**
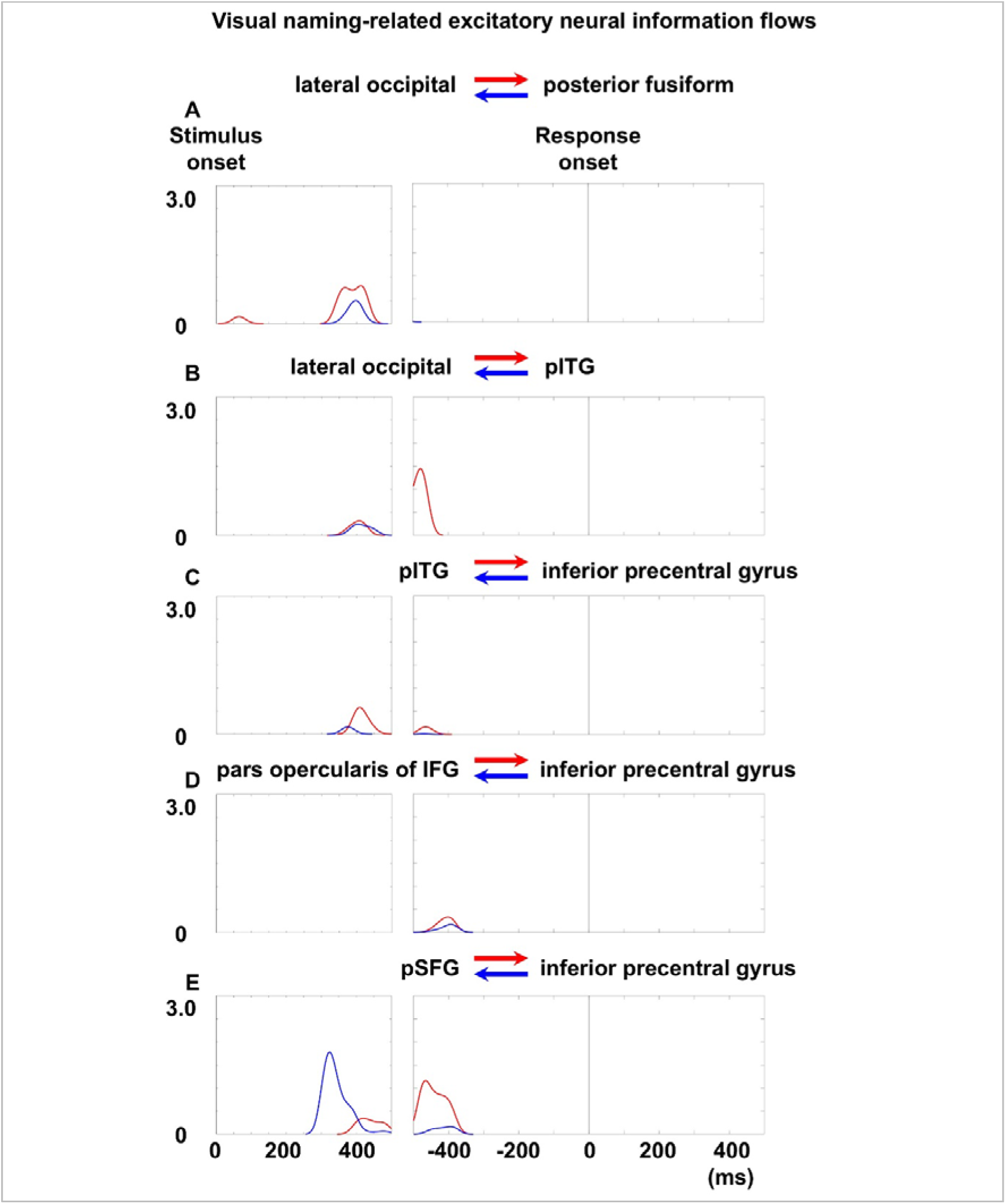
Visual naming–related excitatory neural information flows within the left hemisphere. **A** lateral occipital and posterior fusiform gyri. **B** lateral occipital and posterior inferior temporal gyrus (pITG). **C** pITG and inferior precentral gyrus. **D** pars opercularis of the IFG and inferior precentral gyrus. **E** posterior superior frontal gyrus (pSFG) and inferior precentral gyrus.

During the first 300 ms after picture stimulus onset, we found no evidence of top-down modulation from Broca’s area. Between 50 and 300 ms, bidirectional inhibitory flows emerged across the left IFG, rostral middle frontal gyrus, and <u>medial</u> occipital gyri via frontal U-fibers and the inferior fronto-occipital fasciculus (**Figure 5A**), suggesting transient suppression of frontal cortical activity during early visual analysis. At this stage, neural processing appeared dominated by feed-forward transmissions within posterior object-recognition networks, while lexical retrieval and articulatory planning had yet to be engaged. Consistently, visual naming response times were not correlated with neural information flows involving the left IFG, whereas faster auditory naming responses were associated with greater functional coactivation in the left IFG around stimulus offset [10]. Together, these findings indicate that visual naming initially depends on posterior perceptual mechanisms in the ventral stream, and that activation of Broca’s area and related frontal regions supporting lexical retrieval [65] and motor initiation [11–13] emerges only at later stages.

Approximately 400 ms before responses, excitatory flows shifted toward frontal regions initiating articulation. Bidirectional excitatory interactions emerged across the pars opercularis of the left IFG, pSFG, and inferior precentral gyrus via U-fibers and the frontal aslant fasciculus (**Figures 6D–6E**). At the same time, the pars opercularis of the left IFG remained functionally coactivated with the pITG through the arcuate fasciculus (**Figure 5C**), linking lexical retrieval with motor planning. Excitatory flows during this preparatory phase were associated with stimulation-induced speech arrest (**Figure 7D**), reflecting a transition from lexical to articulatory processing consistent with prior studies [19,65].

**Figure 7.**
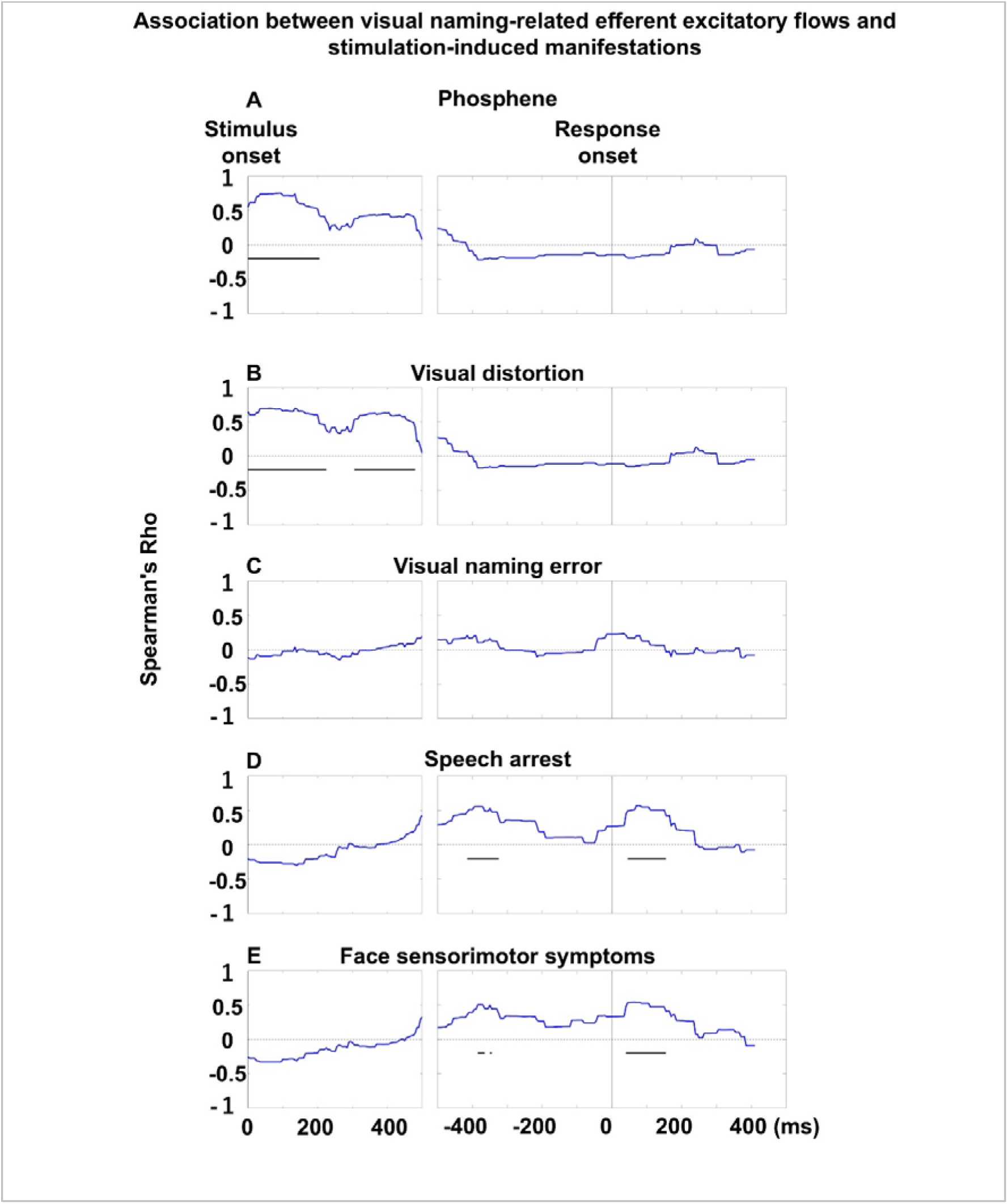
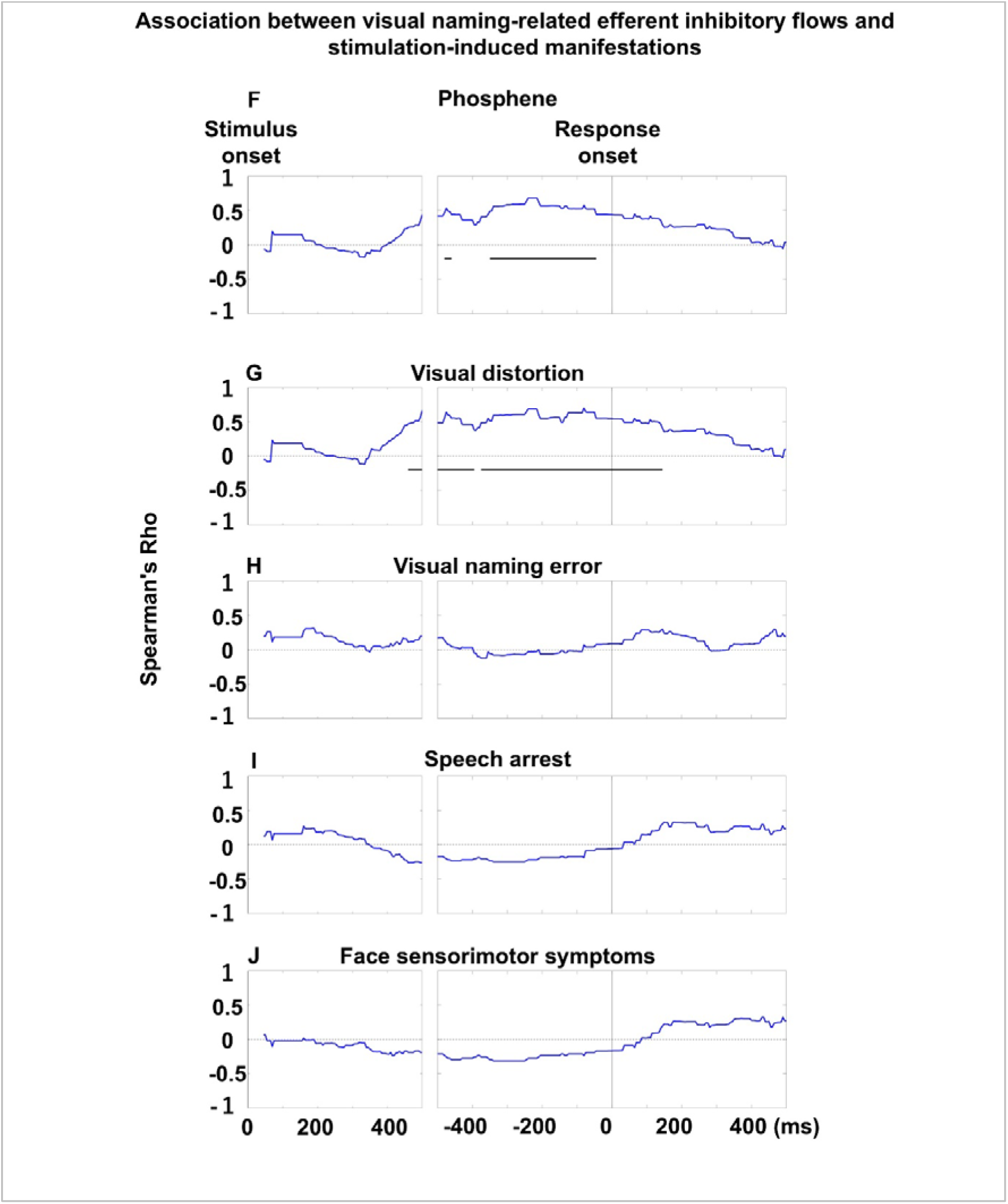
Association between visual naming–related efferent flows and stimulation-induced manifestations. Each plot shows Spearman’s rho, reflecting the correlation between mean flow intensity and the likelihood of stimulation-induced symptoms across time bins. **A–E** Correlations between efferent excitatory flows and stimulation-induced manifestations. Rho values > 0 indicate that, at a given moment, the probability of a given manifestation was higher in regions emitting excitatory flows to other regions via white matter tracts, as determined by transfer entropy–based effective connectivity analysis. Horizontal bars mark intervals of significant correlation. **F-J** Correlations between efferent inhibitory flows and stimulation-induced manifestations. Rho values >0 indicate that the probability of a given manifestation was higher in regions emitting inhibitory flows to other regions. Spearman’s rho was not computed for time bins lacking neural information flows.

**Figure 8.**
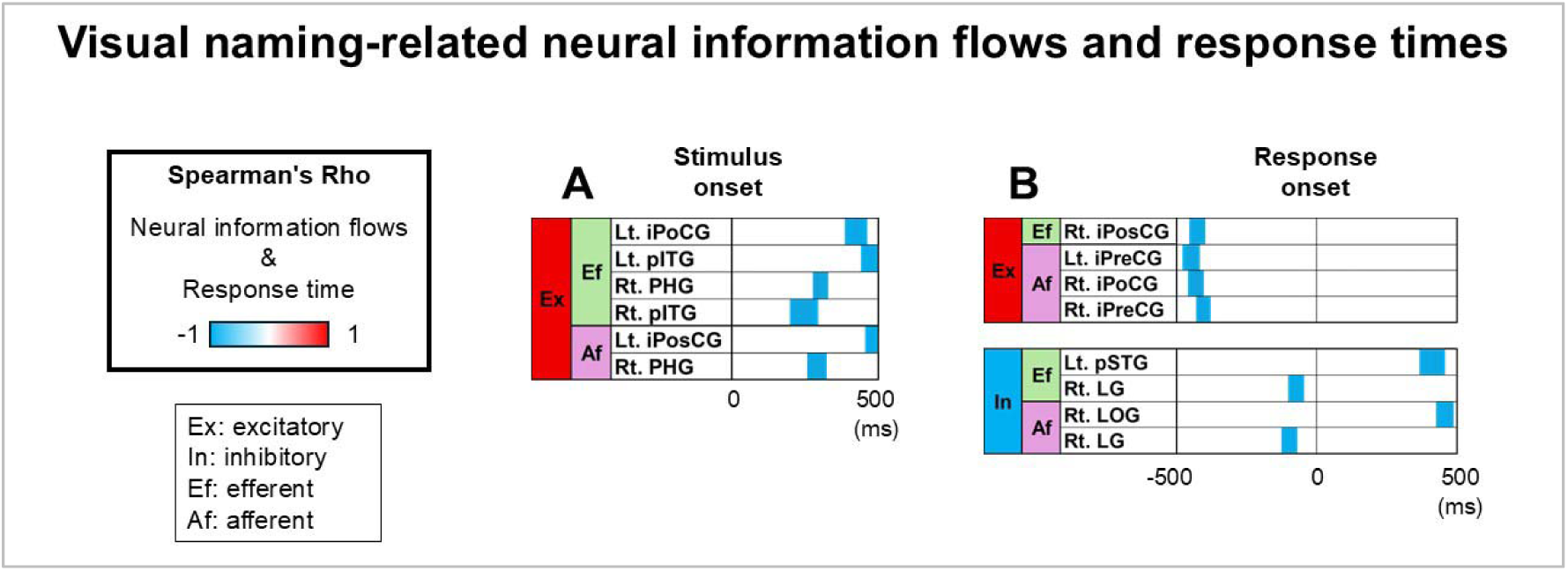
Visual naming-related neural information flows and response times. Spearman’s rho over time illustrating correlations between neural information flow intensity and naming response times. Blue indicates increased flows associated with faster responses; red indicates slower responses. **A** following stimulus onset. **B** around response onset. Full names of all regions of interest are provided in **eTable 2**.

### Utility of the atlases

Our atlases visualize two complementary forms of temporally resolved neural interaction: one maps directional information flow, and the other captures functional coactivation along specific white matter tracts (**eVideos 1-4**). Although validation through postoperative neuropsychological outcomes [31–32] is still needed, both atlases show functional relevance, as supported by converging correlations with direct electrical stimulation results and behavioral response times. During auditory naming, both excitatory flow and coactivation intensity predicted stimulation-induced symptoms with comparable effect sizes (Spearman’s rho ≈ 0.6–0.8) [10]. In visual naming, both measures similarly tracked the likelihood of visual phenomena such as phosphenes and distortions (rho ≈ 0.6–0.9). However, a mechanistic dissociation emerged: only excitatory flows (rho ≈ 0.55; **Figures 7D–7E**) predicted speech arrest and face sensorimotor symptoms, whereas coactivation during visual naming did not. Persistent coactivation near visual naming response onset may obscure correlations (**Figures 5C–5D**), while transient excitatory flows precisely reflect initiation of motor processes. Within individuals, brief surges in both excitatory flow and coactivation around auditory stimulus offset and shortly after picture onset were associated with faster naming responses. Notably, fewer brain regions showed significant associations between excitatory flow and response times—likely reflecting the phasic, time-limited nature of these flows compared to the more sustained synchrony measured by coactivation [10]. Together, these findings underscore the complementary mechanistic value of both atlases: excitatory flow maps delineate rapid, stage-specific causal signaling events, while coactivation maps capture broader, sustained network integration supporting speech processing.

### Limitations

This study is constrained by the spatial limitations inherent to iEEG in presurgical evaluation, where electrode placement is clinically determined and therefore excludes key subcortical structures such as the thalamus, whose signals have been shown to support effective speech decoding [66]. As a result, their potential contributions to neural information flow remain unexamined. Although electrodes in seizure onset zones, spiking regions, and MRI-visible lesions were excluded to minimize direct pathological influence, we also assessed potential remote effects of epilepsy. Consistent with our prior findings [10], patients with left-hemisphere epileptogenic foci exhibited reduced left-lateralization of neural interactions (**eResults**); nonetheless, there was no evidence that such effects distorted our primary conclusions. While spatial sampling constraints and residual pathology warrant consideration, the principal neural interaction patterns reported here appear robust.

The validity of our iEEG-based analyses has been extensively addressed in prior work [9–10]. To derive generalizable and statistically robust estimates of neural dynamics, we computed average high-gamma amplitudes across all patients contributing electrodes to each ROI. To ensure reliability, only ROIs with data from at least seven patients were included. The principal ROIs underpinning our key findings were sampled by more than 20 patients and included over 70 electrodes each (**eTable 2**). Leave-one-out analyses confirmed that no single patient disproportionately influenced the group means (Spearman’s rho ≈ 0.95–0.99; **eFigures 5–6**). High electrode counts per ROI yielded narrow confidence intervals around the mean high-gamma amplitudes (**eFigures 3–4**), supporting the generalization of observed modulations to the broader patient population. This approach parallels practical strategies in developmental neuroscience: for example, while the ideal framework for modeling changes in cortical metabolism or synaptic density from ages 1 to 20 would involve longitudinal tracking within individuals, in practice, normative trajectories are inferred by averaging cross-sectional data across individuals sampled at discrete developmental stages [35,67–68]. Assessing temporal relationships between electrode sites within individual patients offers the advantage of minimizing confounds introduced by inter-individual variability in neural activation. However, such within-subject analyses carry distinct limitations. Given the restricted spatial coverage of nonepileptic electrodes in any single patient, findings are often derived from sparse sampling, with some cortico-cortical pathways represented by as few as three individuals or fewer. One strategy to mitigate these sampling constraints involves aggregating signals across broader anatomical extents. Yet, this approach hinges on the assumption that neural interaction dynamics are spatially homogeneous across the expanded ROI—a presumption that may not hold for functionally heterogeneous areas. Some investigators may define regions of interest based on the similarity of iEEG-derived neural signatures; however, this method needs to be applied cautiously to avoid circular inference [69]. To preserve anatomical specificity and analytic rigor, our study defined regions of interest strictly based on established gyral boundaries using the Desikan–Killiany atlas [34].

We urge caution when interpreting neural interaction measures at time windows temporally distal from stimulus presentations or the behavioral response. Inter-individual differences in response timing can blur the functional specificity of high-gamma signals in these late intervals. For example, while the moment of response onset is consistently linked to articulatory execution across patients, the cortical activity 500 ms prior to the response likely differs considerably between individuals; such variability can degrade the fidelity of signals when averaging across patients. However, in our data we did not observe a marked inflation in confidence interval width for these temporally distant windows (**eFigures 3–4**). Since confidence interval width serves as a proxy for inter-individual consistency, this finding suggests that our group-level estimates remained robust despite potential timing variability across patients. Importantly, convergent evidence from electrical stimulation mapping [11–14] validated our iEEG-based dynamic tractography atlases, reinforcing that the observed neural interactions reflect functionally meaningful network dynamics rather than statistical artifacts.

## Supporting information

eVideo 4

eVideo 1

eVideo 2

eVideo 3

## Acknowledgments

We are grateful to Sandeep Sood, MD, Alanna Carlson, MS, LLP, Jamie MacDougall, RN, BSN, CPN at Children’s Hospital of Michigan for the collaboration and assistance in performing the studies described above.

## Data availability

The data are publicly available through OpenNeuro: https://openneuro.org/datasets/ds006910/versions/1.0.1 https://openneuro.org/datasets/ds006914/versions/1.0.3

## Code availability

The codes are openly available on GitHub: https://github.com/rkwsu/Project_Auditory_Visual_EC

## Supplementary Figures: eFigures

**eFigure 1.**
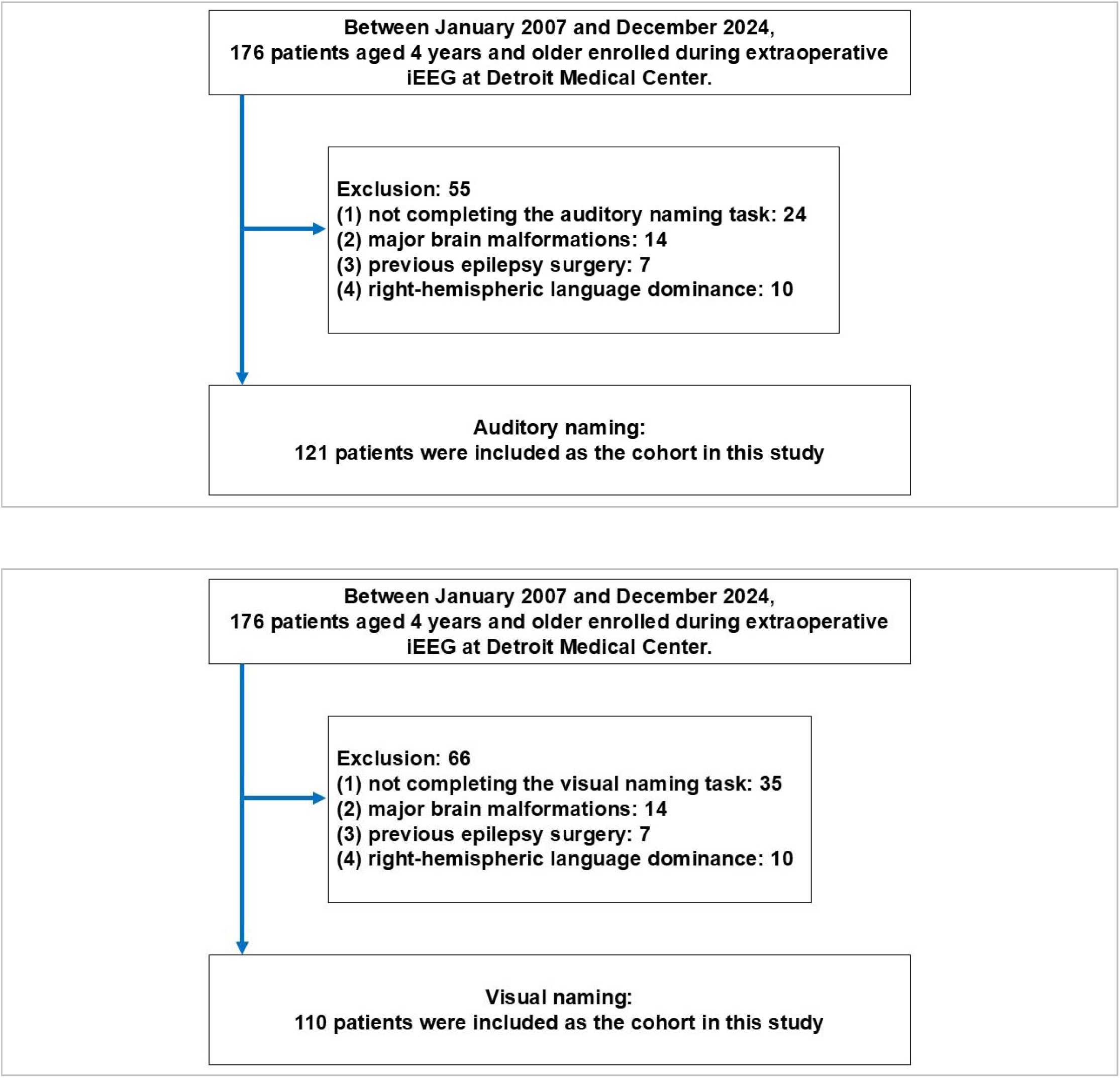
Flowcharts showing the selection of patients who met eligibility criteria. A total of 176 native English-speaking patients aged ≥4 years were enrolled. Of these, 24 did not complete the auditory naming task and 35 did not complete the visual naming task. An additional 14 patients were excluded due to major brain malformations, 7 due to a history of epilepsy surgery, and 10 due to right-hemispheric language dominance. Thus, intracranial EEG data were available from 121 patients for auditory naming and 110 patients for visual naming, with 104 patients contributing data for both tasks.

**eFigure 2.**
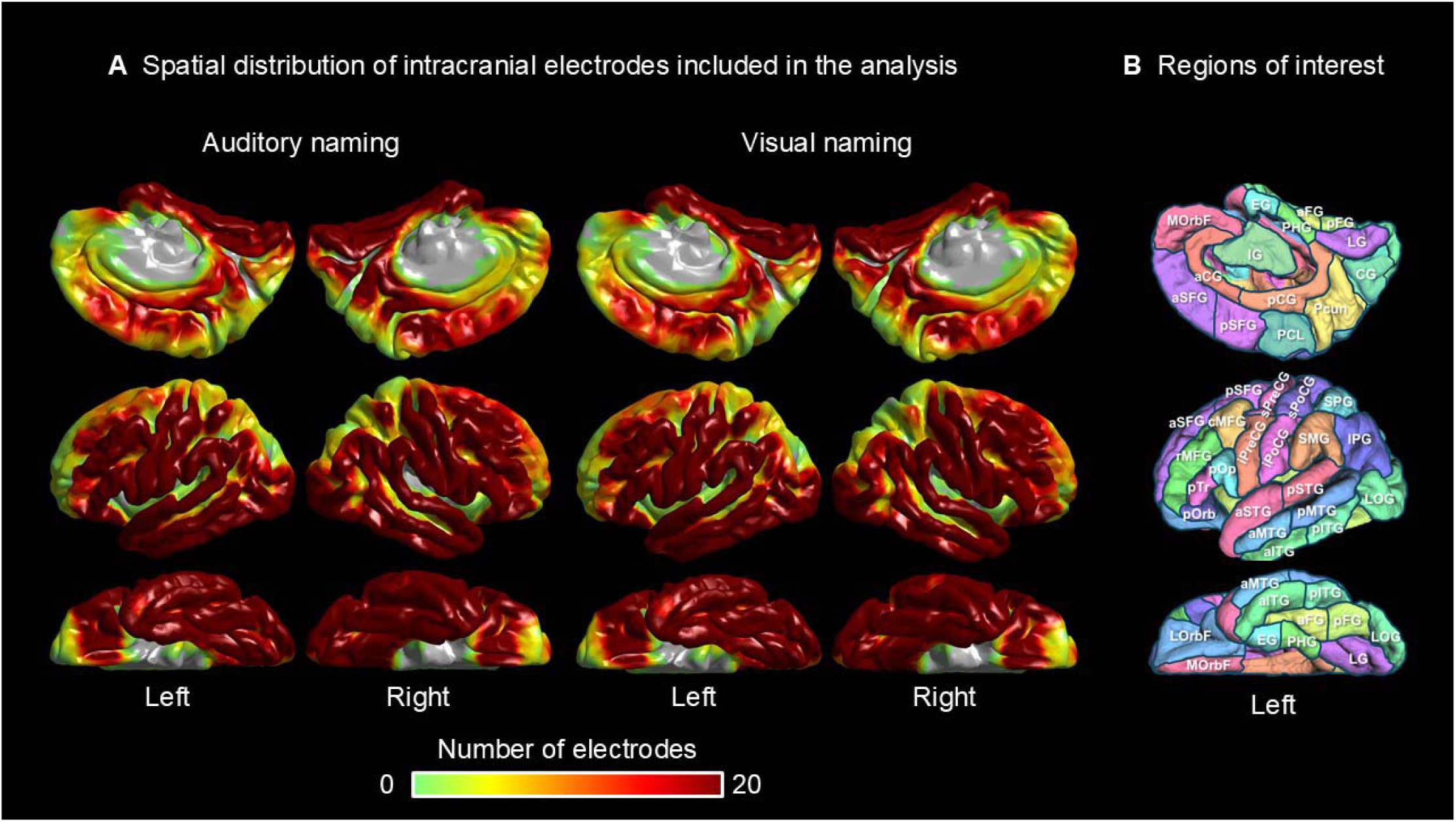
Intracranial electrode coverage. **A** Number of artifact-free, non-epileptic sites analyzed for high-gamma activity during auditory (left) and picture naming (right). **B** Left hemispheric ROIs based on the FreeSurfer standard brain space; full names listed in **eTable 2**.

**eFigure 3.**
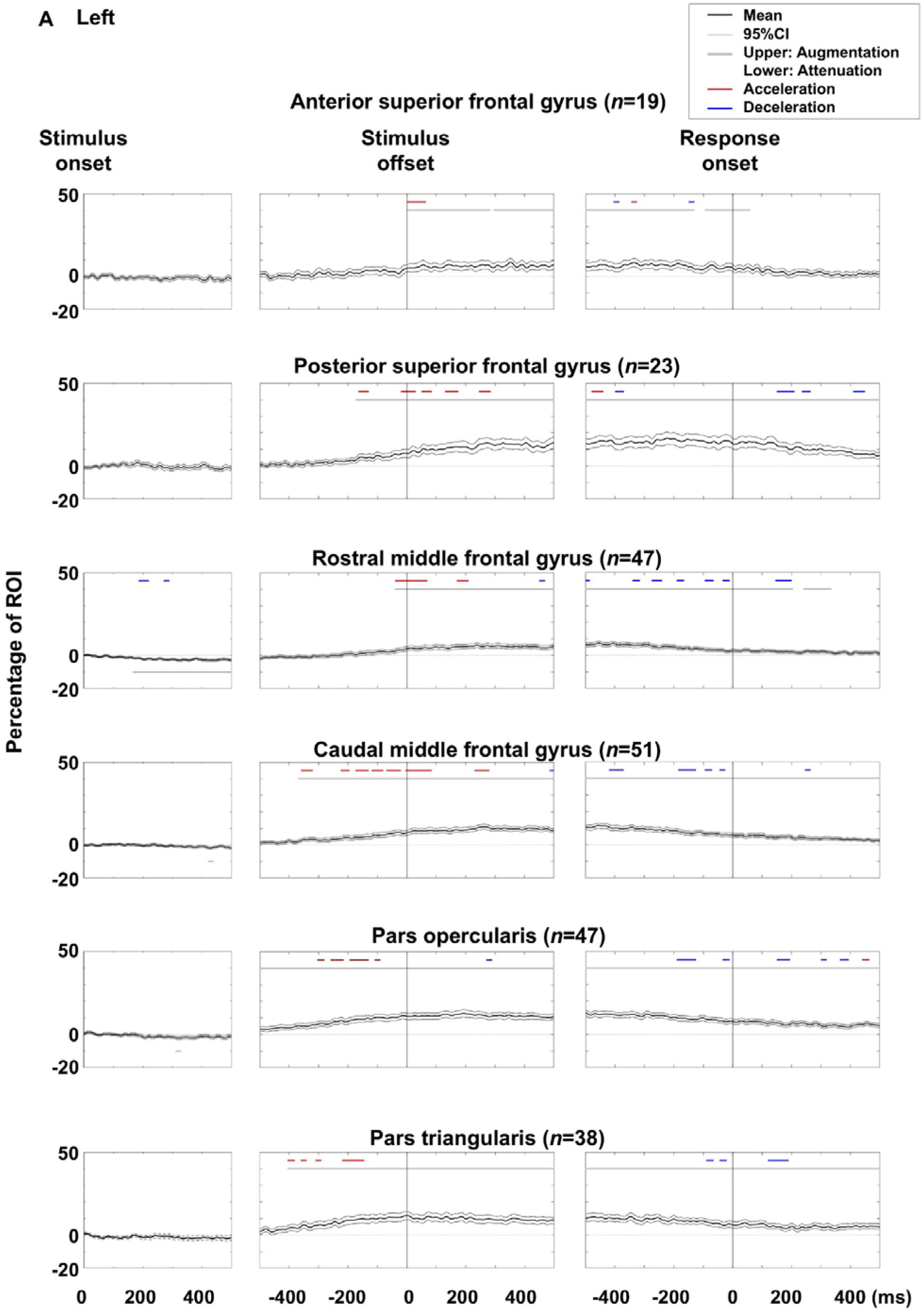
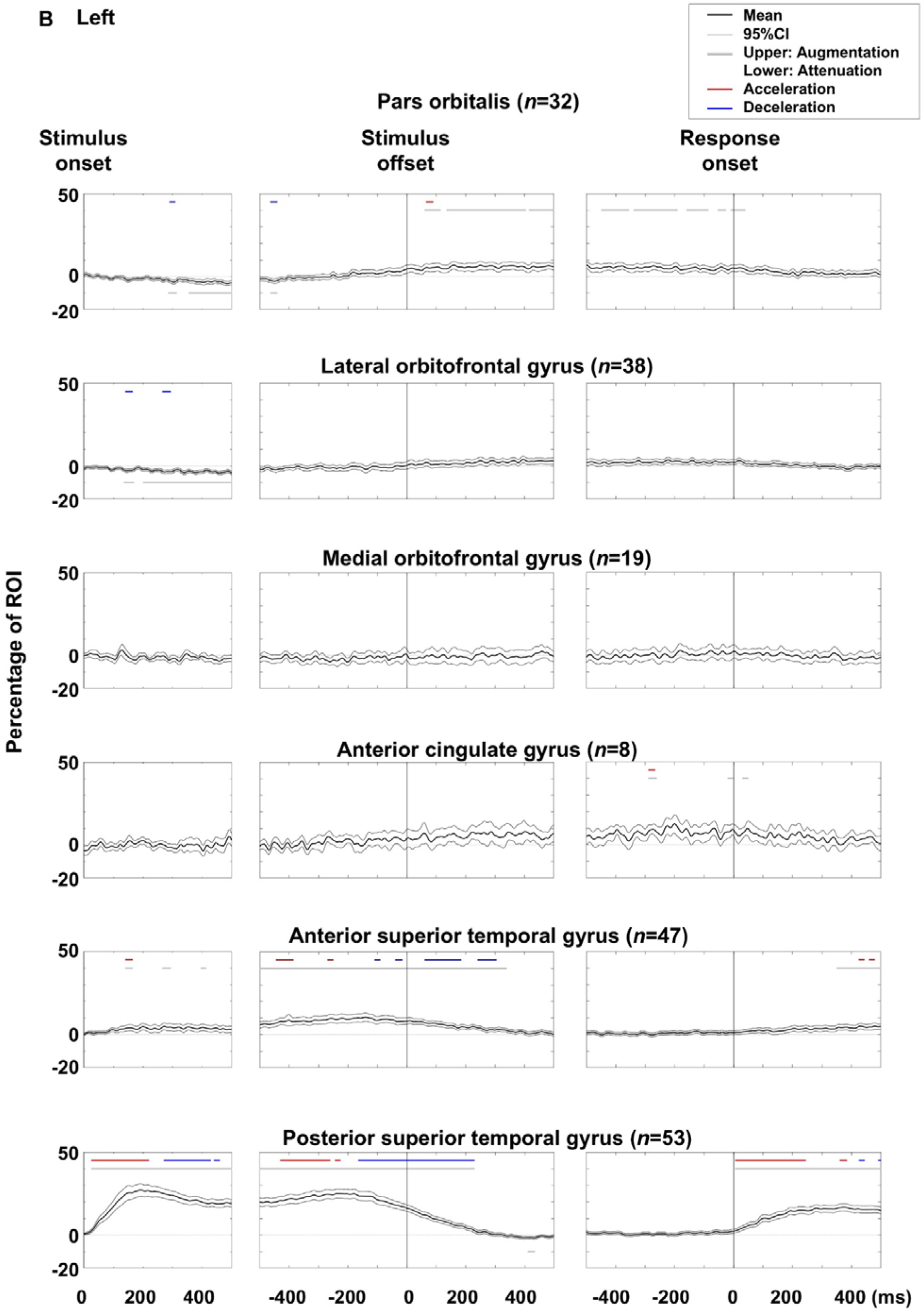
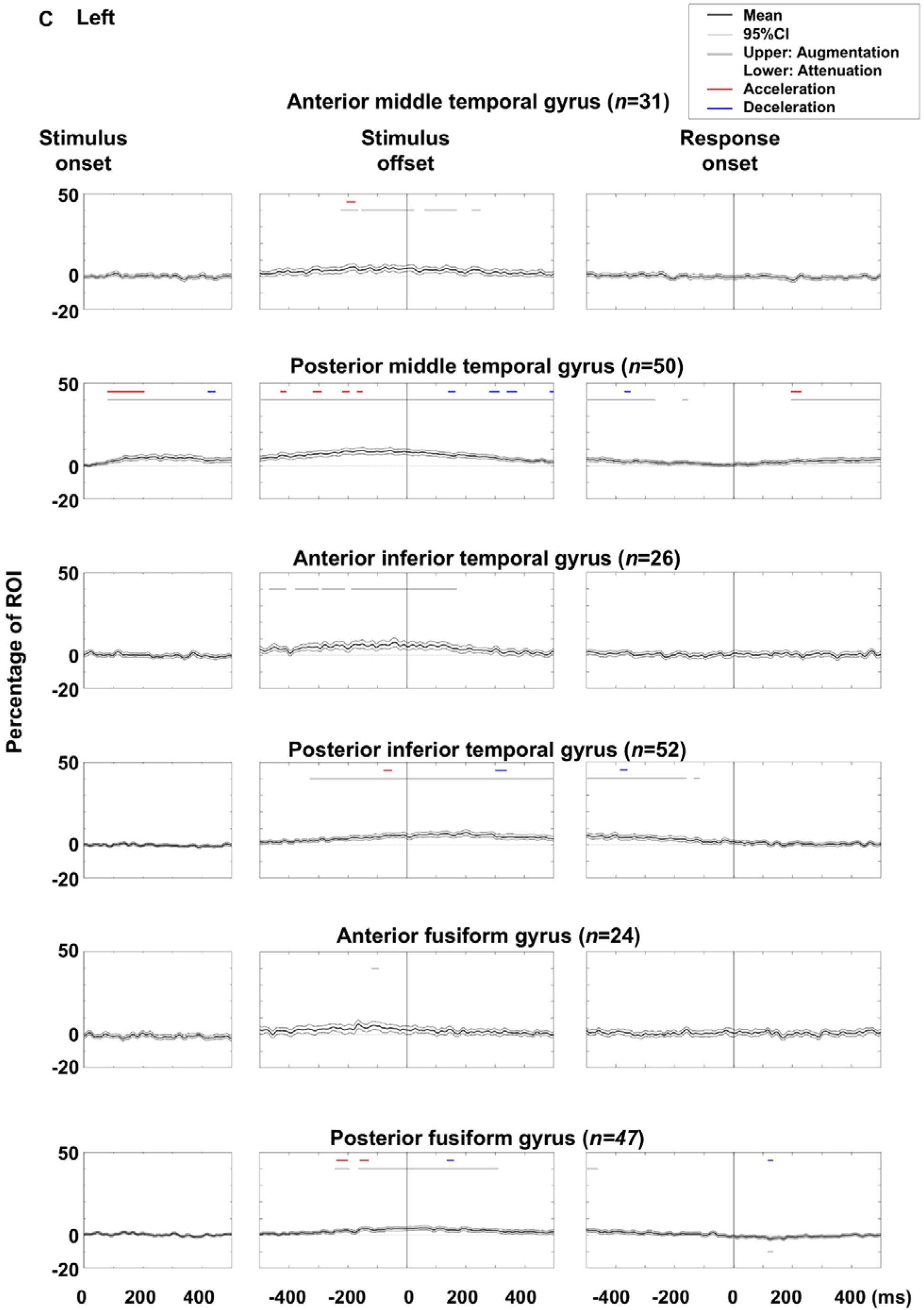
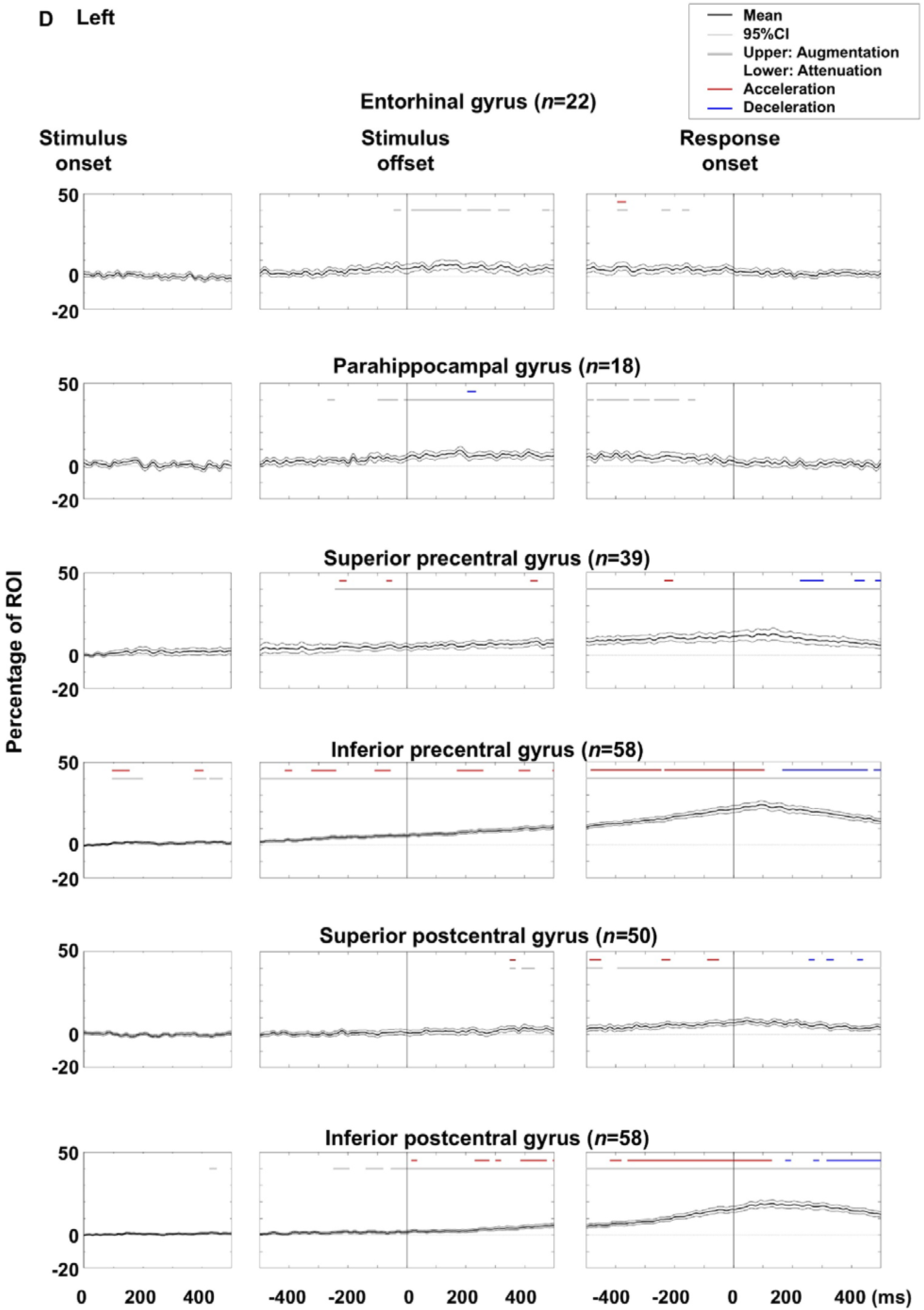
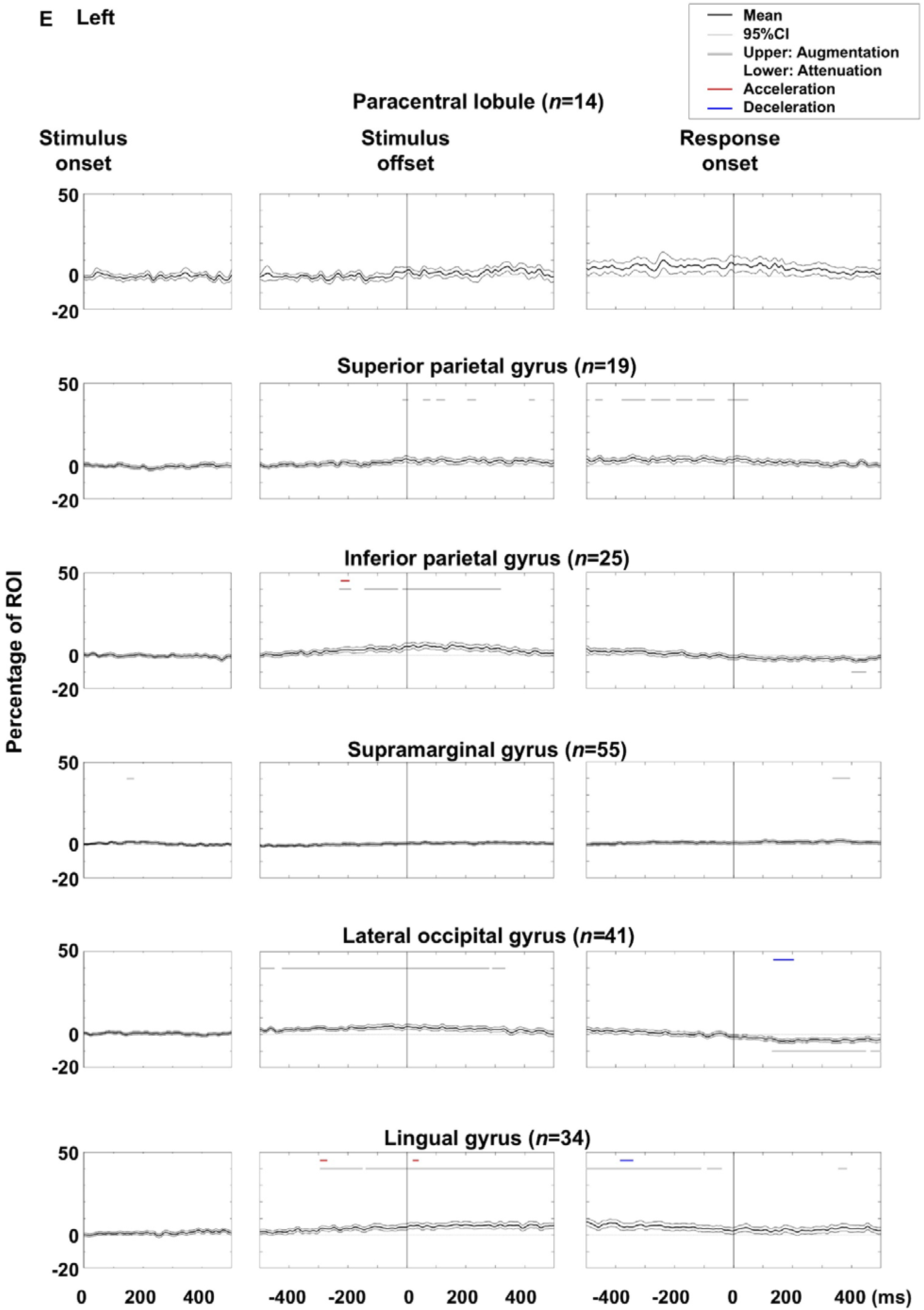
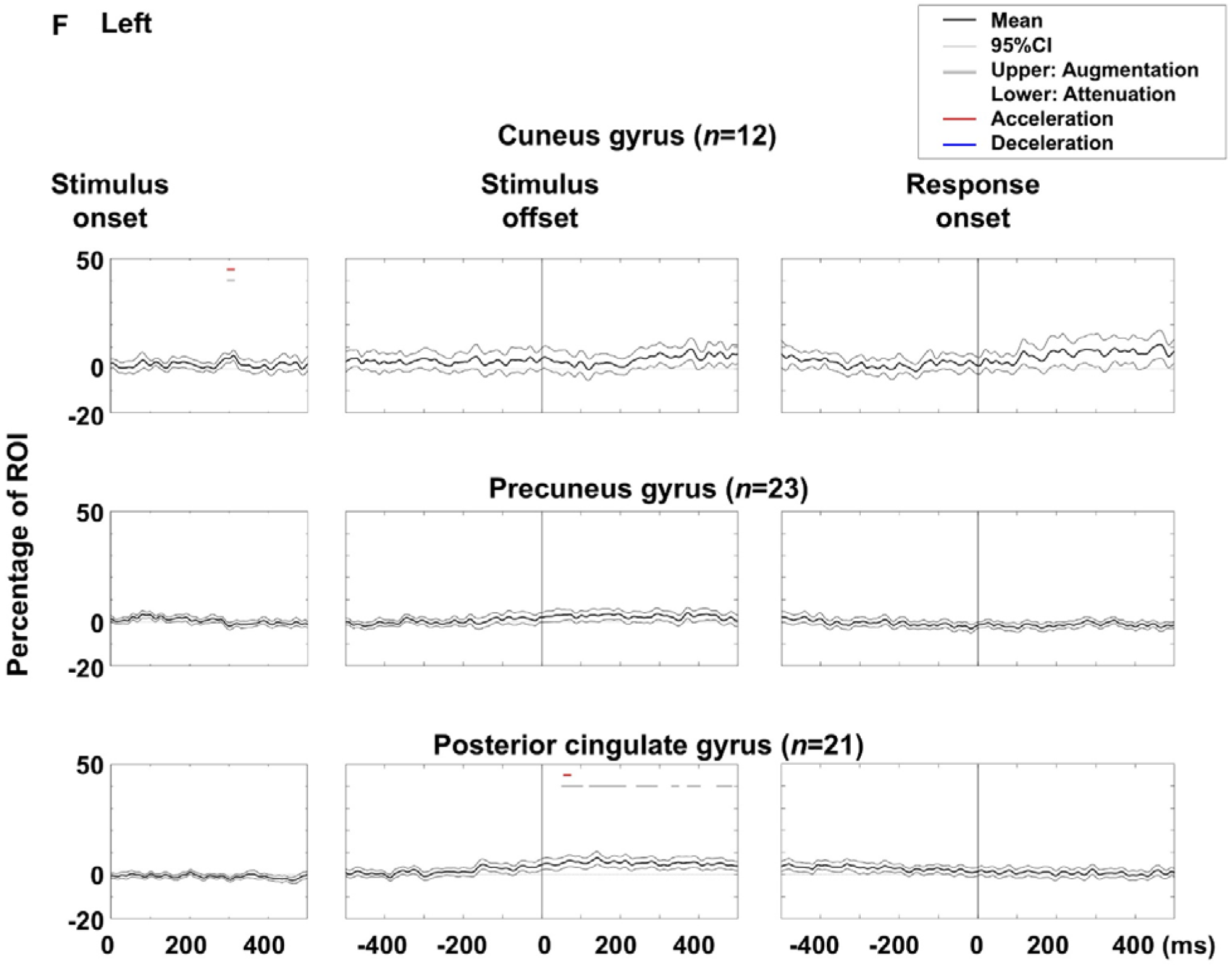
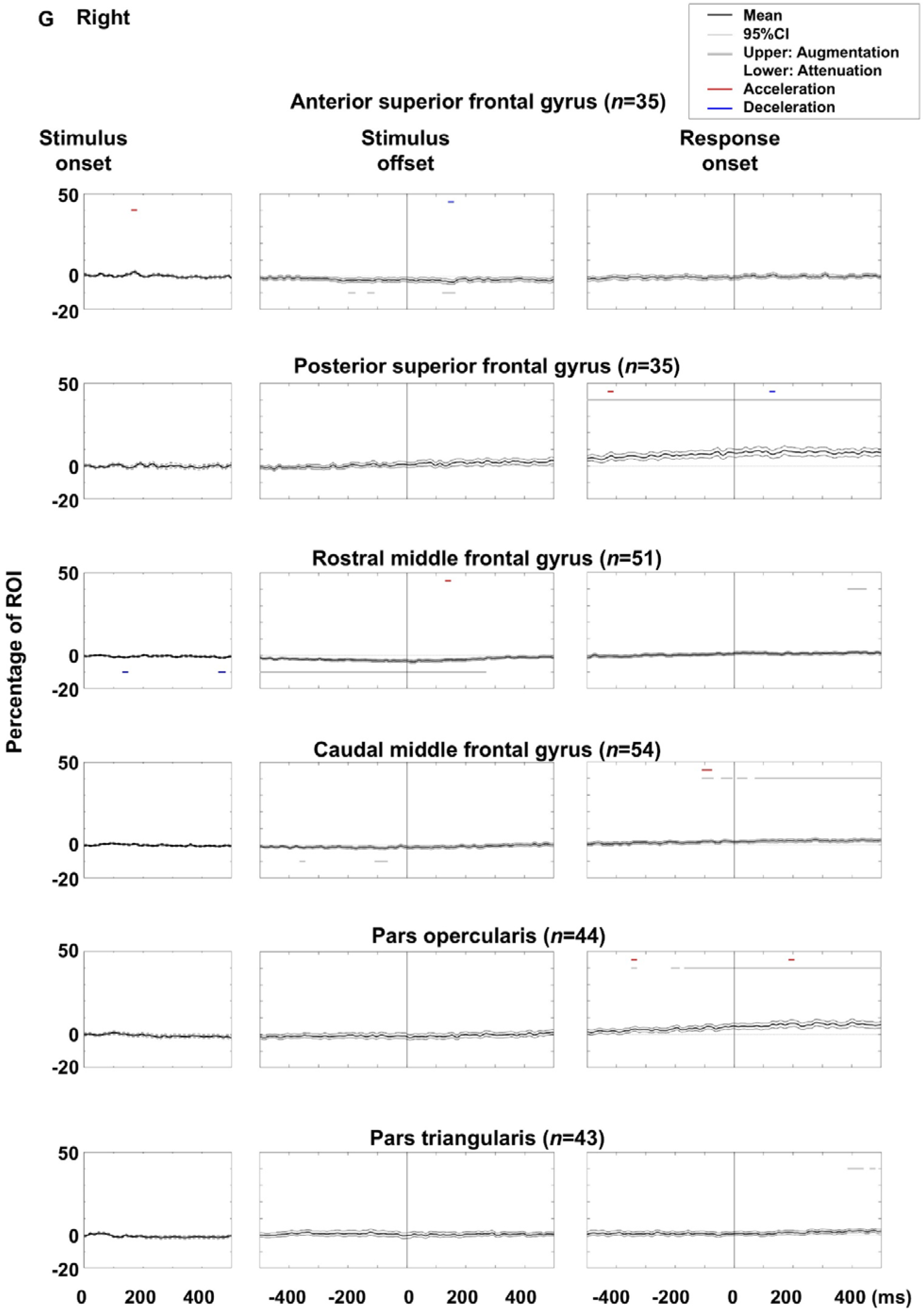
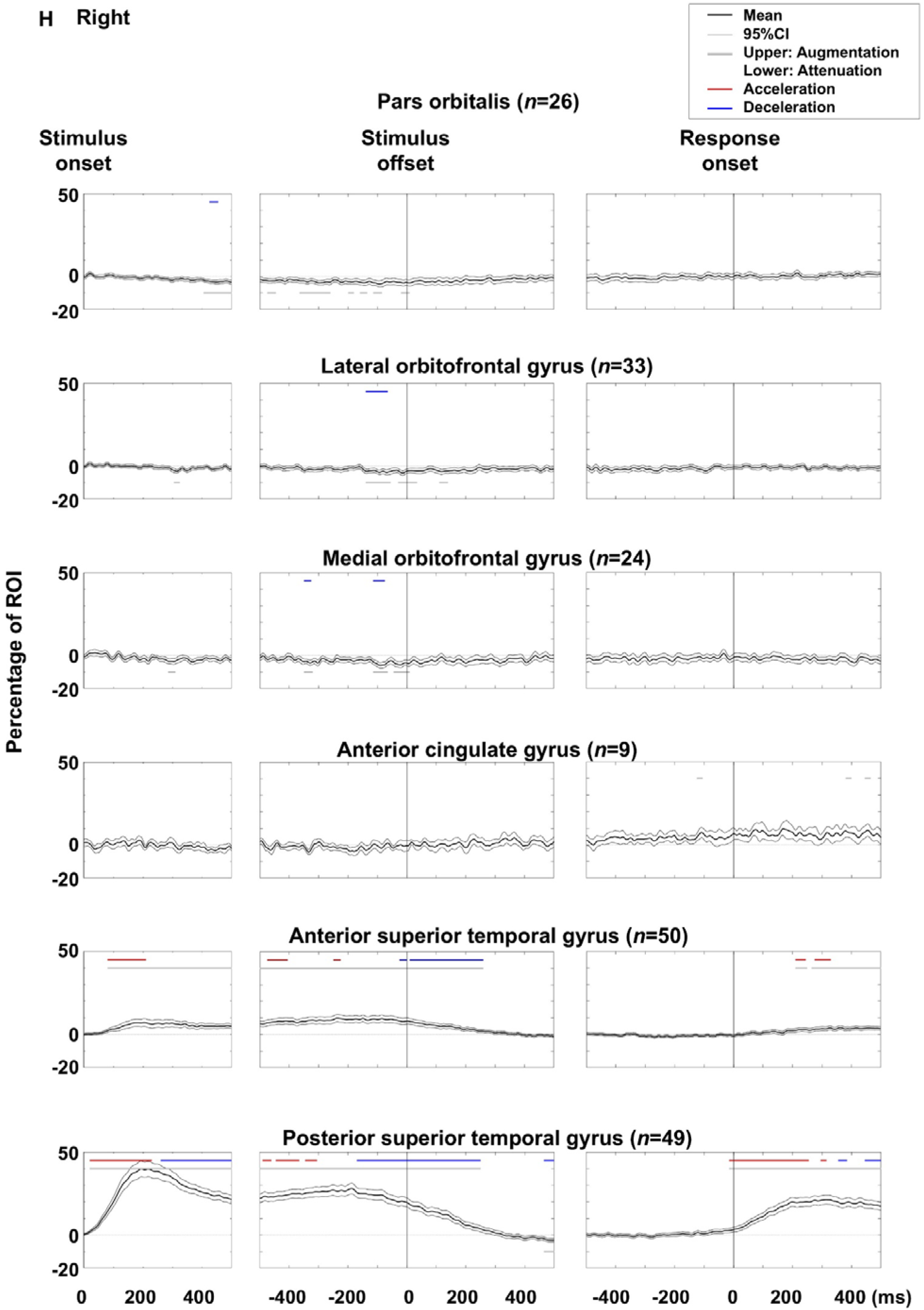
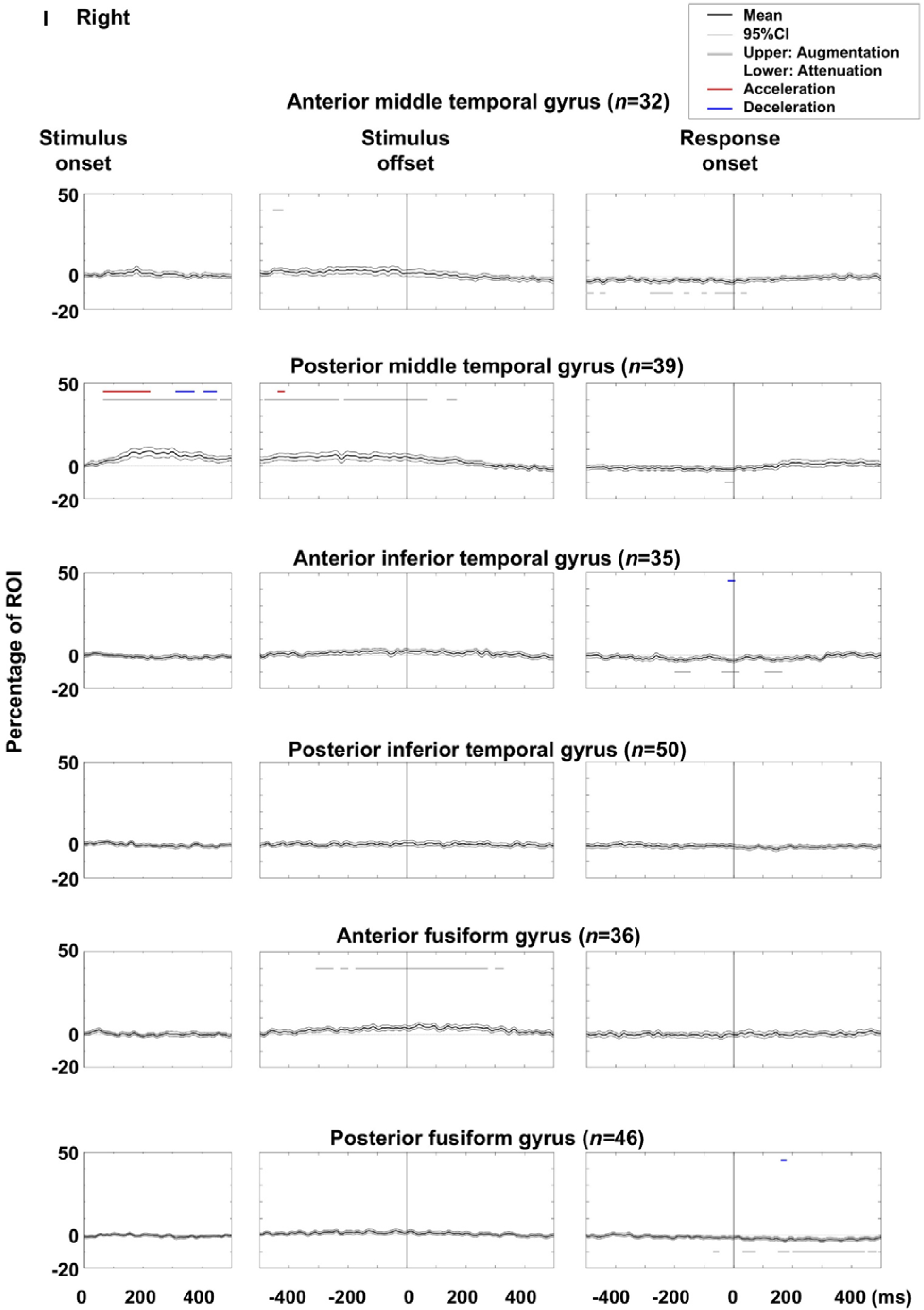
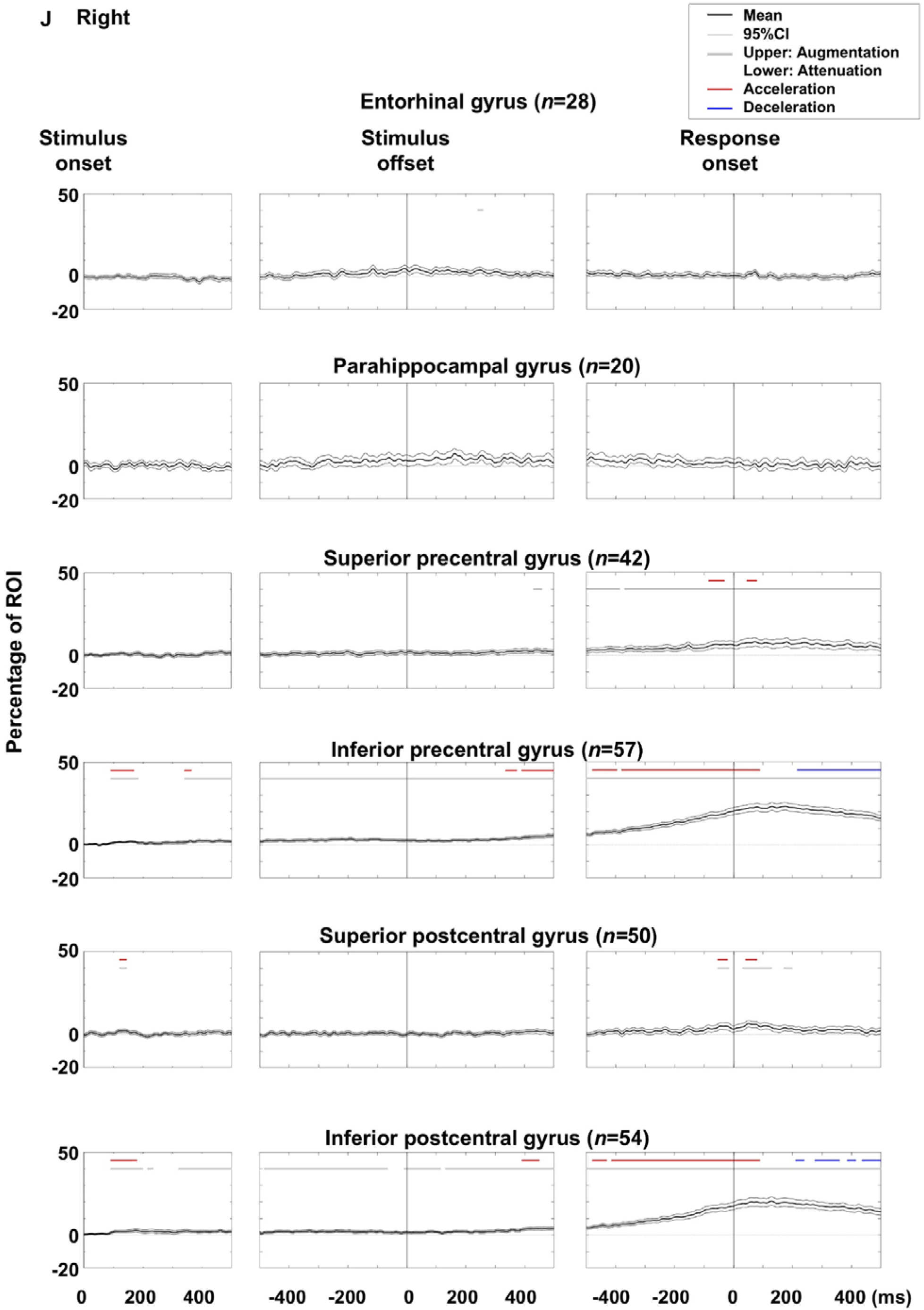
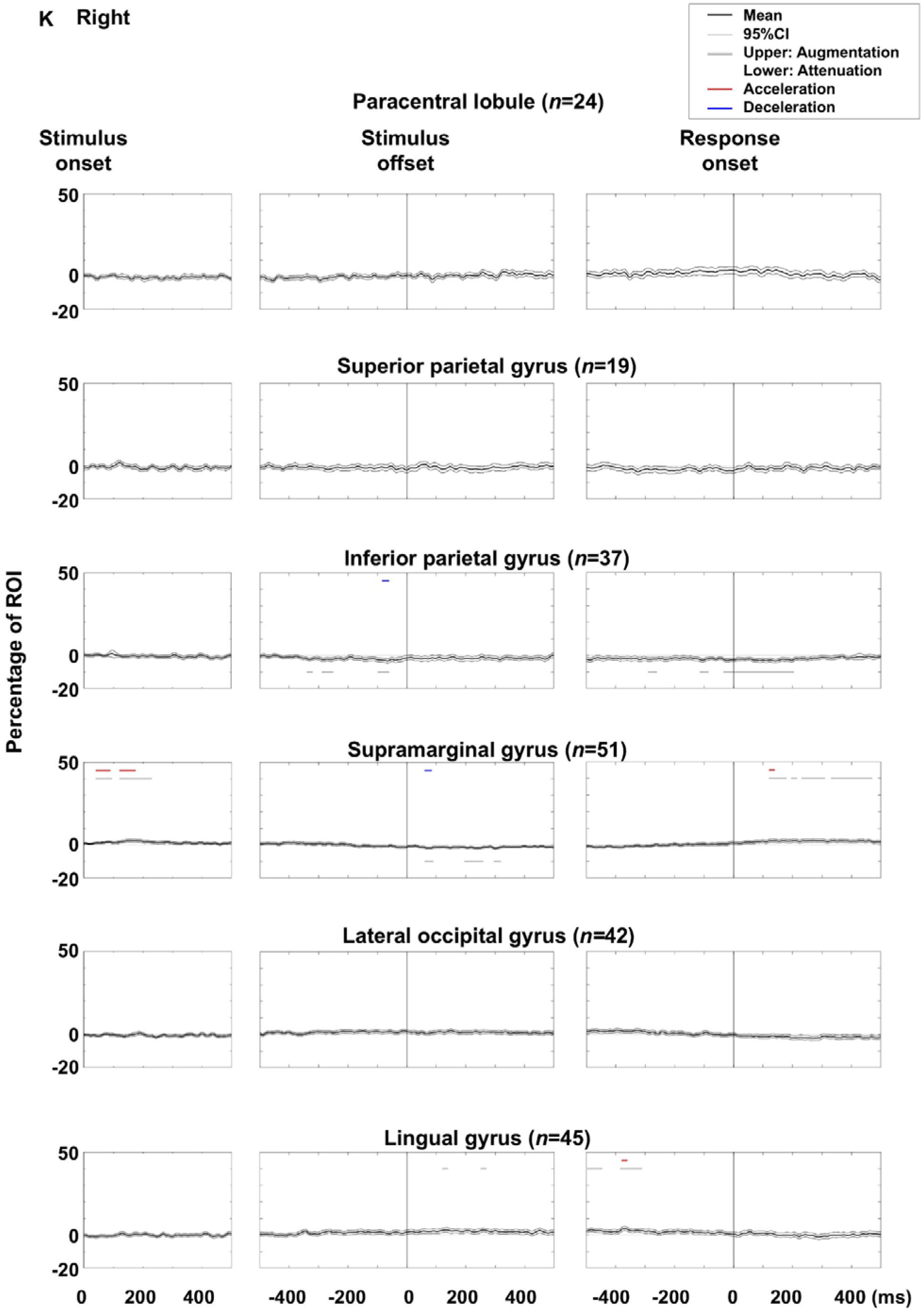
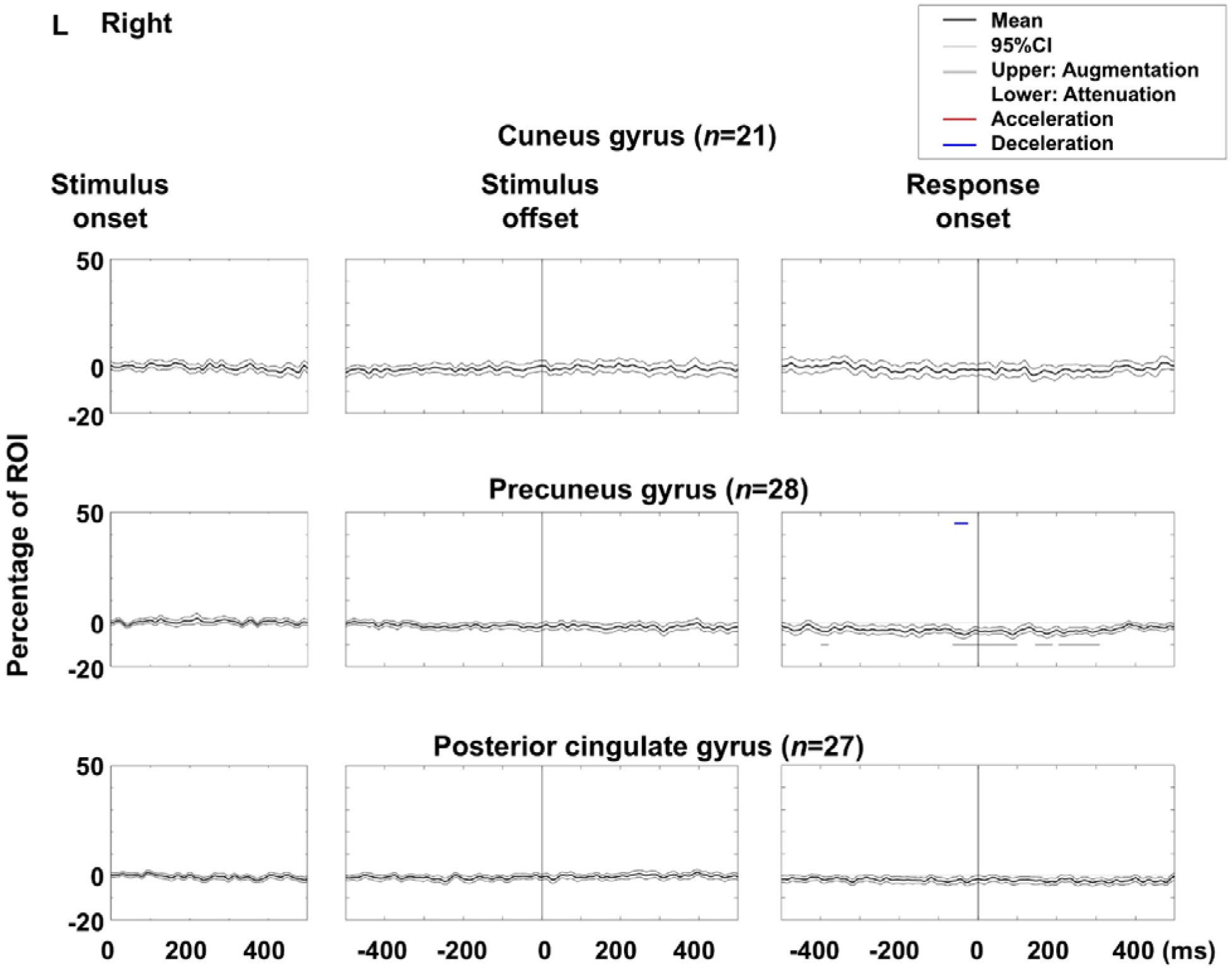
Auditory naming-related high-gamma dynamics: distinguishing augmentation from acceleration and attenuation from deceleration. Mean high-gamma amplitude (% change relative to baseline) at each region of interest (ROI) is shown with 95% confidence intervals. Upper and lower horizontal bars denote time bins with significant high-gamma augmentation and attenuation, respectively, while red and blue bars indicate significant high-gamma acceleration and deceleration. **A–F** Left hemispheric ROIs. **G–L** Right hemispheric ROIs.

**eFigure 4.**
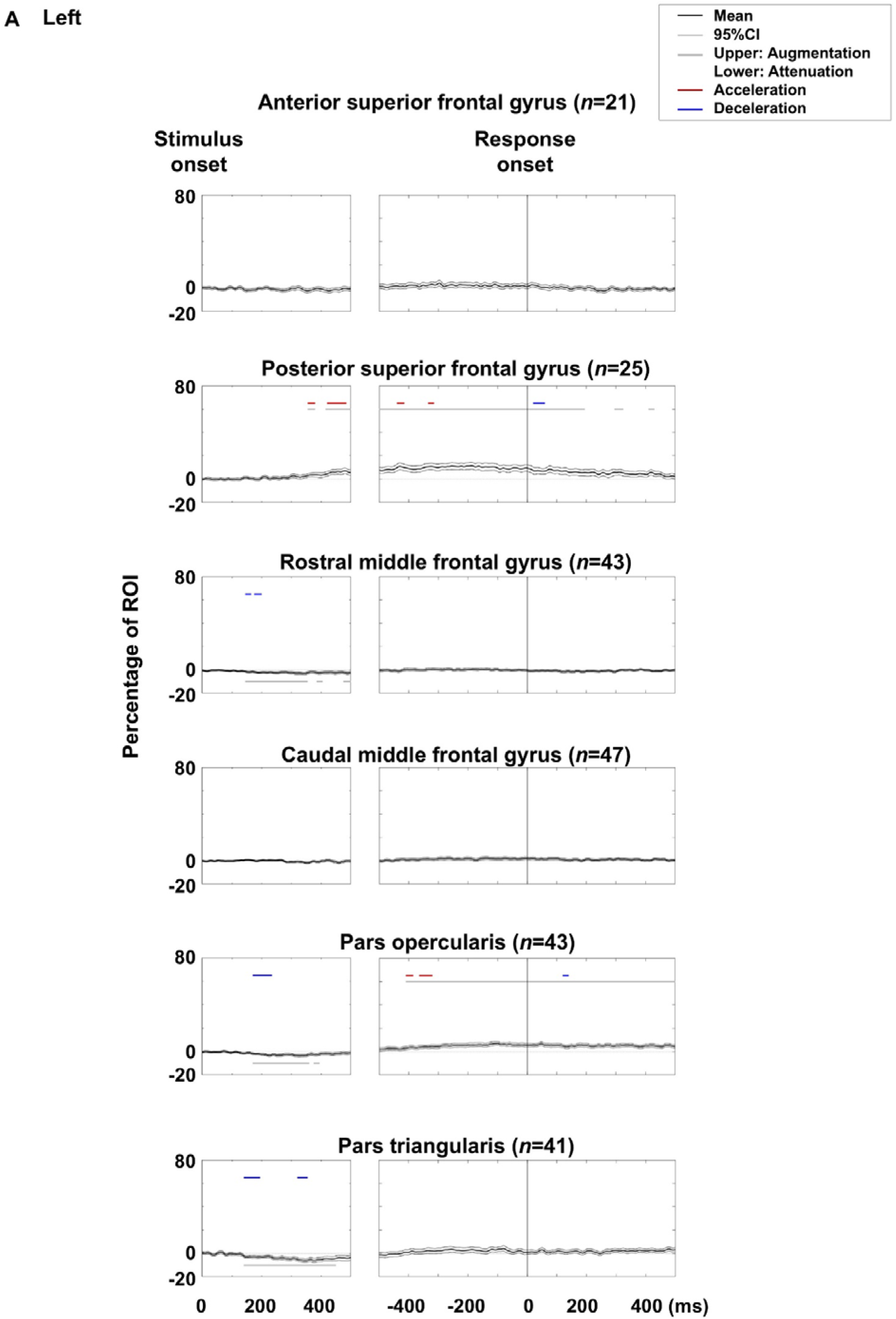
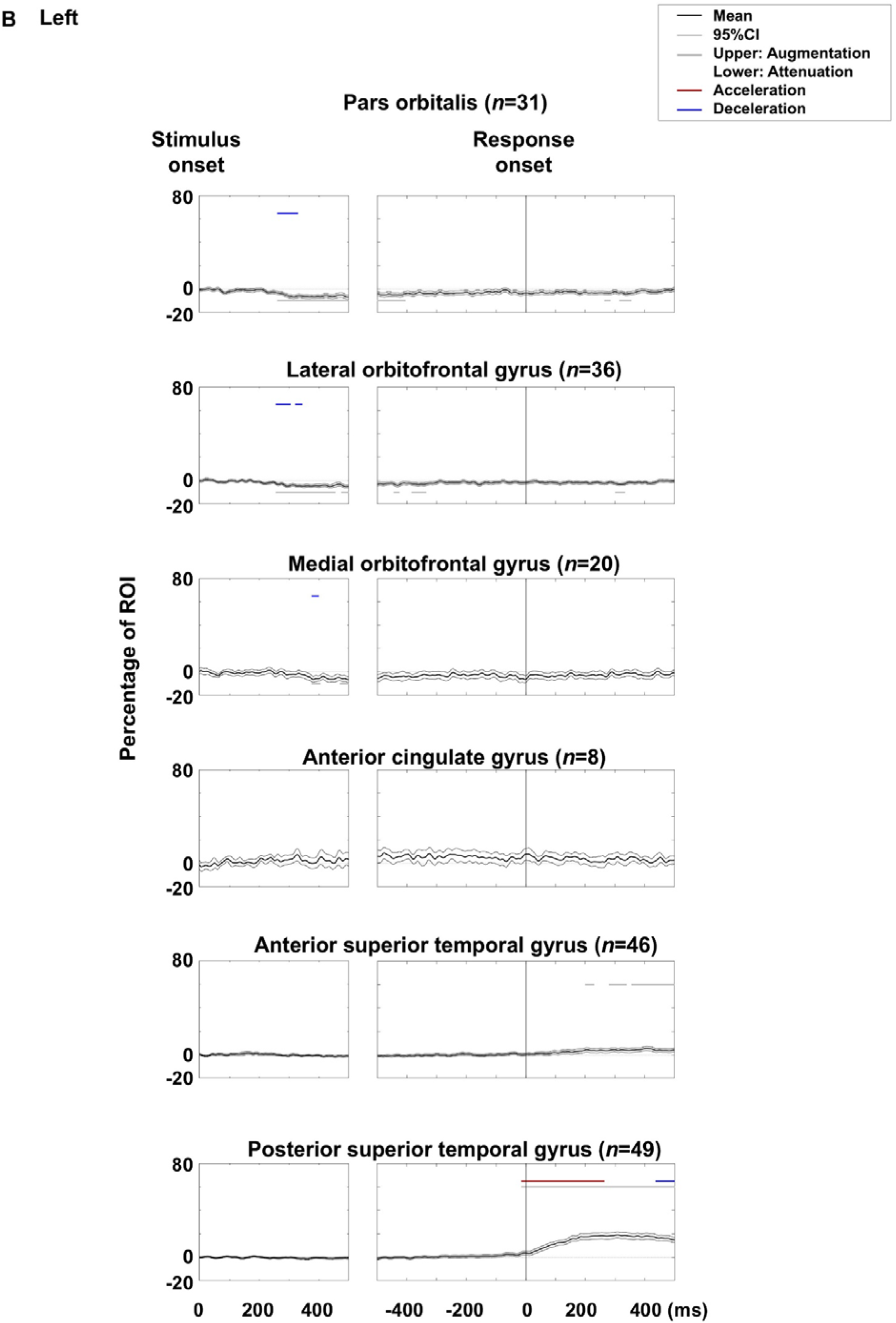
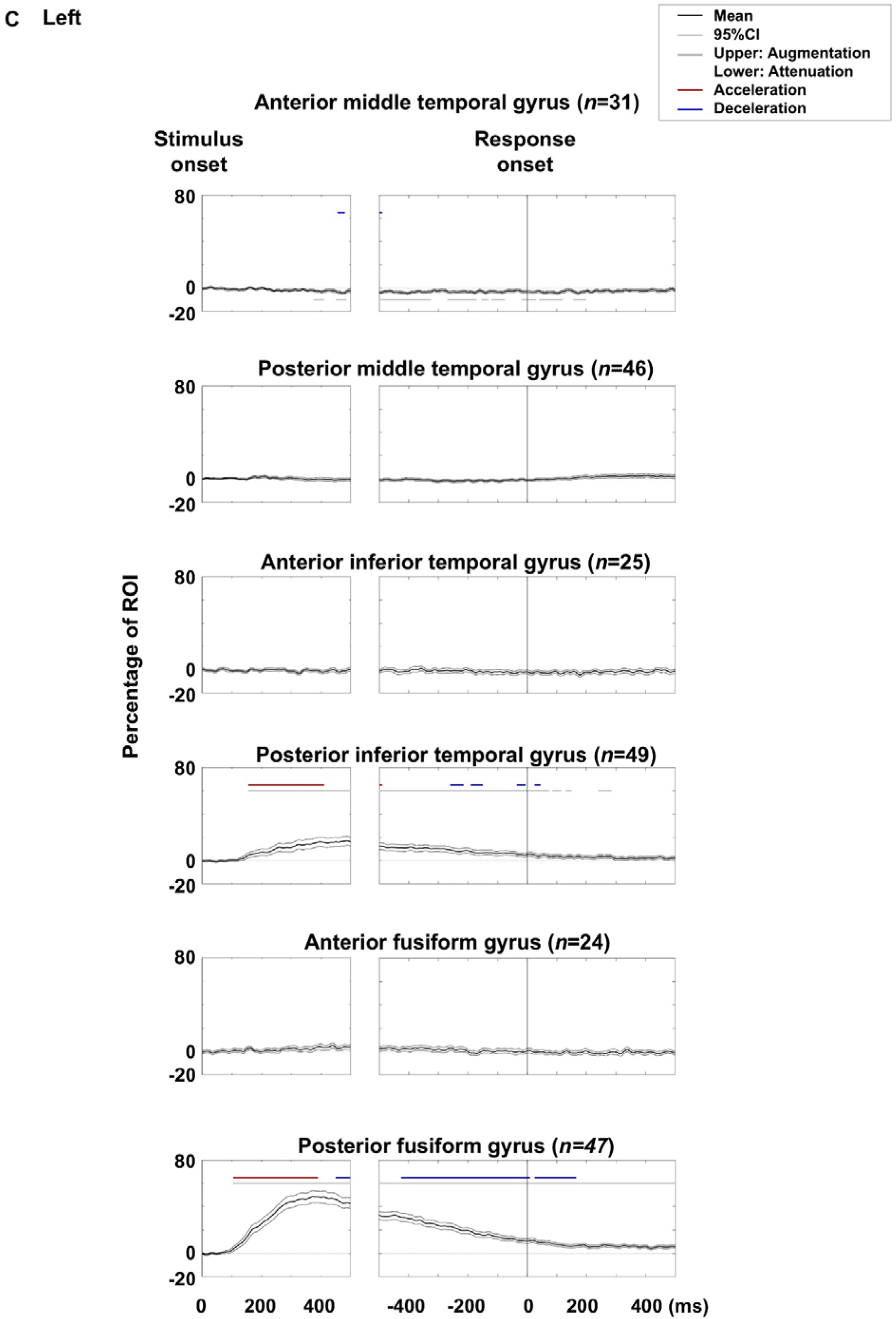
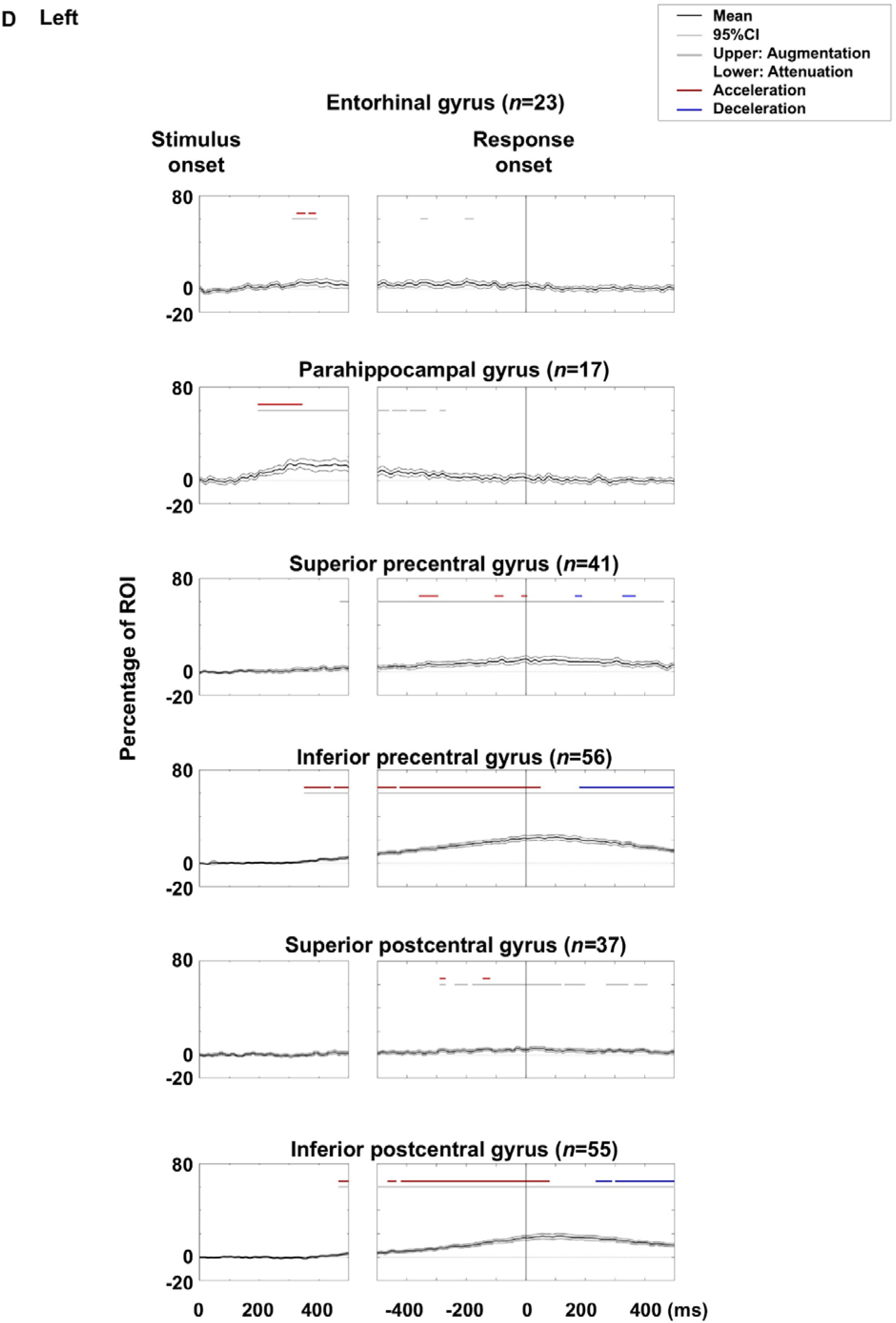
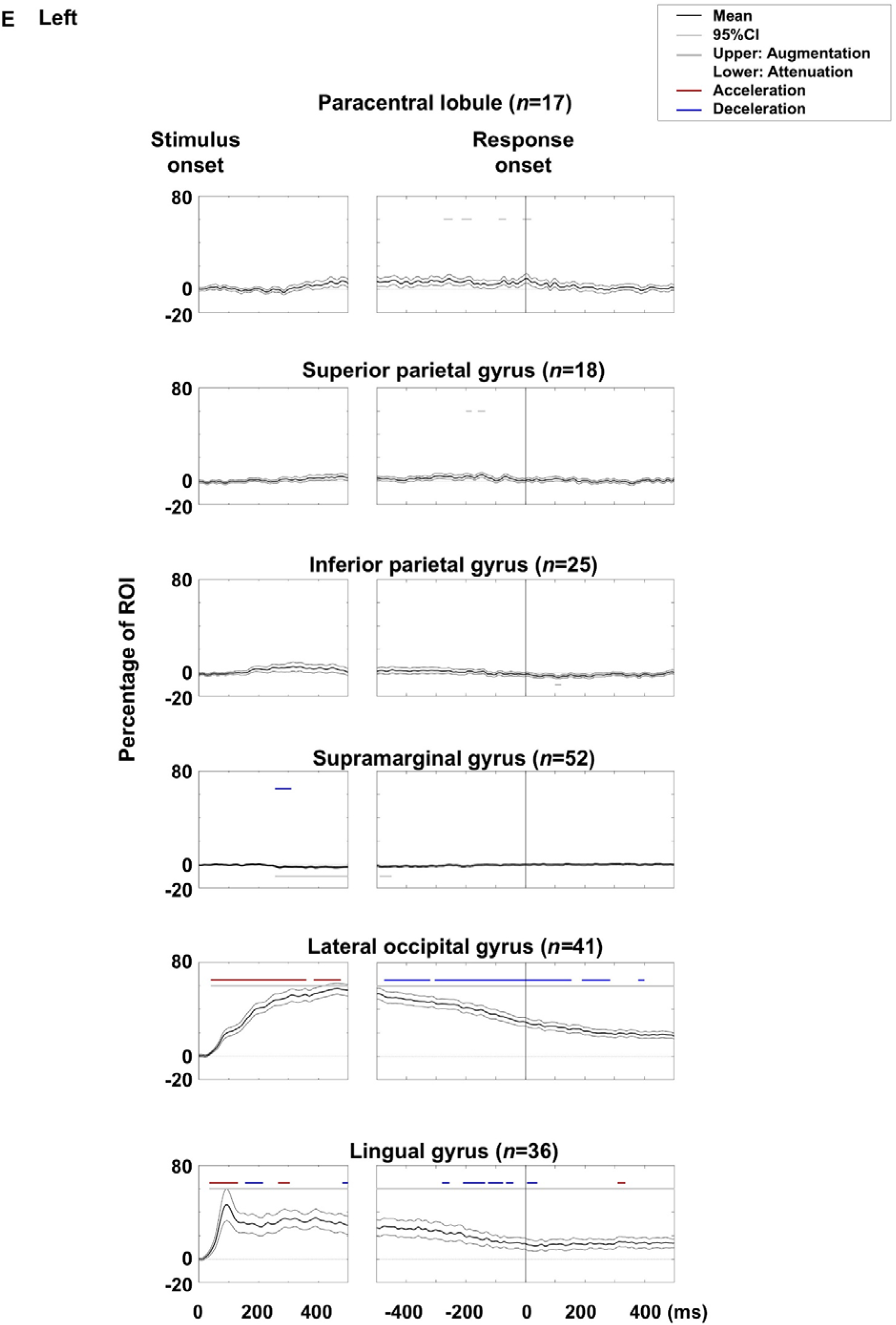
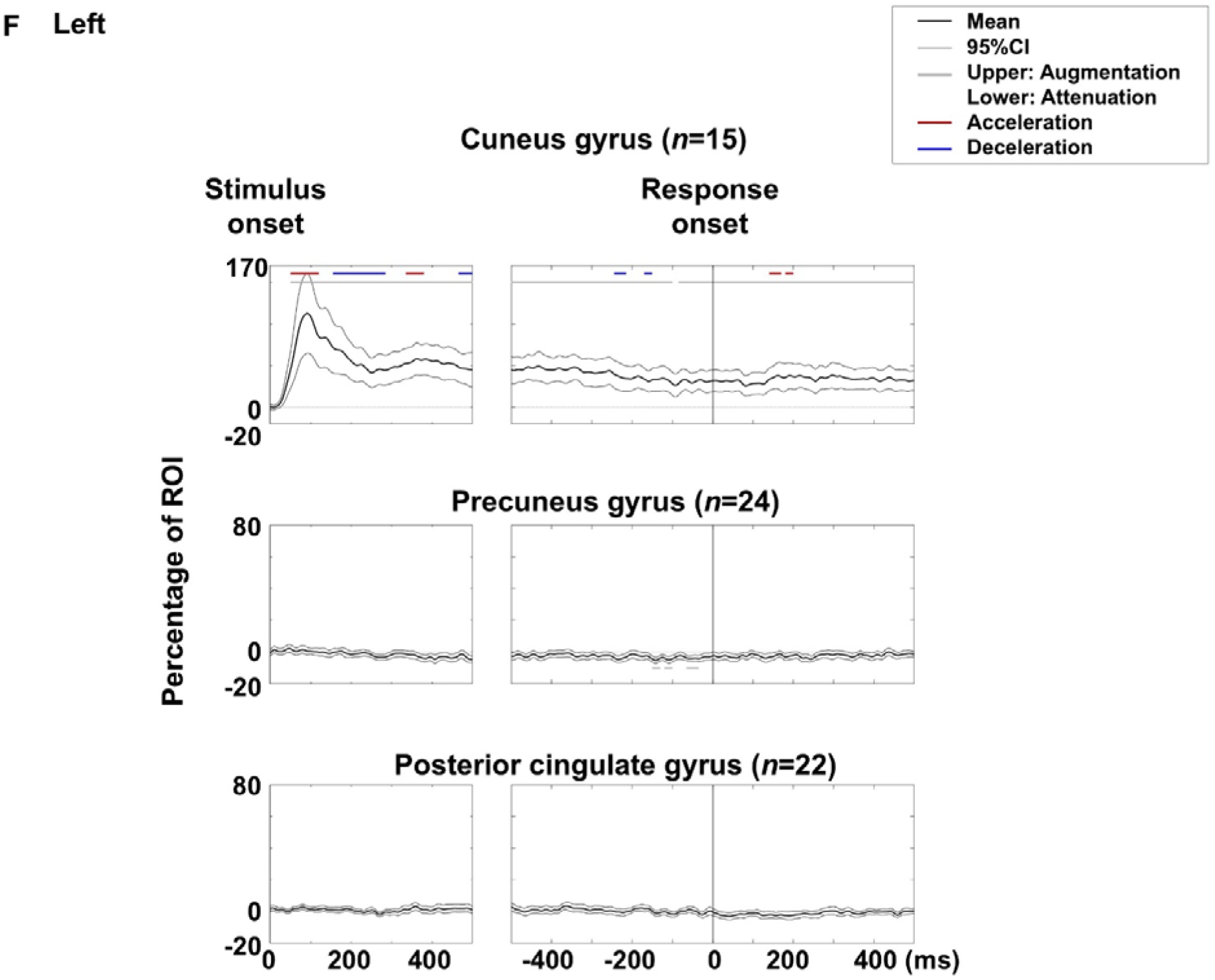
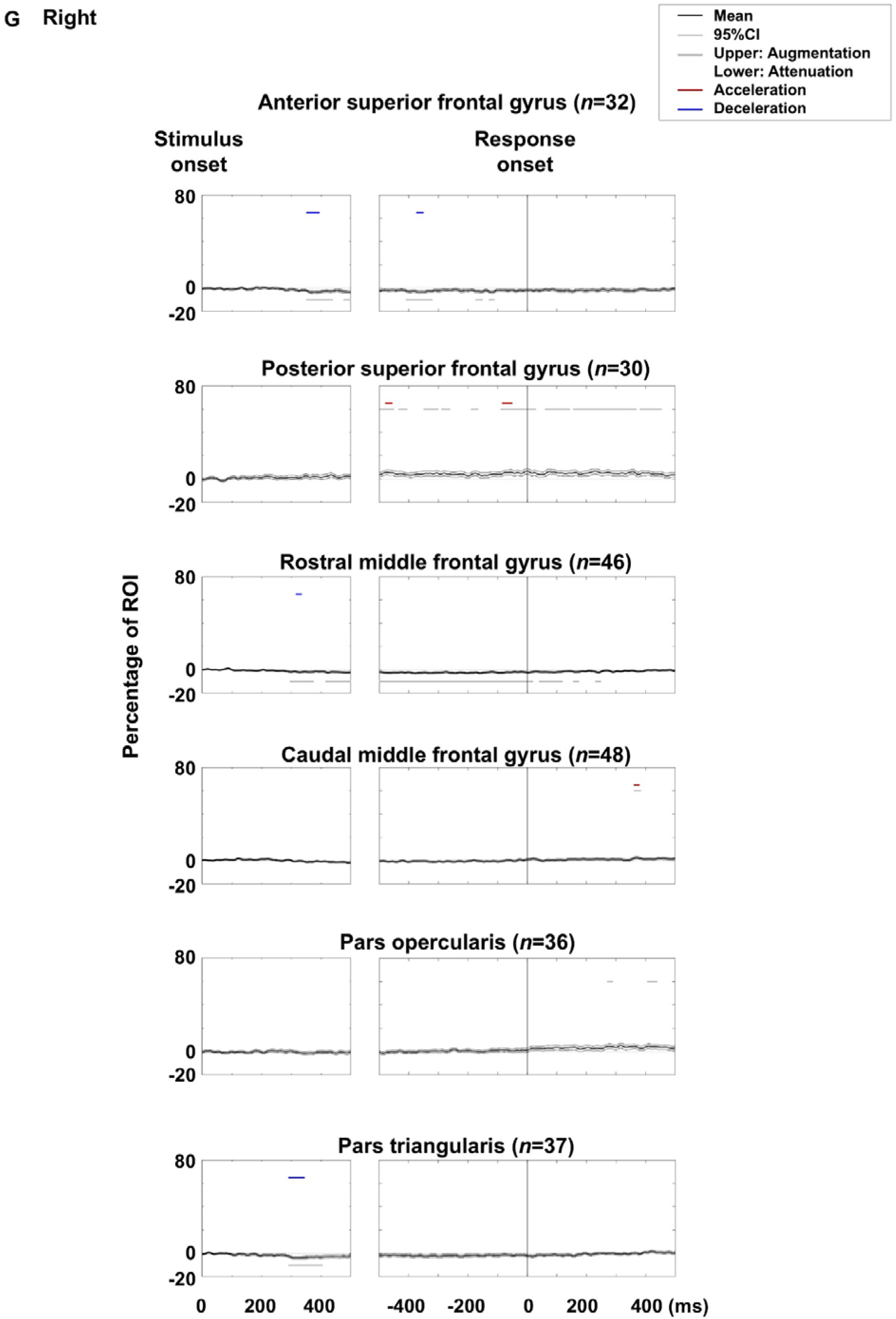
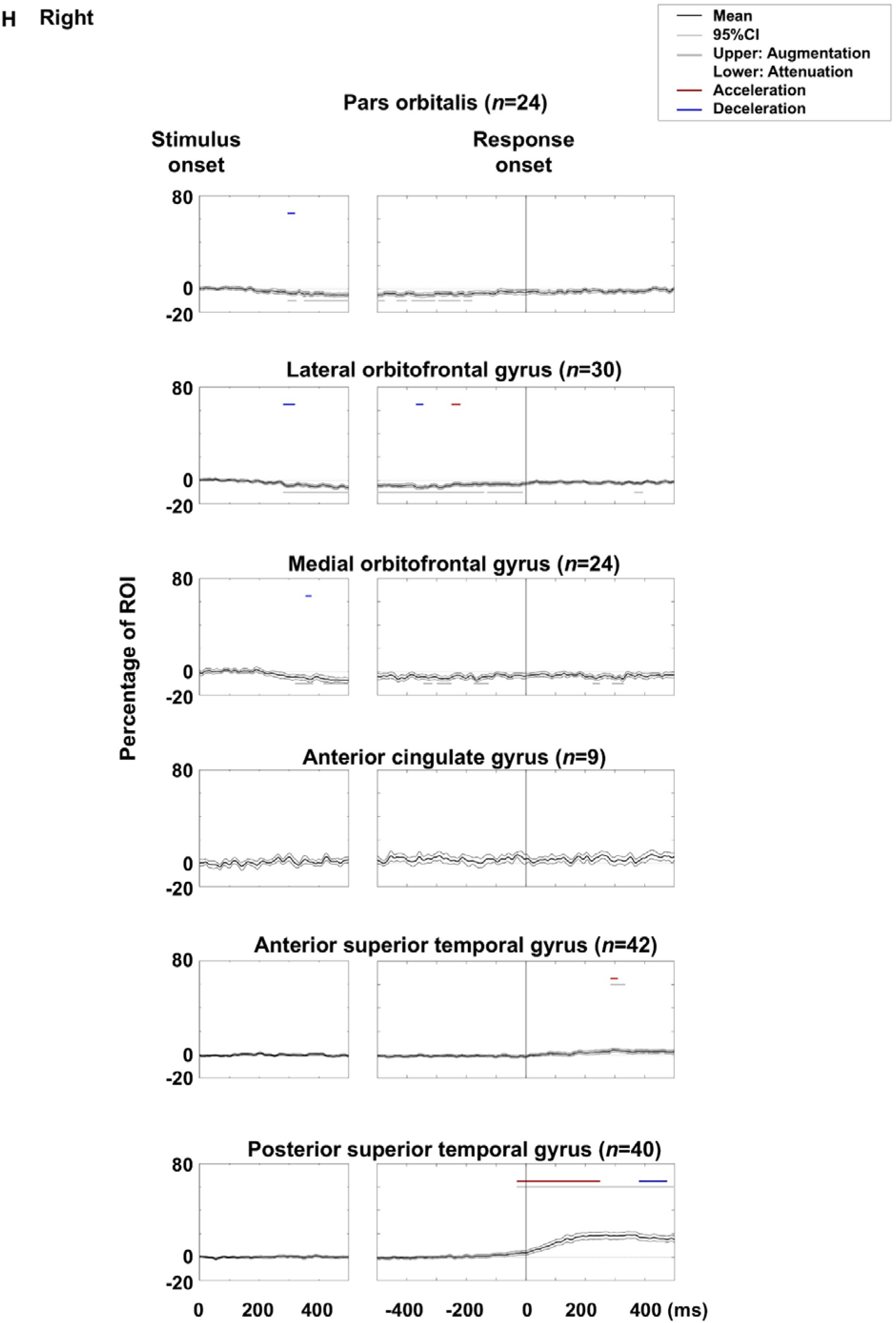
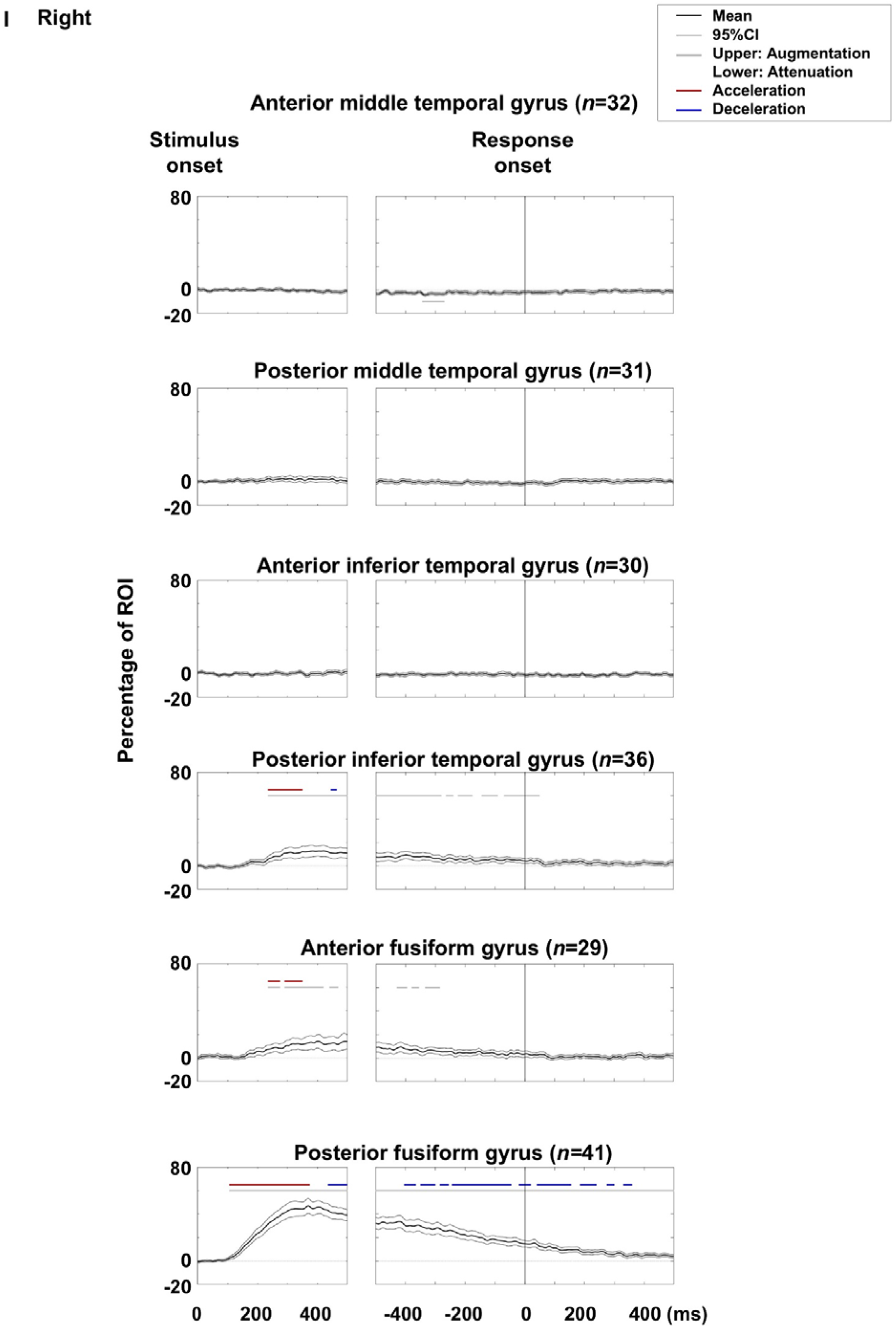
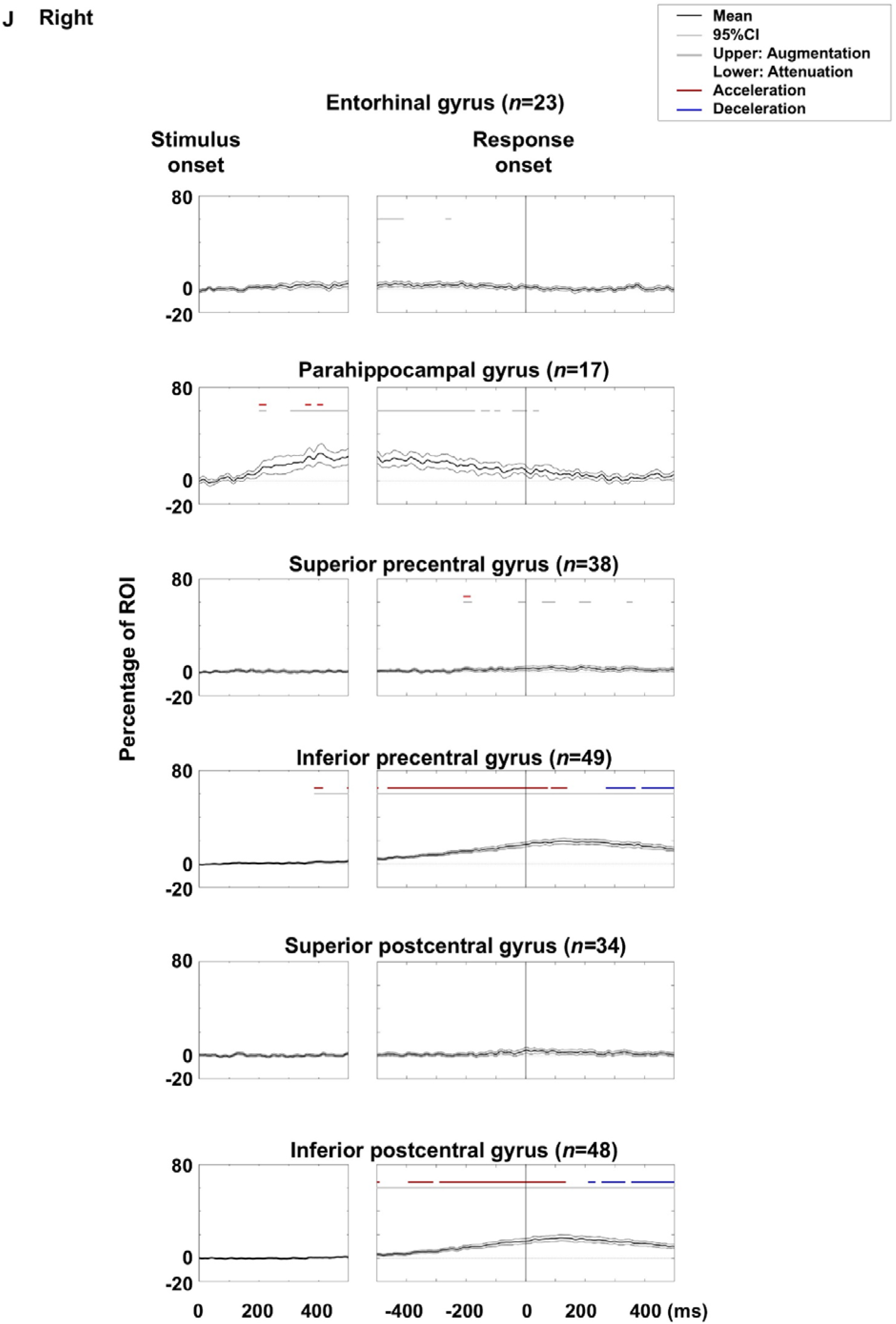
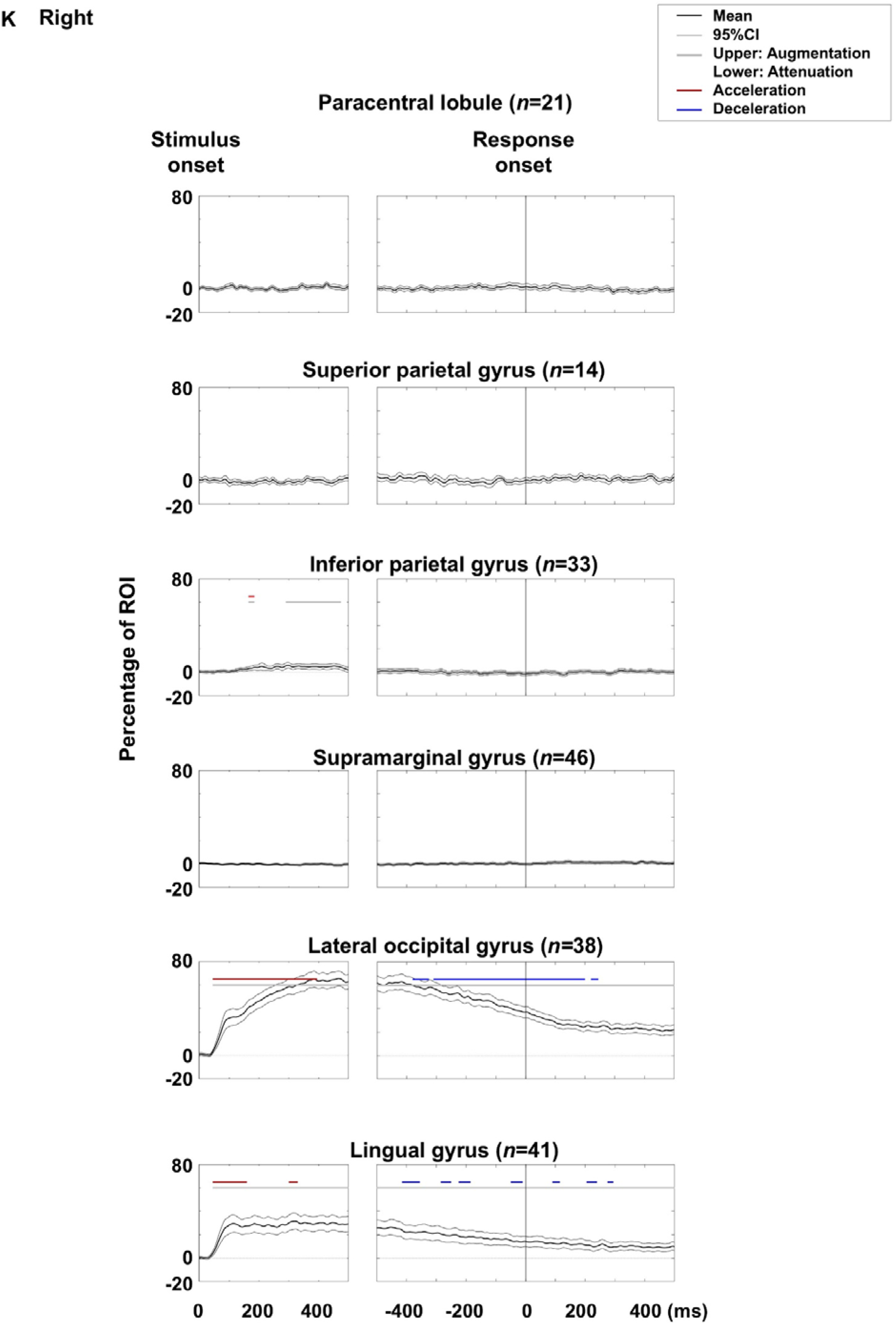
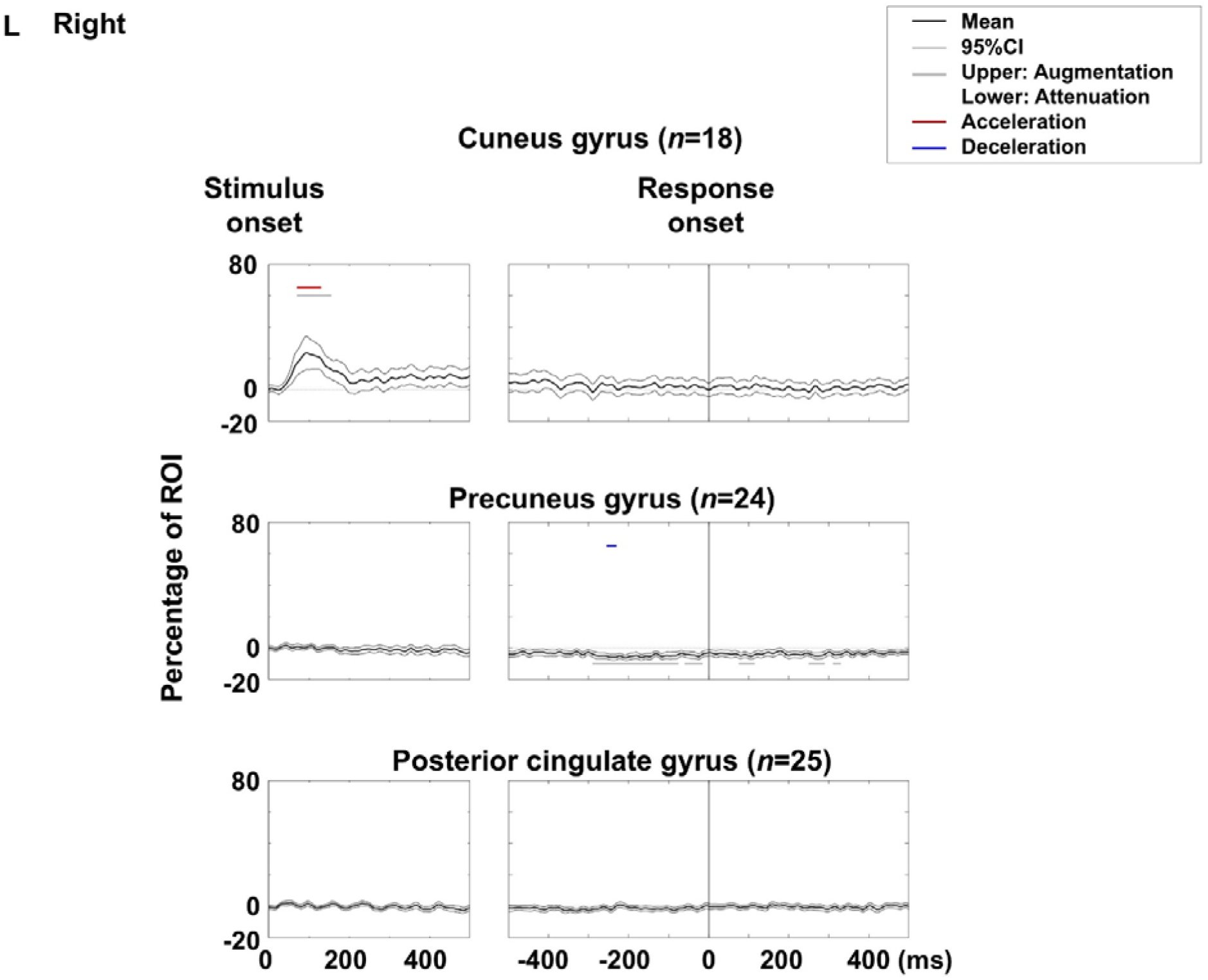
Visual naming-related high-gamma dynamics: distinguishing augmentation from acceleration and attenuation from deceleration. Mean high-gamma amplitude (% change relative to baseline) at each region of interest (ROI) is shown with 95% confidence intervals. Upper and lower horizontal bars denote time bins with significant high-gamma augmentation and attenuation, respectively, while red and blue bars indicate significant high-gamma acceleration and deceleration. **A–F** Left hemispheric ROIs. **G–L** Right hemispheric ROIs.

**eFigure 5.**
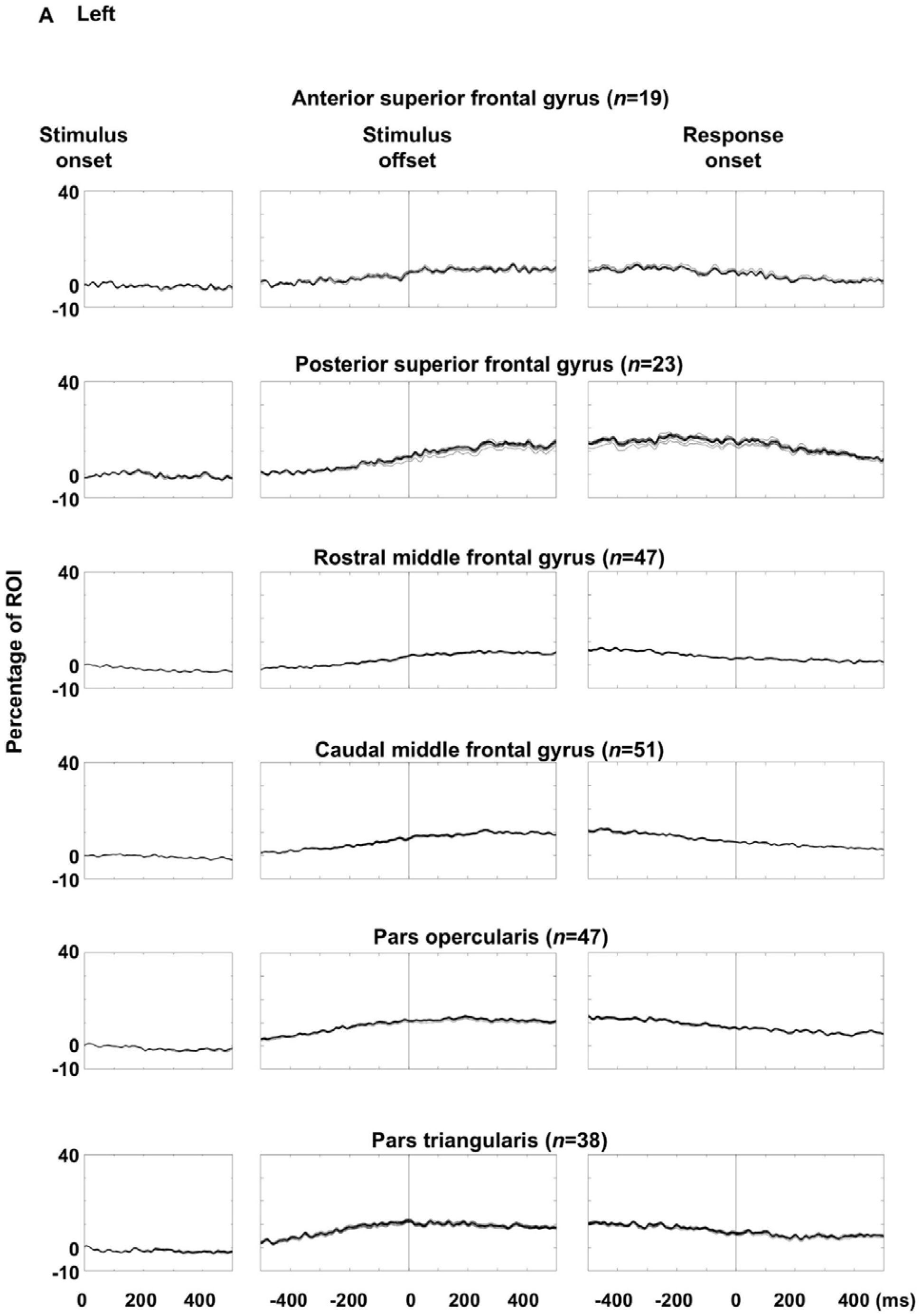
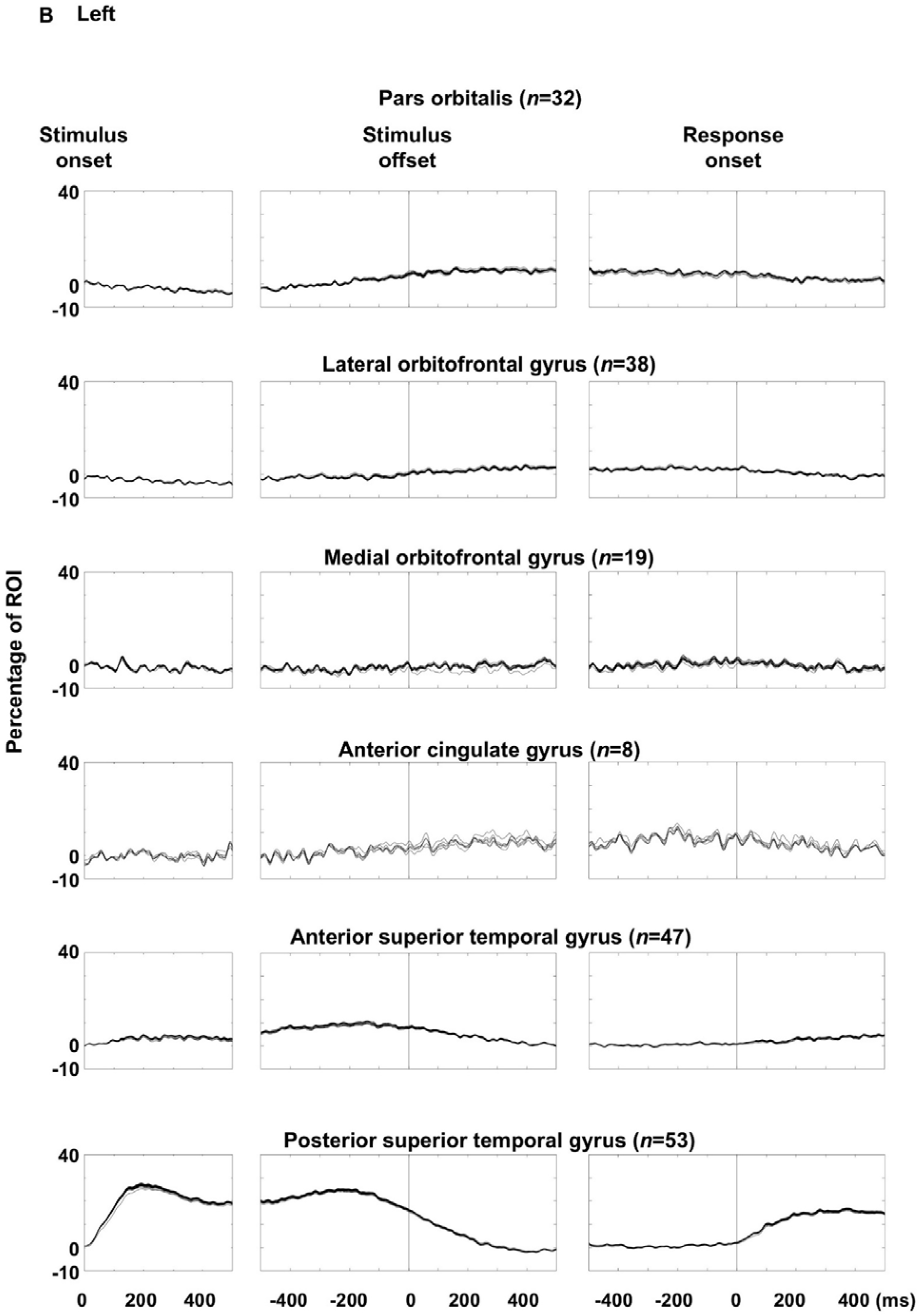
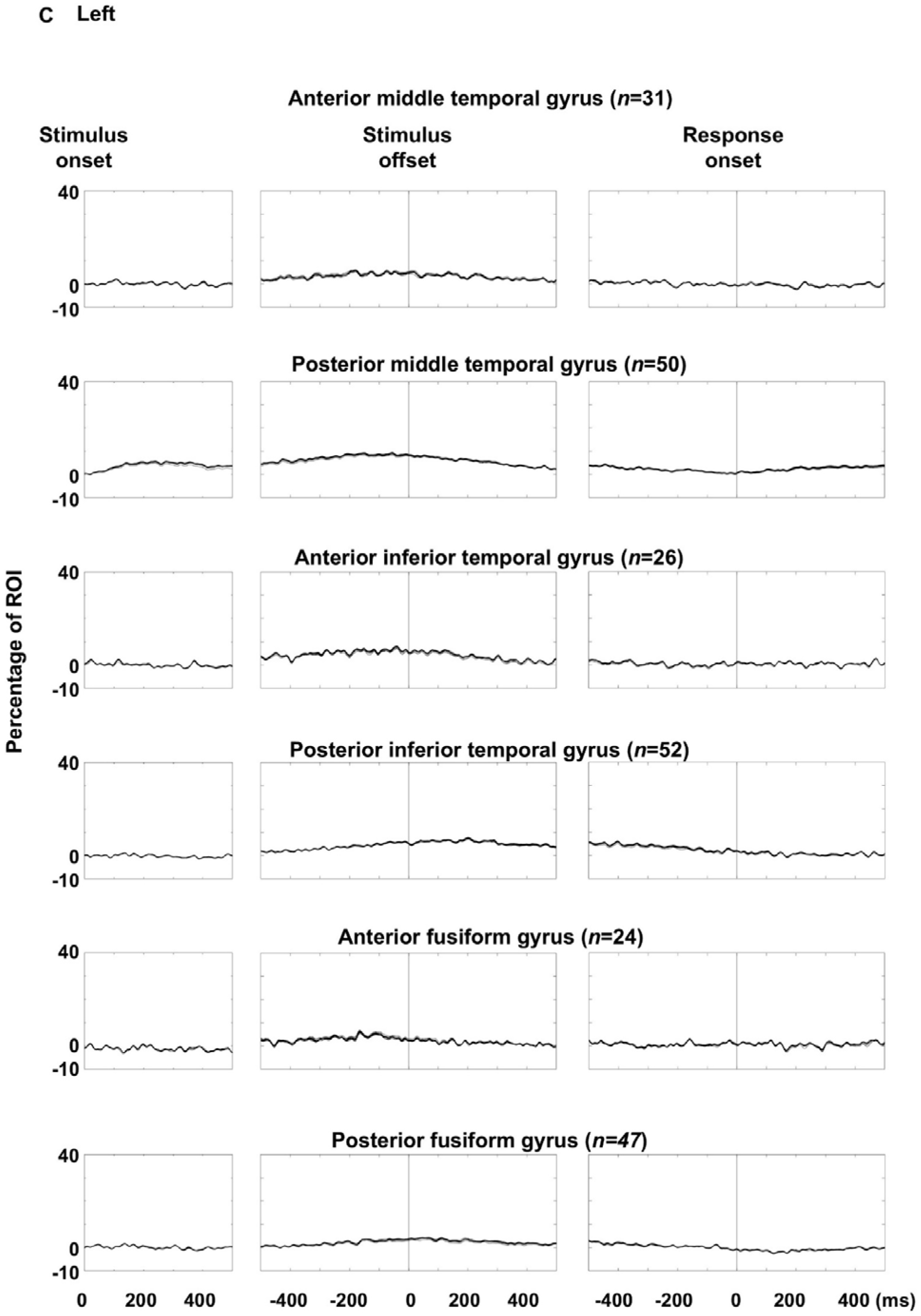
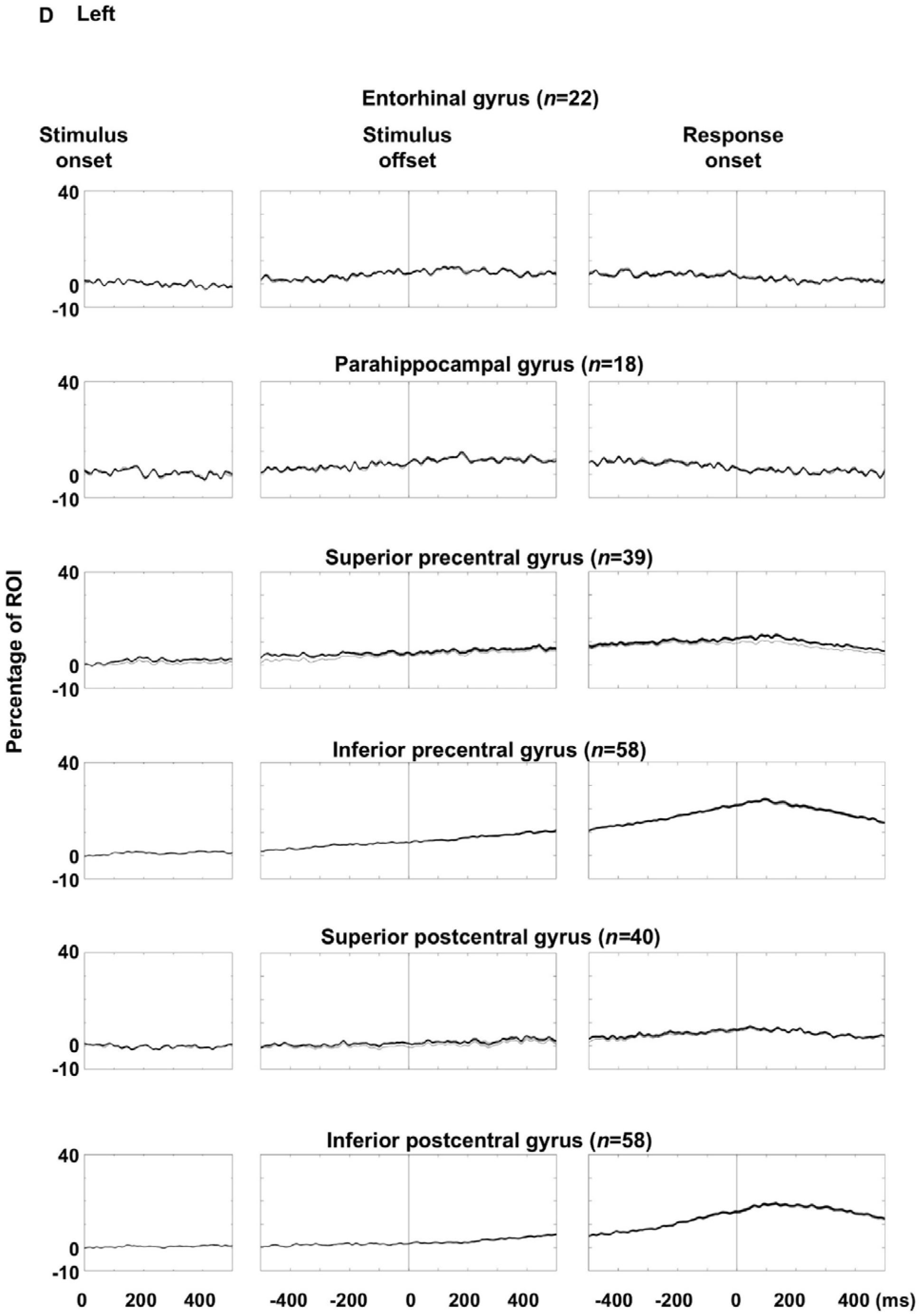
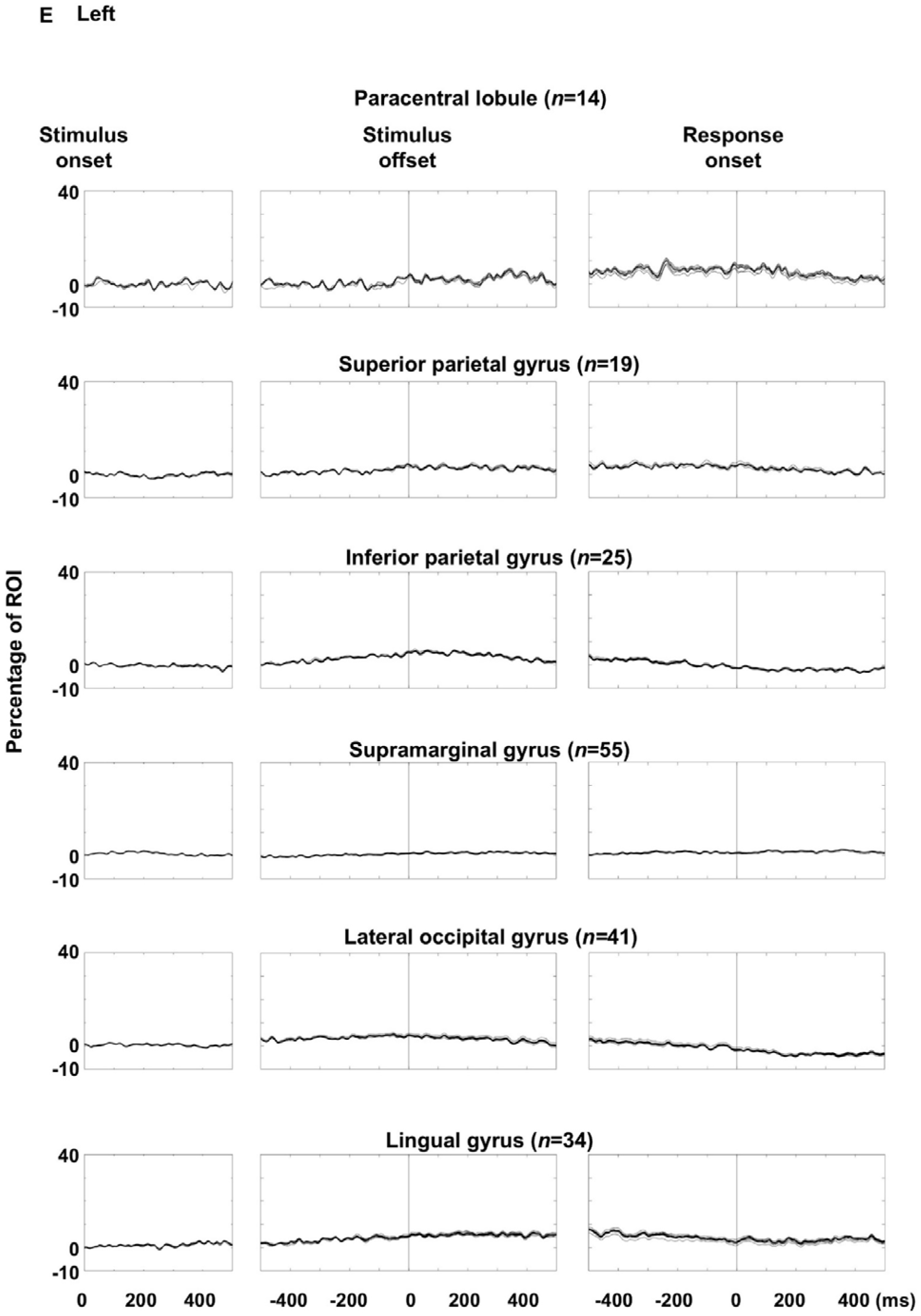
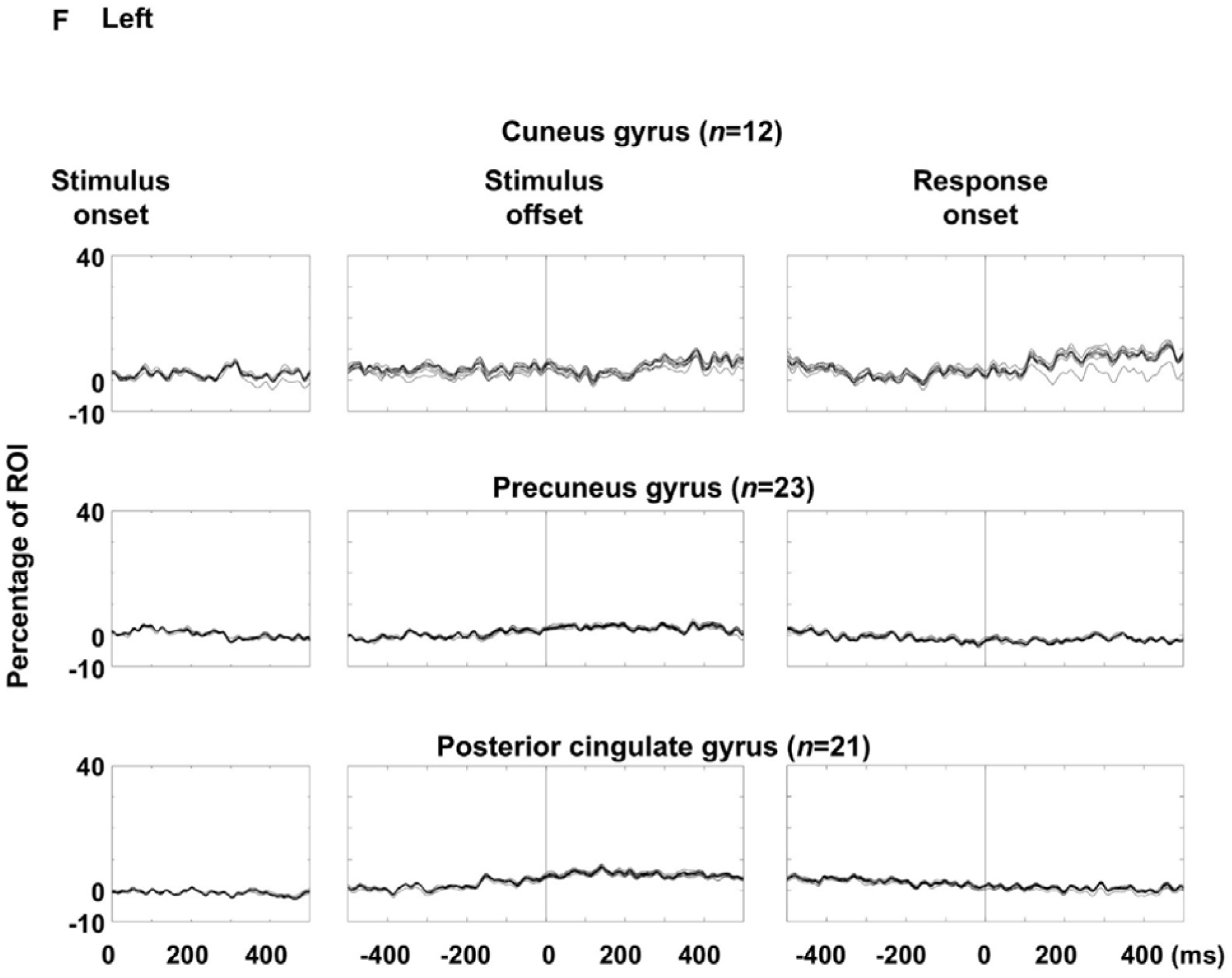
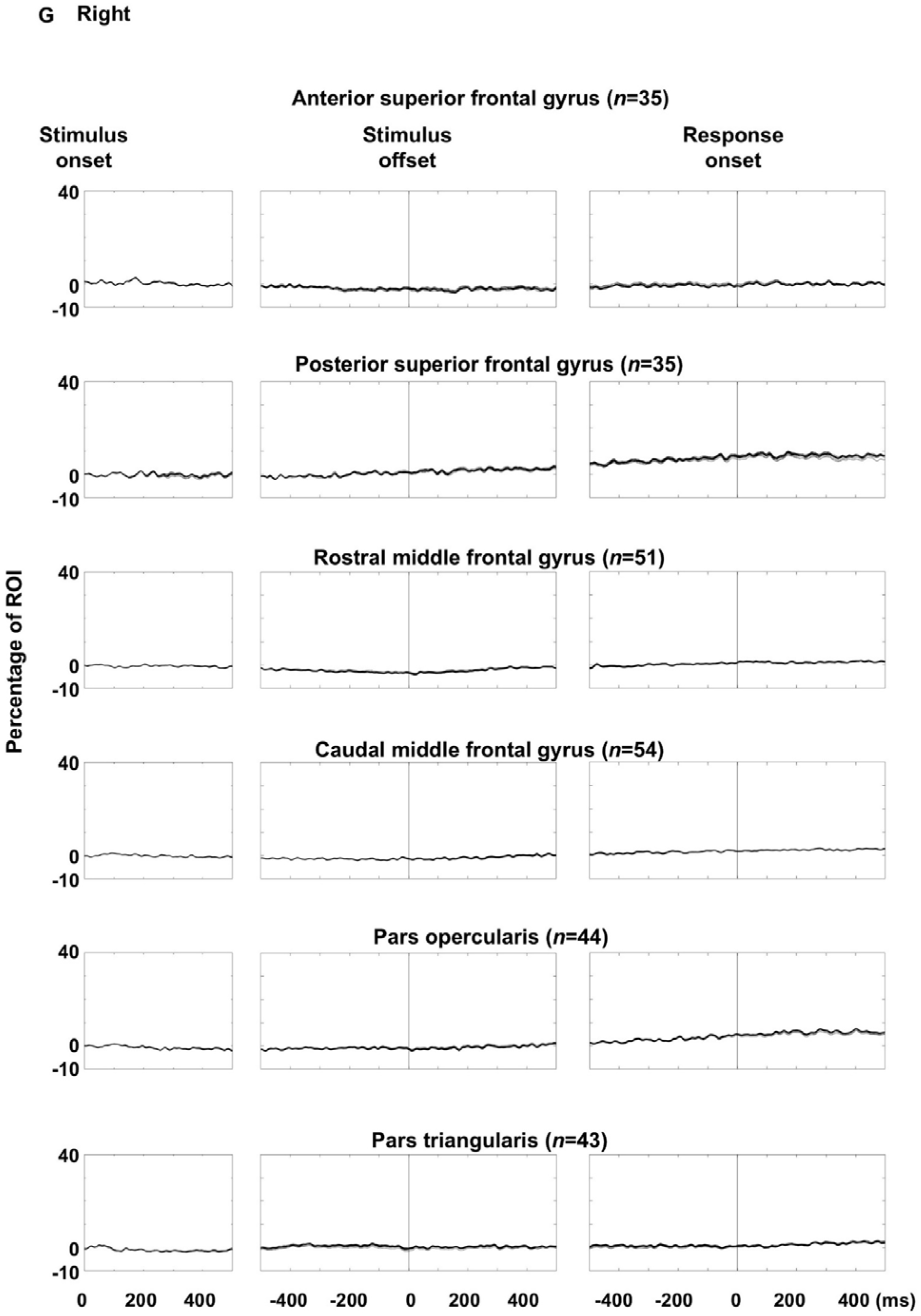
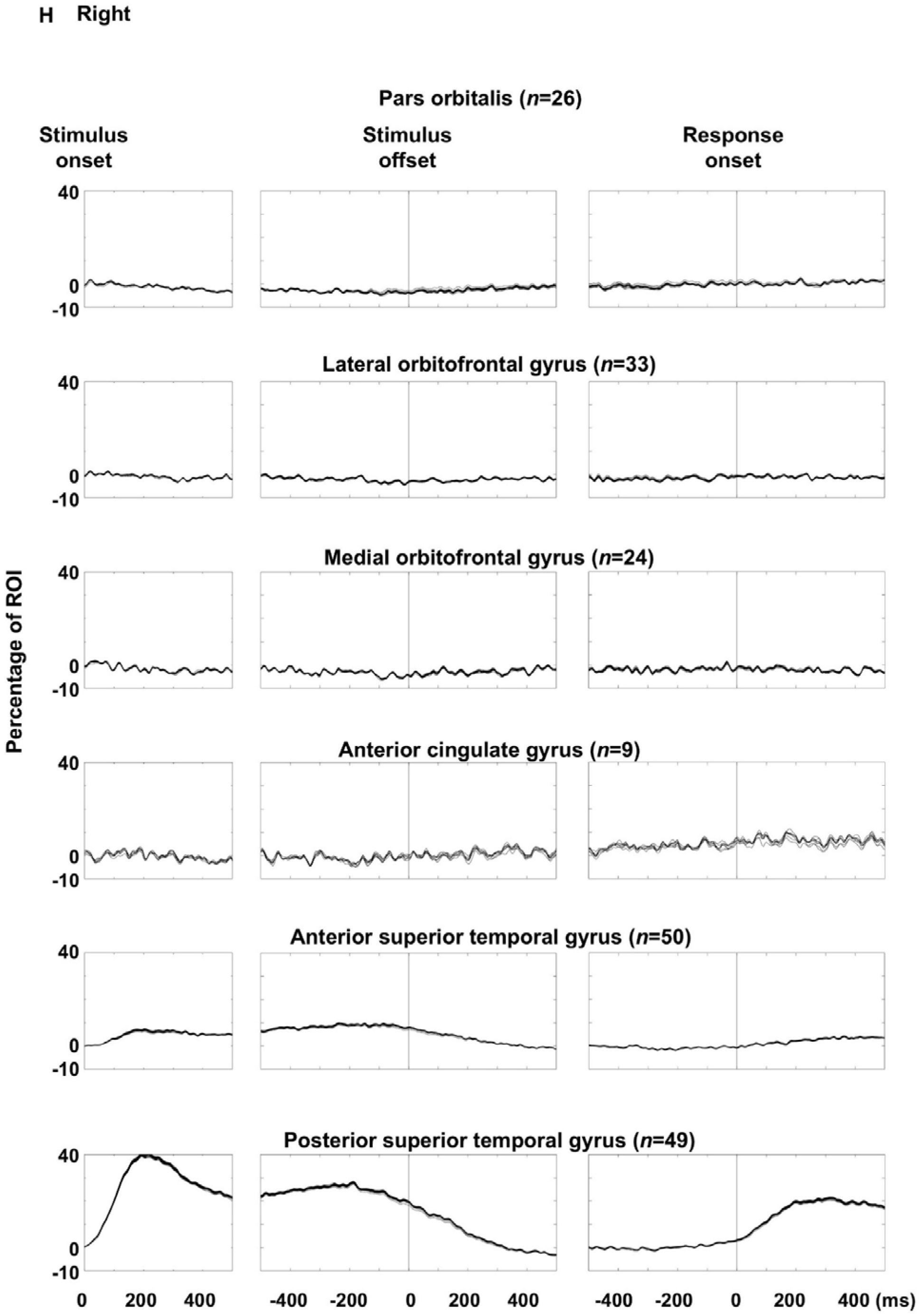
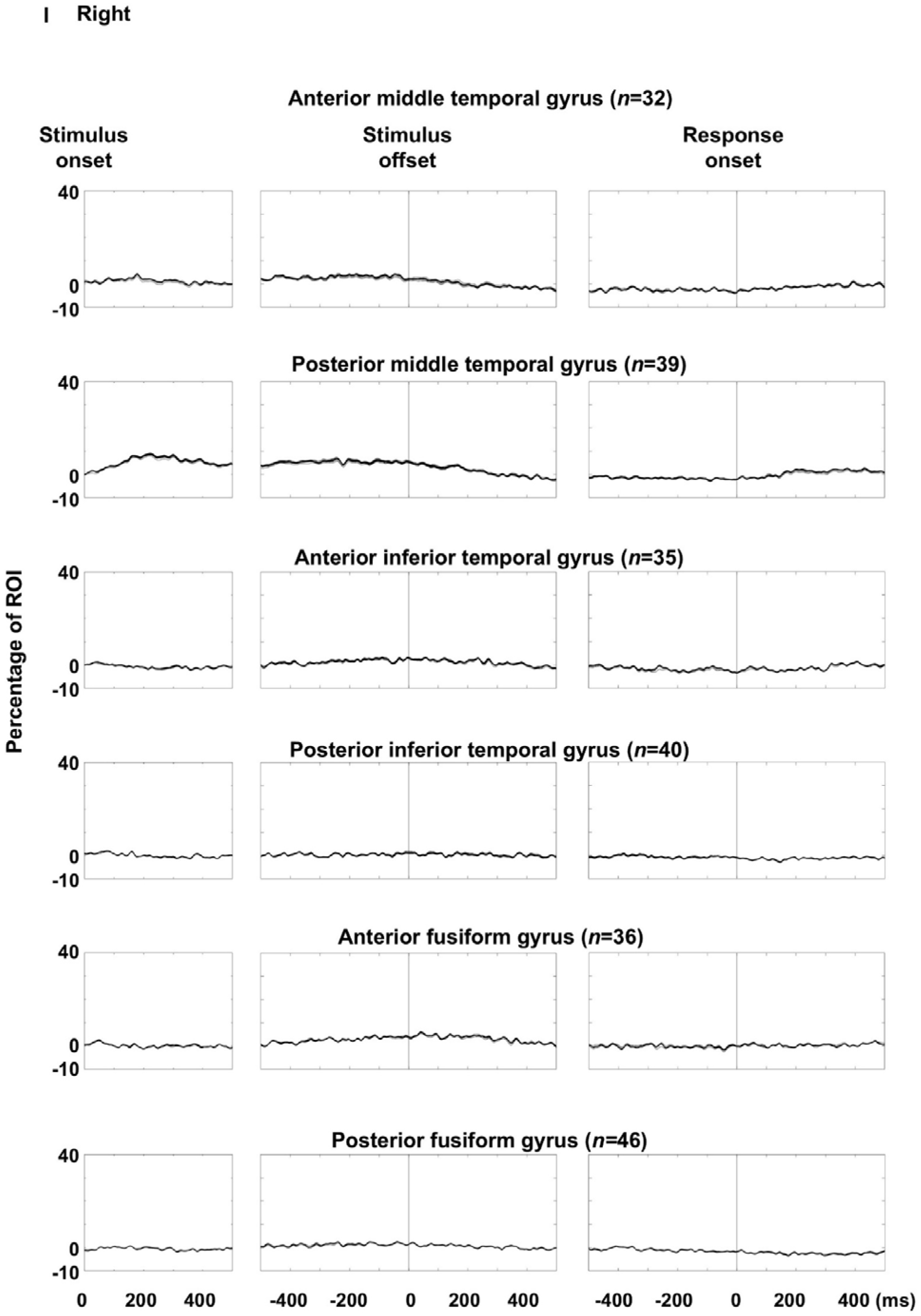
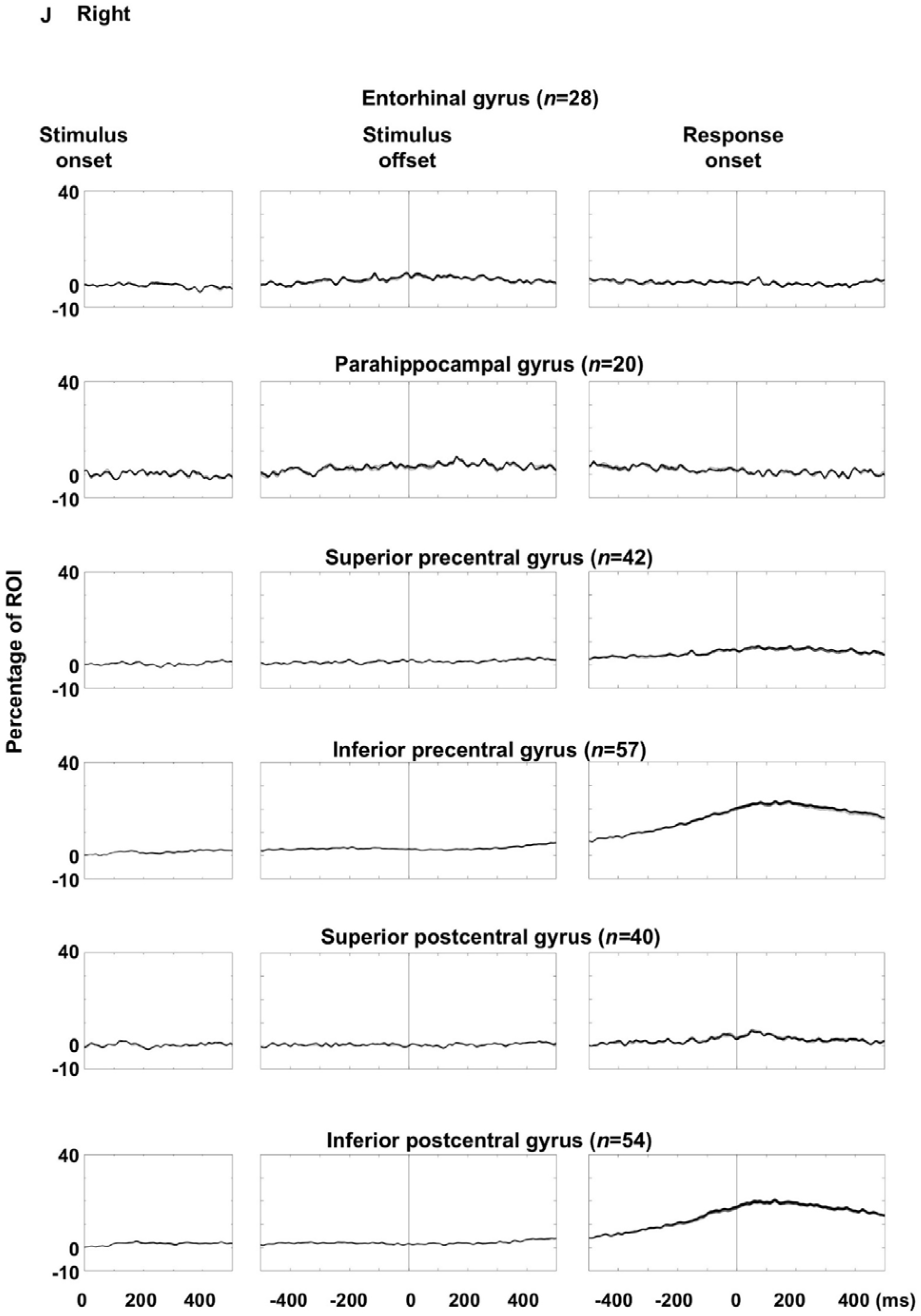
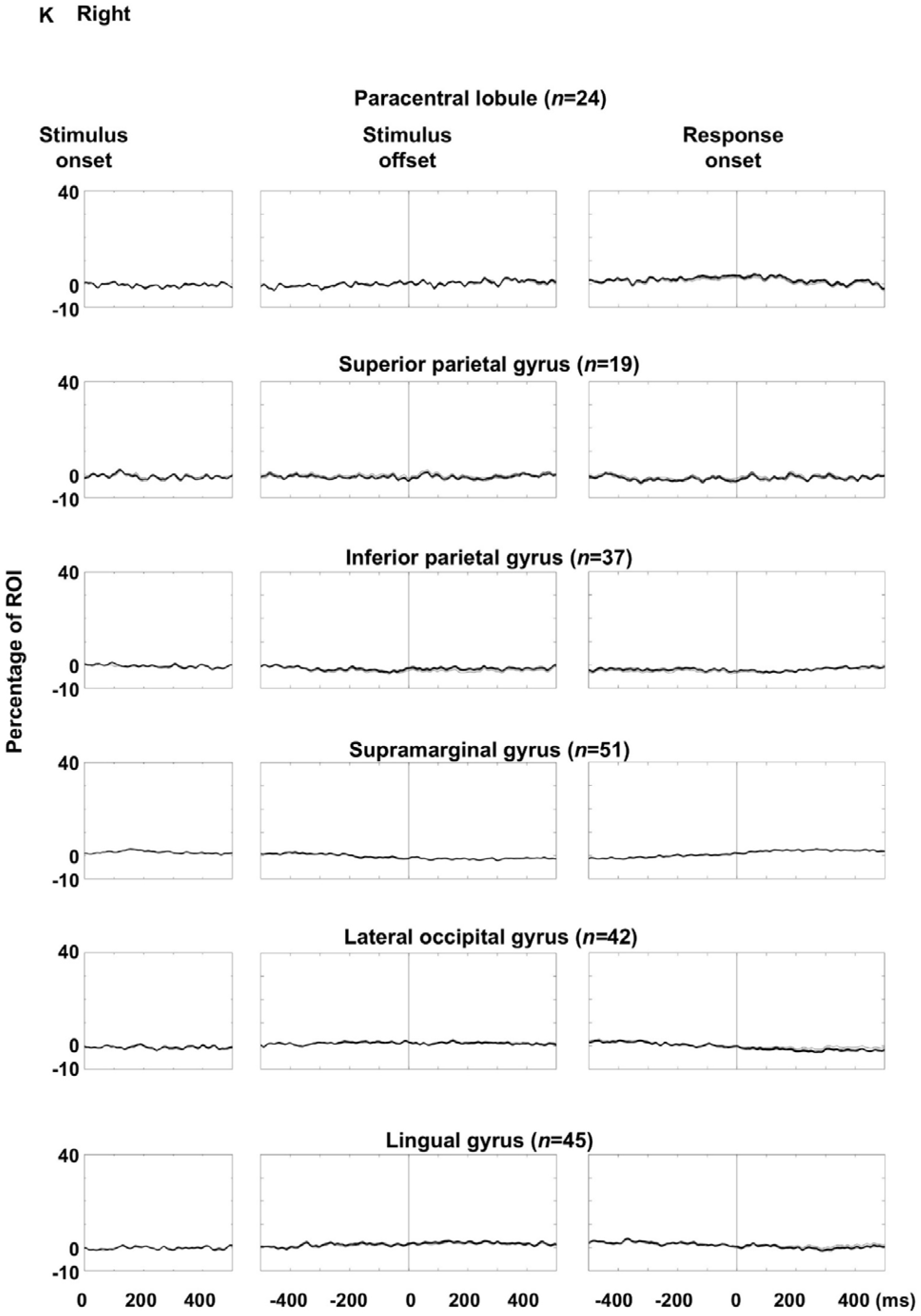
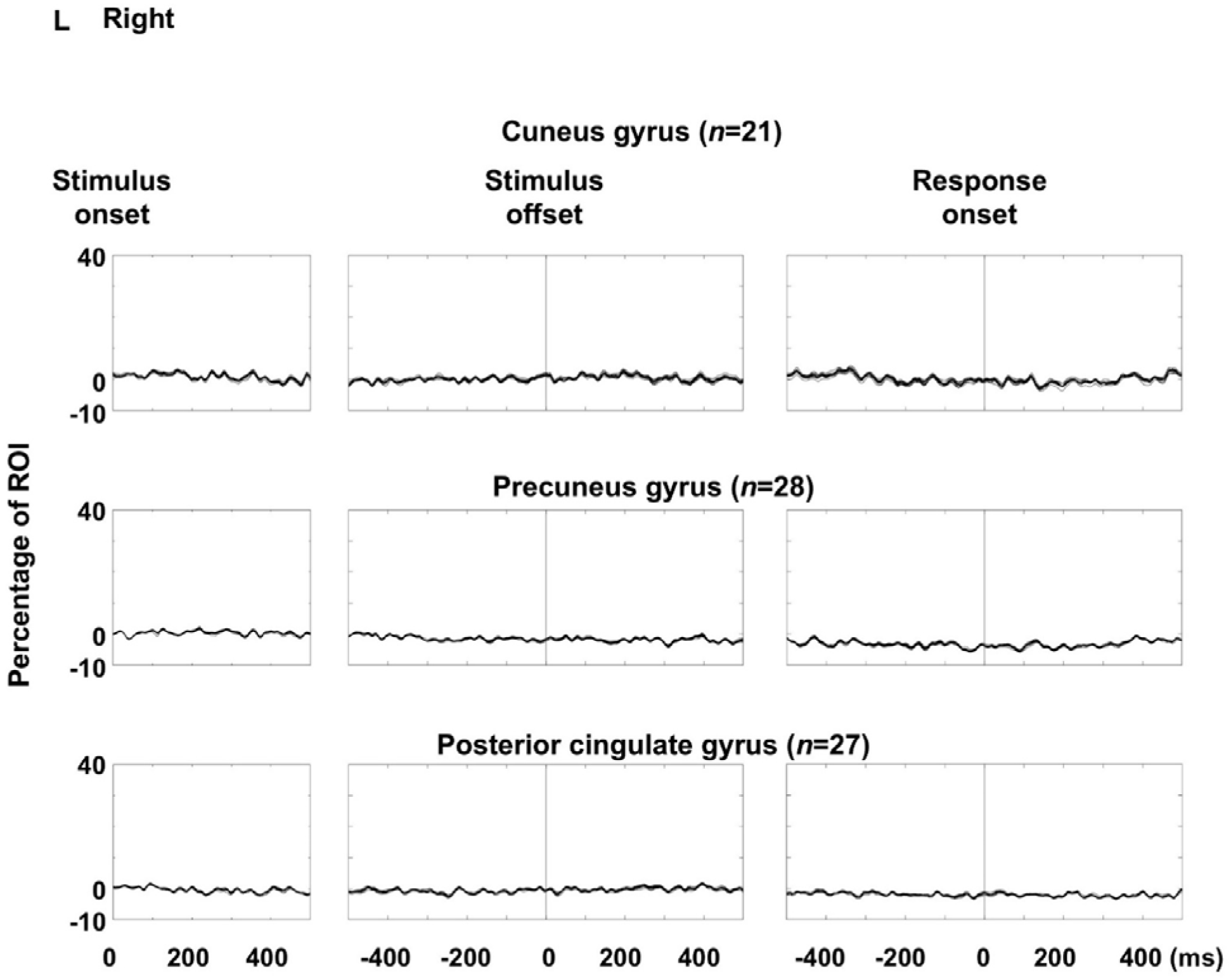
Auditory naming-related high-gamma amplitude modulations: leave-one-patient-out analysis. Each plot shows the dynamics of high-gamma amplitude at each region of interest (ROI; % change relative to baseline) averaged across all patients except one excluded patient. **A–F** Left hemispheric ROIs. **G–L** Right hemispheric ROIs. To confirm that high-gamma amplitude modulations were robust and not disproportionately influenced by individual patients, we applied a leave-one-patient-out approach, computing Spearman’s correlation coefficient (rho) repeatedly while excluding each patient in turn. Across the 66 ROIs, the mean rho values ranged from 0.95 to 0.99. One-sample t-tests demonstrated that mean rho values were significantly greater than zero at all ROIs (all p < 0.00001).

**eFigure 6.**
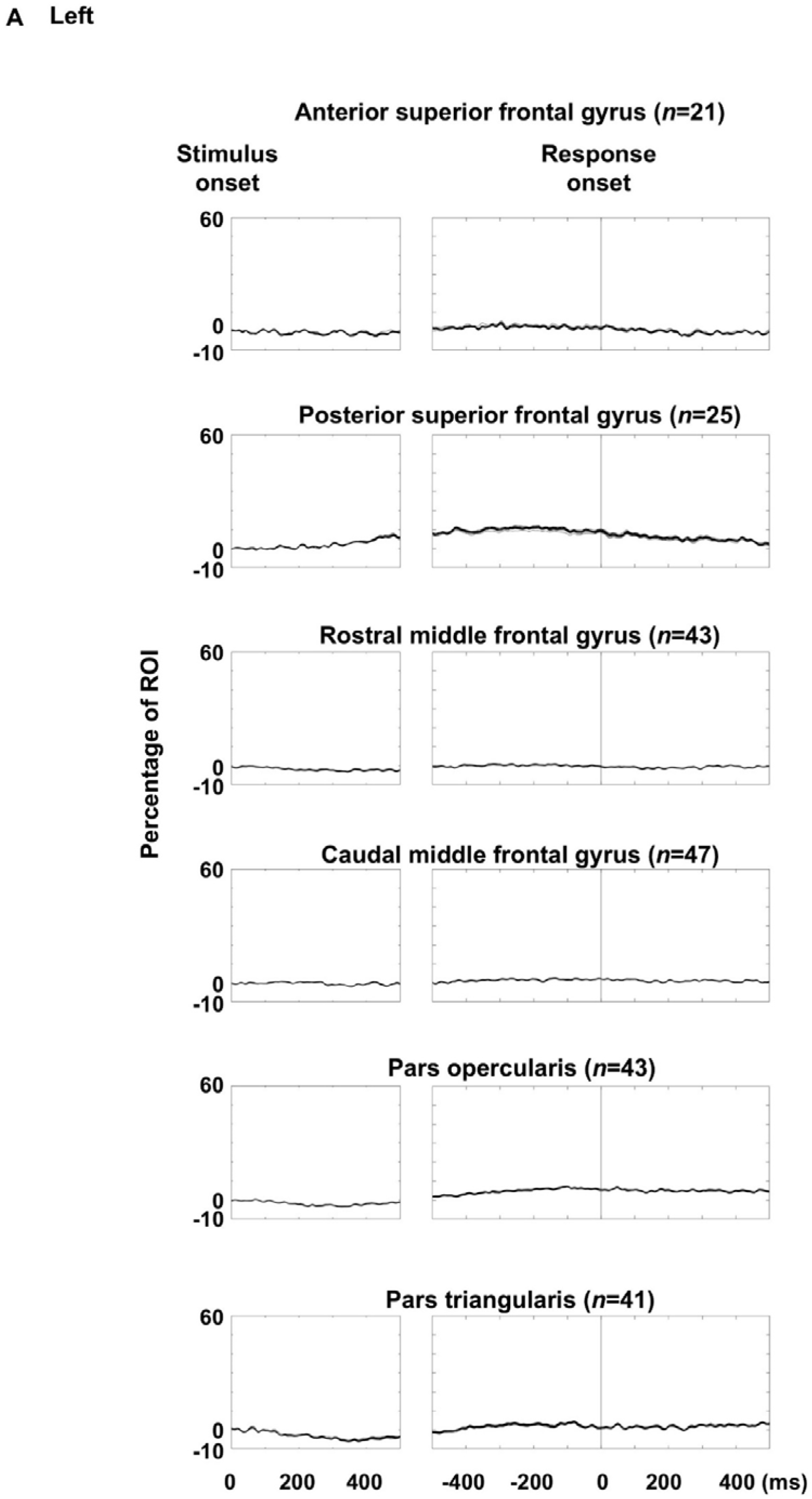
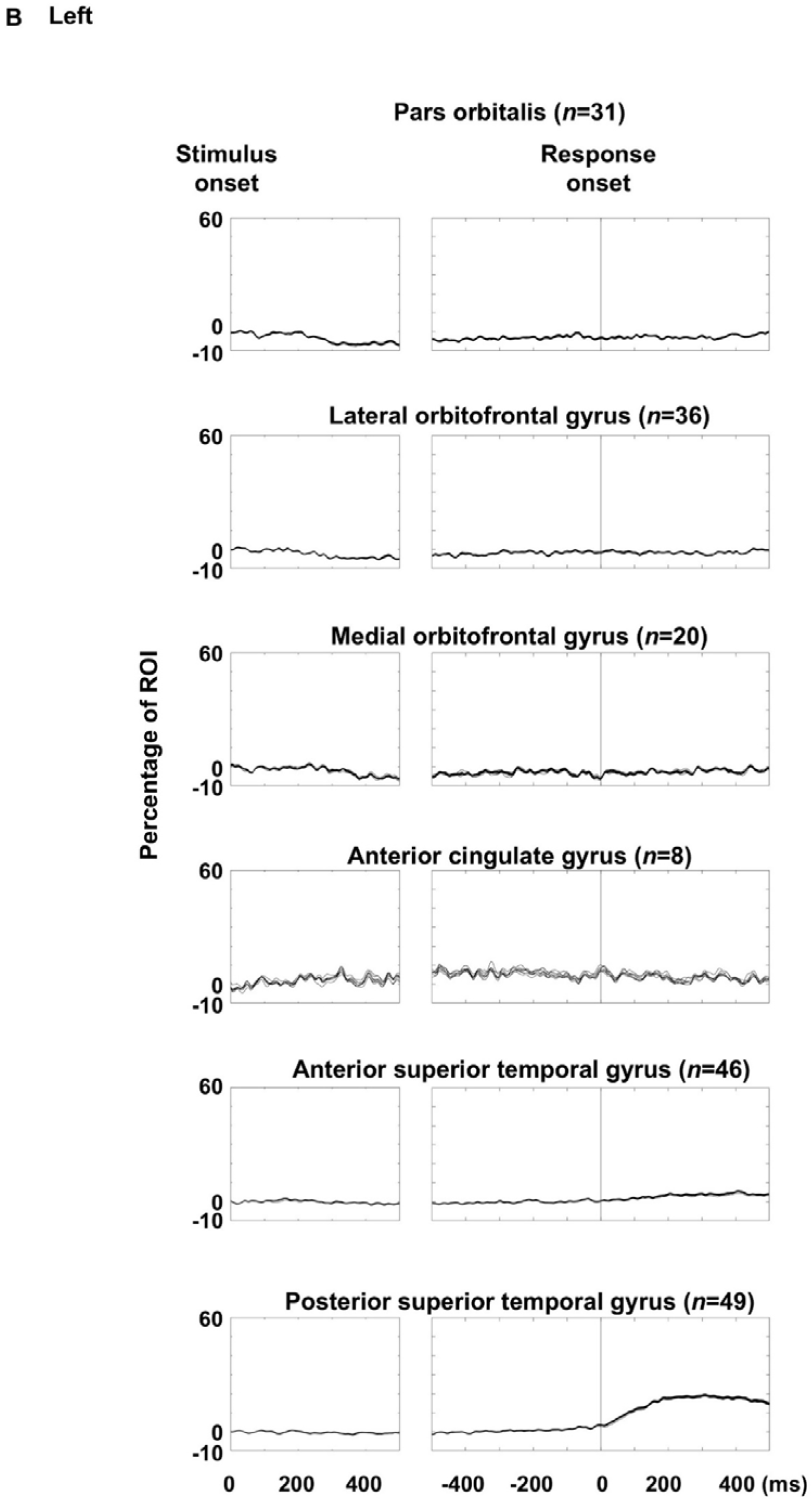
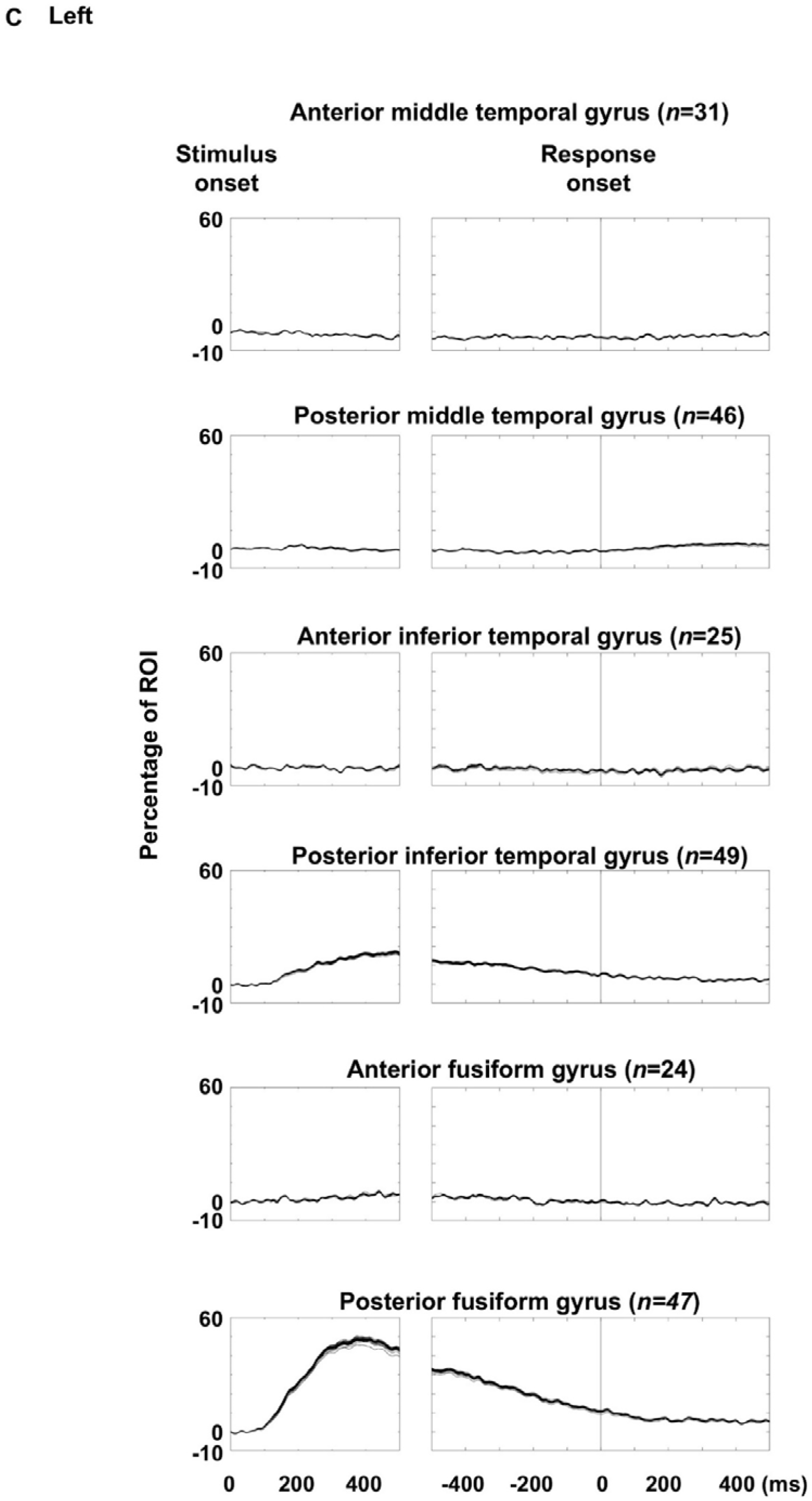
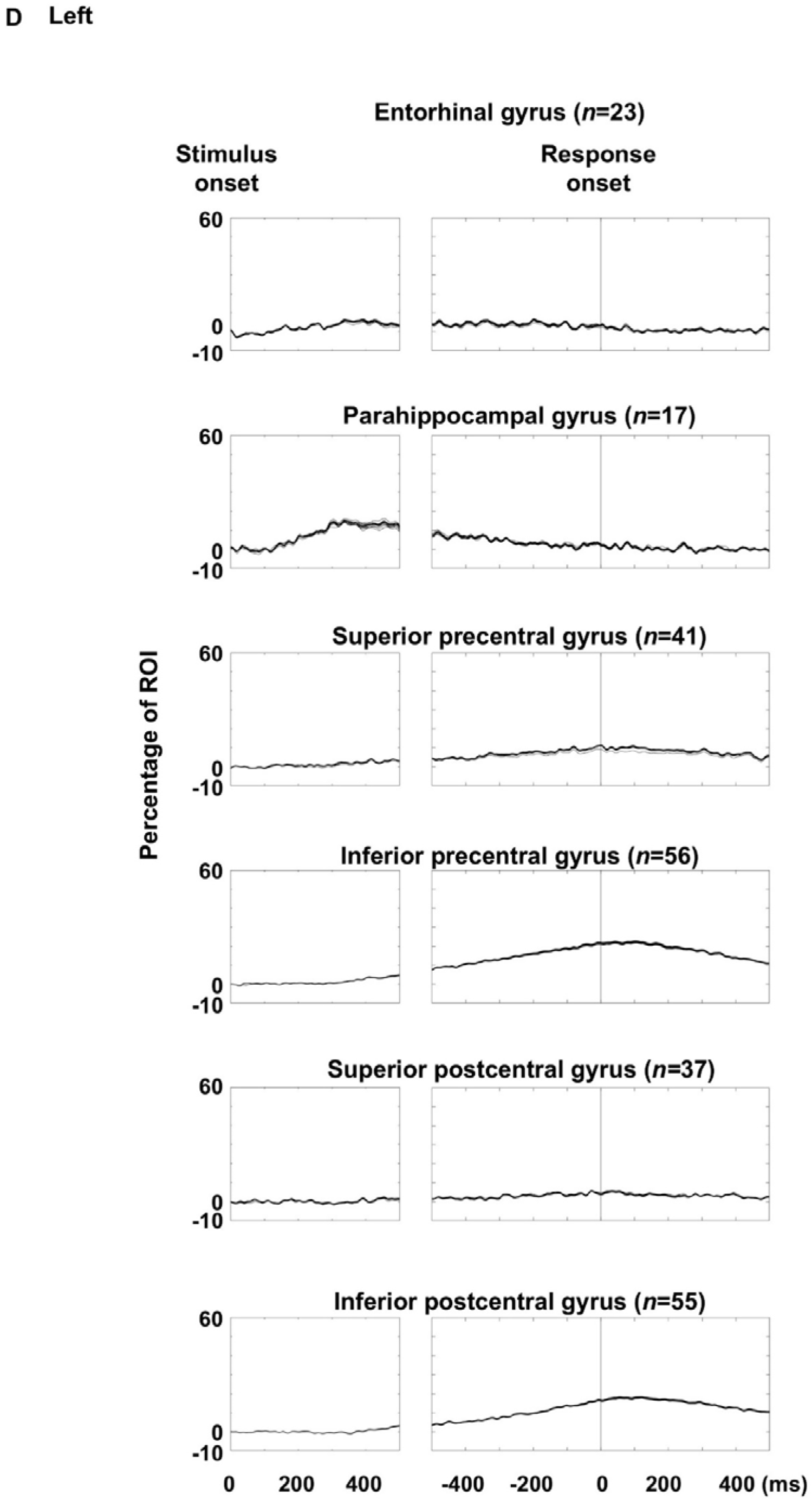
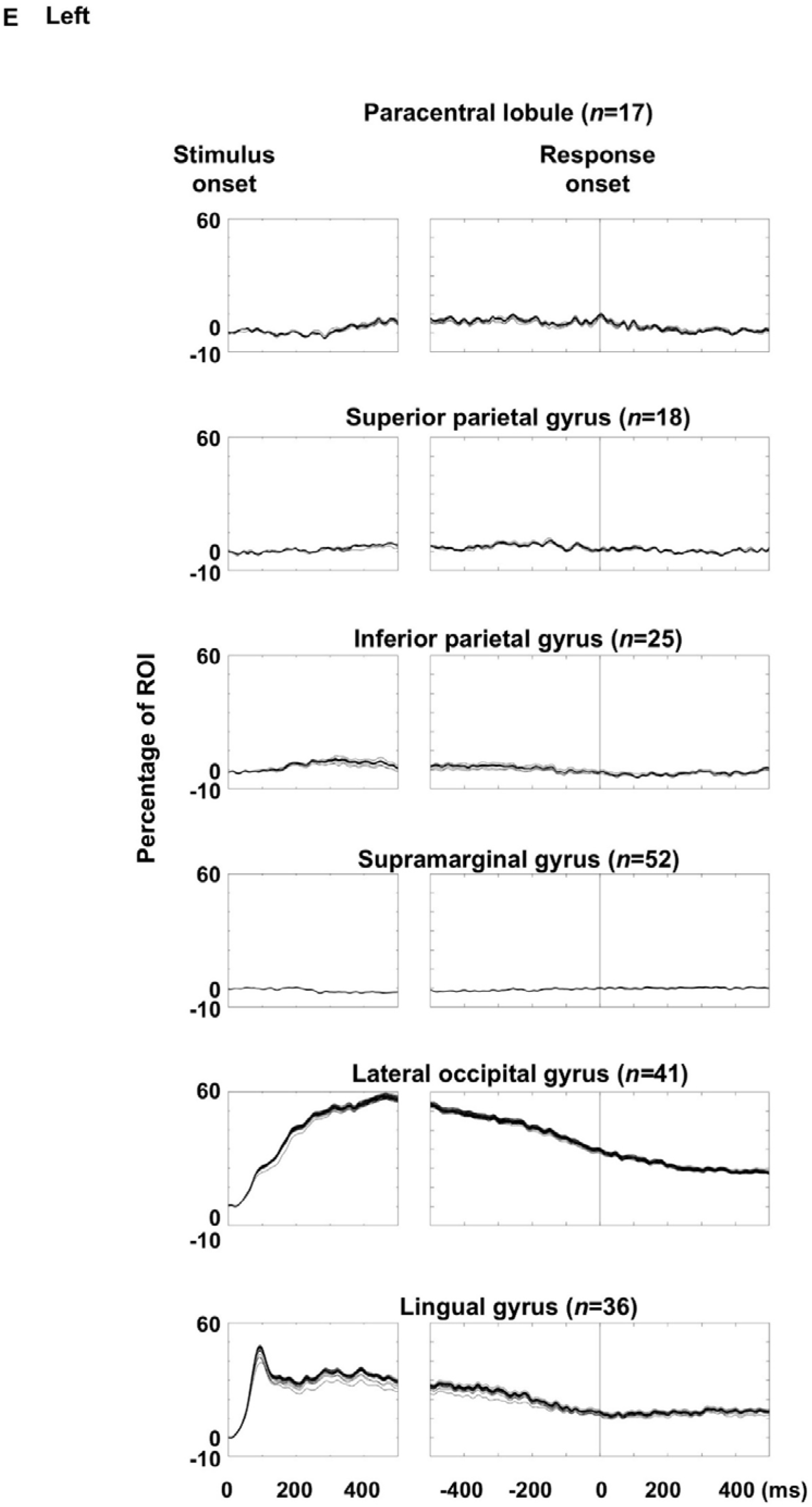
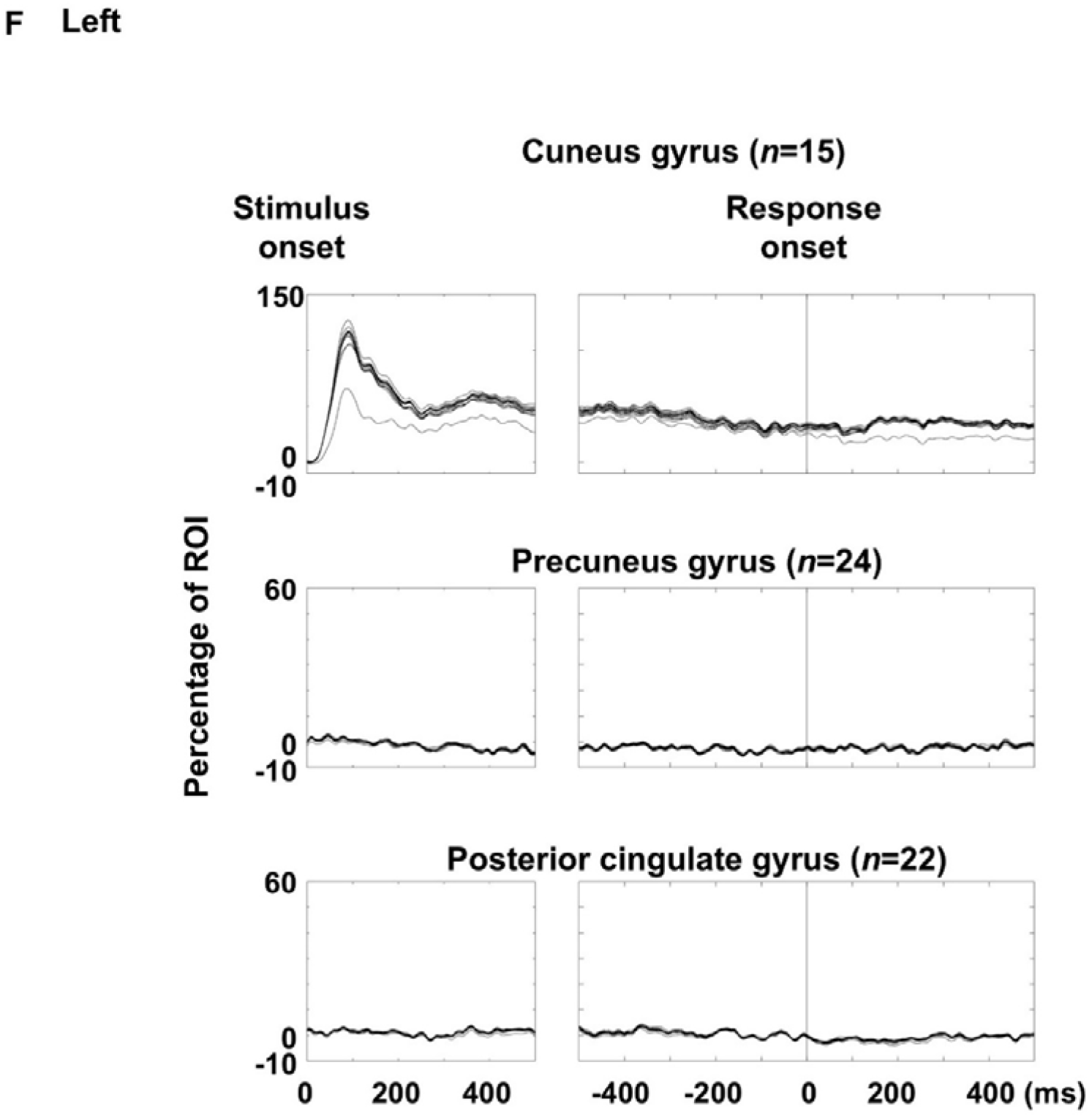
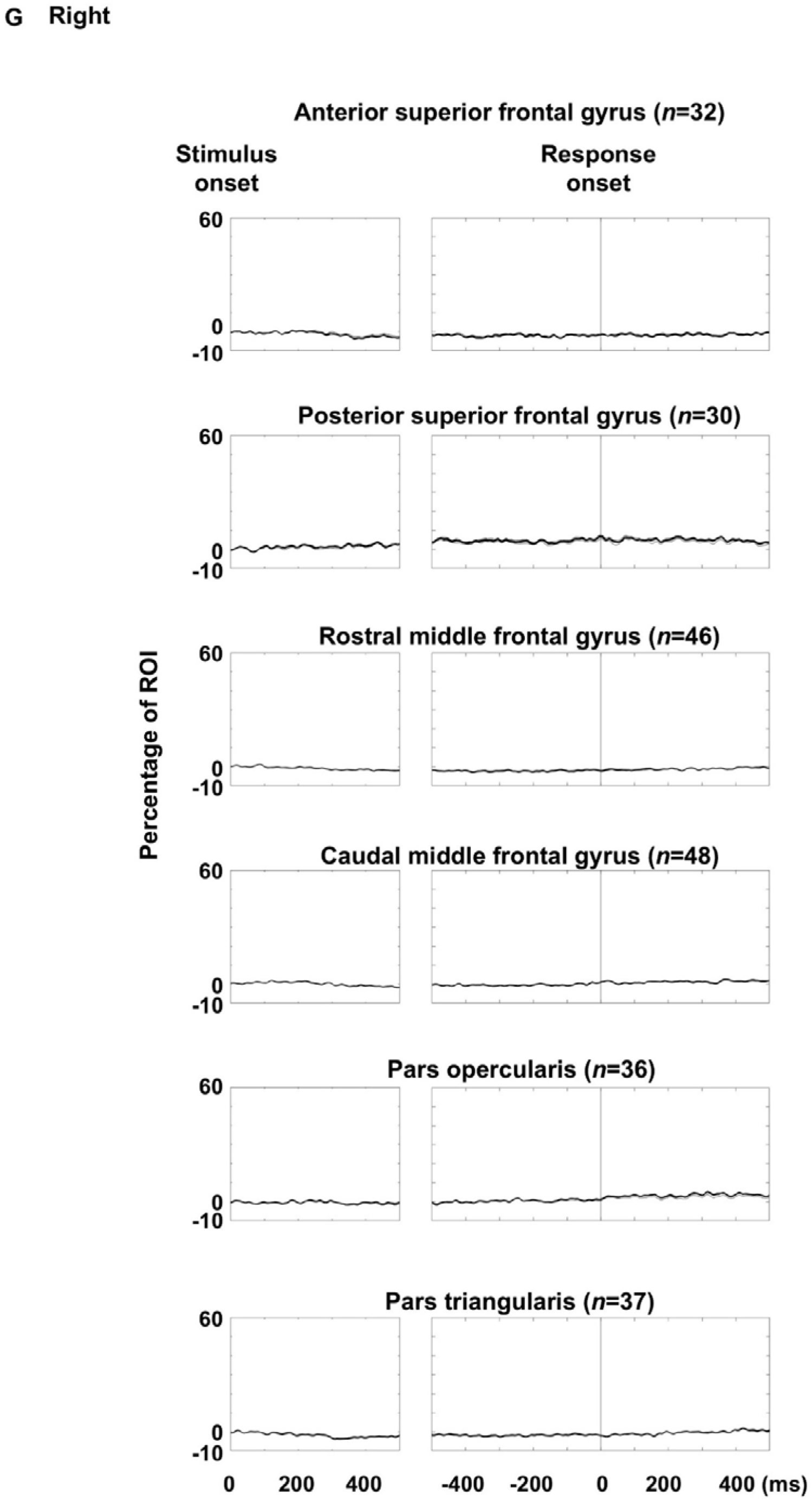
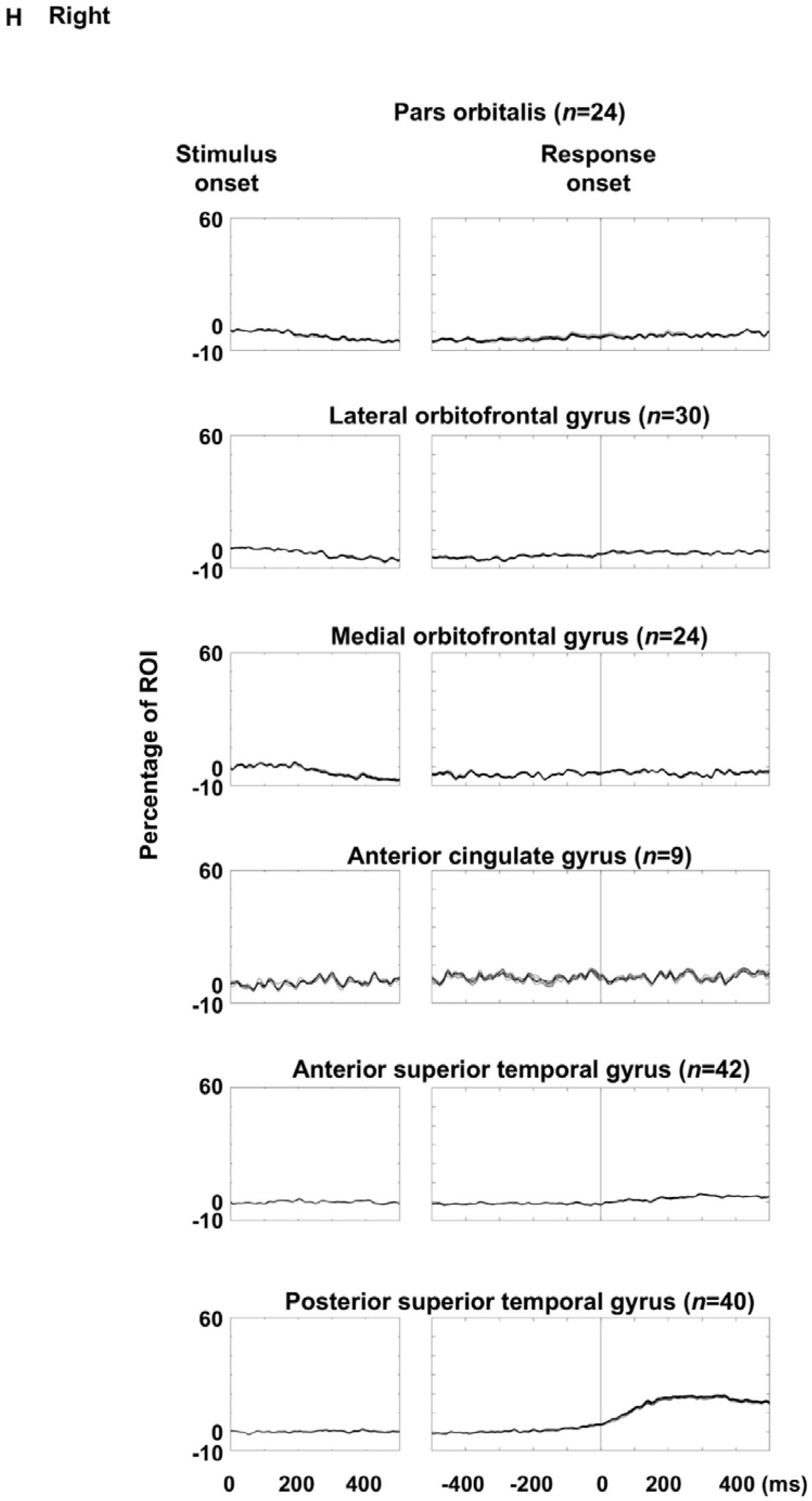
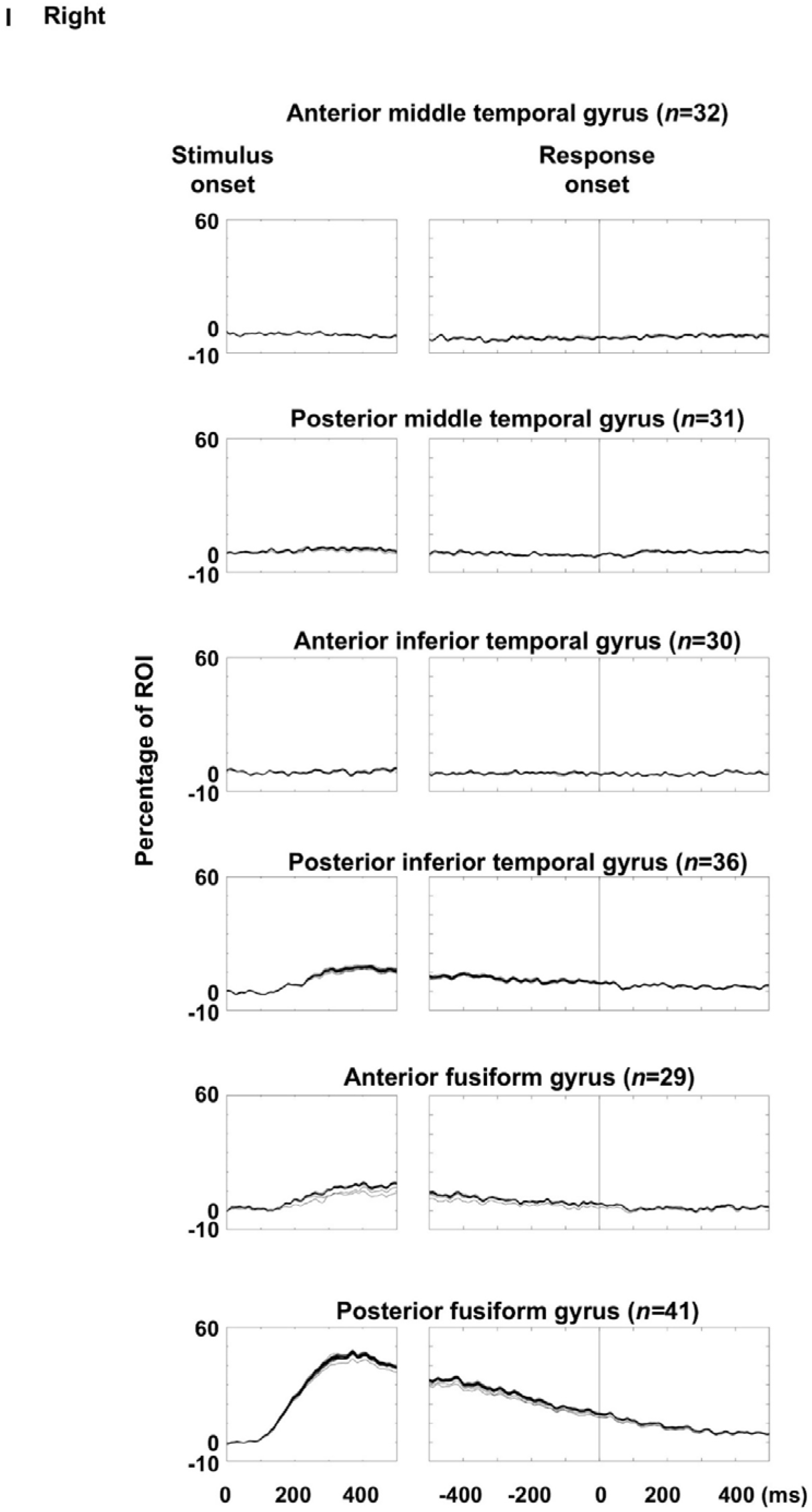
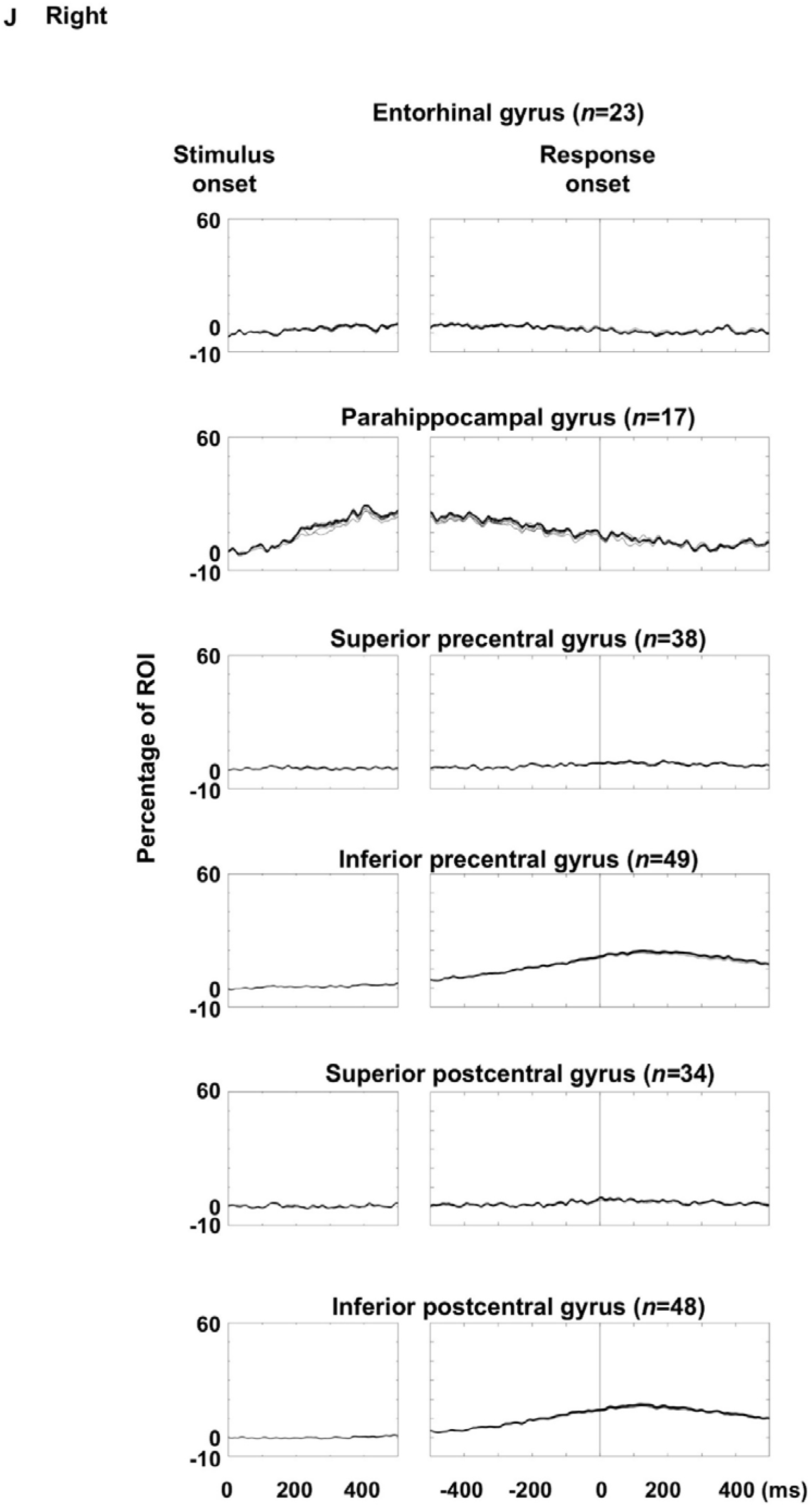
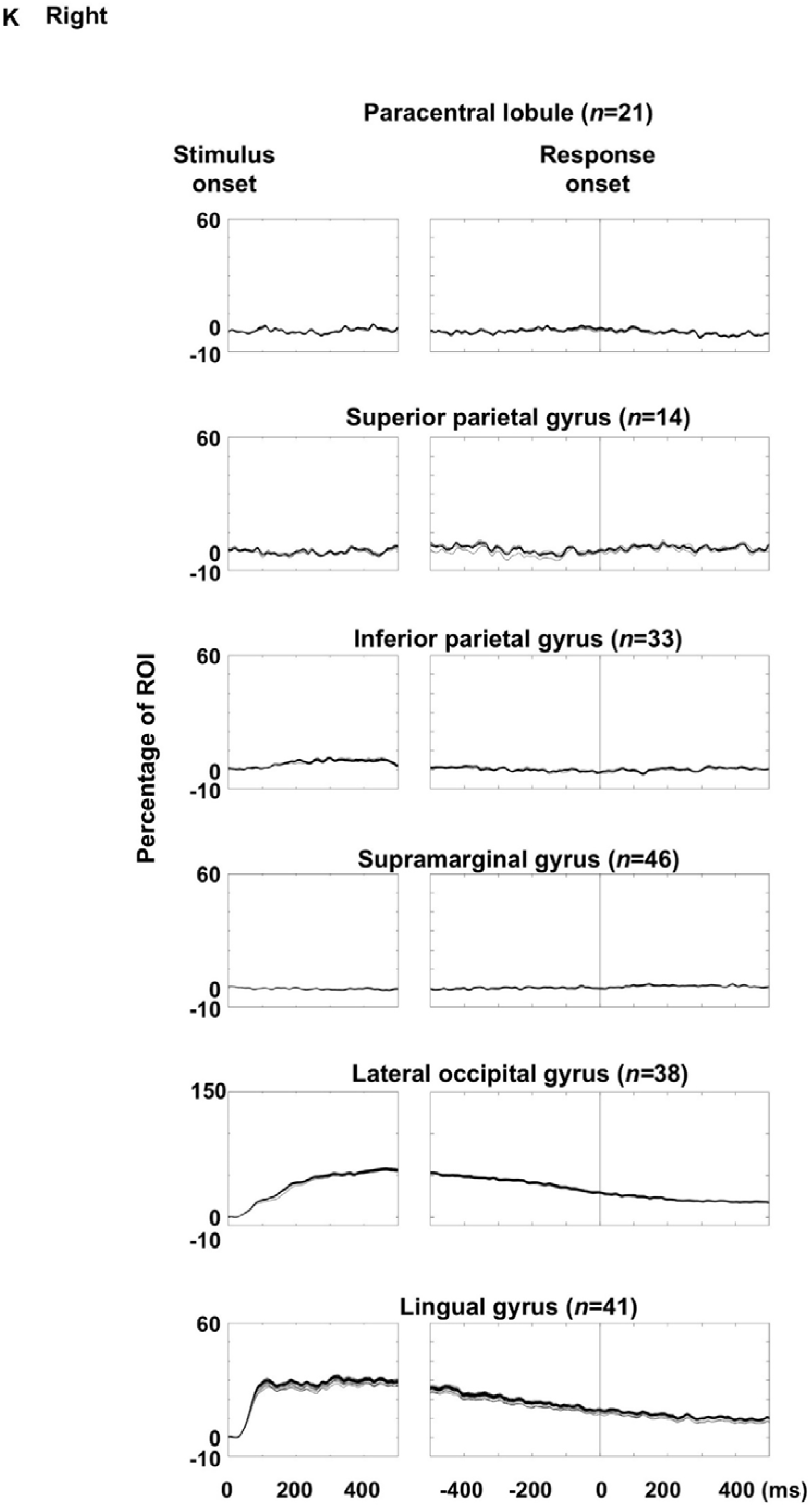
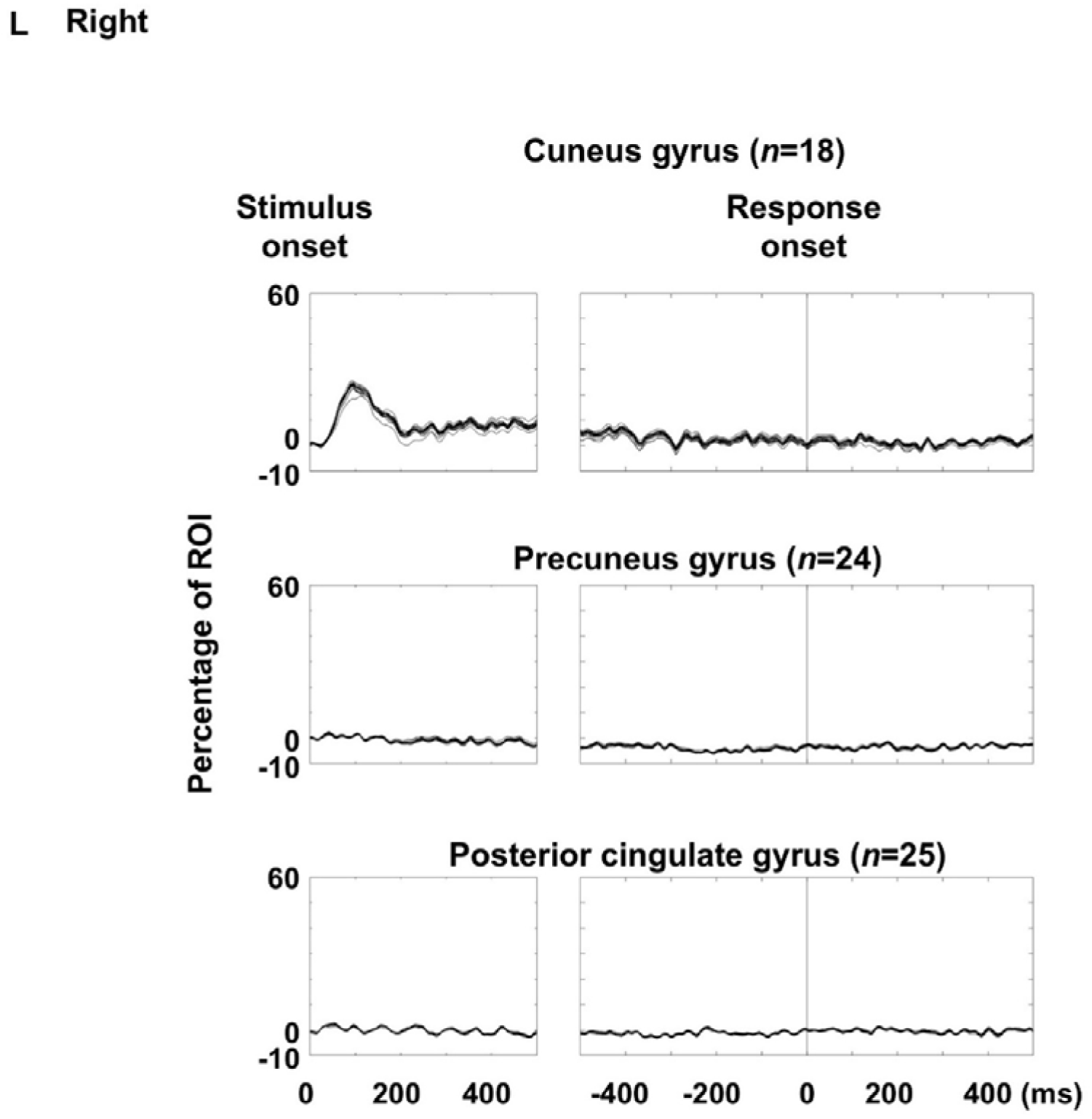
Visual naming-related high-gamma amplitude modulations: leave-one-patient-out analysis. Each plot shows the dynamics of high-gamma amplitude at each region of interest (ROI; % change relative to baseline) averaged across all patients except one excluded patient. **A–F** Left hemispheric ROIs. **G–L** Right hemispheric ROIs. To confirm that high-gamma amplitude modulations were robust and not disproportionately influenced by individual patients, we applied a leave-one-patient-out approach, computing Spearman’s correlation coefficient (rho) repeatedly while excluding each patient in turn. Across the 66 ROIs, the mean rho values ranged from 0.96 to 0.99. One-sample t-tests demonstrated that mean rho values were significantly greater than zero at all ROIs (all p < 0.00001).

**eFigure 7.**
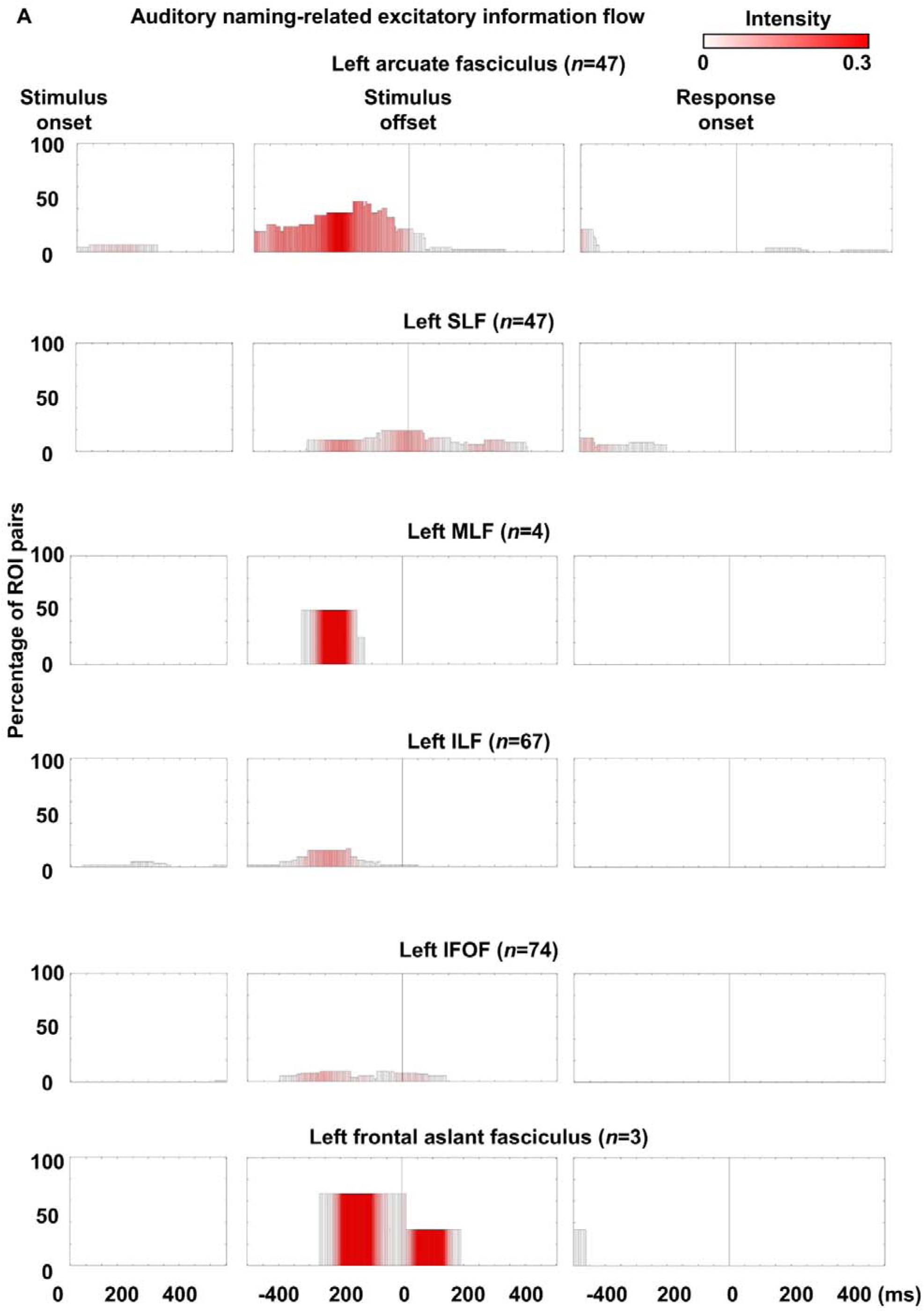
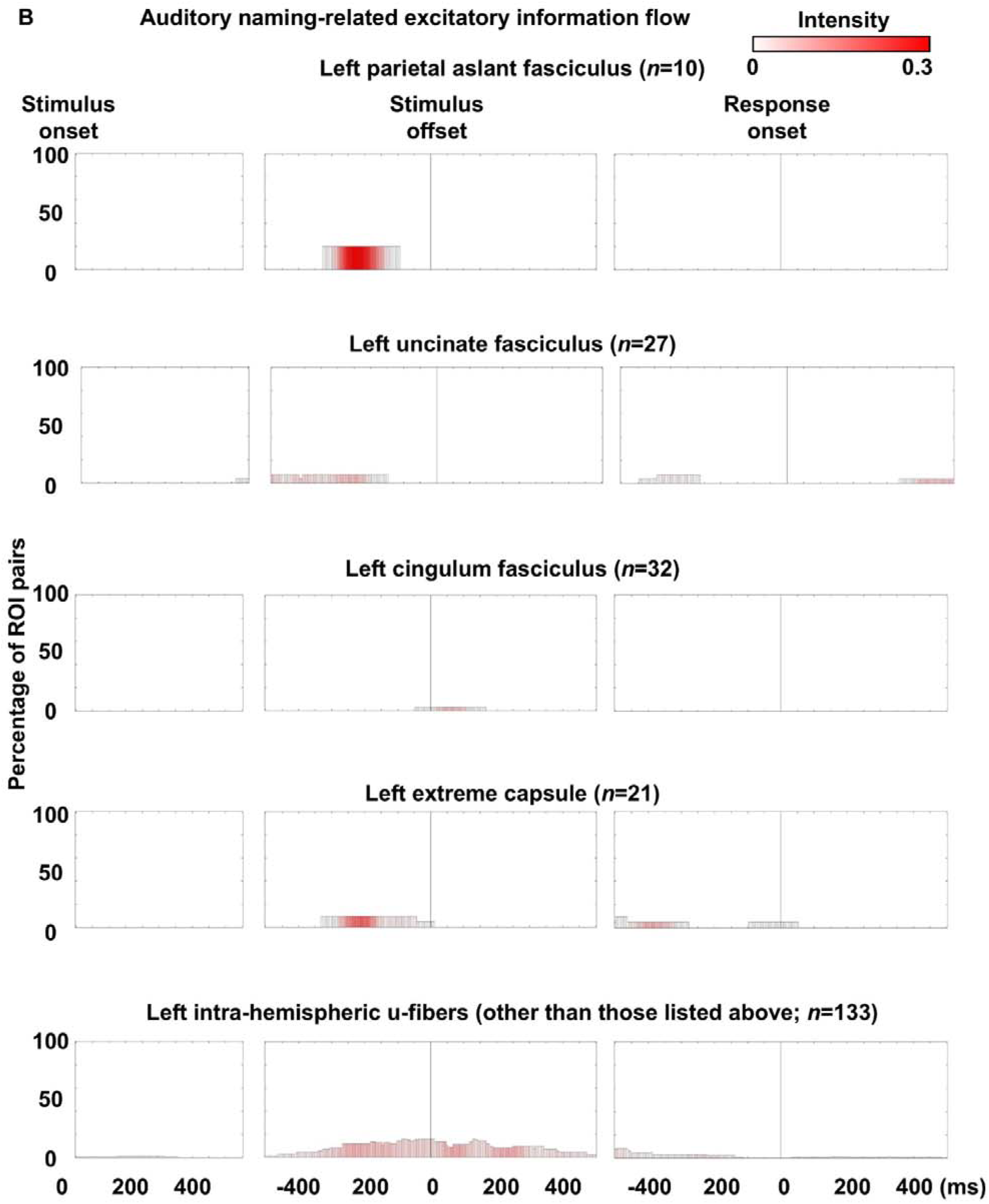
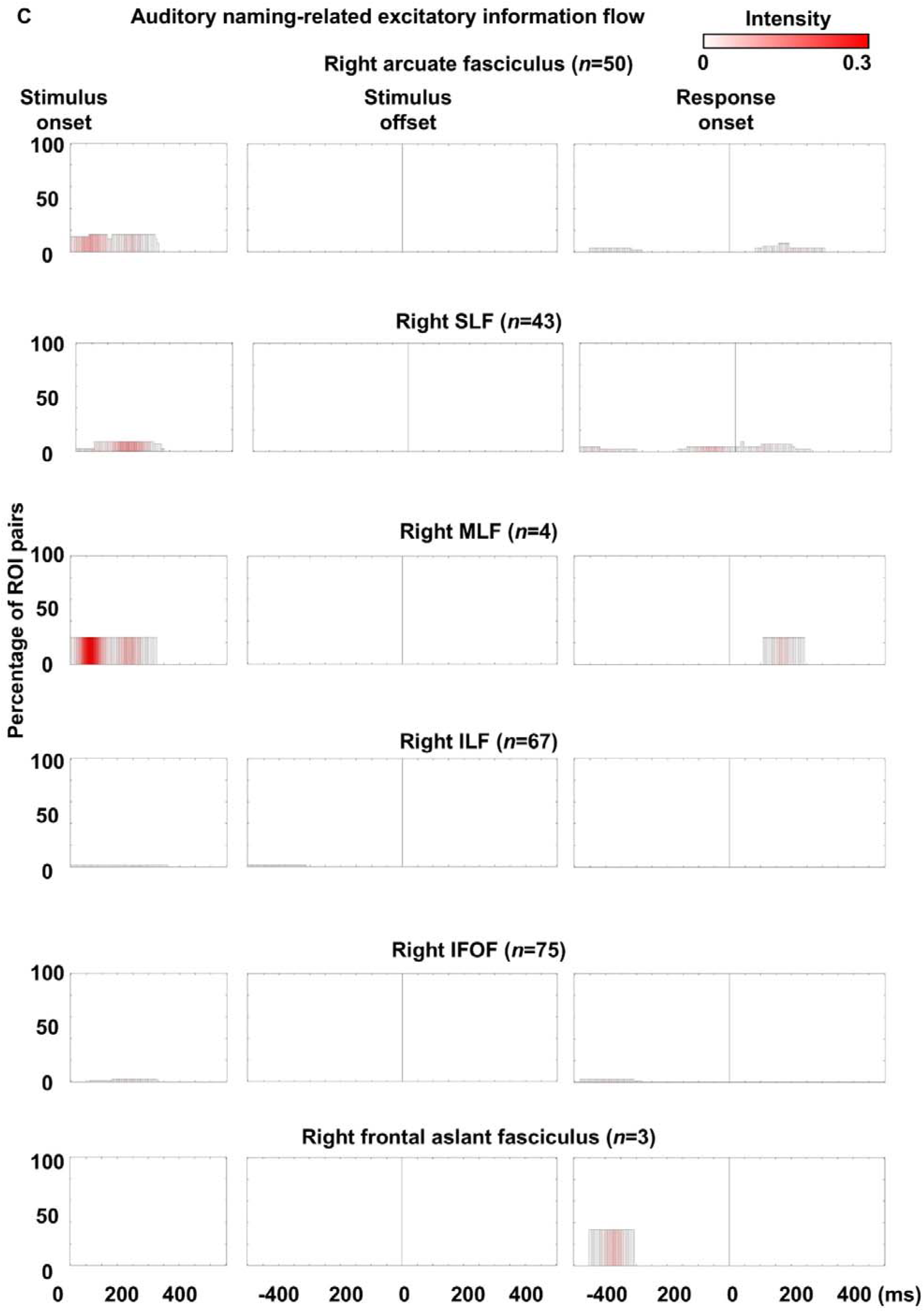
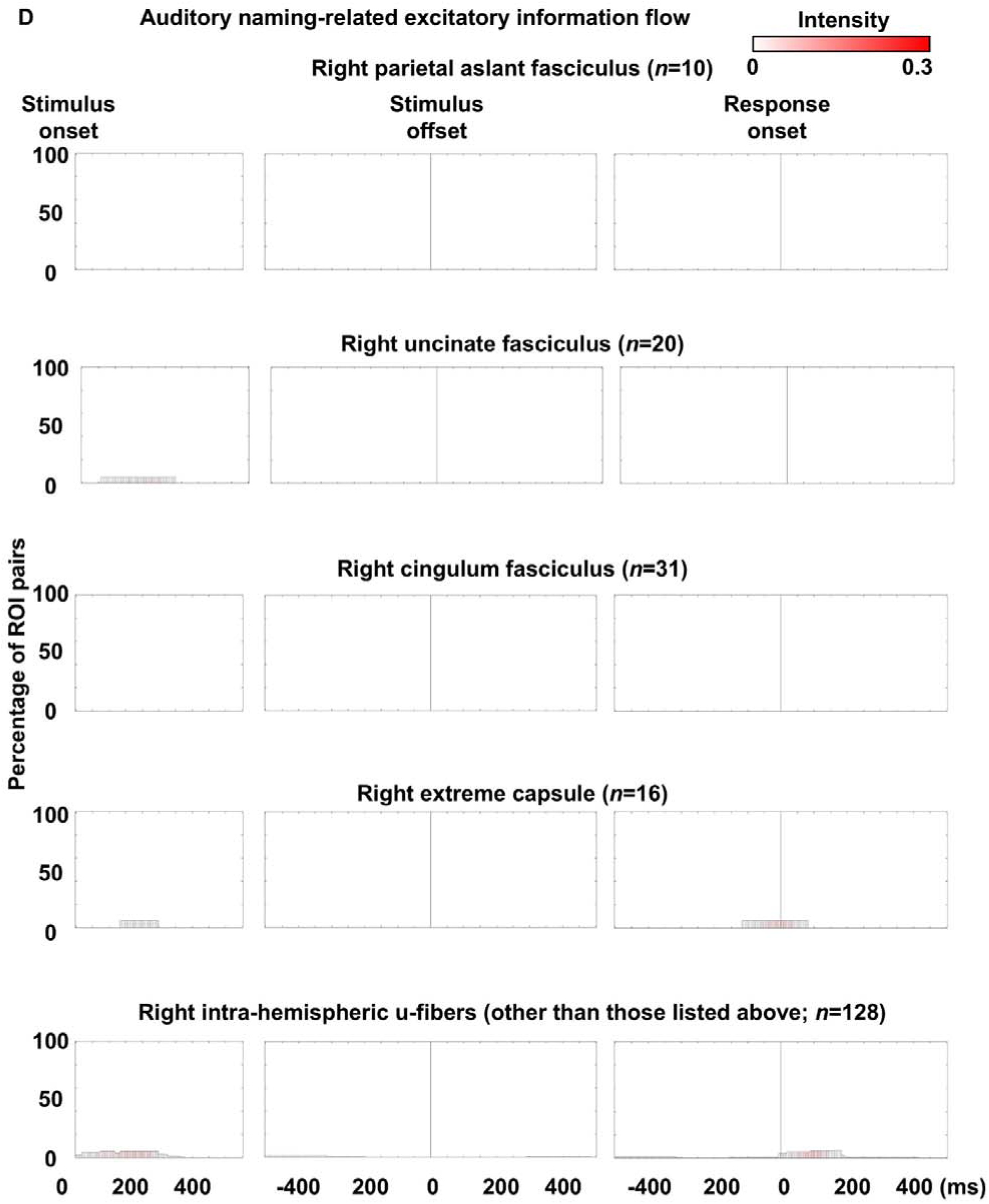
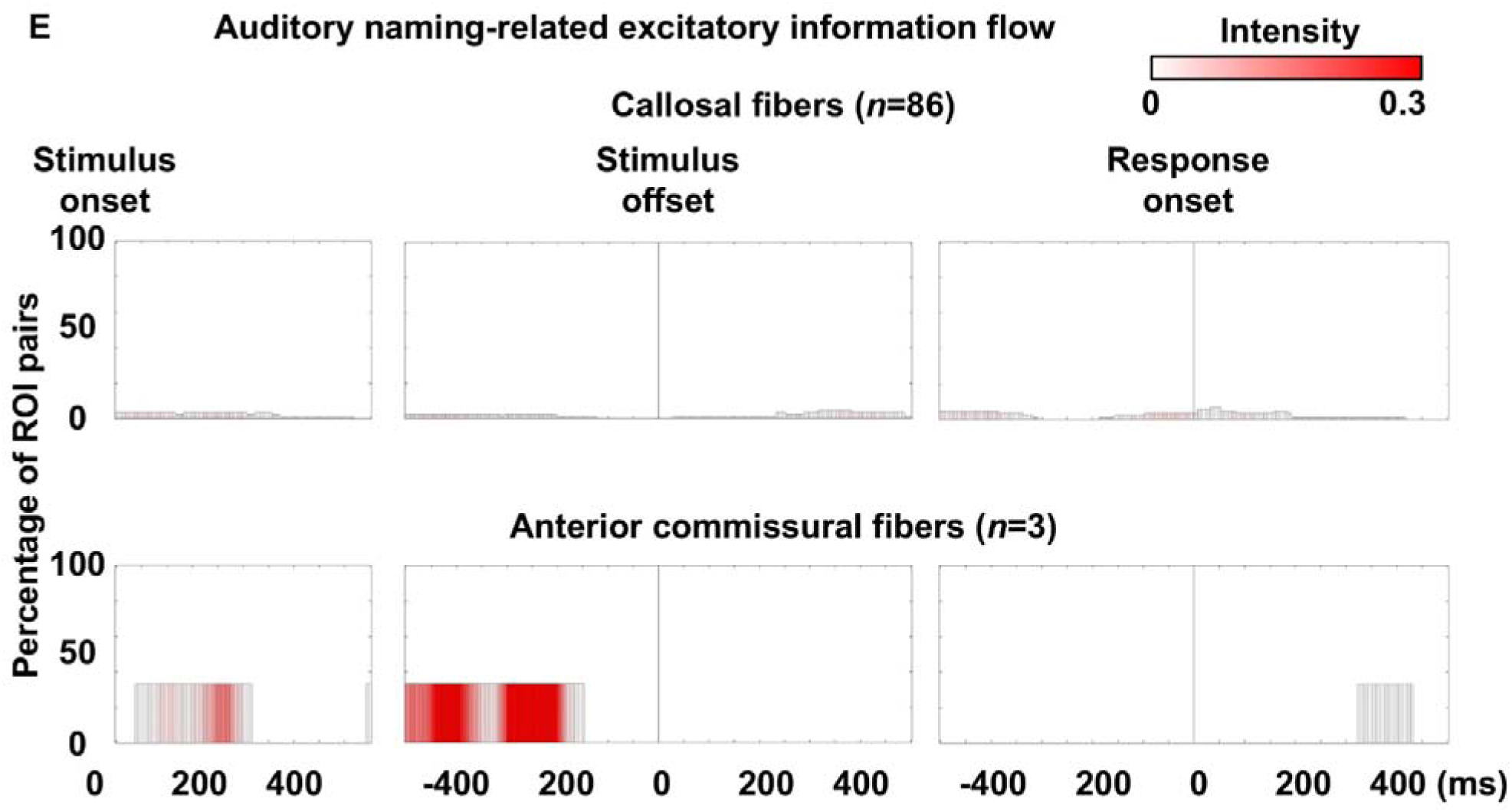
Auditory naming-related excitatory neural information flows through each fasciculus. Bar height represents the proportion of white matter pathways within each fasciculus exhibiting excitatory flows per time bin, as determined by transfer entropy–based effective connectivity. Bar color indicates mean flow intensity.

**eFigure 8.**
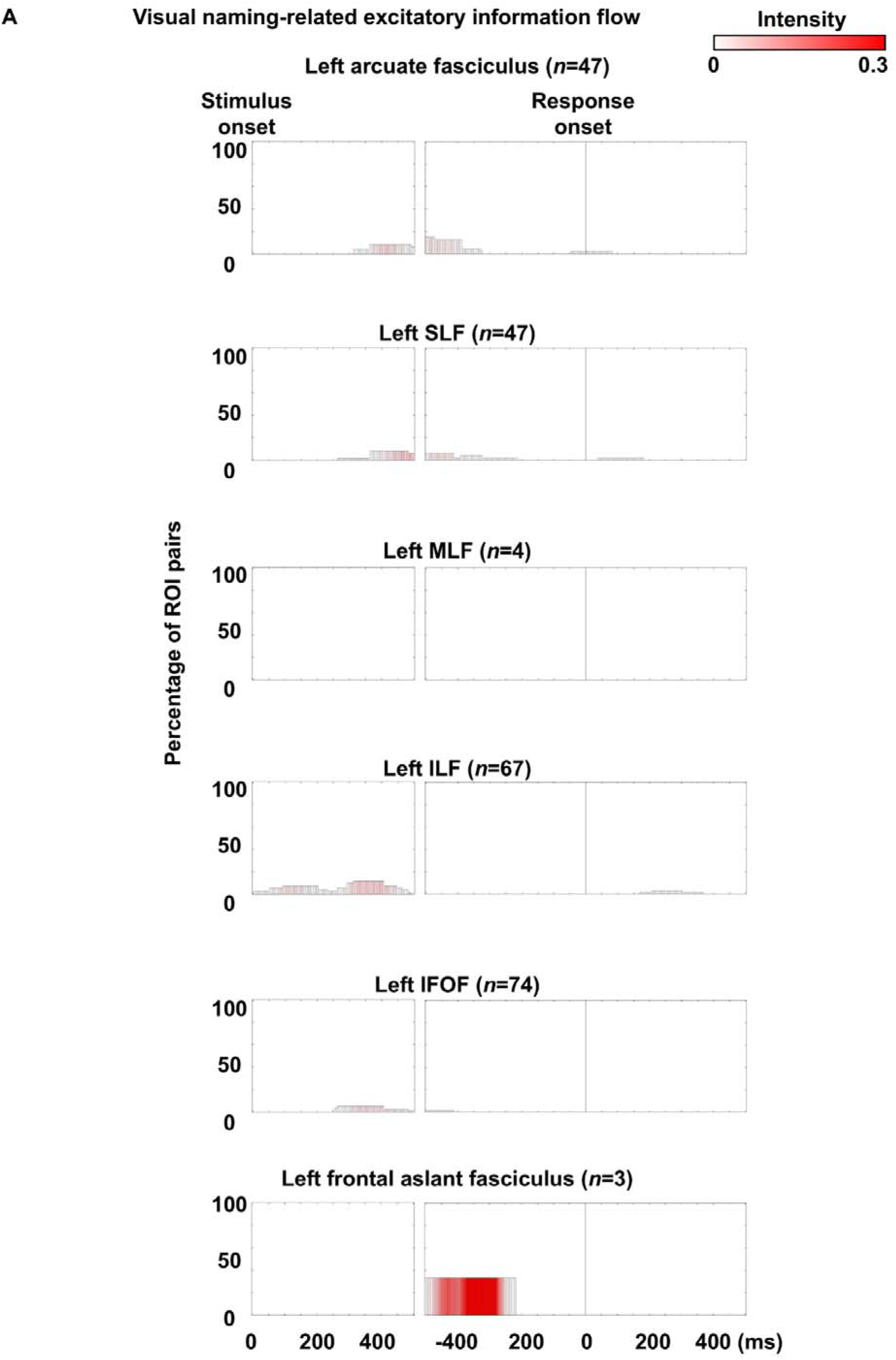
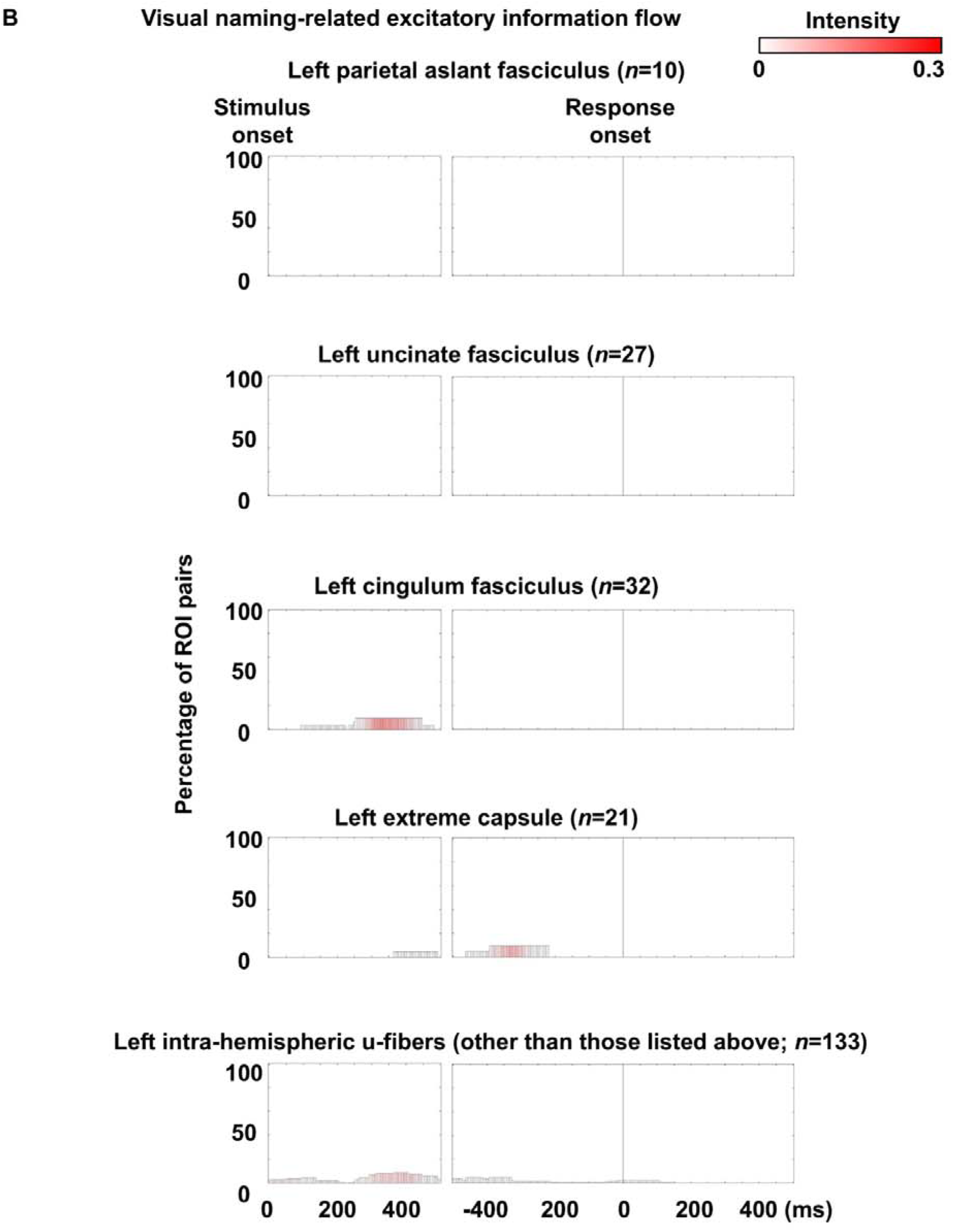
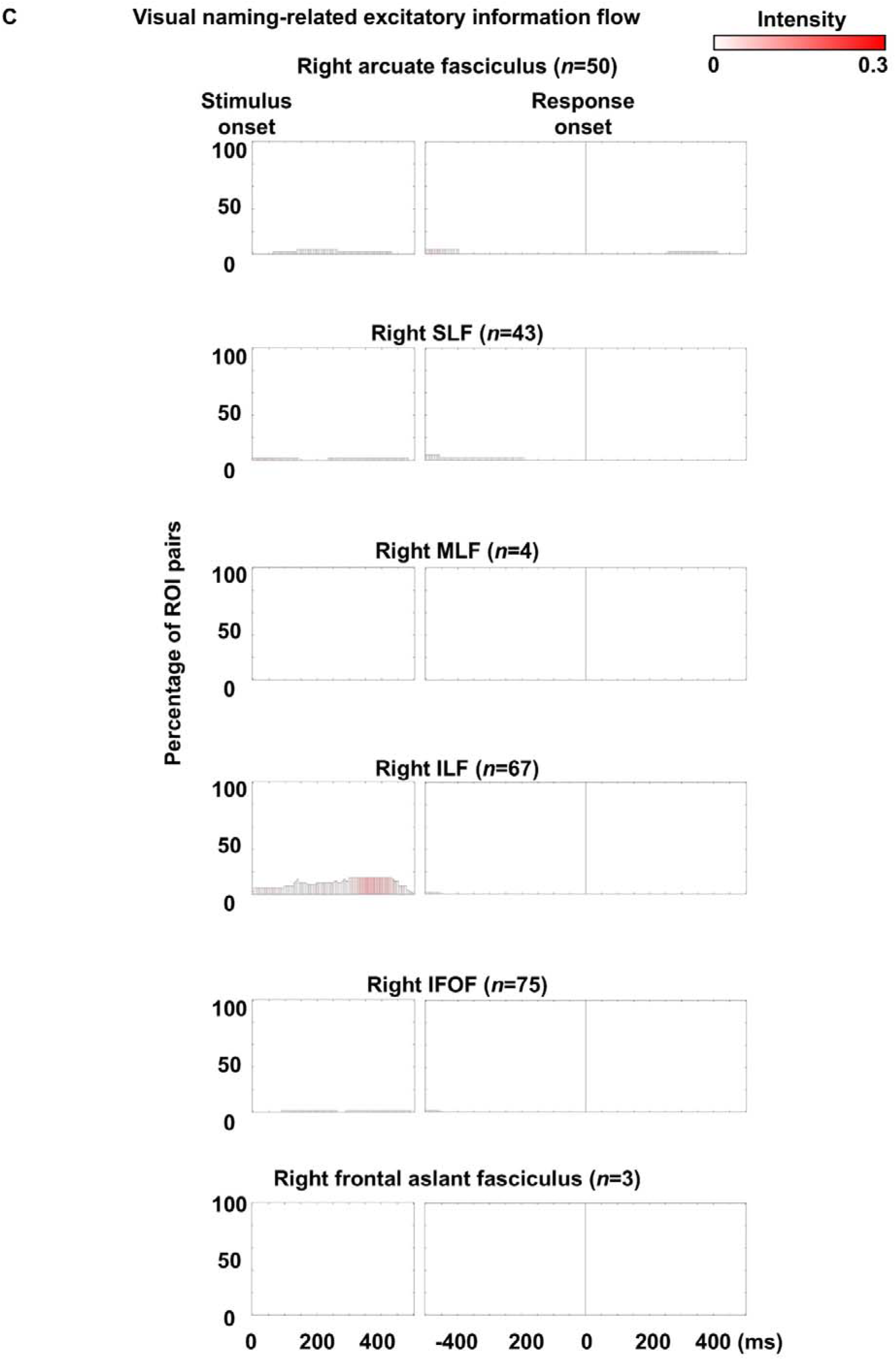
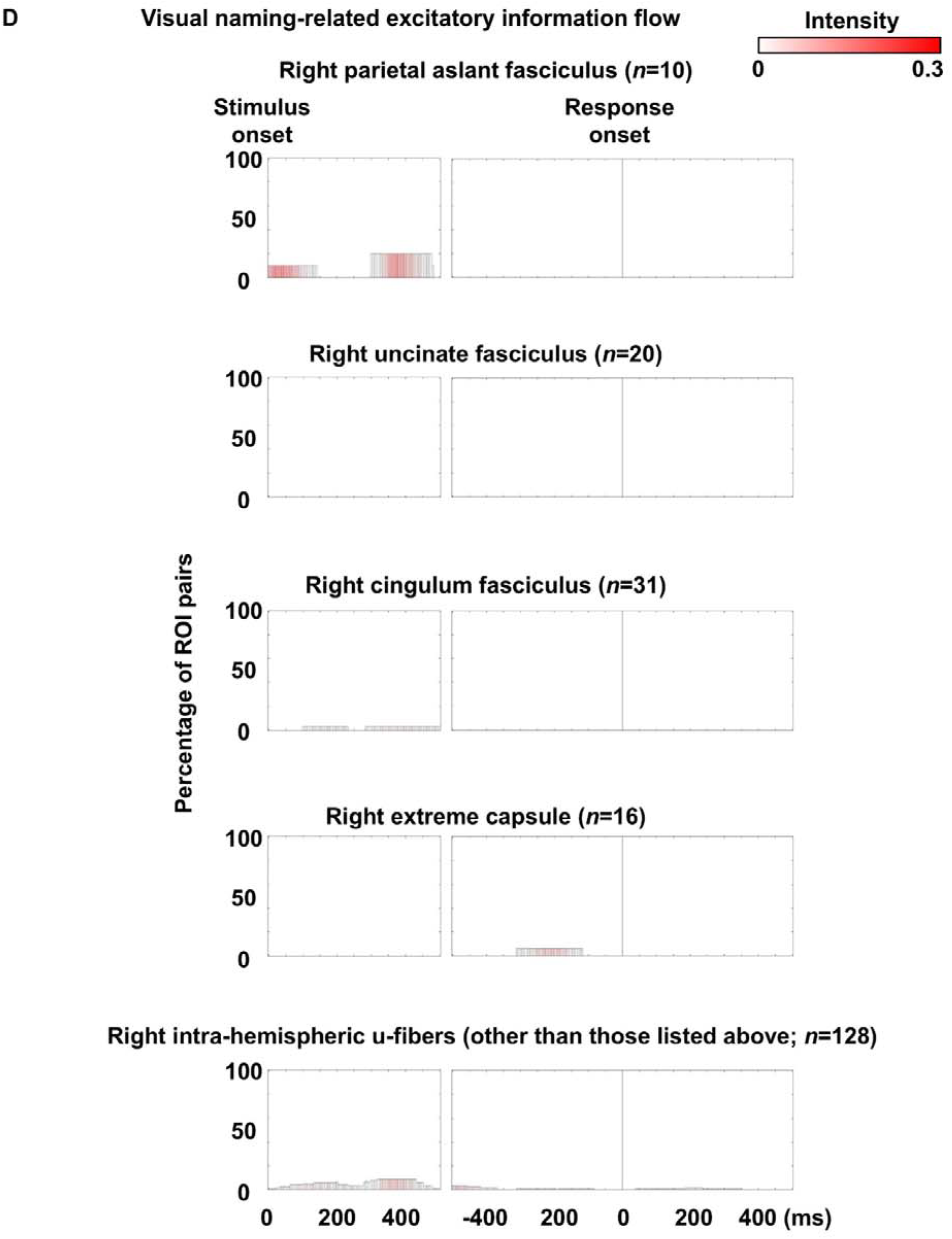
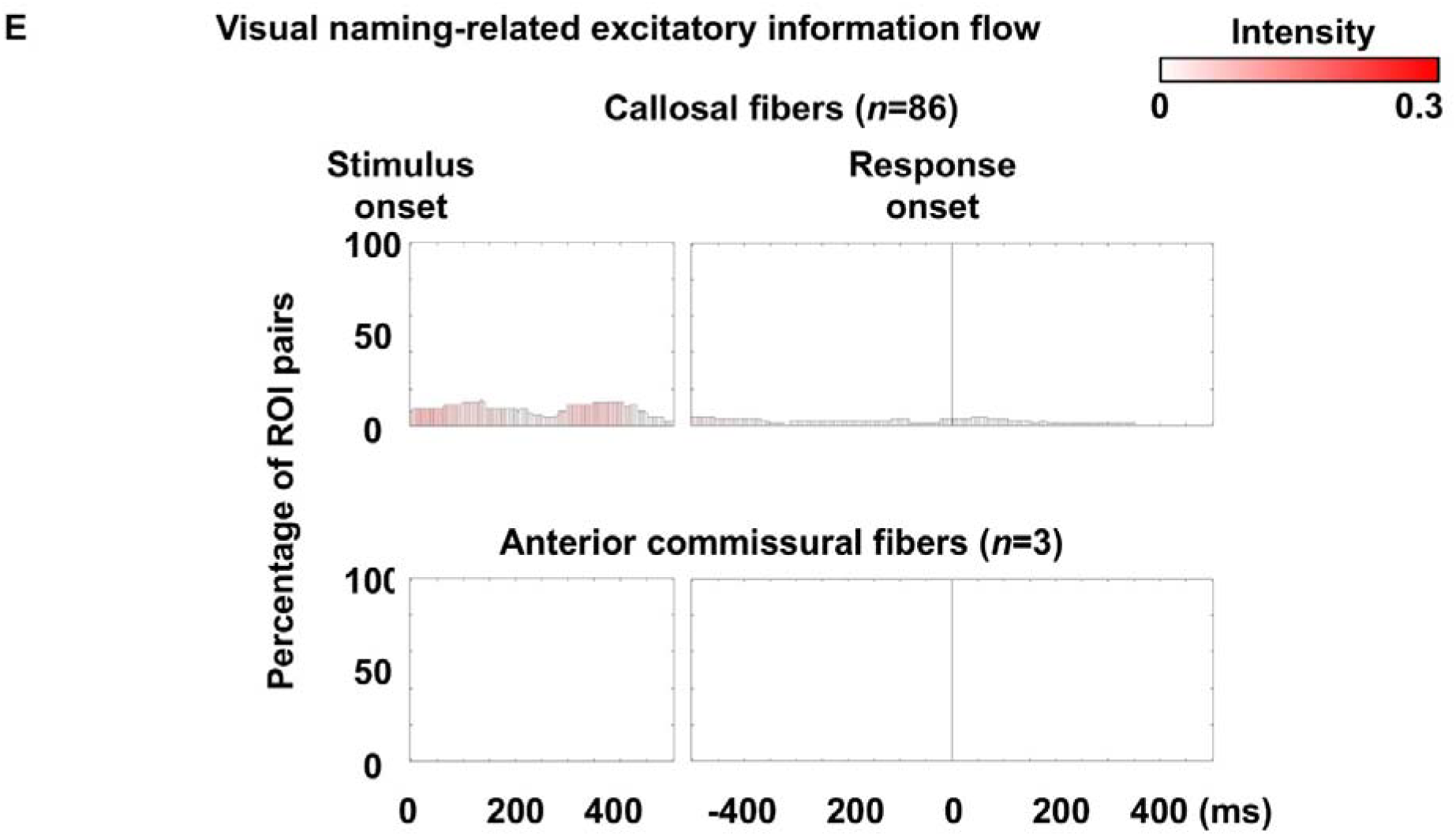
Visual naming-related excitatory neural information flows through each fasciculus. Bar height represents the proportion of white matter pathways within each fasciculus exhibiting excitatory flows per time bin, as determined by transfer entropy–based effective connectivity. Bar color indicates mean flow intensity.

**eFigure 9.**
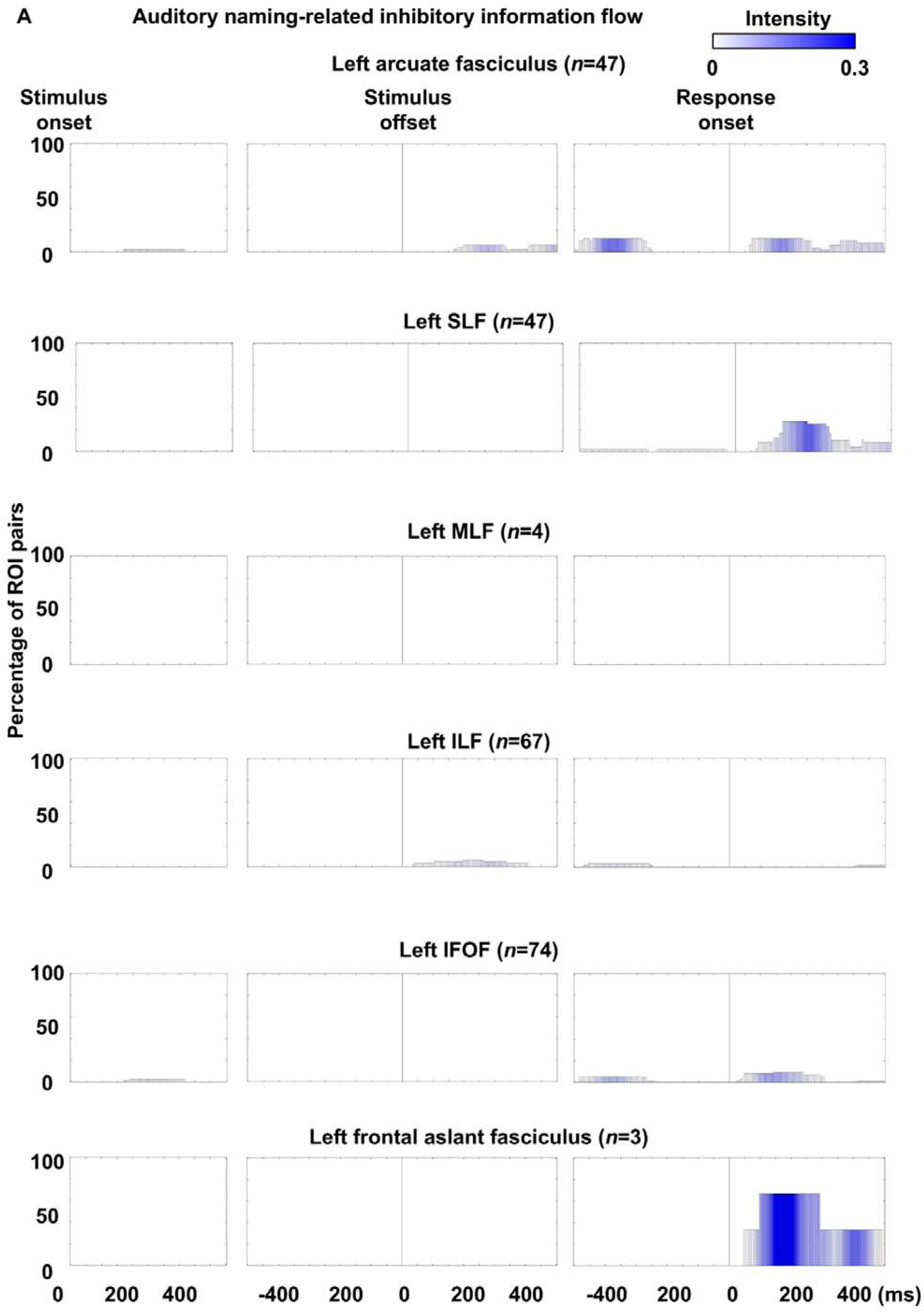
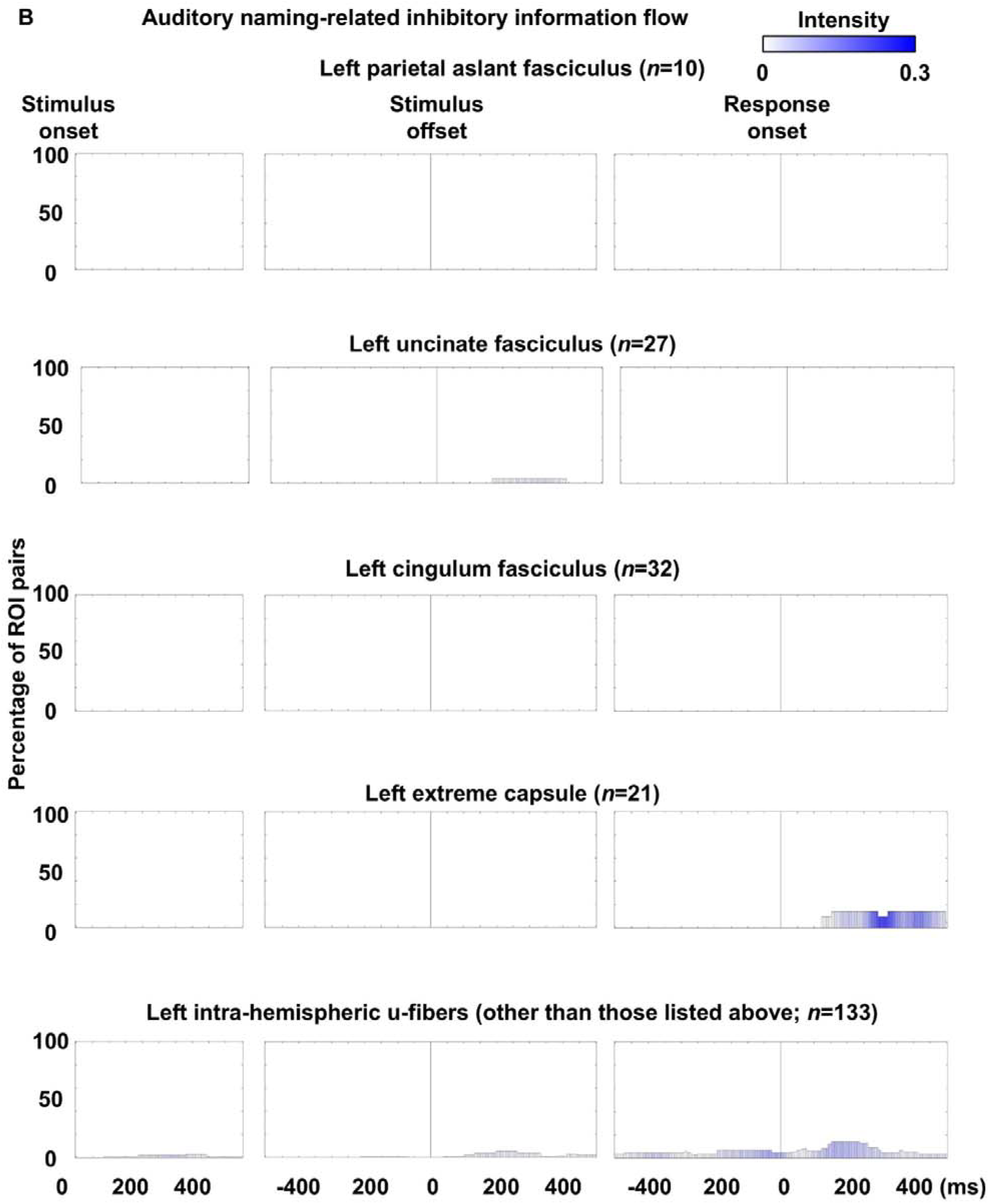
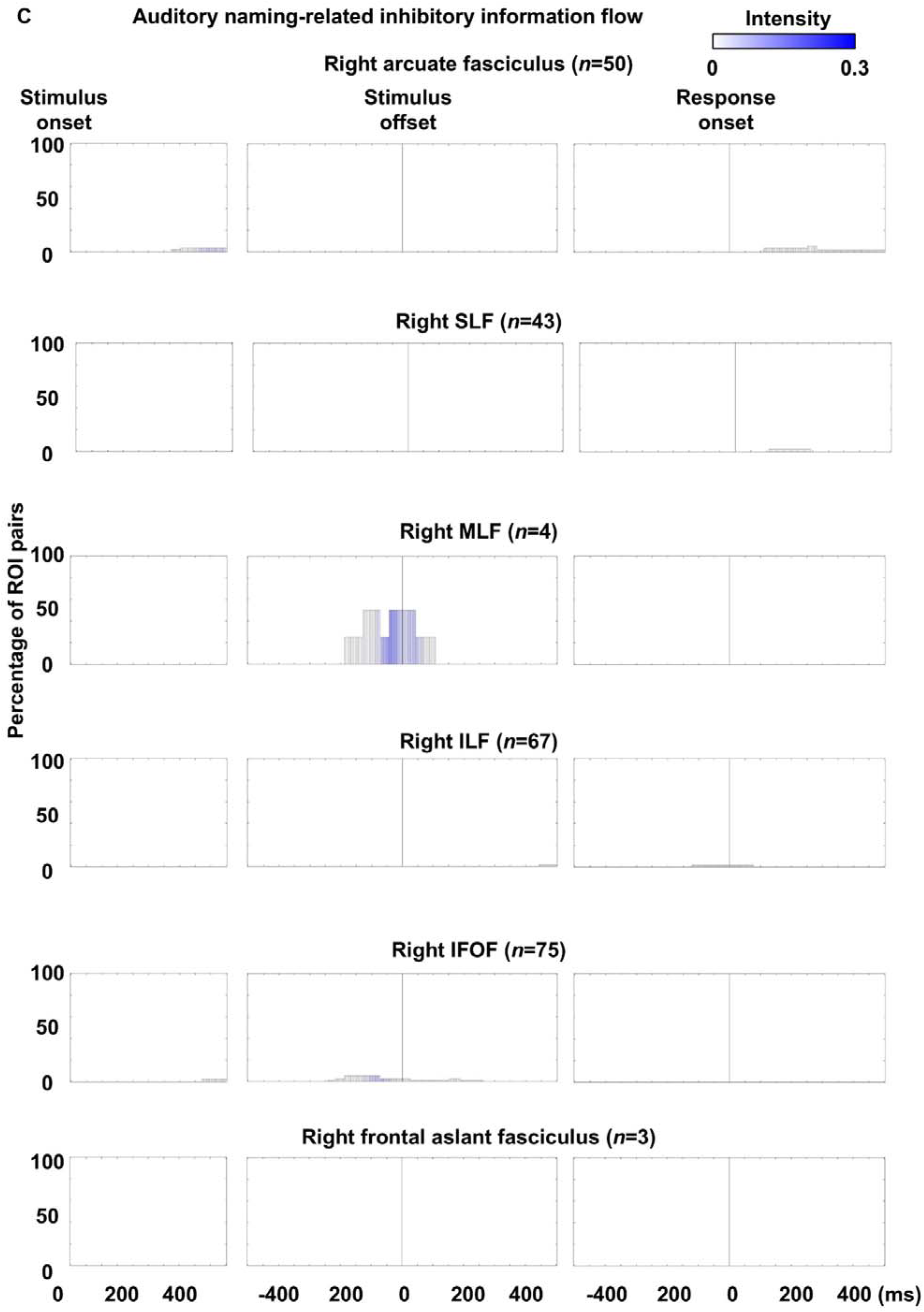
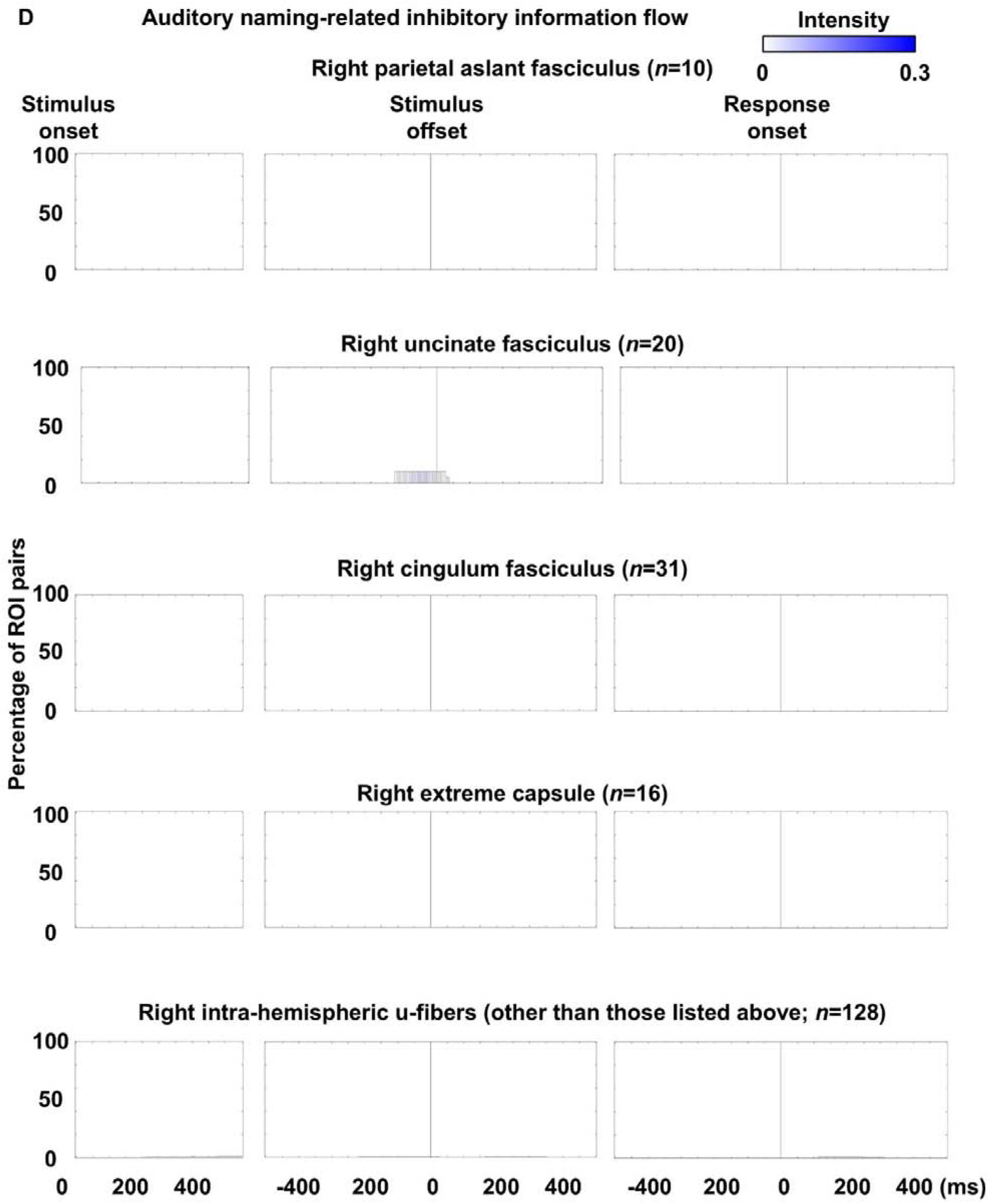
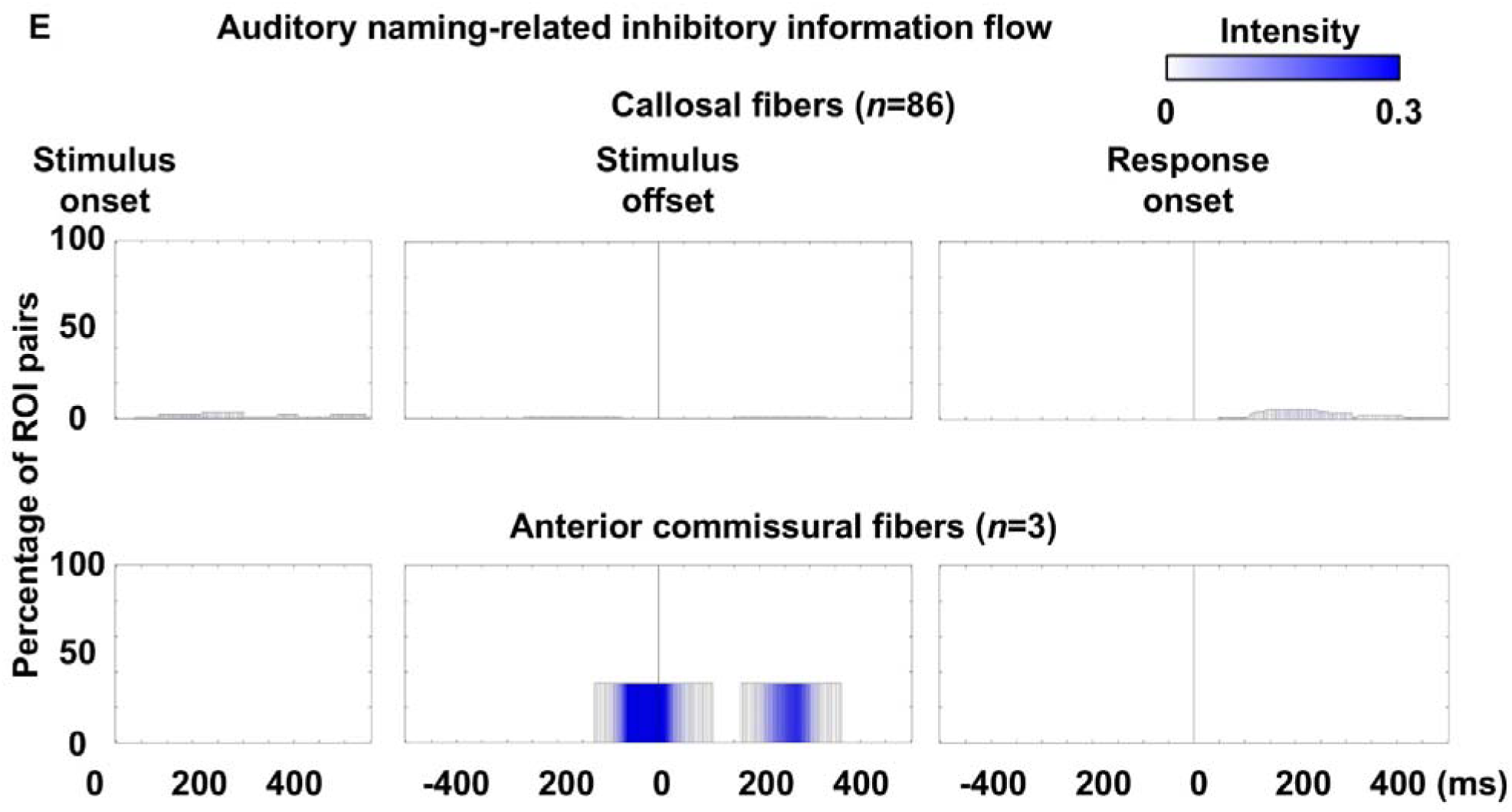
Auditory naming-related inhibitory neural information flows through each fasciculus. Bar height represents the proportion of white matter pathways within each fasciculus exhibiting inhibitory flows per time bin, as determined by transfer entropy–based effective connectivity. Bar color indicates mean flow intensity.

**eFigure 10.**
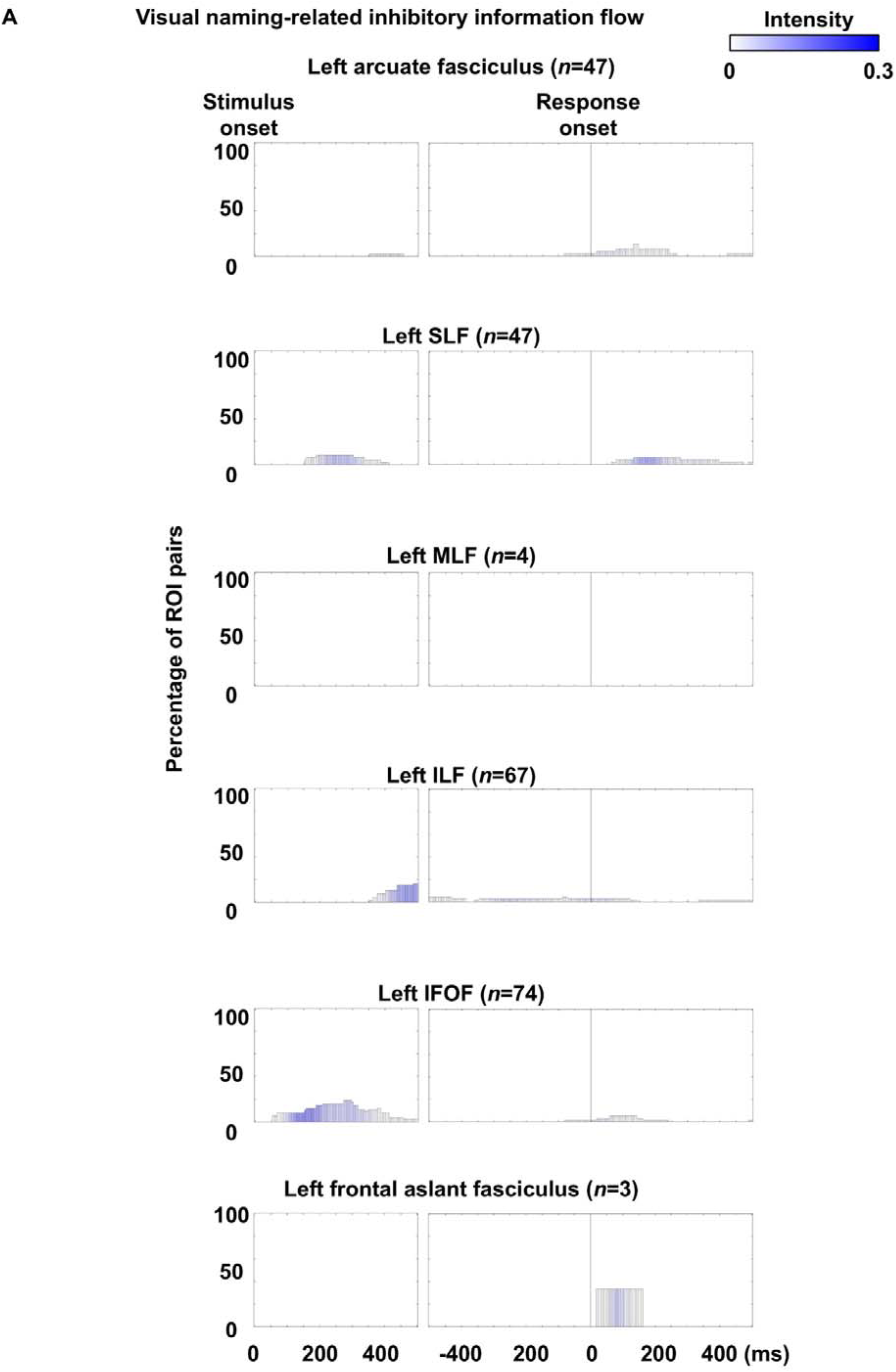
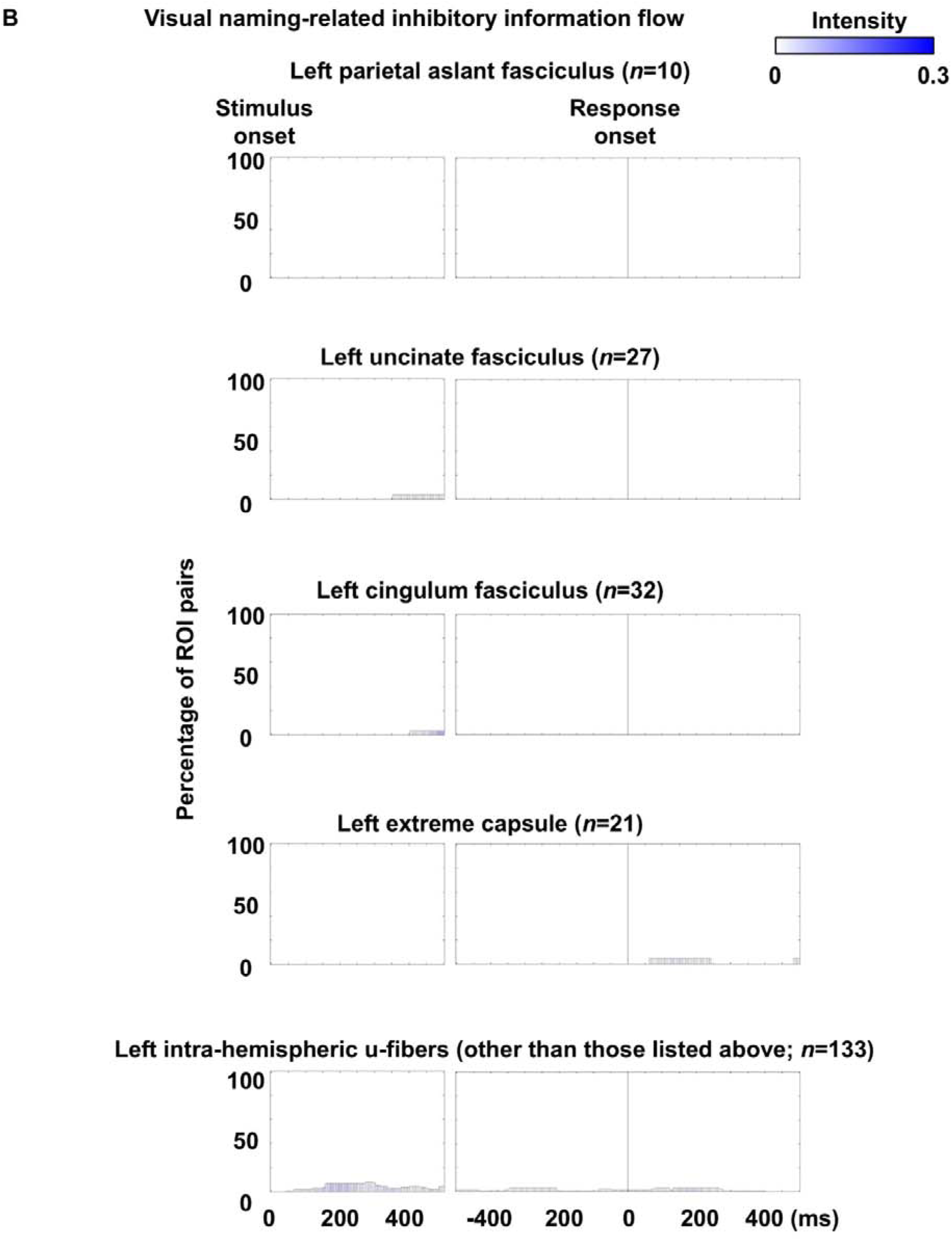
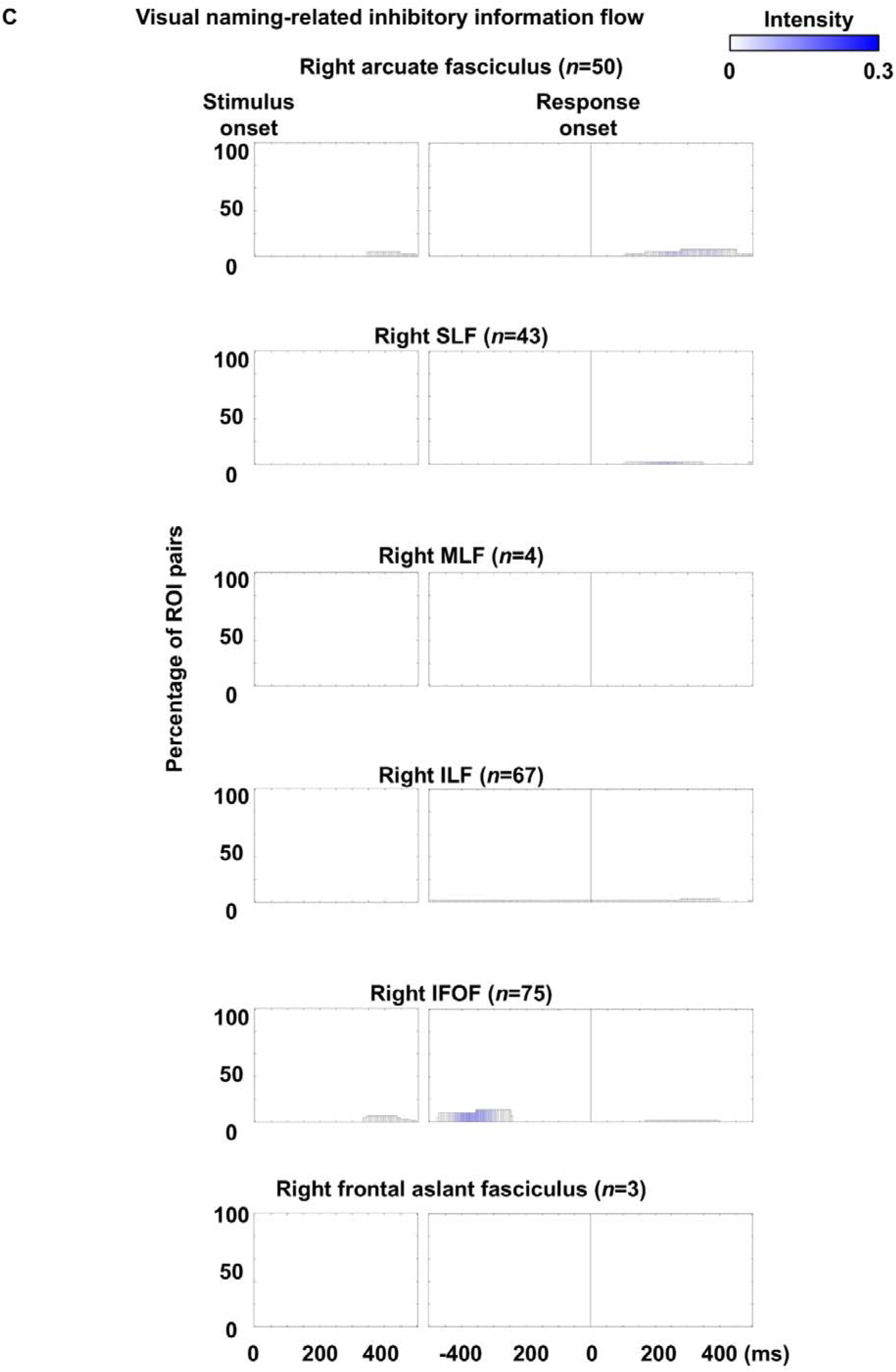
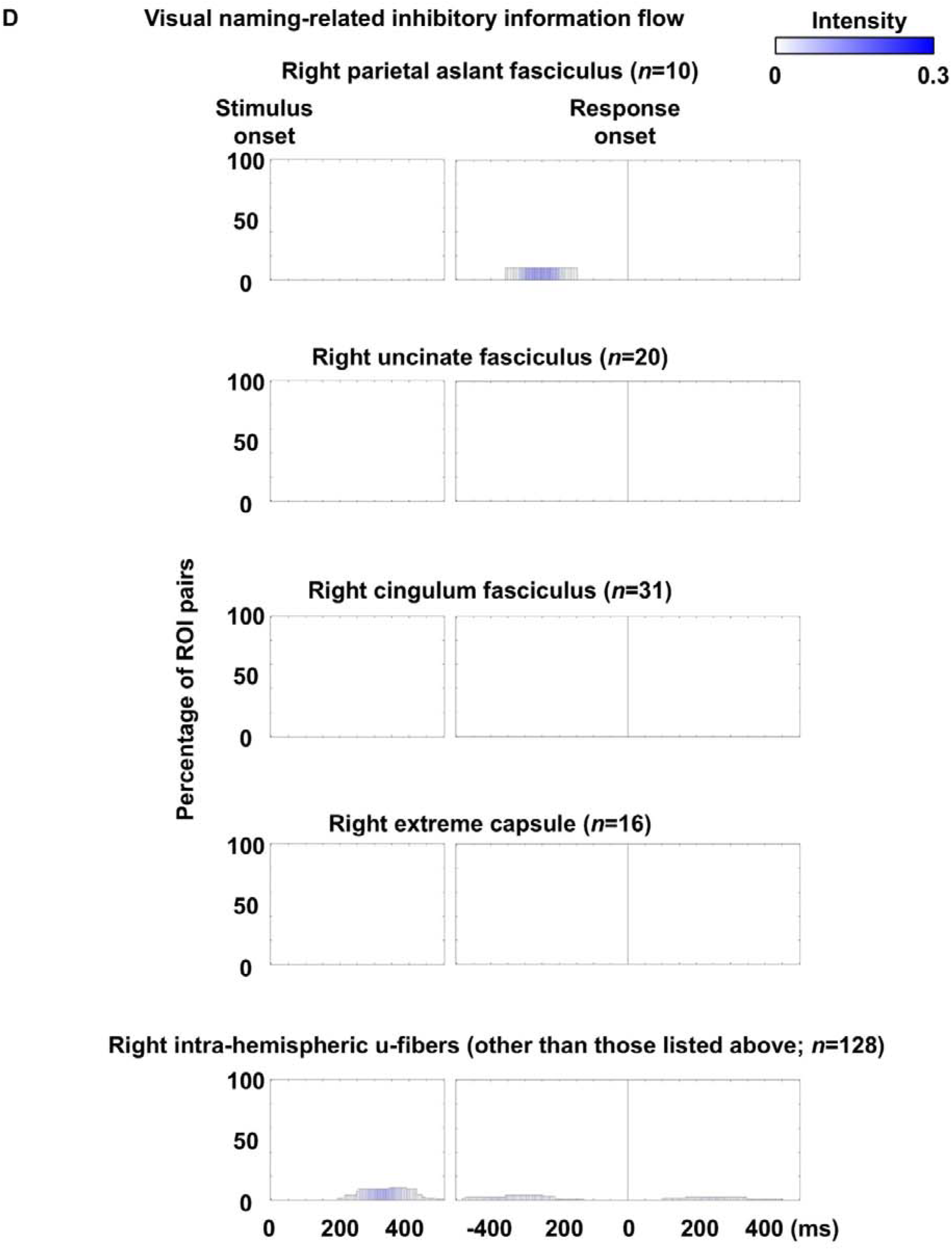
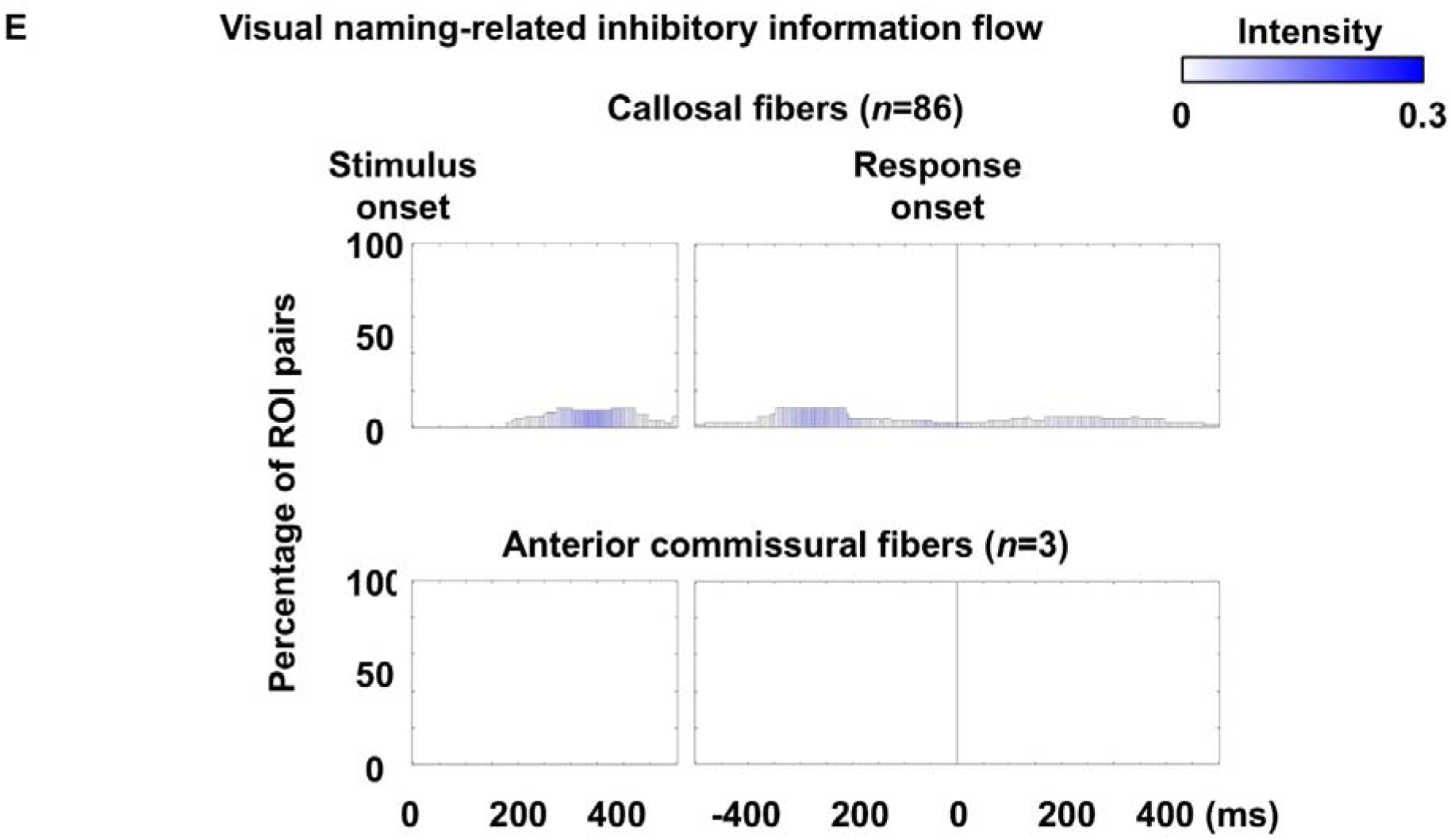
Visual naming-related inhibitory neural information flows through each fasciculus. Bar height represents the proportion of white matter pathways within each fasciculus exhibiting inhibitory flows per time bin, as determined by transfer entropy–based effective connectivity. Bar color indicates mean flow intensity.

**eFigure 11.**
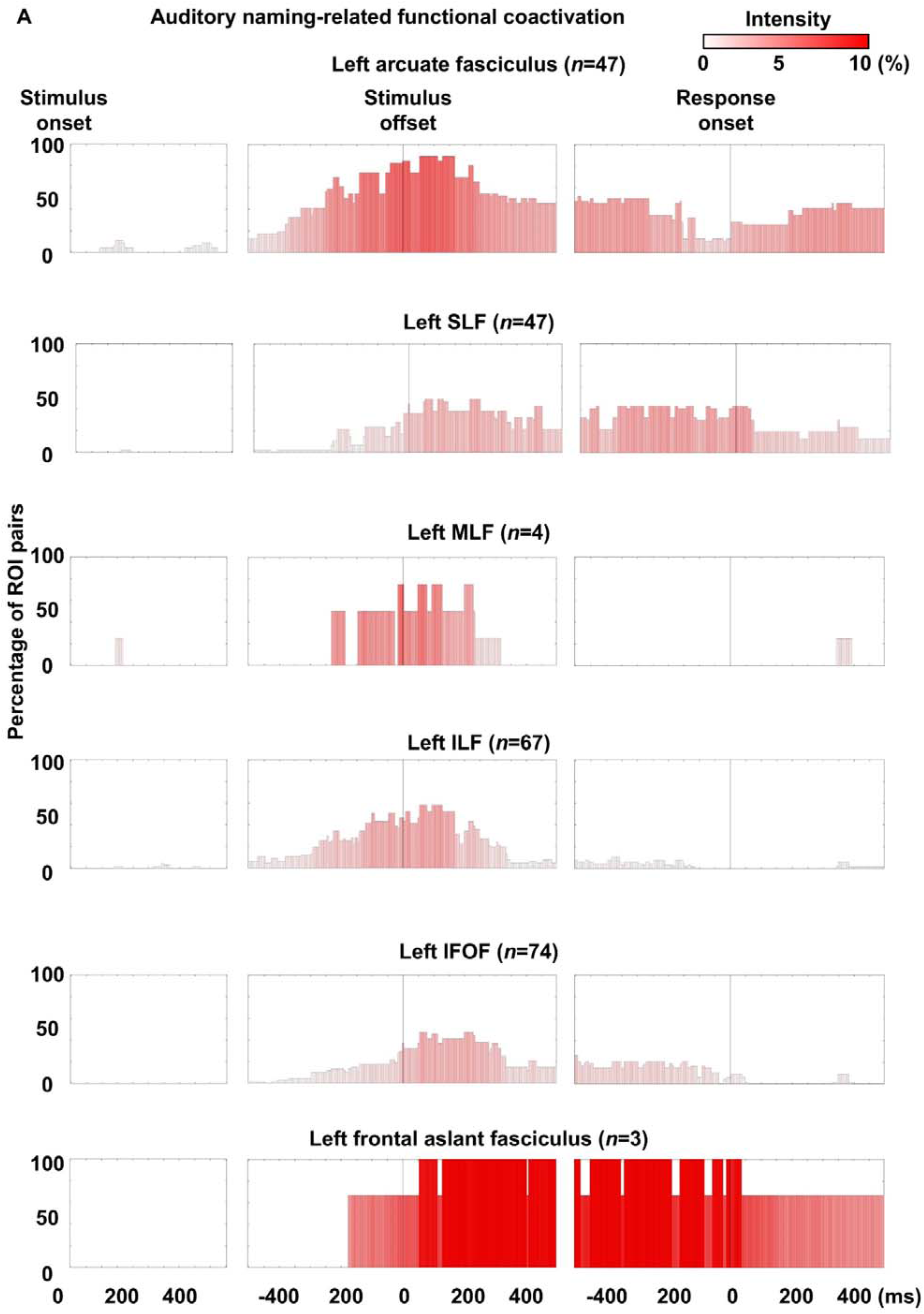
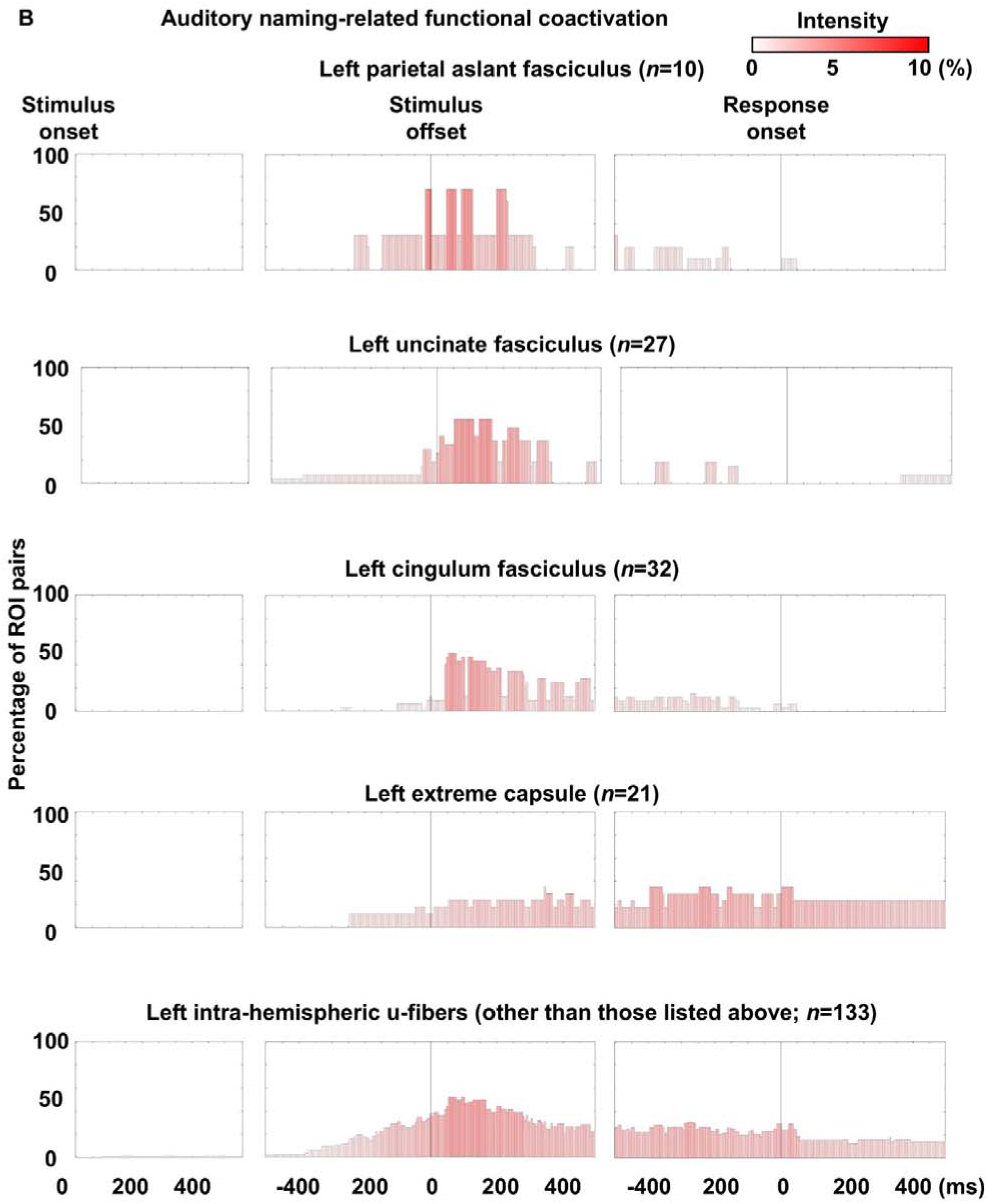
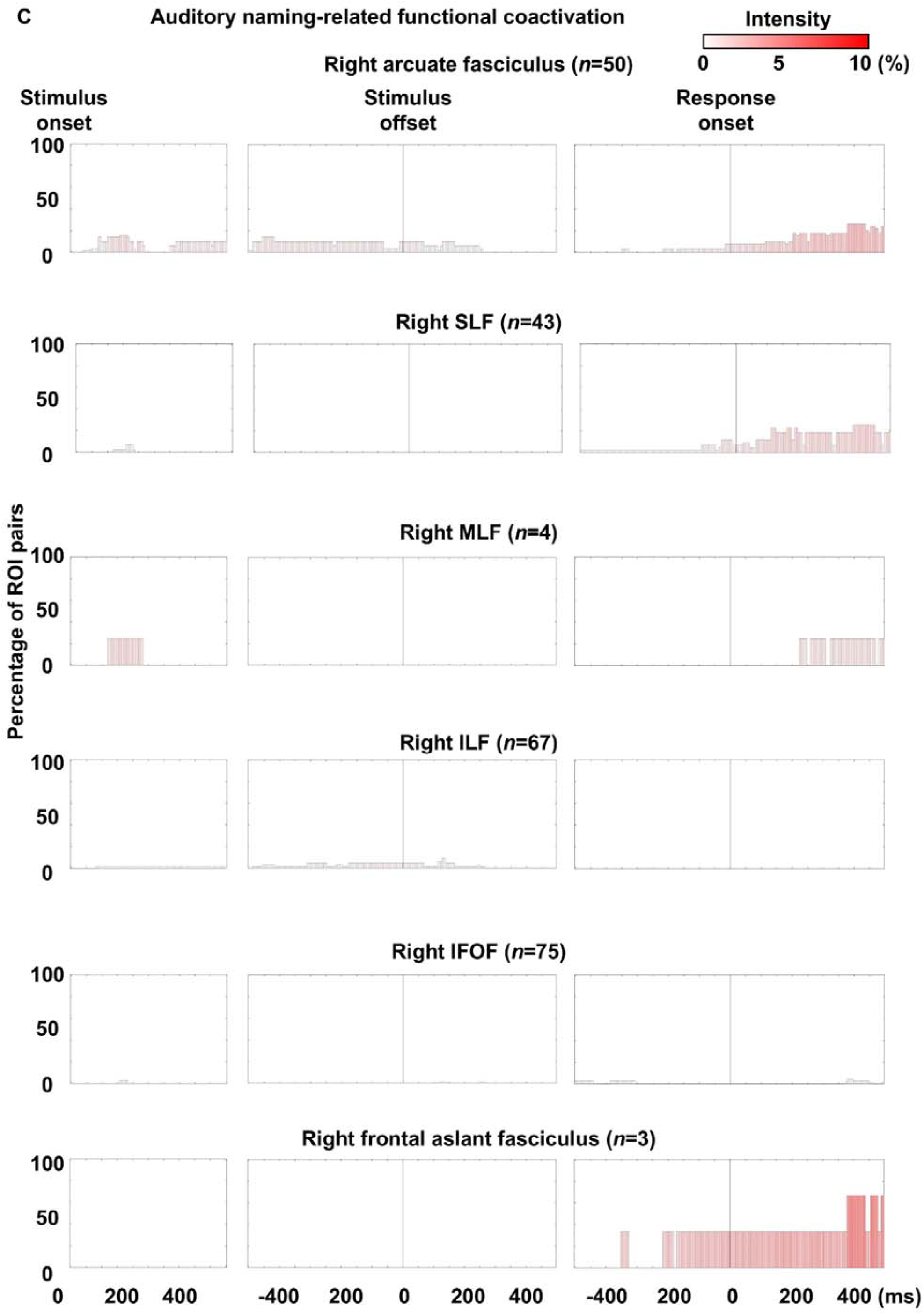
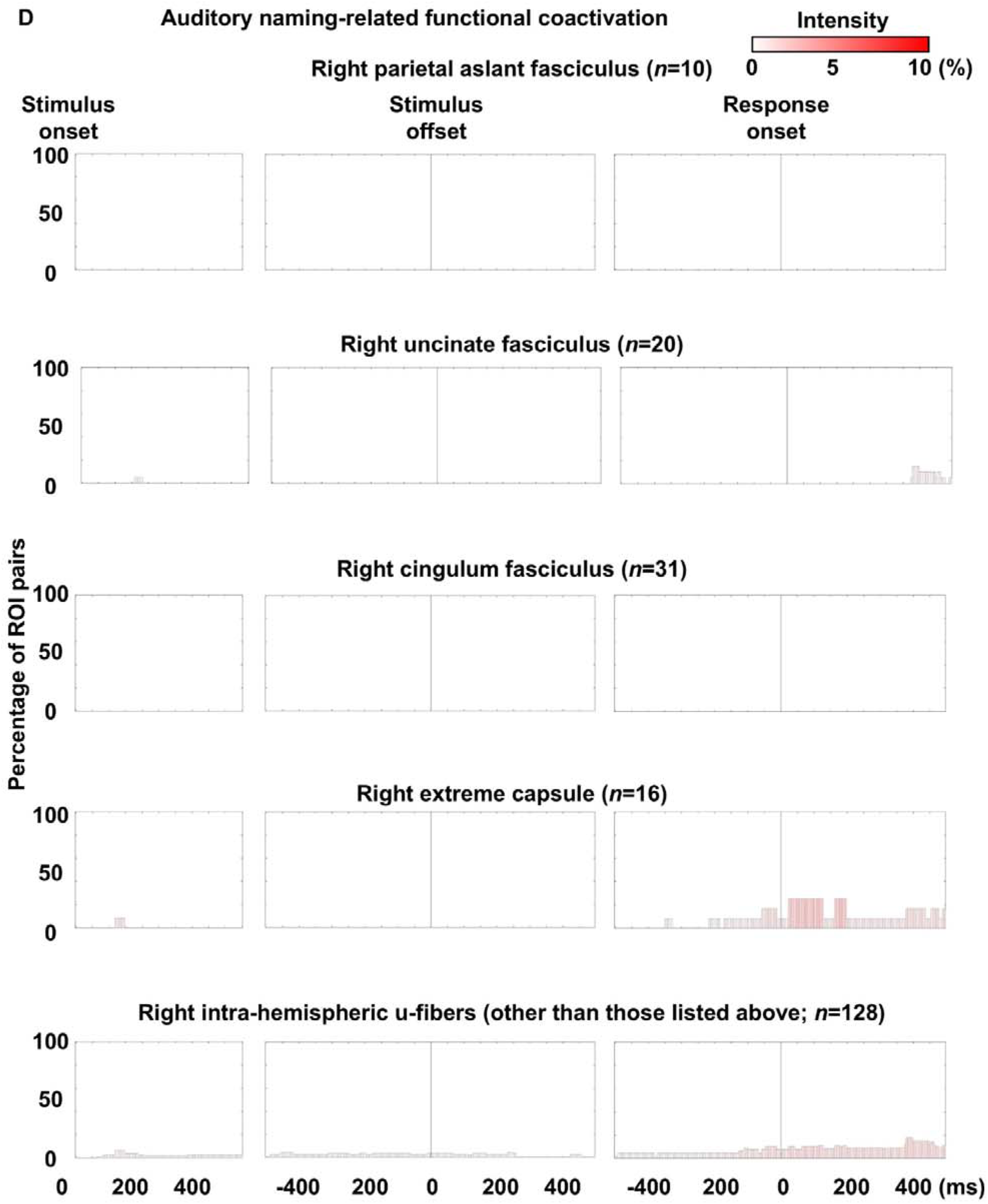
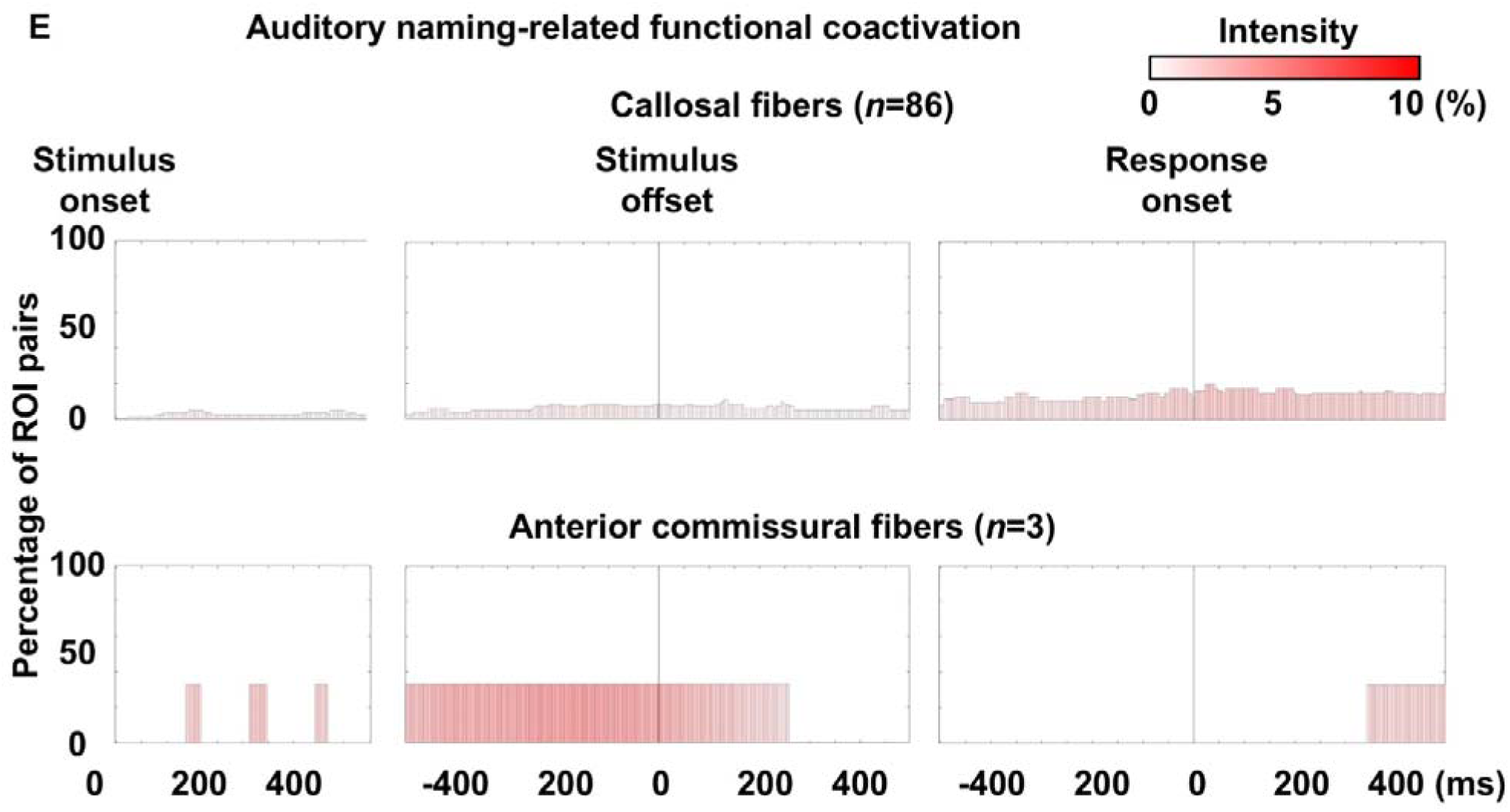
Auditory naming-related functional coactivation through each fasciculus. Bar height represents the proportion of white matter pathways within each fasciculus exhibiting functional coactivation per time bin. Bar color indicates mean coactivation intensity.

**eFigure 12.**
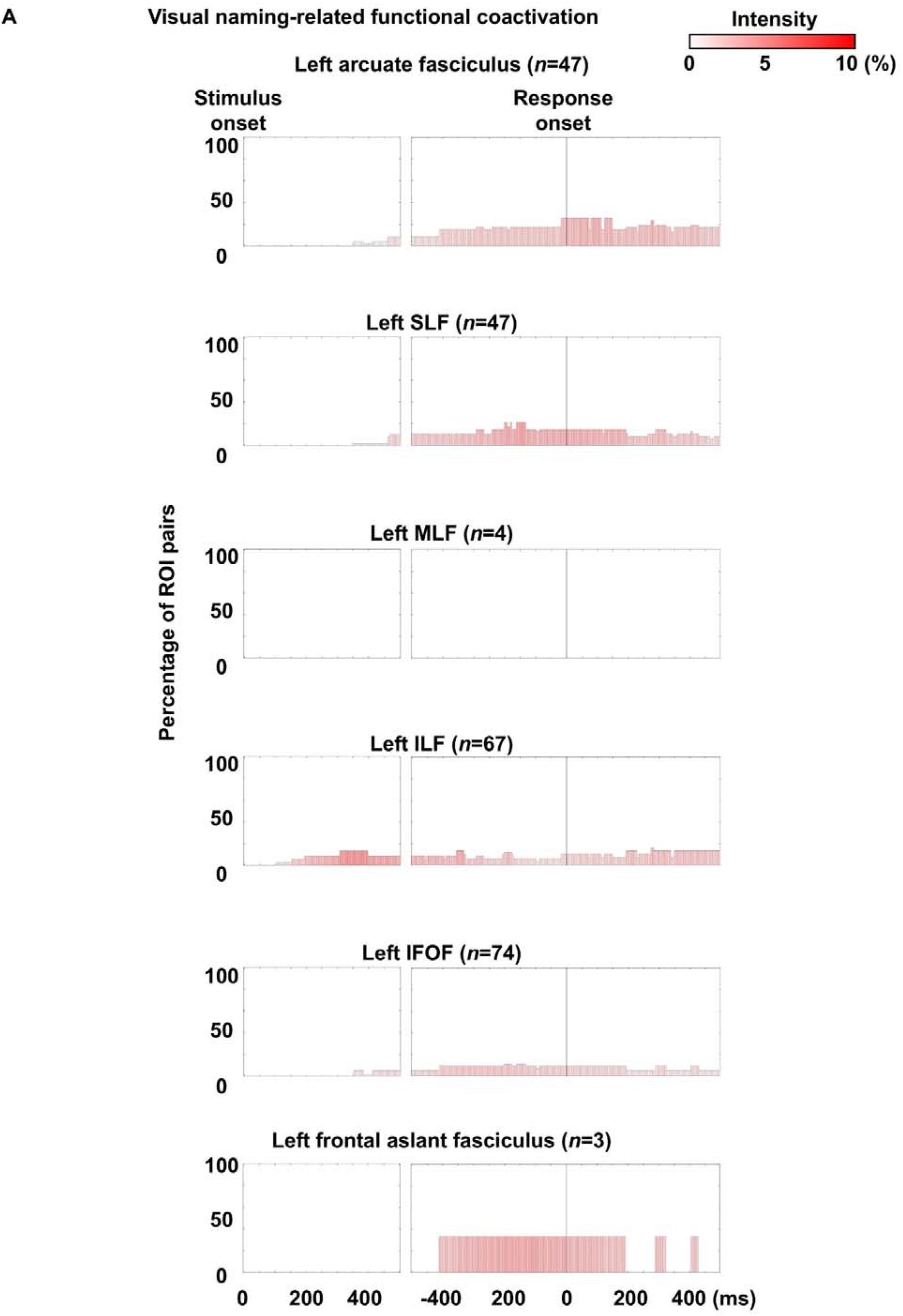
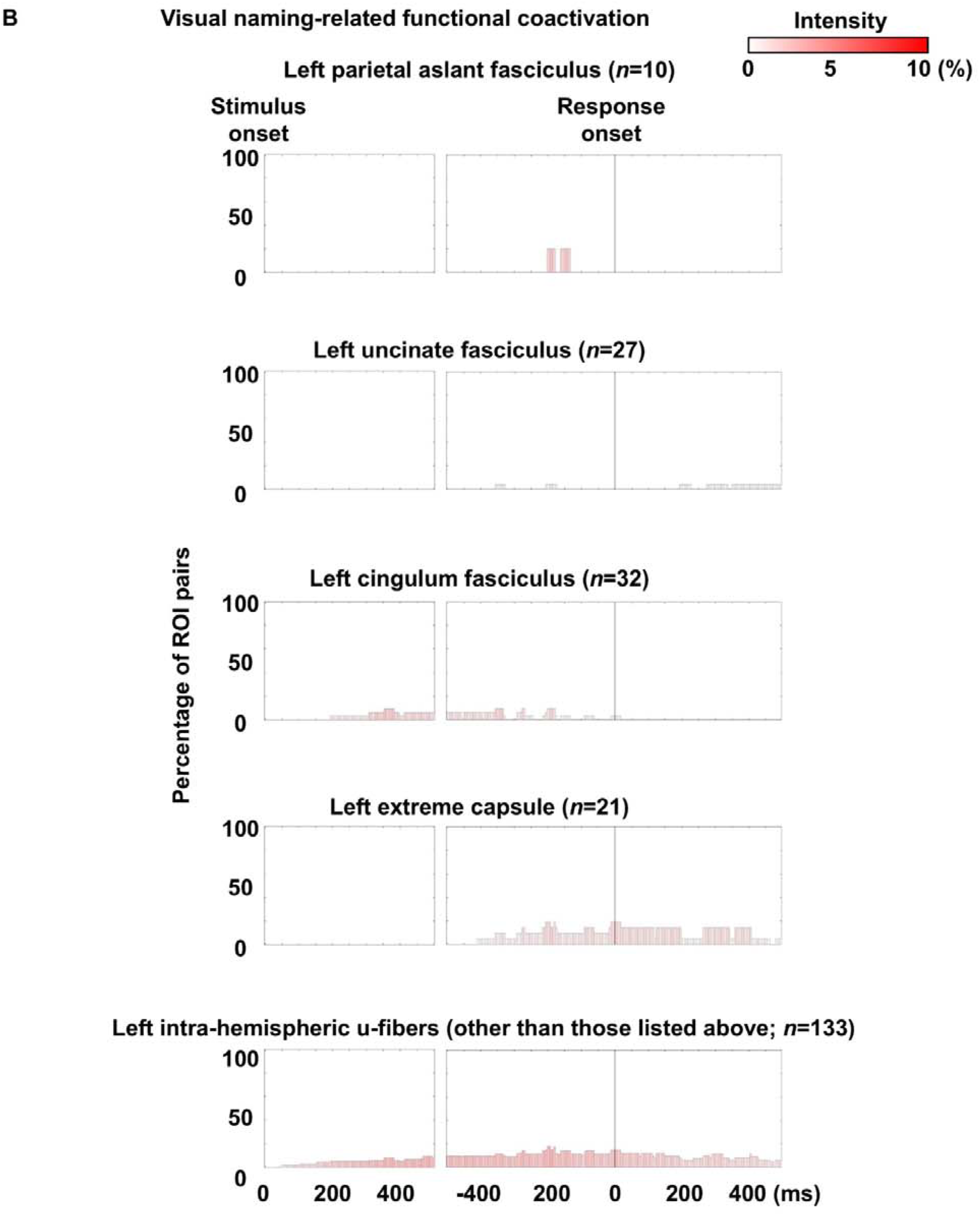
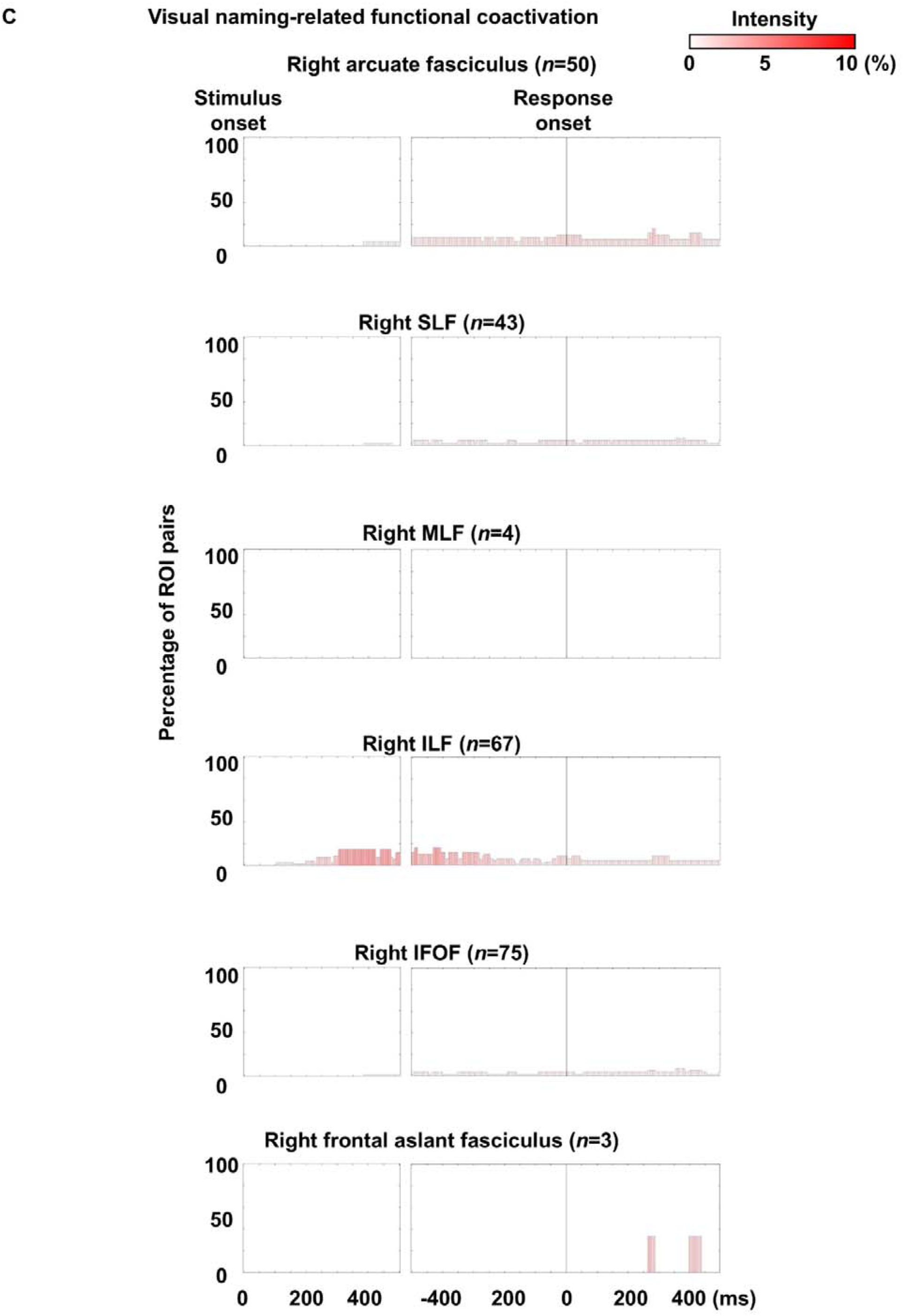
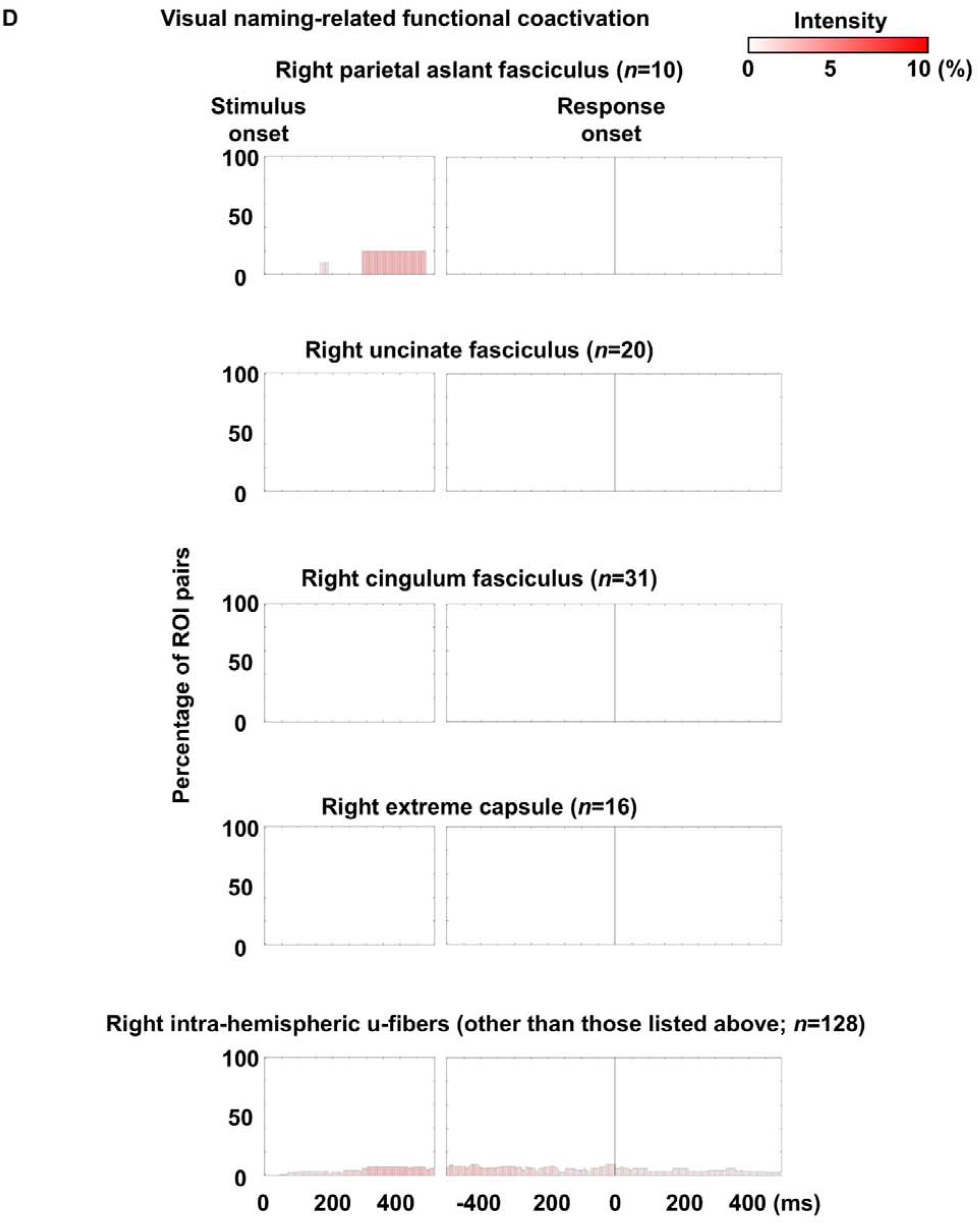
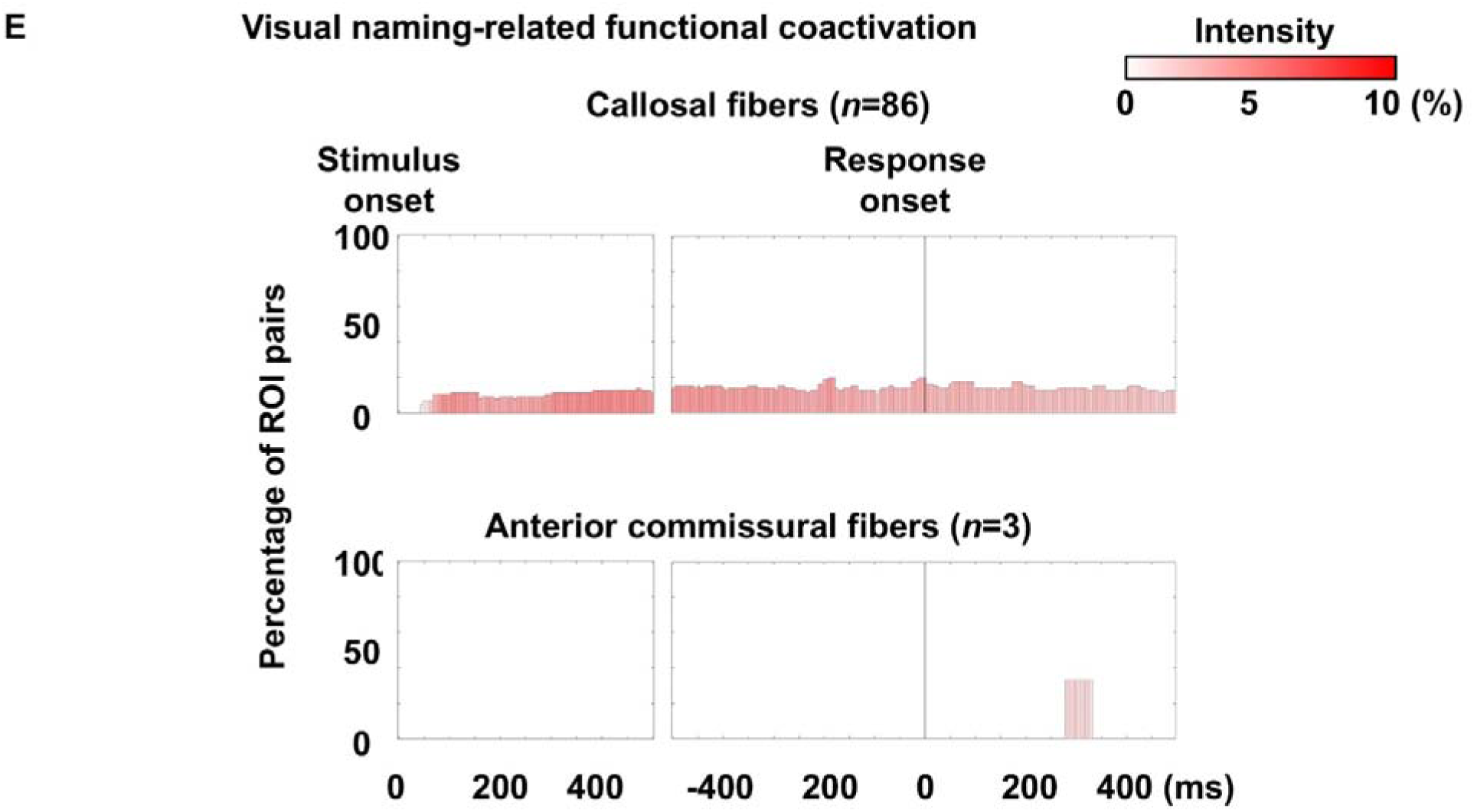
Visual naming-related functional coactivation through each fasciculus. Bar height represents the proportion of white matter pathways within each fasciculus exhibiting functional coactivation per time bin. Bar color indicates mean coactivation intensity.

**eFigure 13.**
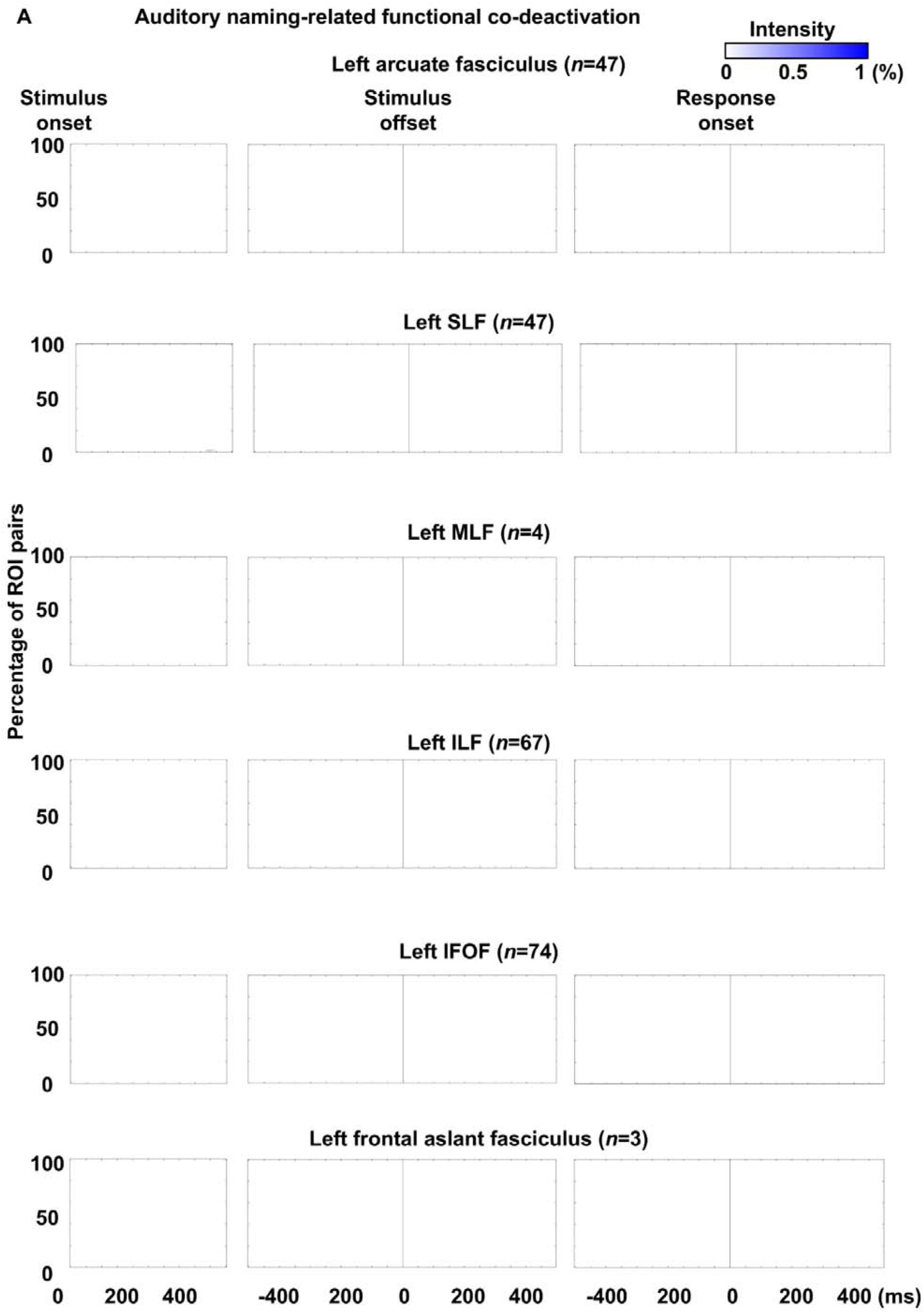
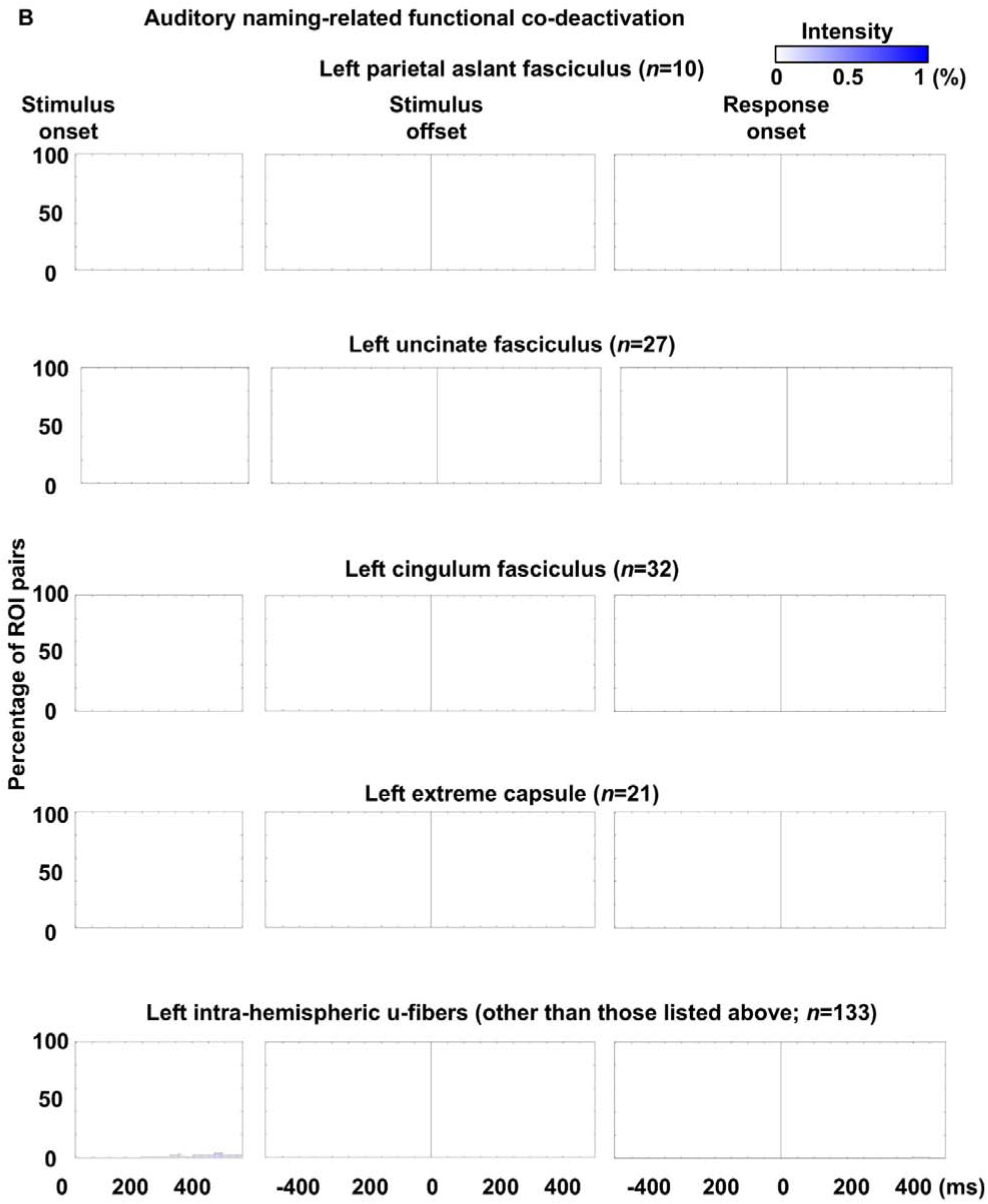
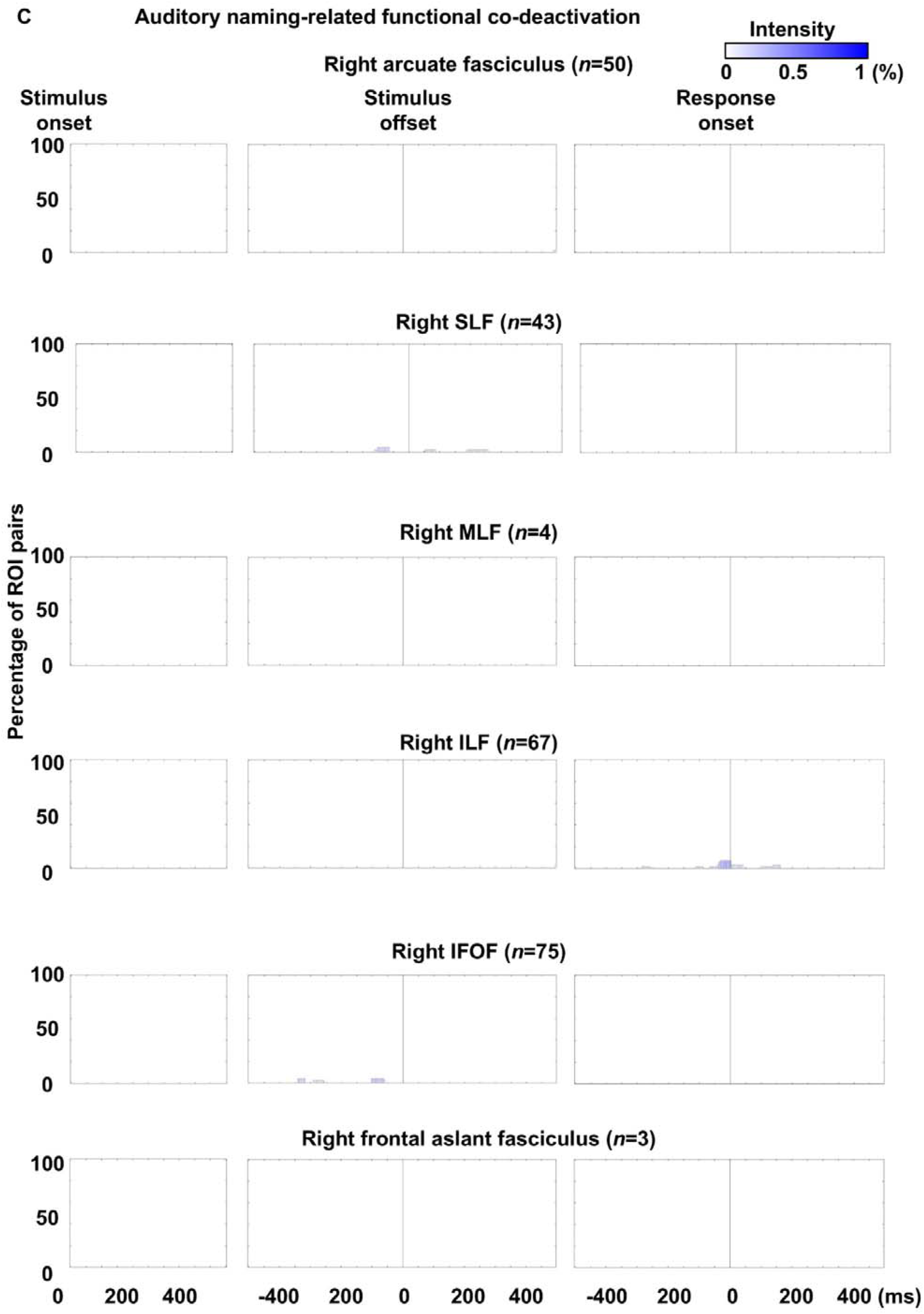
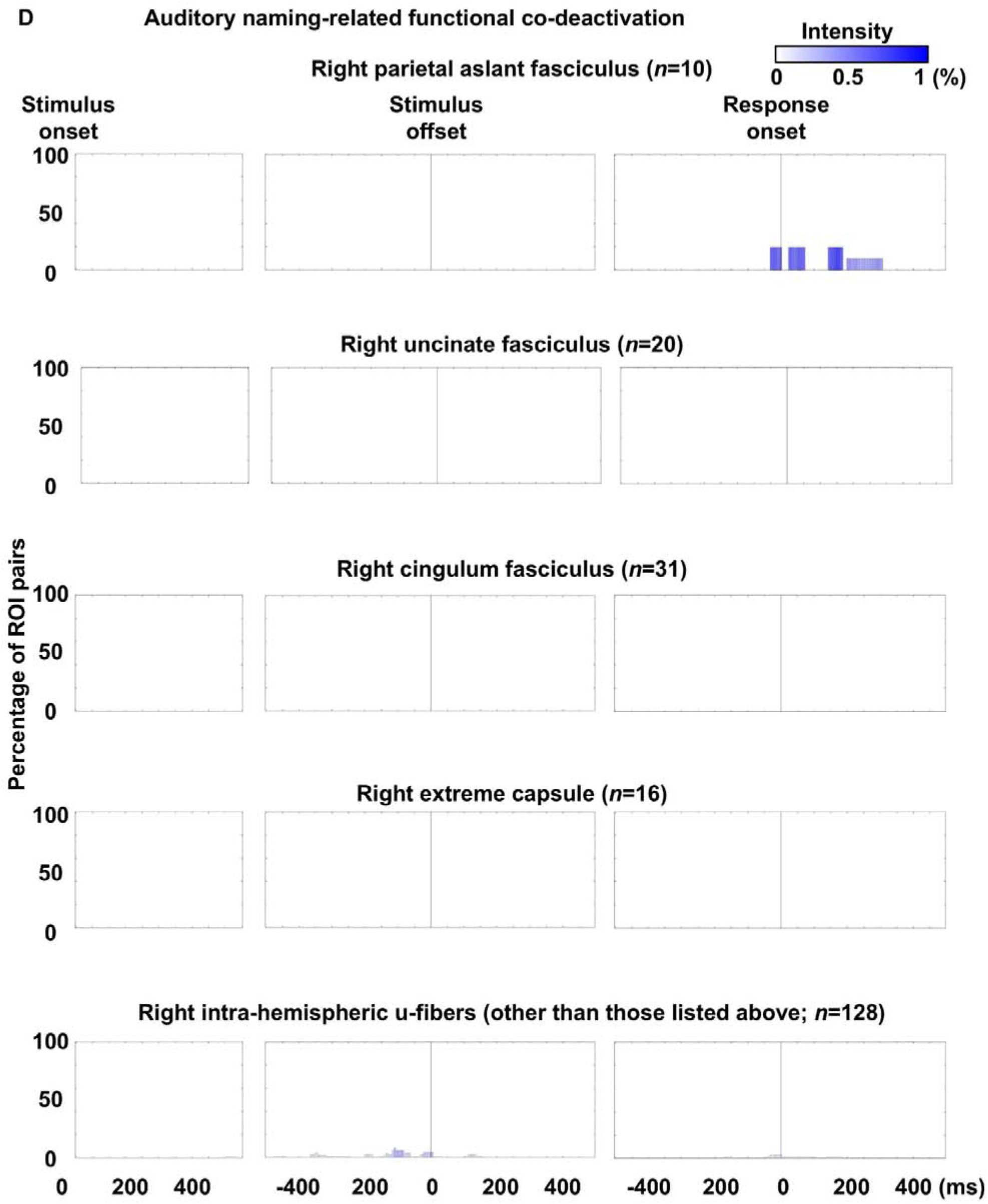
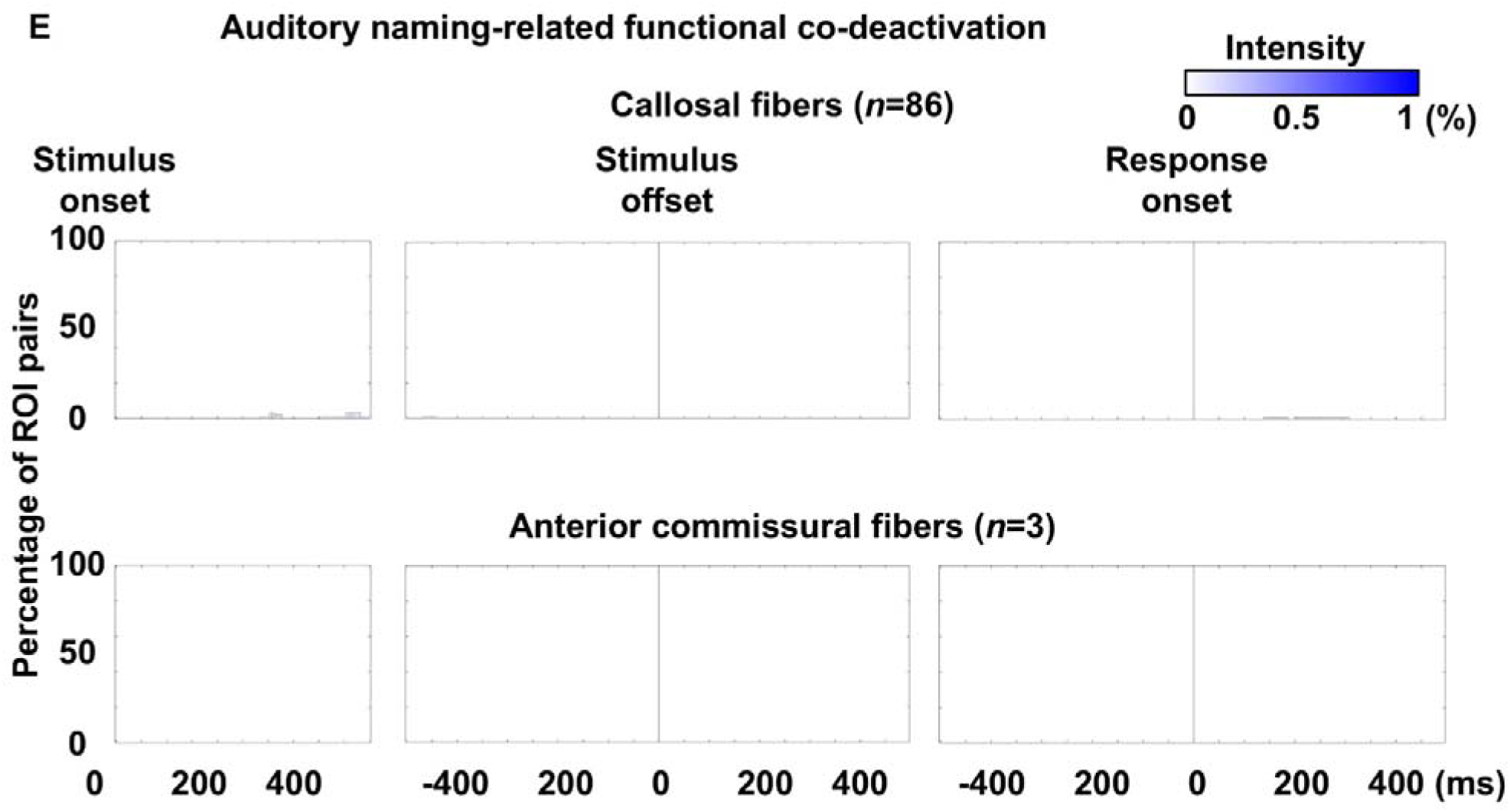
Auditory naming-related functional co-deactivation through each fasciculus. Bar height represents the proportion of white matter pathways within each fasciculus exhibiting functional co-deactivation per time bin. Bar color indicates mean co-deactivation intensity.

**eFigure 14.**
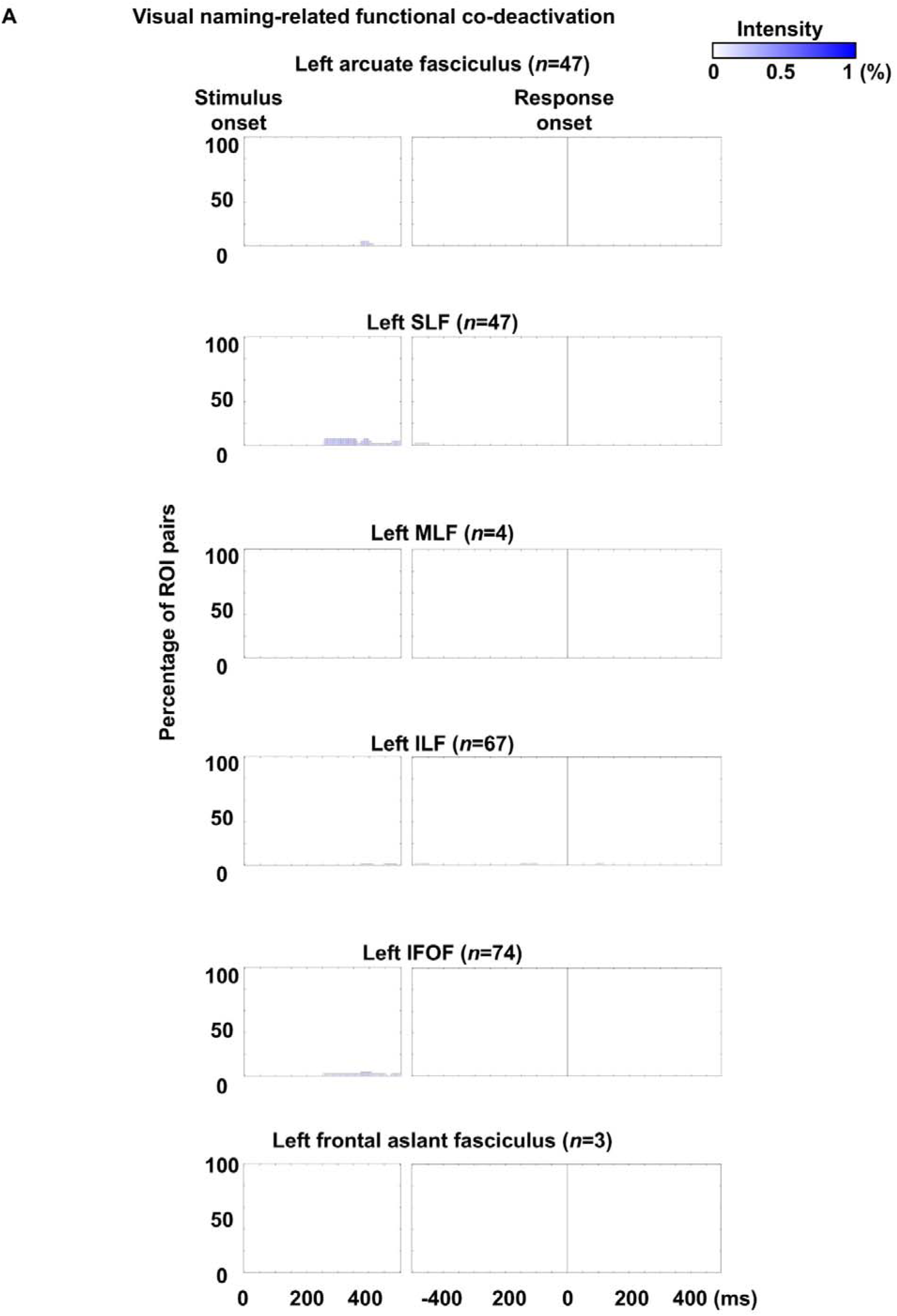
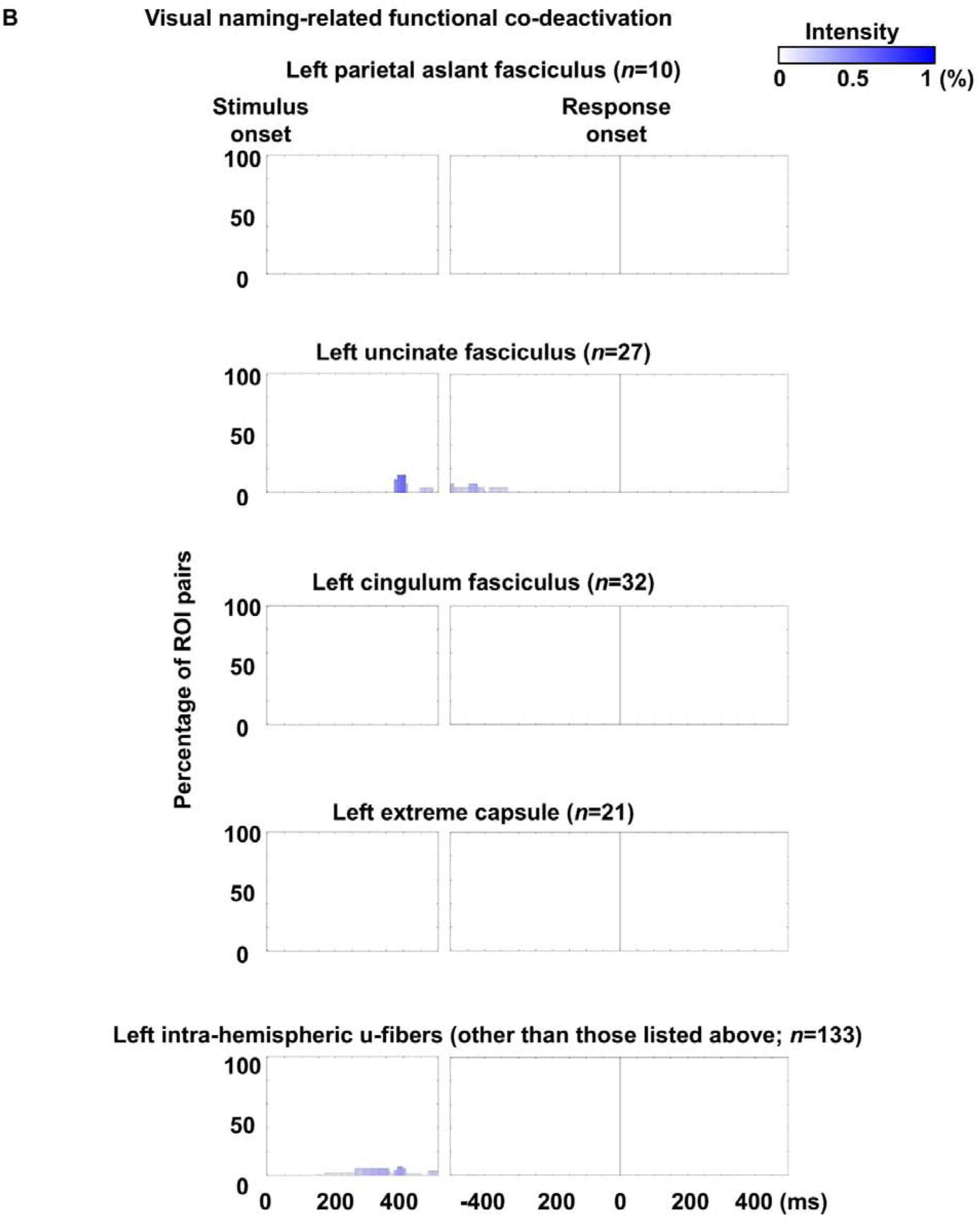
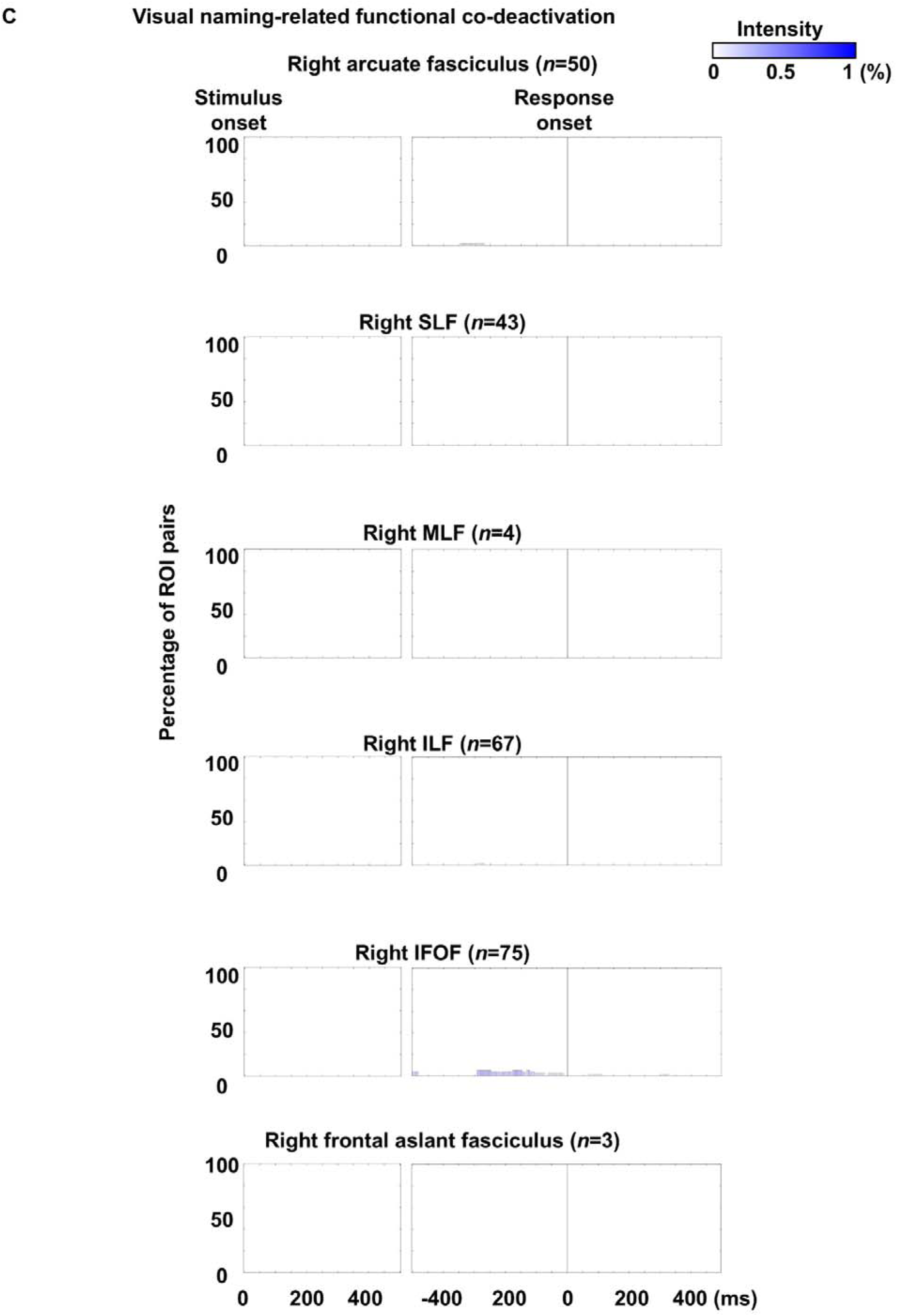
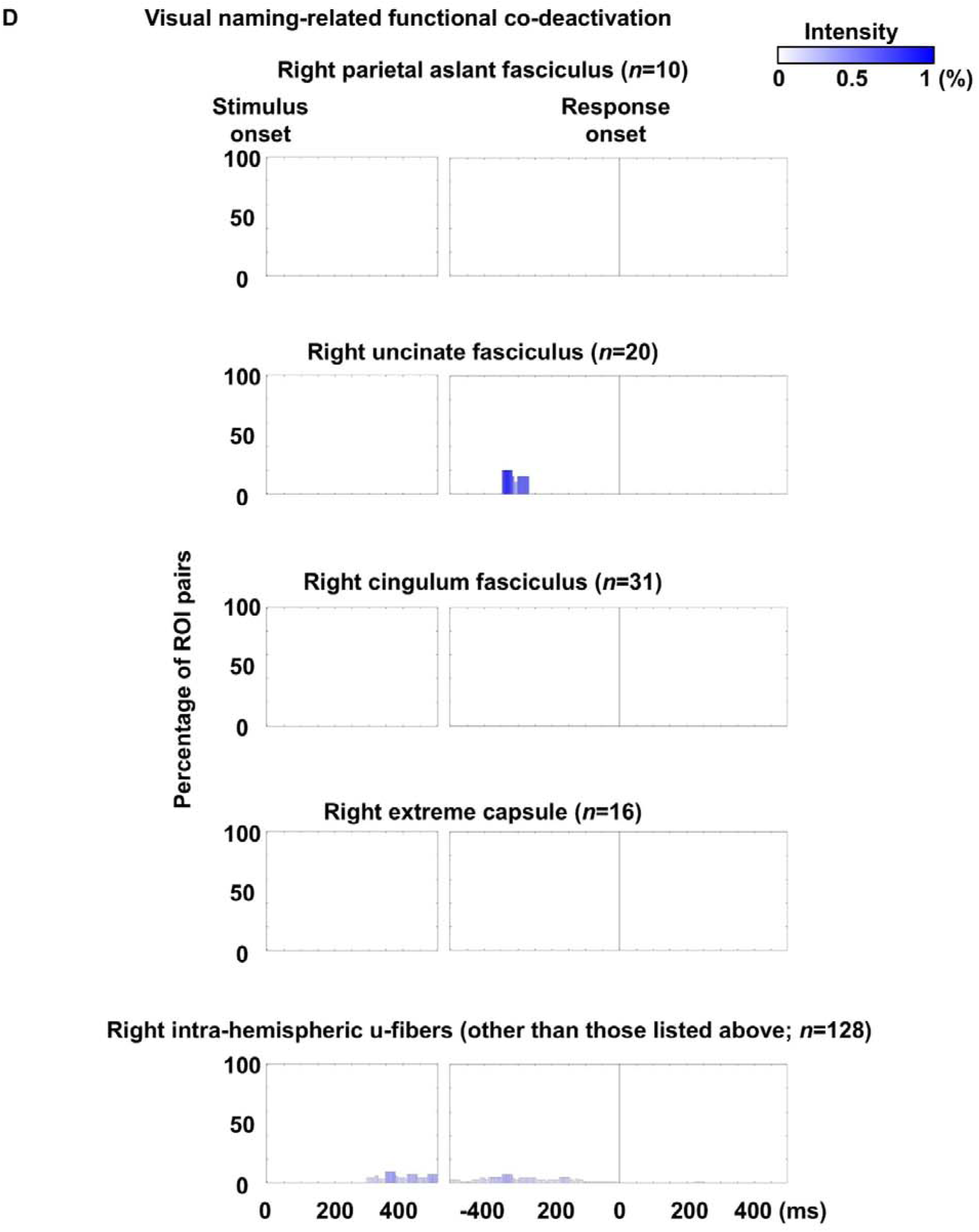
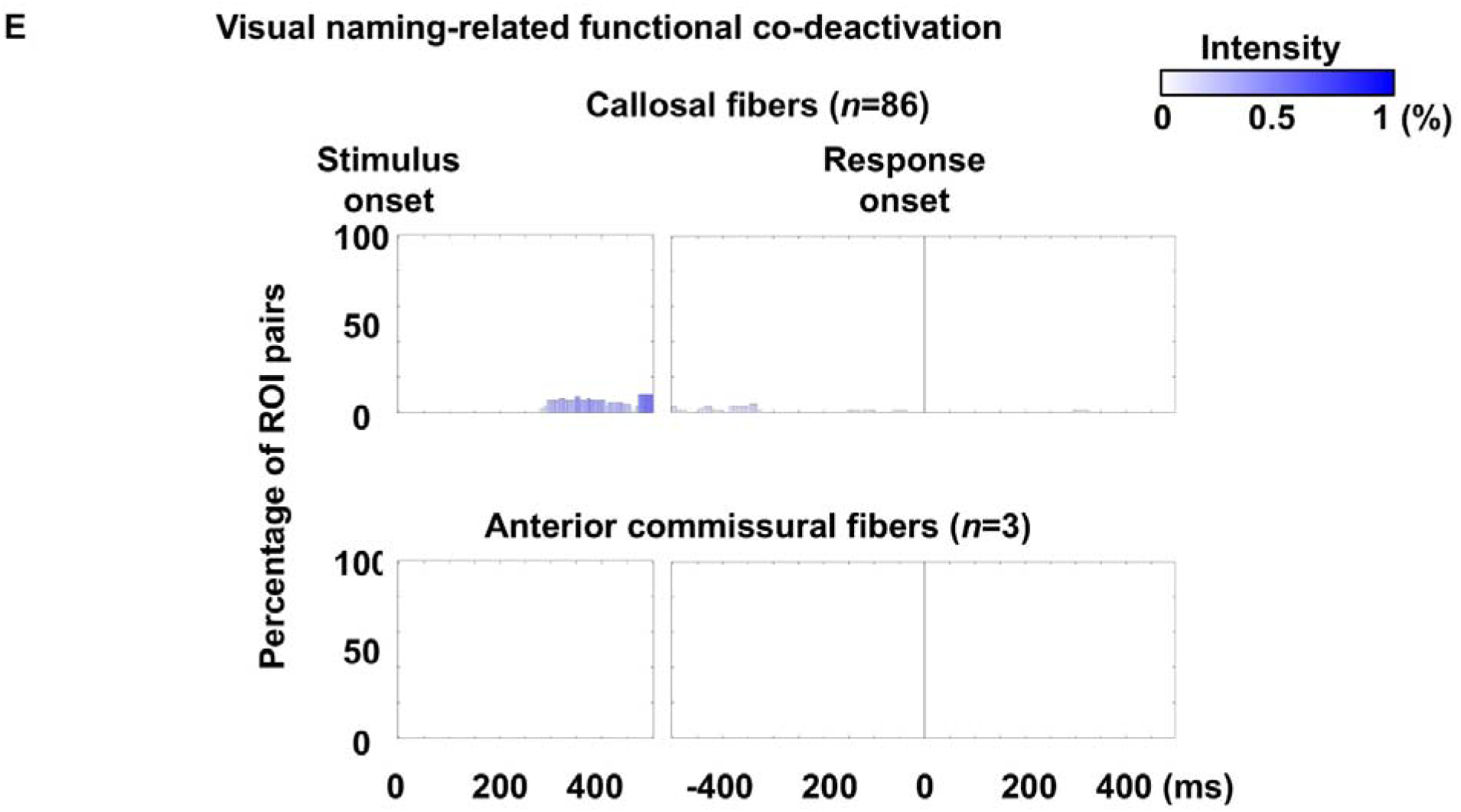
Visual naming-related functional co-deactivation through each fasciculus. Bar height represents the proportion of white matter pathways within each fasciculus exhibiting functional co-deactivation per time bin. Bar color indicates mean co-deactivation intensity.

**eFigure 15.**
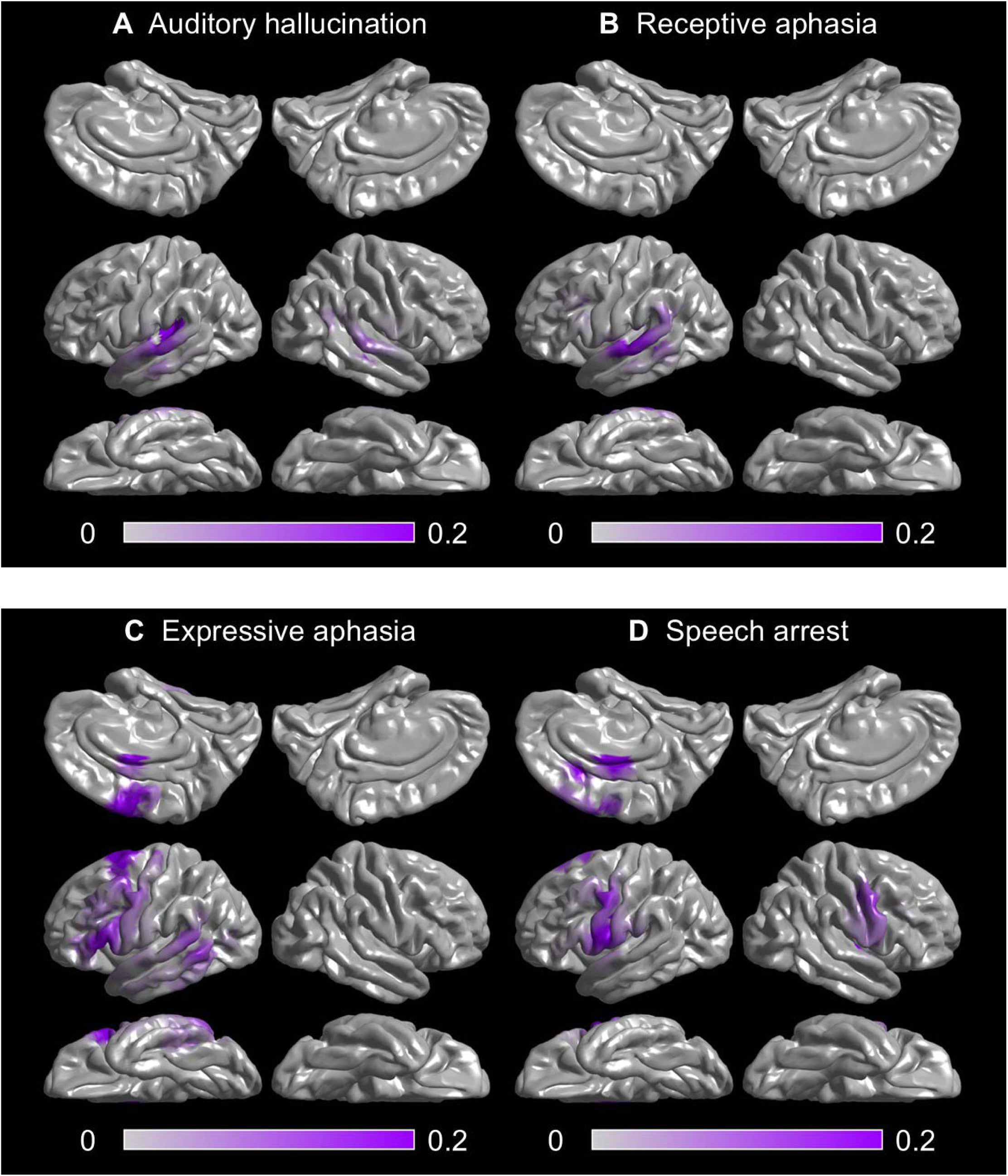
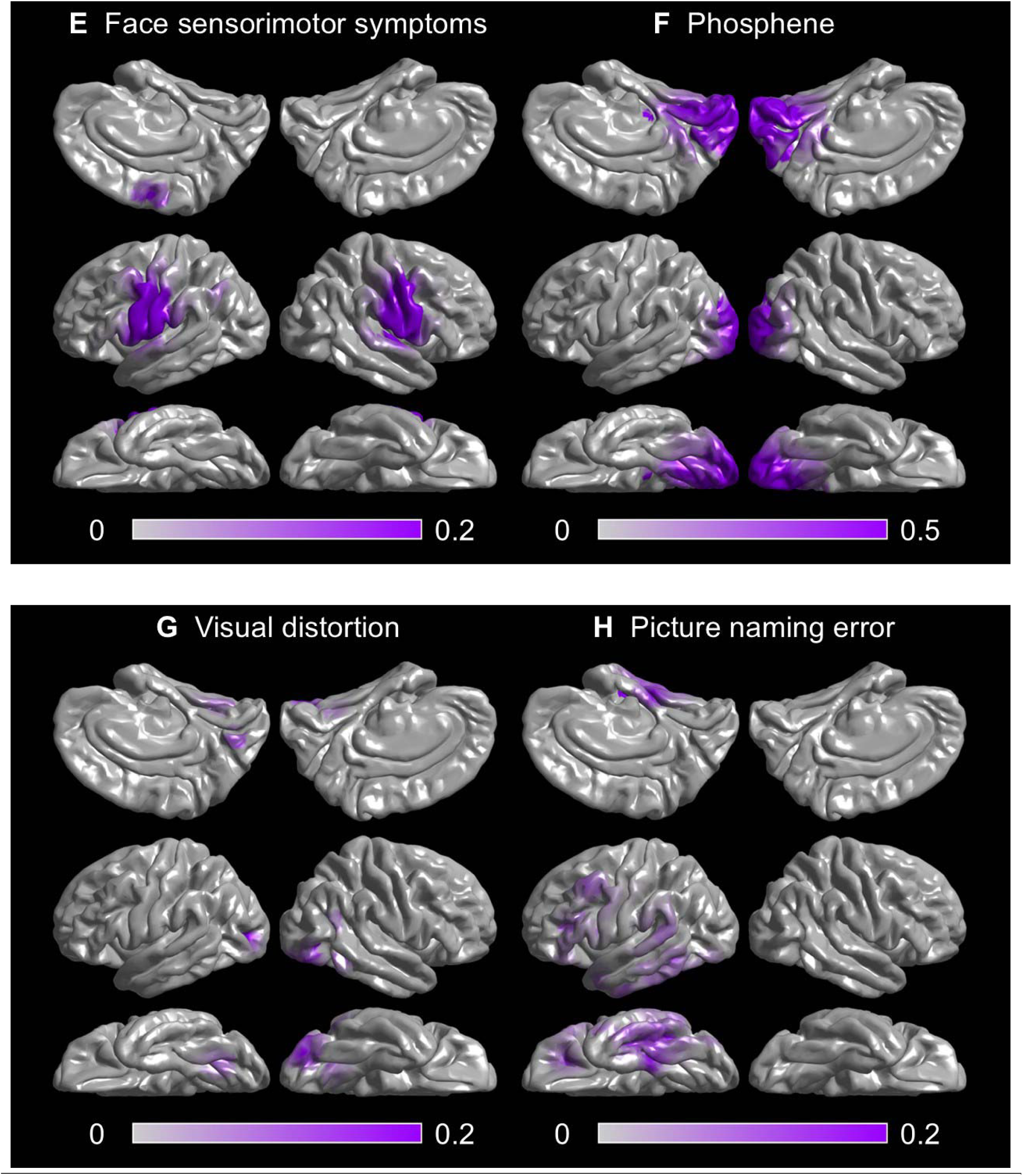
Probability map of stimulation-induced symptoms. **A** Auditory hallucinations. **B** Receptive aphasia. **C** Expressive aphasia. **D** Speech arrest. **E** Face sensorimotor symptoms. **F** Phosphene. **G** Visual distortion. **H** Picture naming error.

**eFigure 16.**
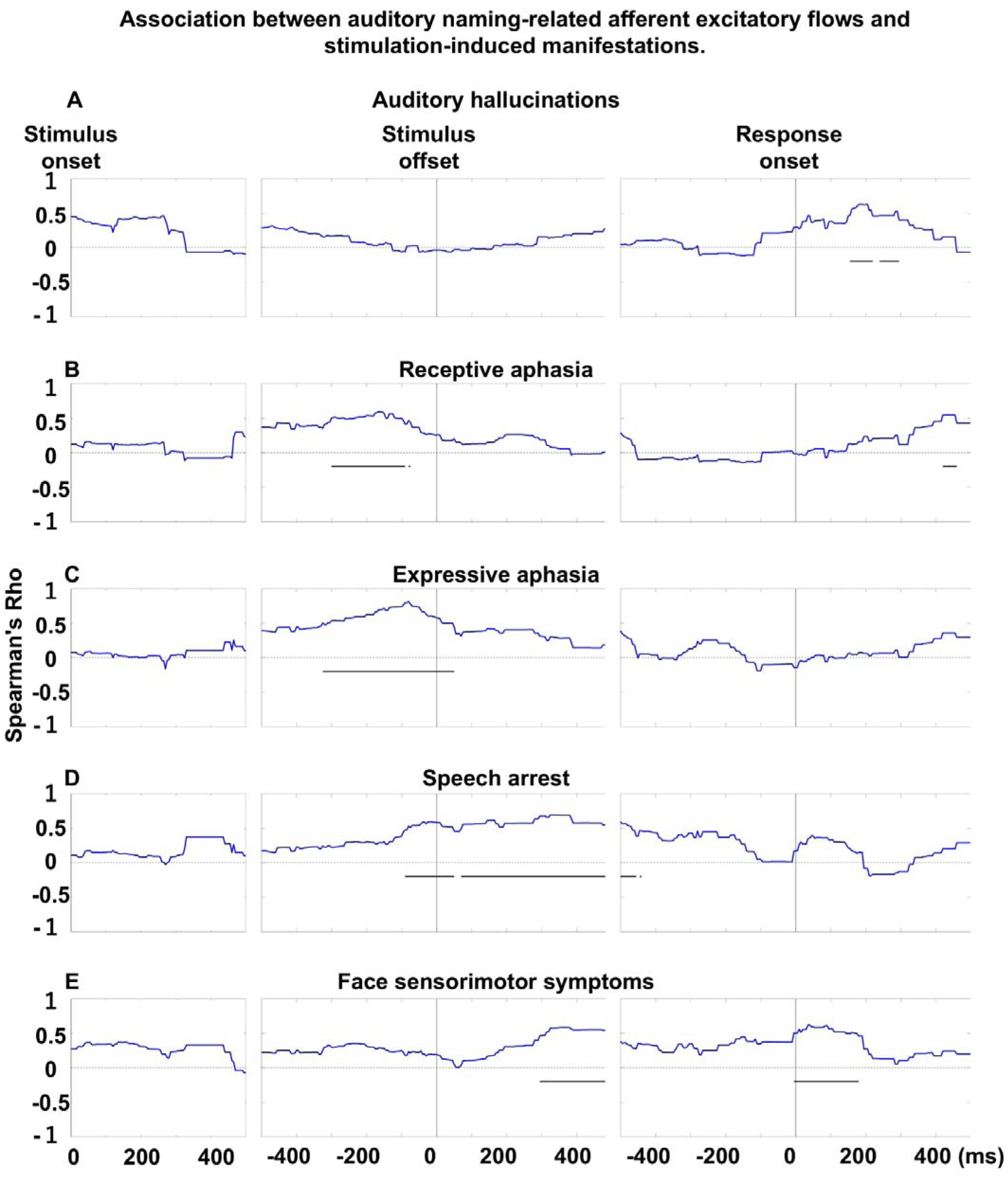
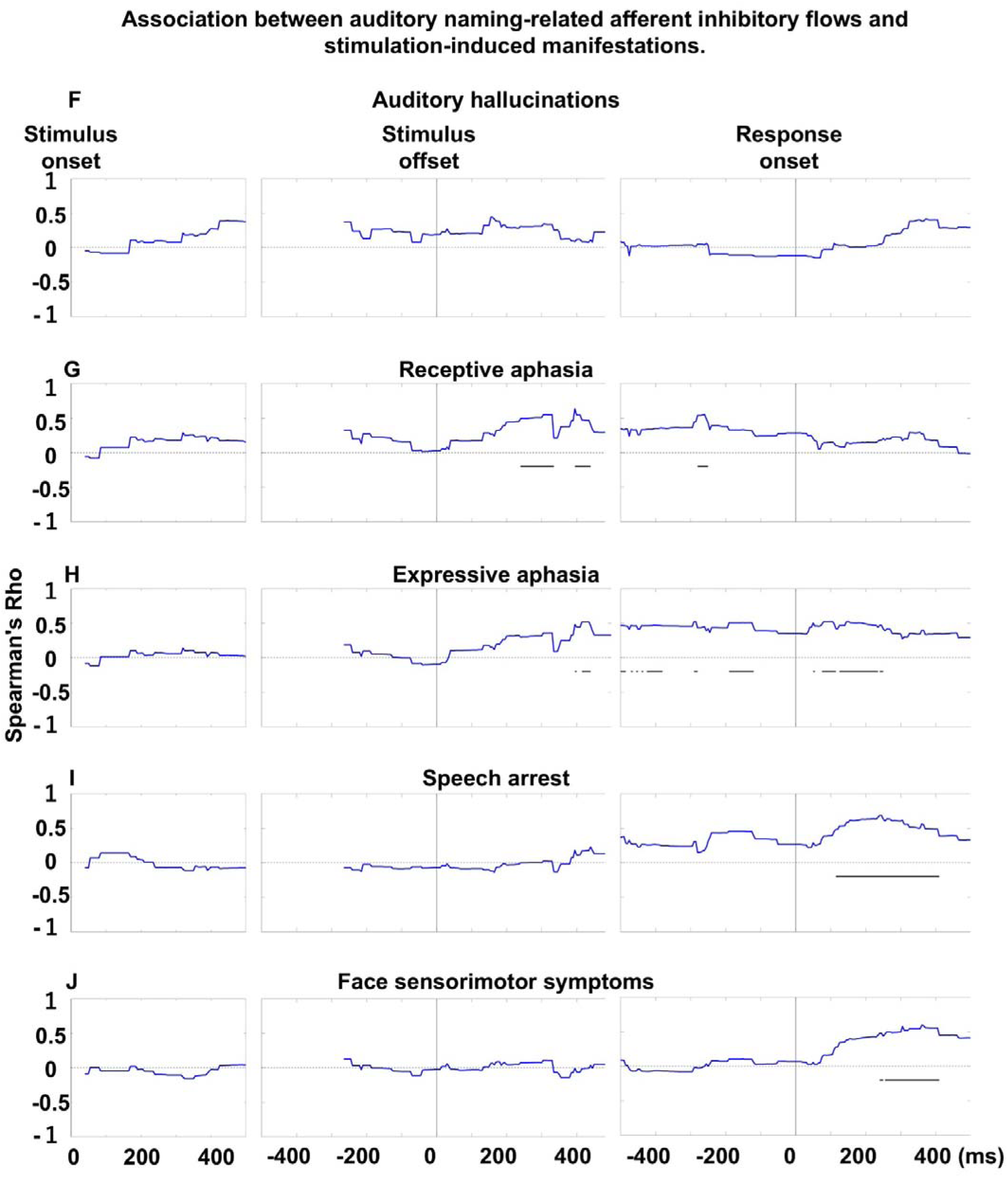
Association between auditory naming–related afferent flows and stimulation-induced manifestations. Each plot shows Spearman’s rho, reflecting the correlation between mean flow intensity and the likelihood of stimulation-induced symptoms across time bins. **A–E** Correlations between excitatory flows and stimulation-induced manifestations. **F–J** Correlations between inhibitory flows and stimulation-induced manifestations. Horizontal bars mark intervals of significant correlation.

**eFigure 17.**
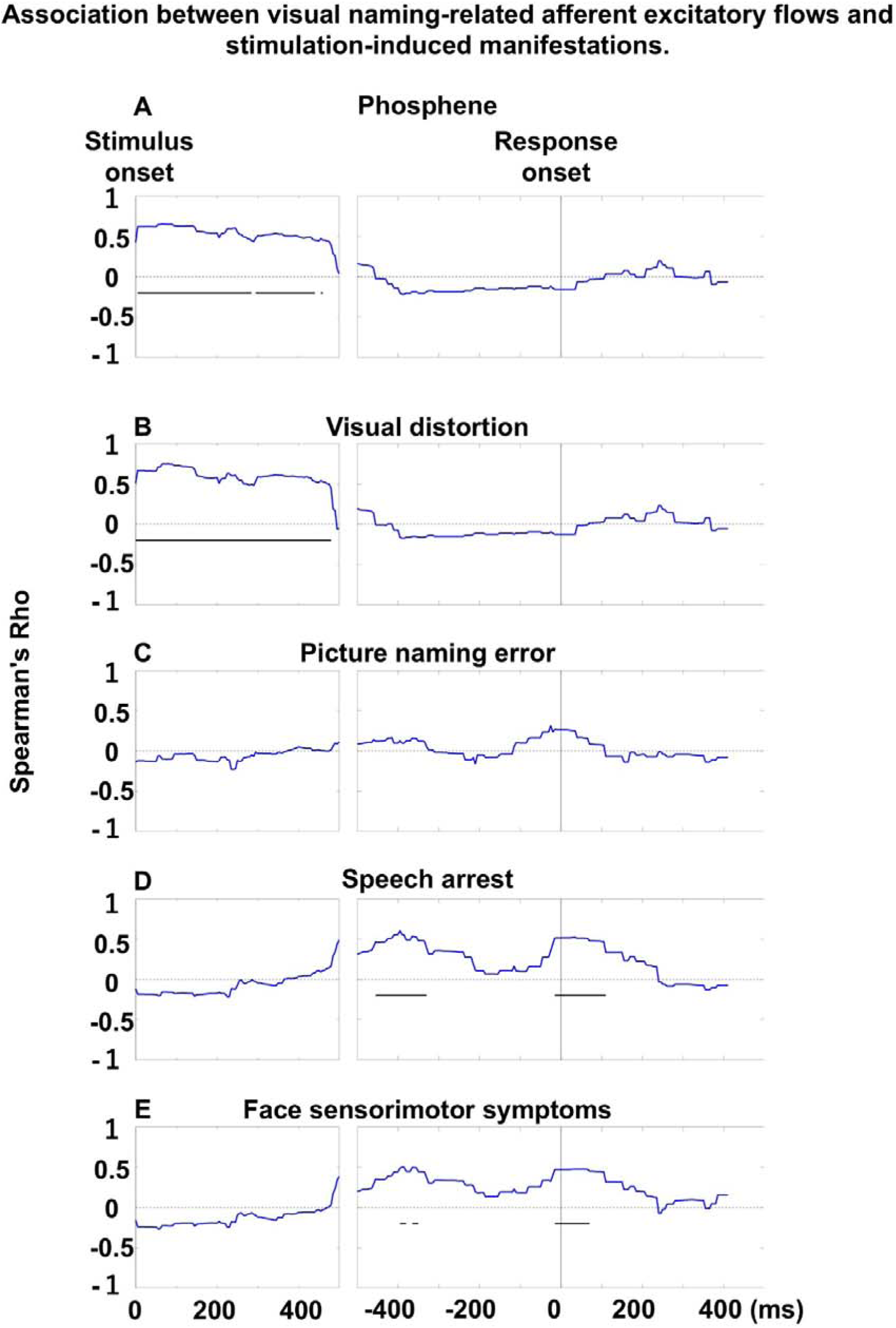
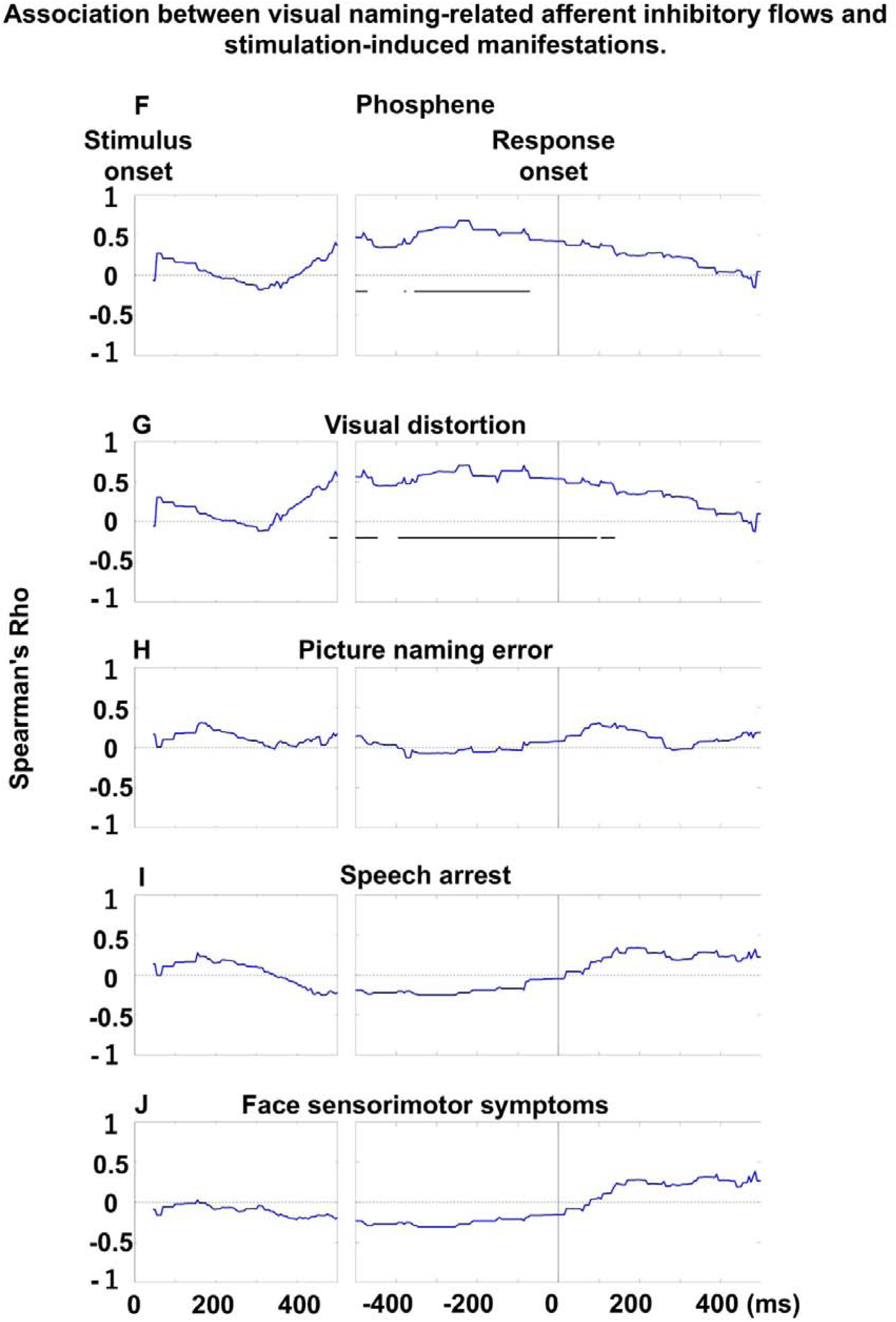
Association between visual naming–related afferent flows and stimulation-induced manifestations. Each plot shows Spearman’s rho, reflecting the correlation between mean flow intensity and the likelihood of stimulation-induced symptoms across time bins. **A–E** Correlations between excitatory flows and stimulation-induced manifestations. **F–J** Correlations between inhibitory flows and stimulation-induced manifestations. Horizontal bars mark intervals of significant correlation.

**eFigure 18.**
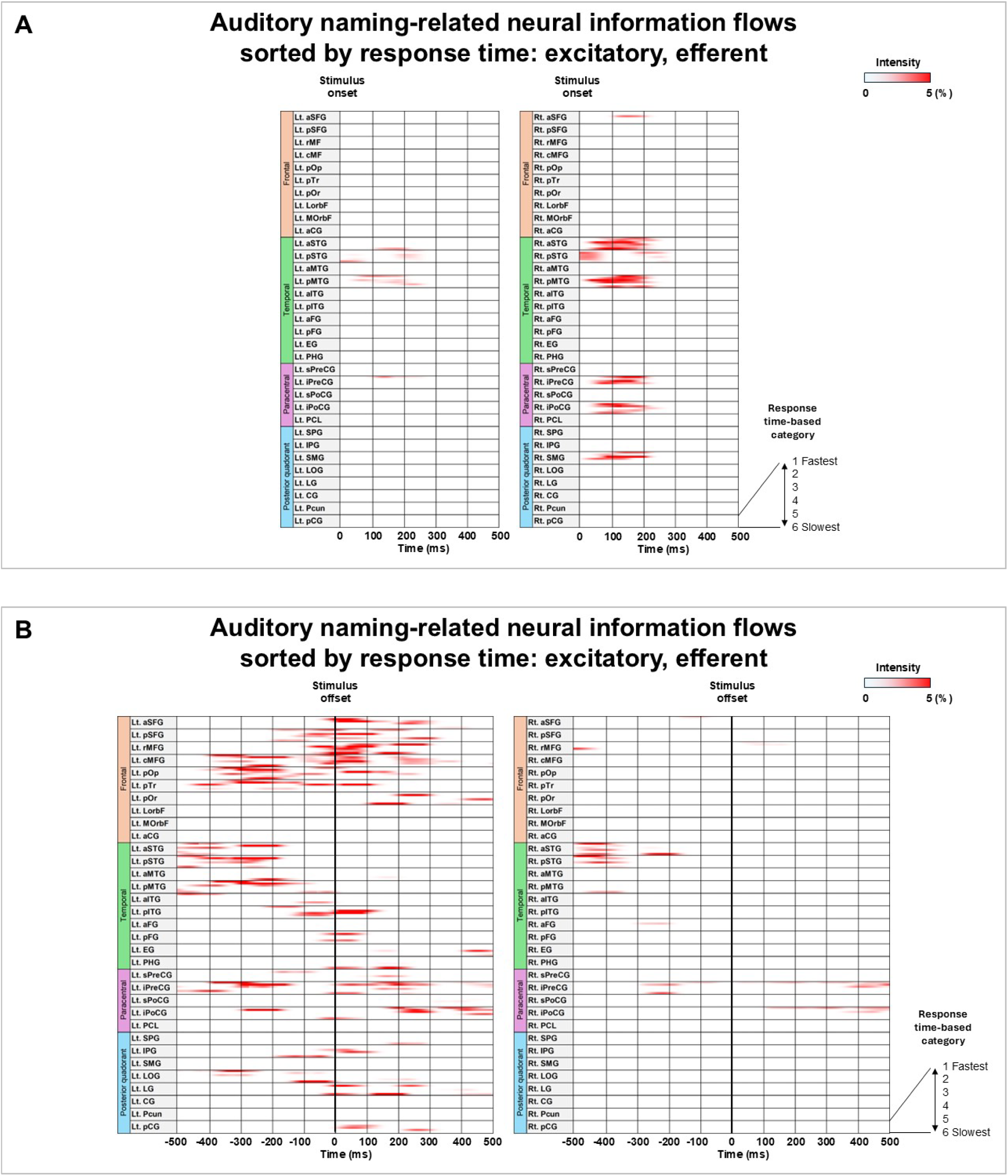
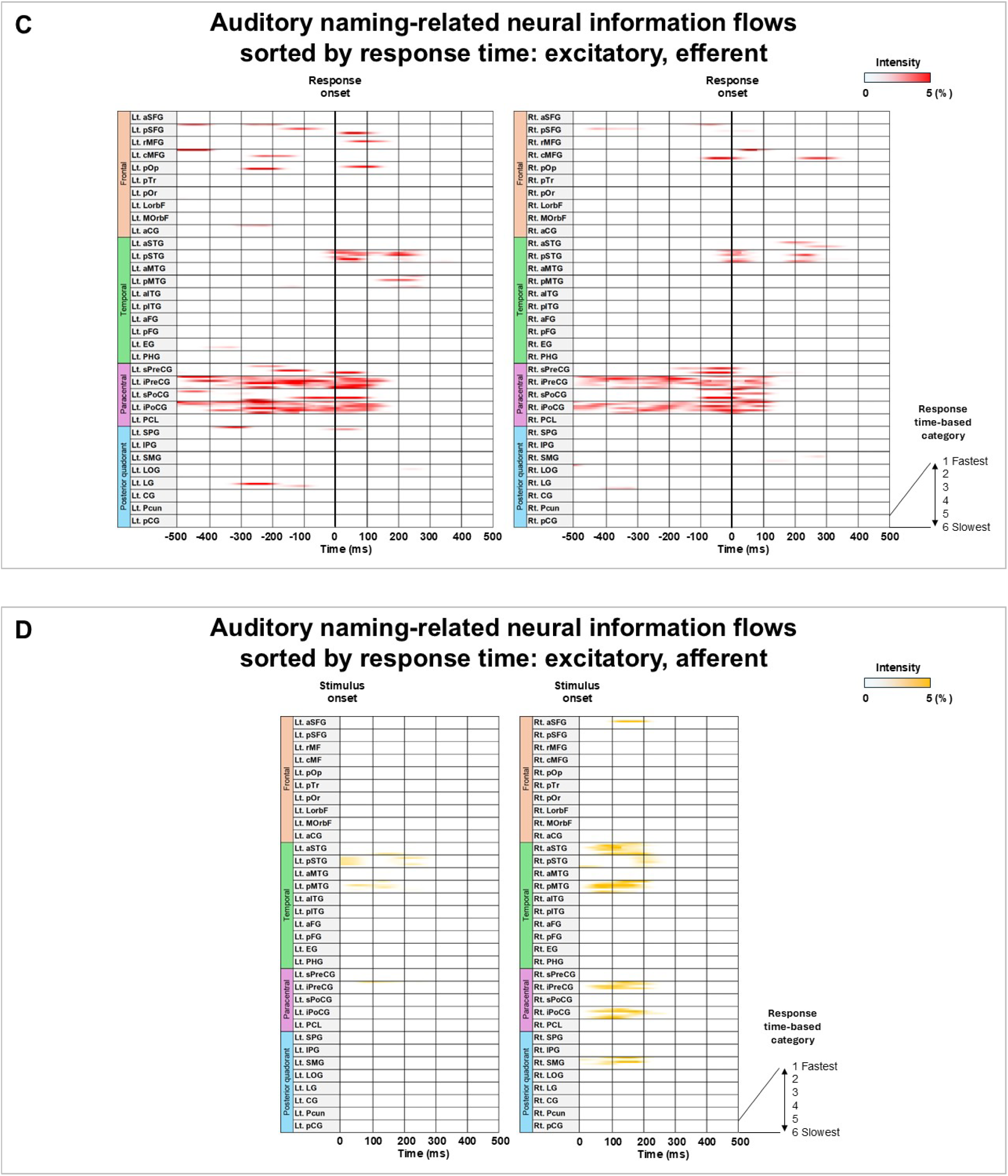
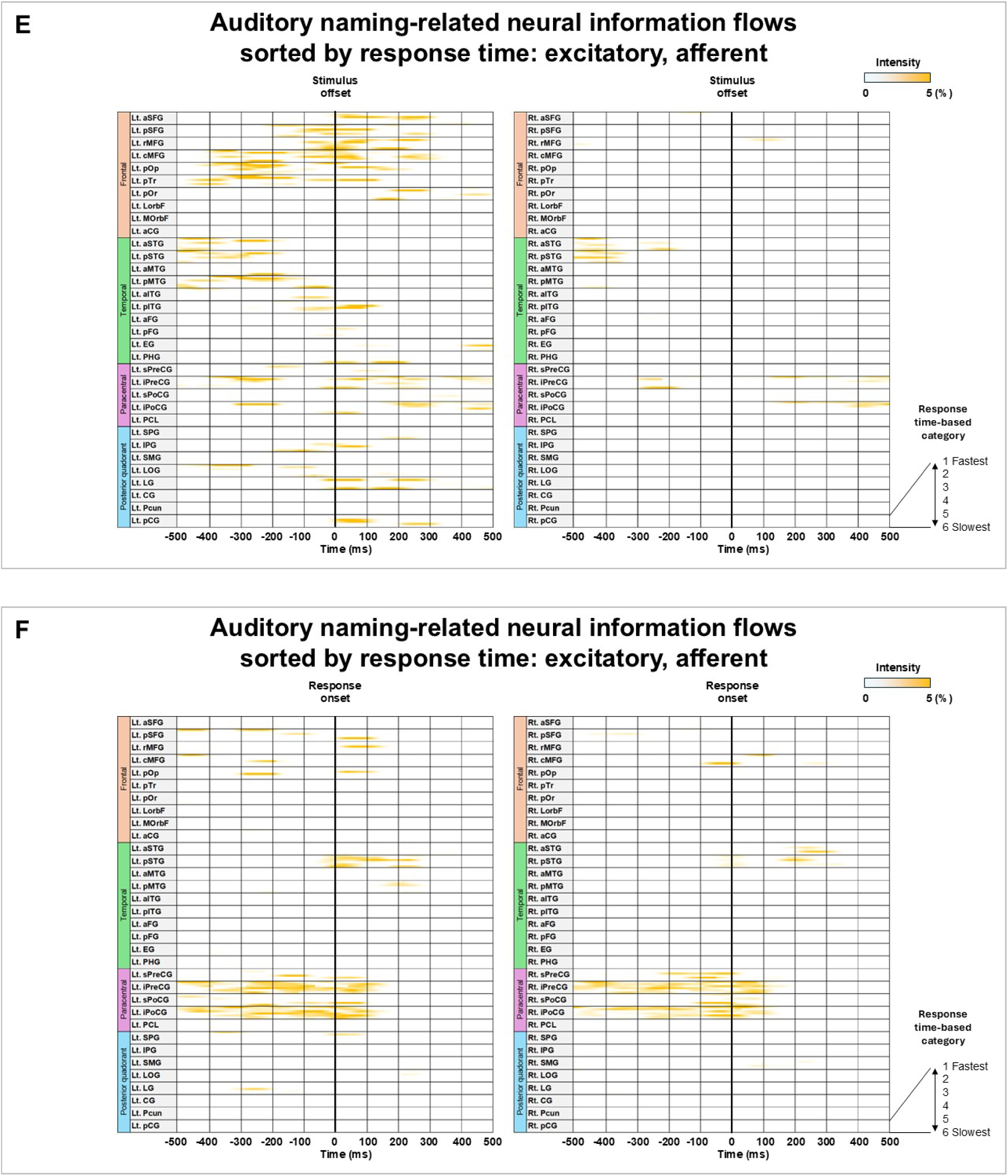
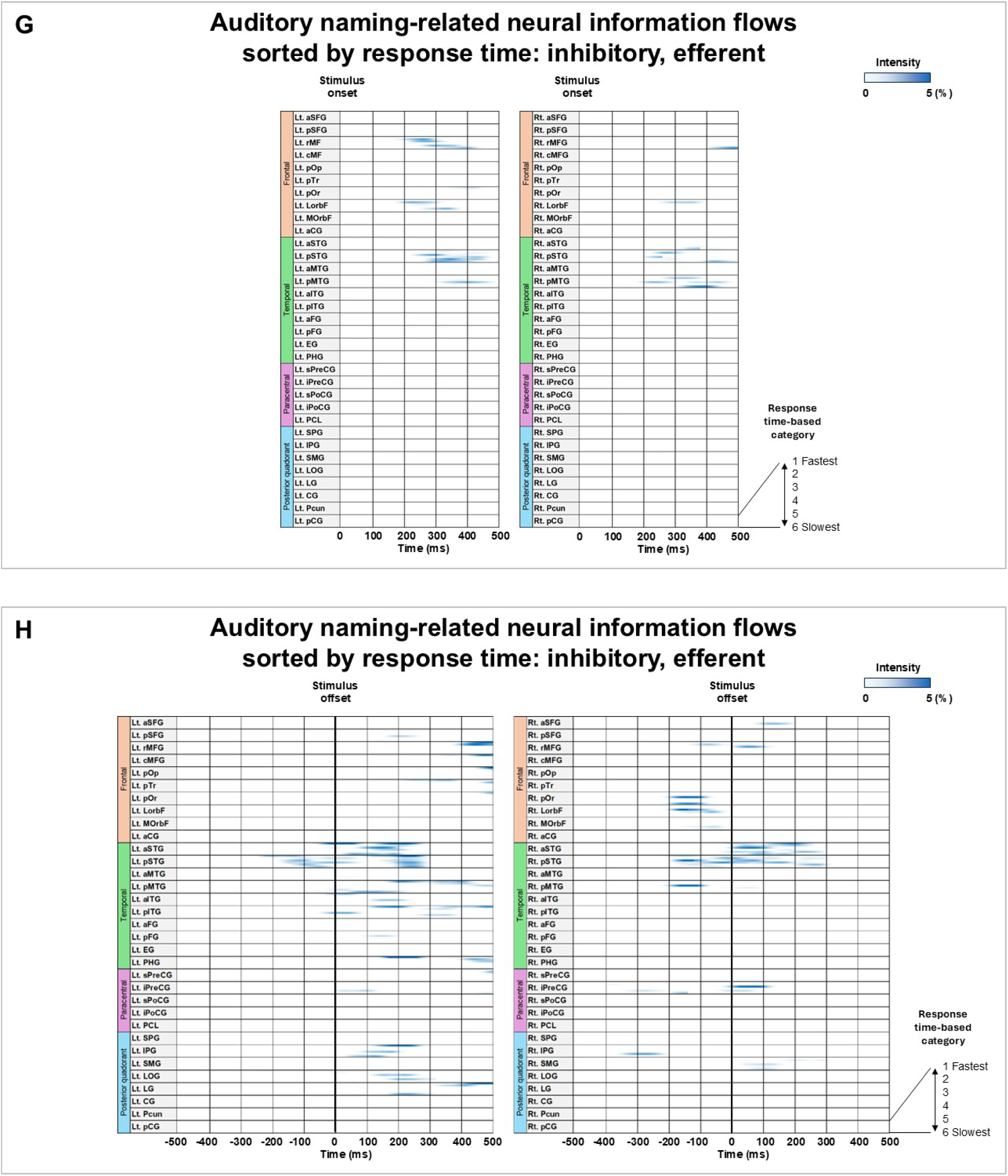
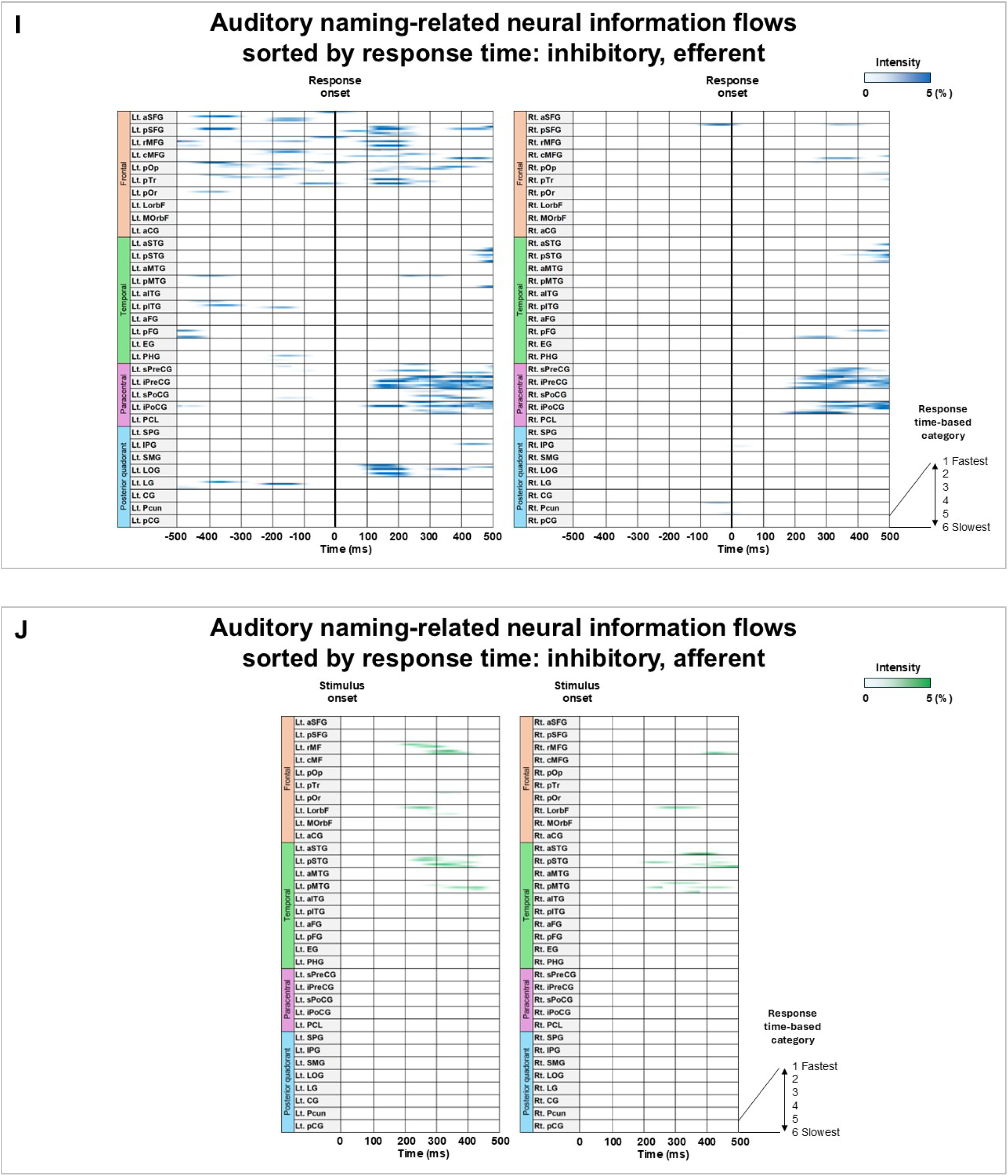
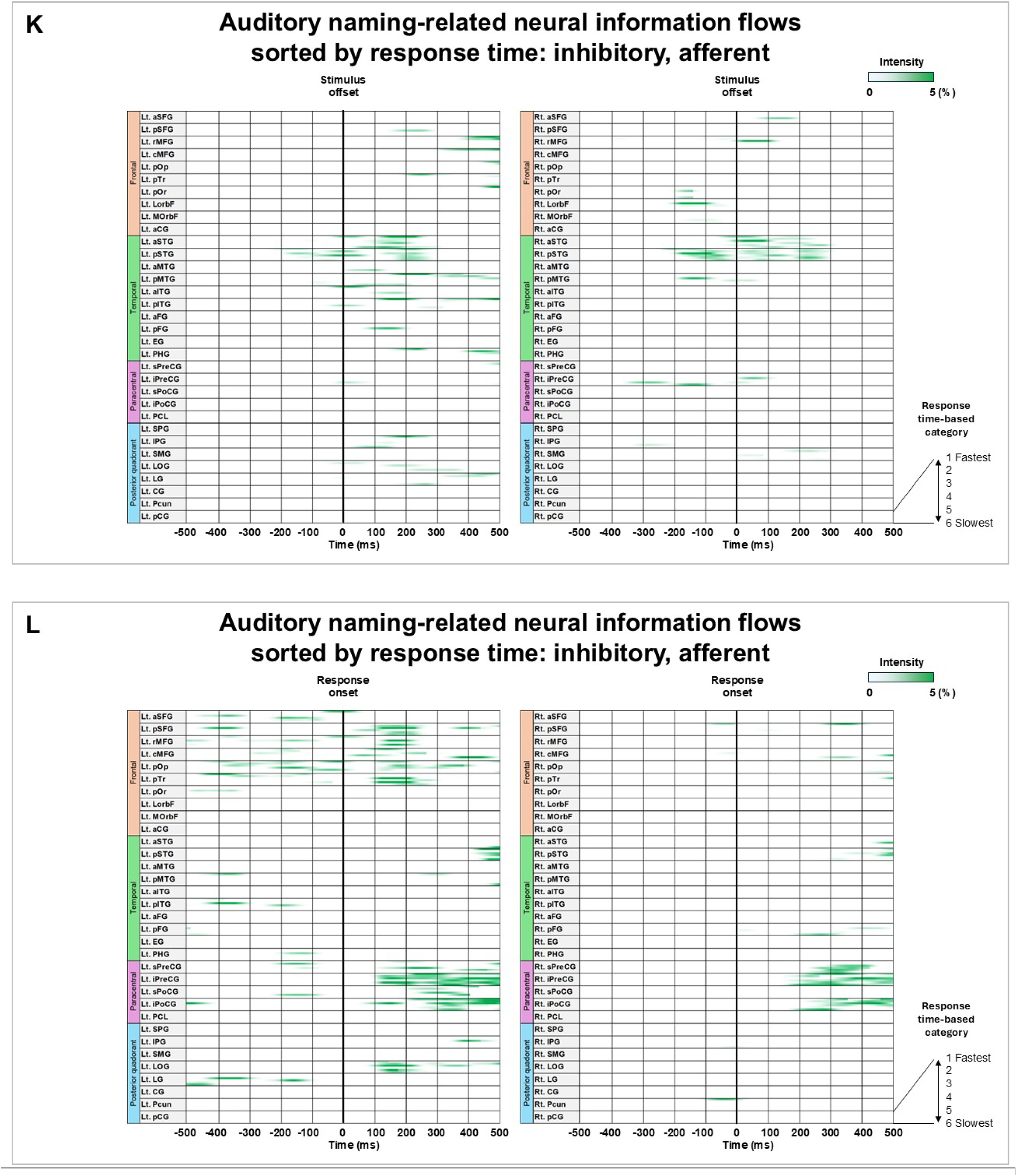
Auditory naming–related neural information flows sorted by response time. Neural information flow intensity is displayed as a function of time. To account for inter-patient variability, each patient’s trials were stratified into six response time categories (fastest to slowest). **A–C** excitatory efferent flow. **D–F** excitatory afferent flow. **G–I** inhibitory efferent flow. **J–L** inhibitory afferent flow.

**eFigure 19.**
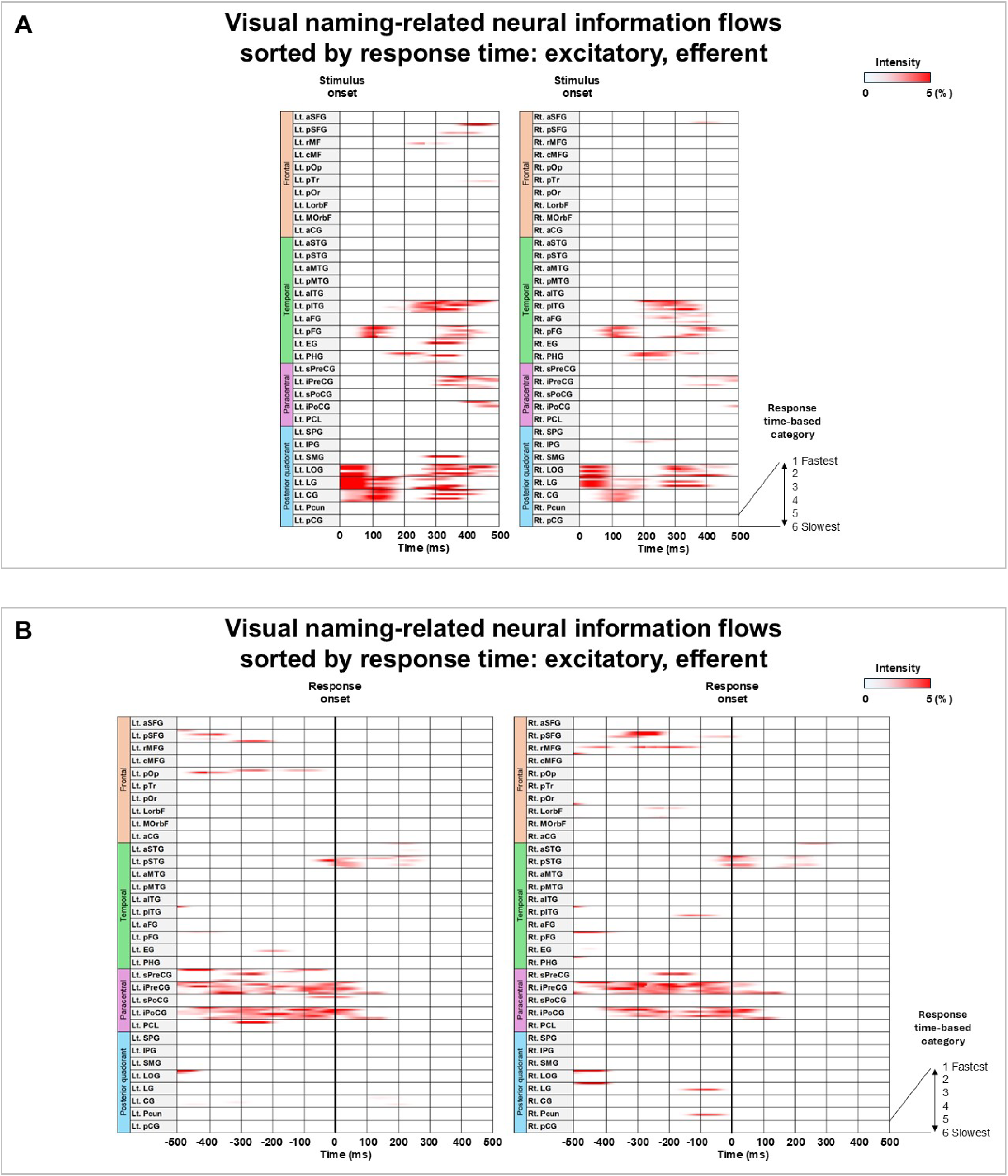
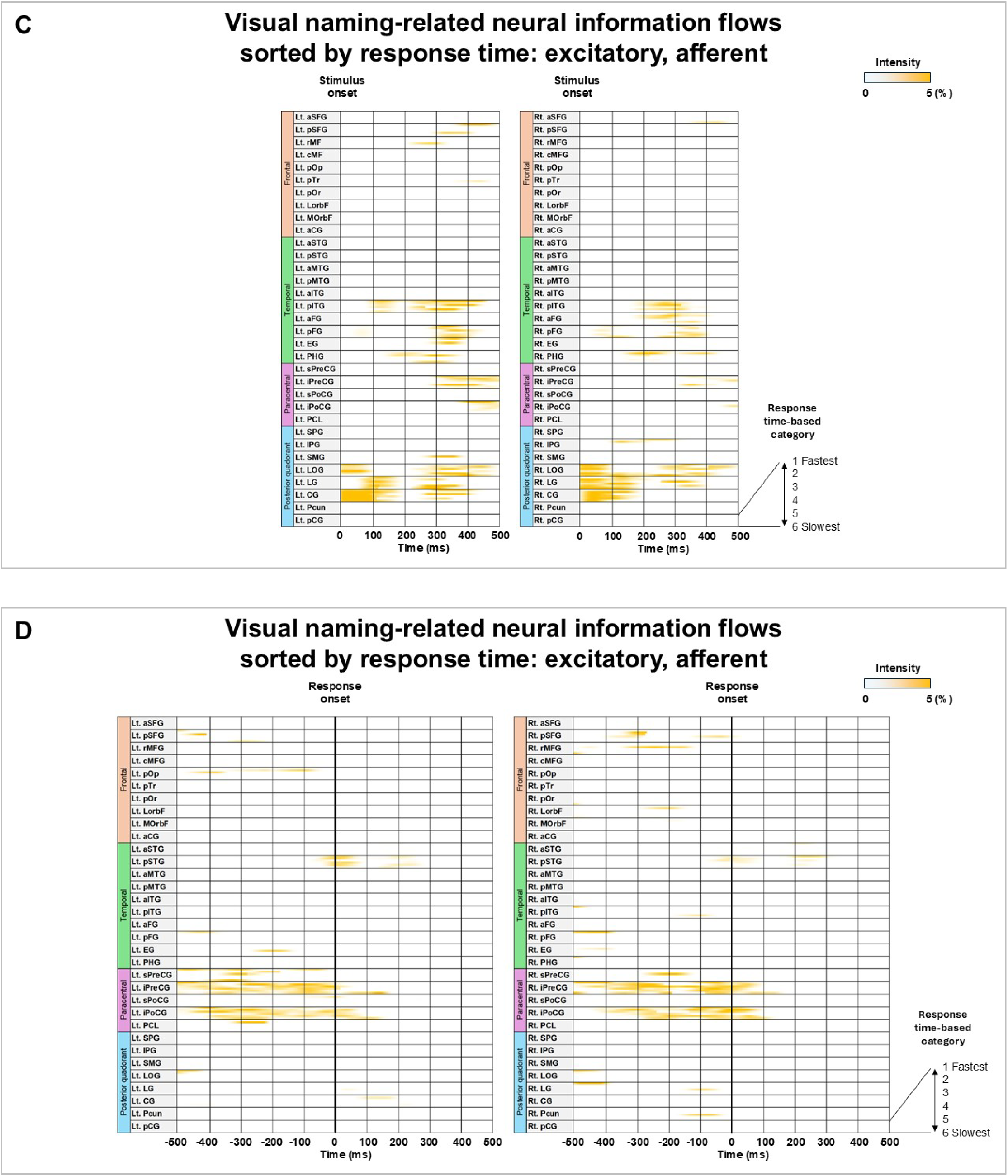
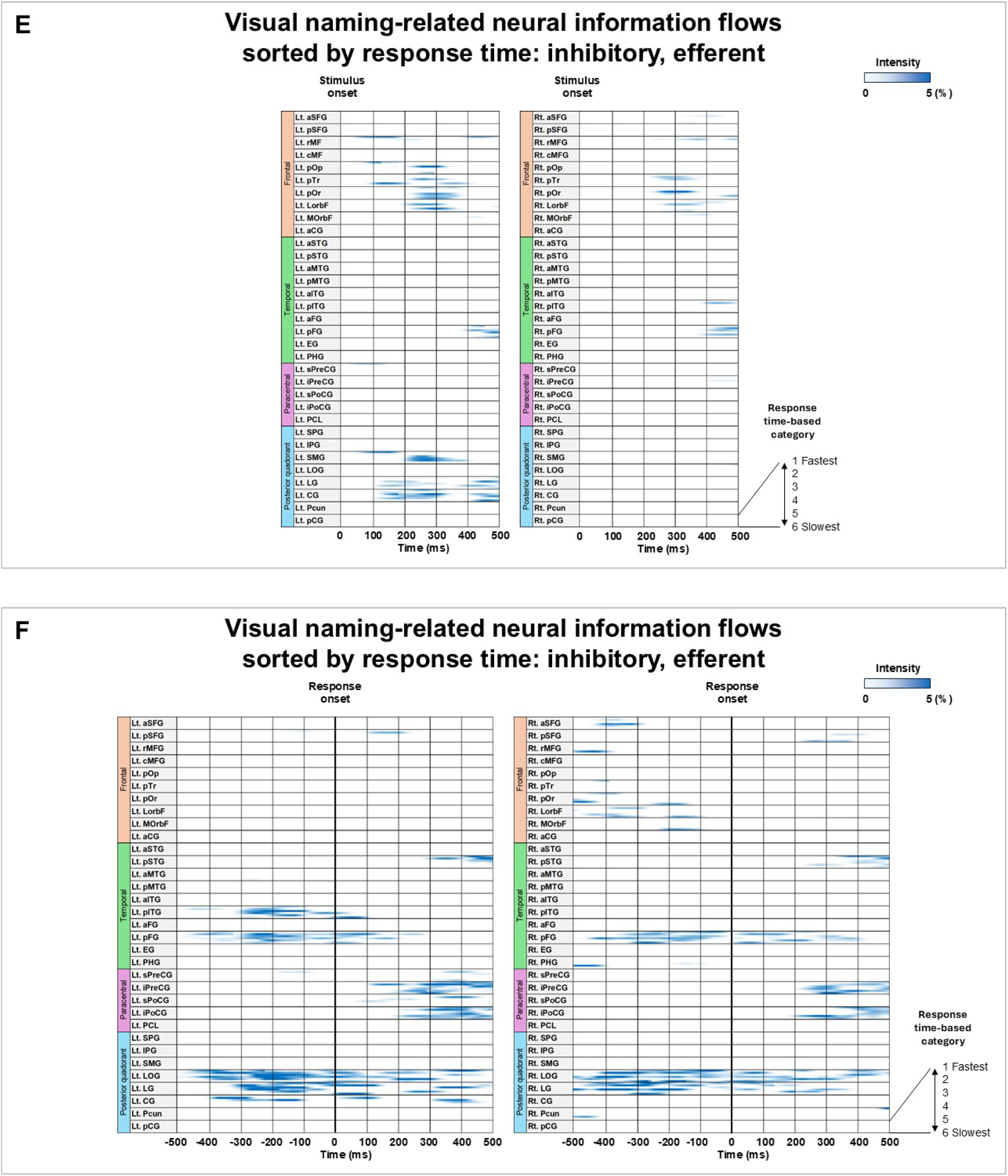
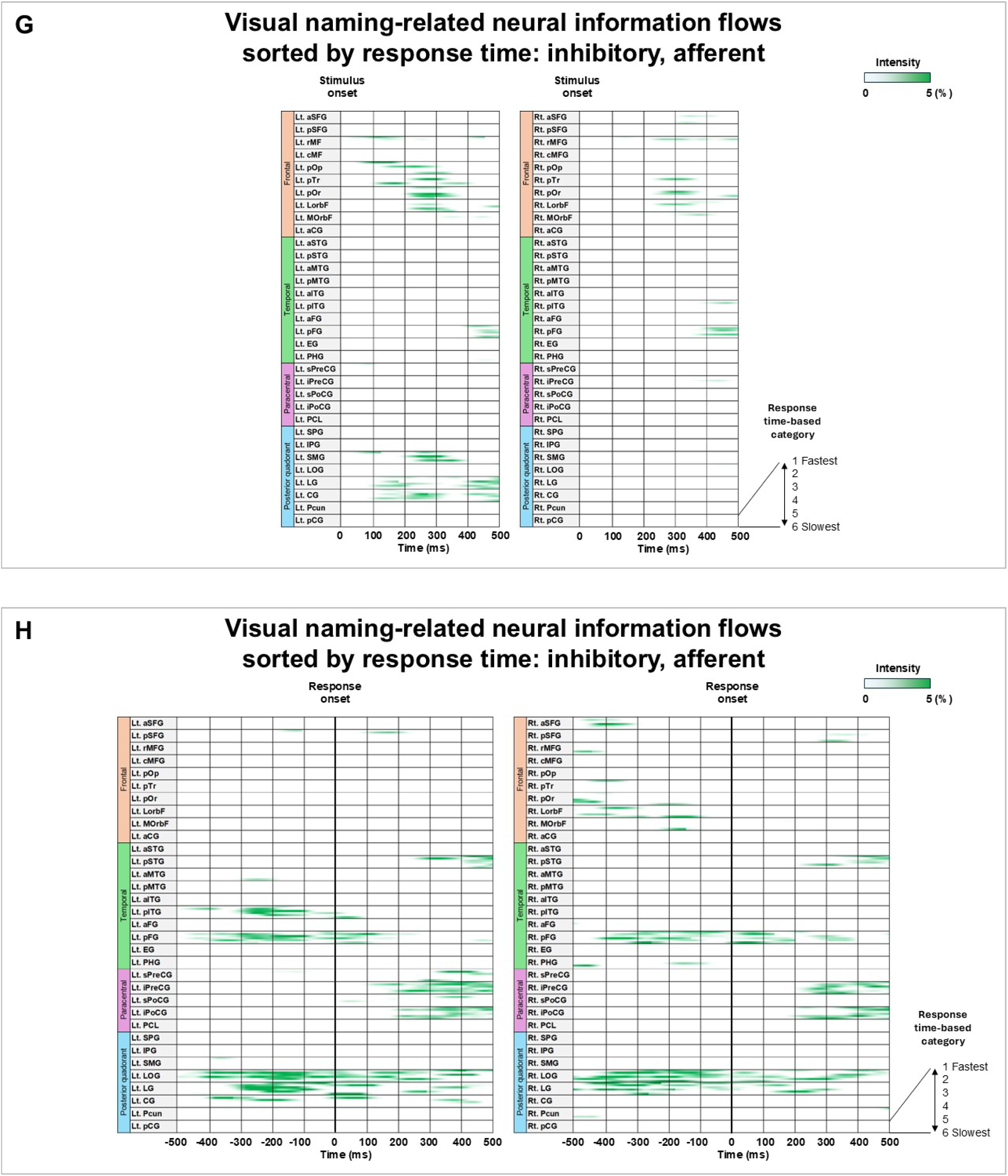
Visual naming–related neural information flows sorted by response time. Neural information flow intensity is displayed as a function of time. To account for inter-patient variability, each patient’s trials were stratified into six response time categories (fastest to slowest). **A–B** excitatory efferent flow. **C–D** excitatory afferent flow. **E–F** inhibitory efferent flow. **G–H** inhibitory afferent flow.

## Supplementary Results: eResults

### Effects of patient profiles on response times and neural dynamics

We implemented mixed-effects modeling to evaluate how the following variables affected response latency and high-gamma activity. Fixed effect predictors included the square root of age (√years), handedness, left-hemispheric epileptogenicity, presence of MRI-detectable lesions, and the number of antiseizure medications, with polytherapy used as an index of greater cognitive burdens. Median response time served as the dependent variable, with intercept and patient included as random effects. Analyses were performed in SPSS version 28 (IBM, Chicago, IL), and statistical significance was defined as two-sided p < 0.05. To evaluate neural responses, the same framework was applied to median high-gamma amplitudes across electrodes within each ROI, calculated in 100-ms bins from stimulus presentation through 500 ms after response, within a 2,500-ms analysis window. Significance thresholds were adjusted using Bonferroni correction (p < 0.05 across 66 ROIs and 25 time bins).

We found that increased √age was associated with shorter median response times in auditory (mixed model estimate: –0.20; 95%CI: –0.32 to –0.08; p-value: 0.001) and picture naming tasks (estimate: –0.19; 95%CI: –0.32 to –0.07; p-value: 0.003). A higher number of antiseizure medications was independently associated with a longer median response time in the auditory naming task (estimate: 0.14; 95%CI: 0.01 to 0.26; p-value: 0.03). These effects were independent of other fixed effect variables, none of which showed significant associations (p-values: 0.08–0.99).

The associations between patient profiles and regional high-gamma amplitude observed in our previous report (Kochi et al., in press) were identically replicated in the present cohort of 127 patients. Specifically, left-handedness was linked to a 15.1% increase in auditory naming–related high-gamma amplitude within the right entorhinal cortex (300–400 ms after stimulus offset; 95% CI: 9.9–20.3%; p = 7.8 × 10). Left-hemispheric epileptogenicity corresponded to a 29.6% increase in picture naming–related high-gamma amplitude at the right posterior superior temporal gyrus (0–100 ms before response onset; 95% CI: 17.7–41.5%; p = 6.6 × 10). No other fixed variables exhibited significant effects.

eReference: Kochi R, Kanno A, Uda H, Hatano K, Endo H, Cools M, Rothermel R, Luat AF, Asano E. Whole-brain dynamic causal tractography atlas of auditory and visual speech networks derived from intracranial neurophysiological data. Neurology (in press)

## Supplementary Tables: eTables

**eTable 1.**
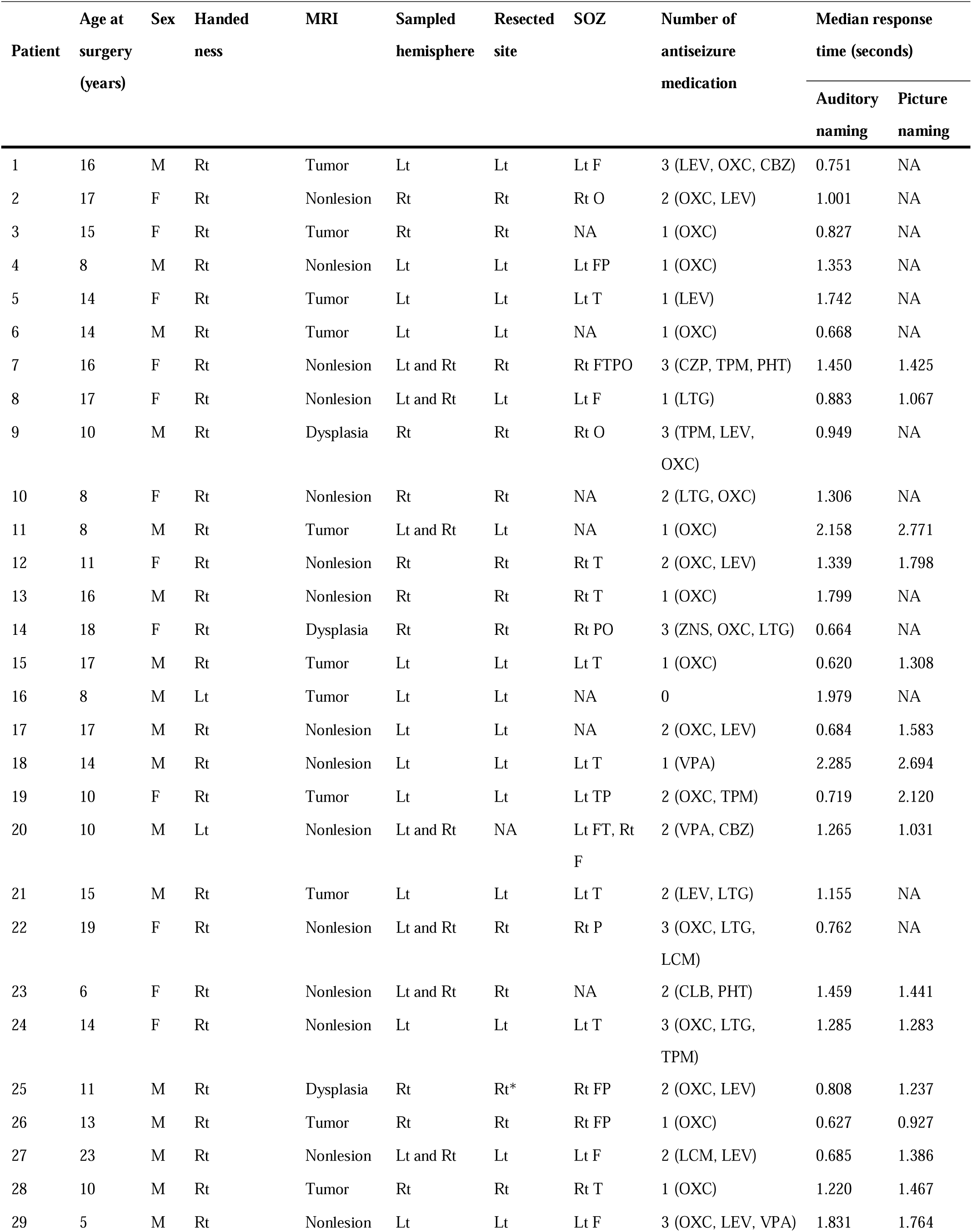
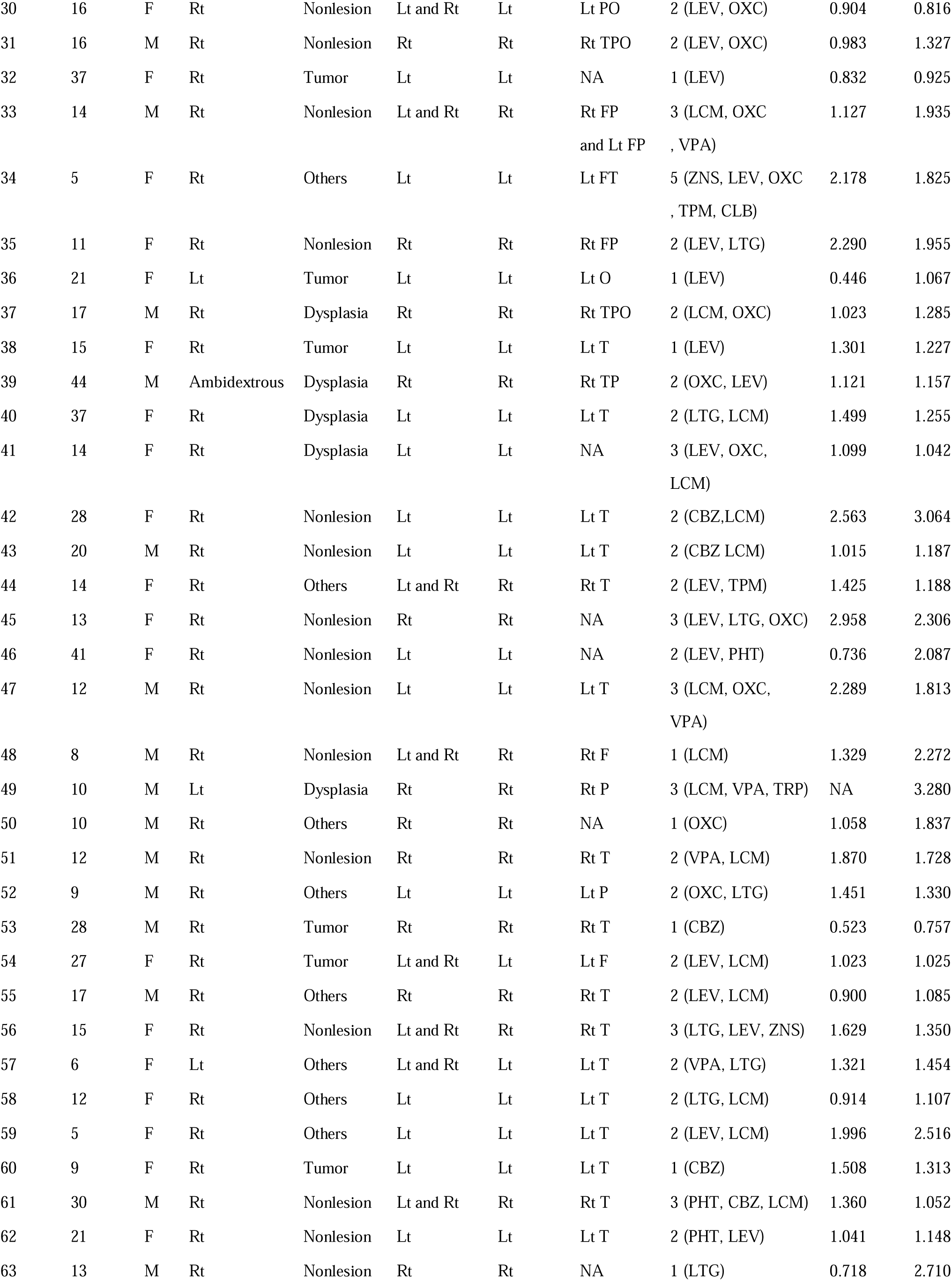
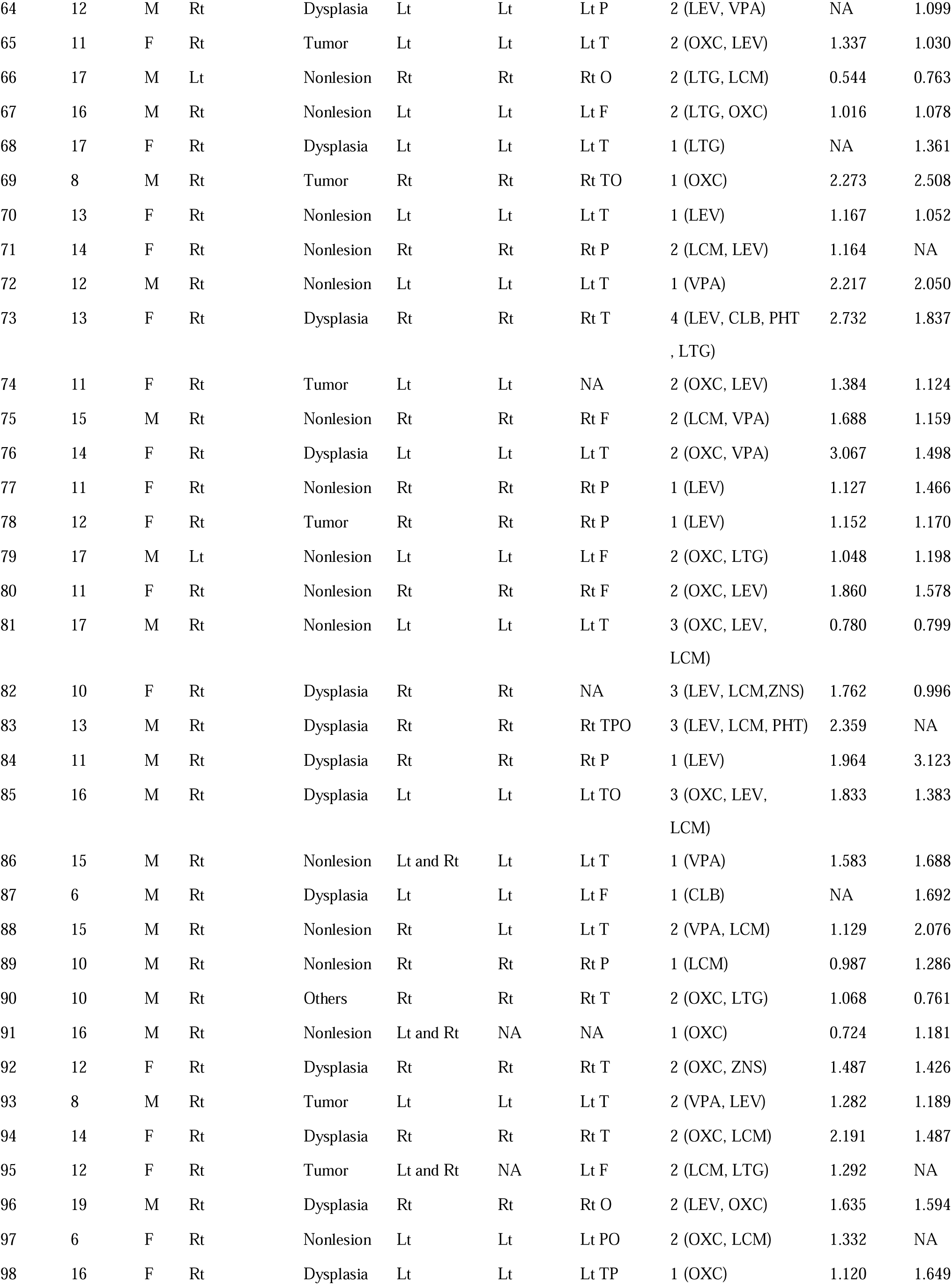
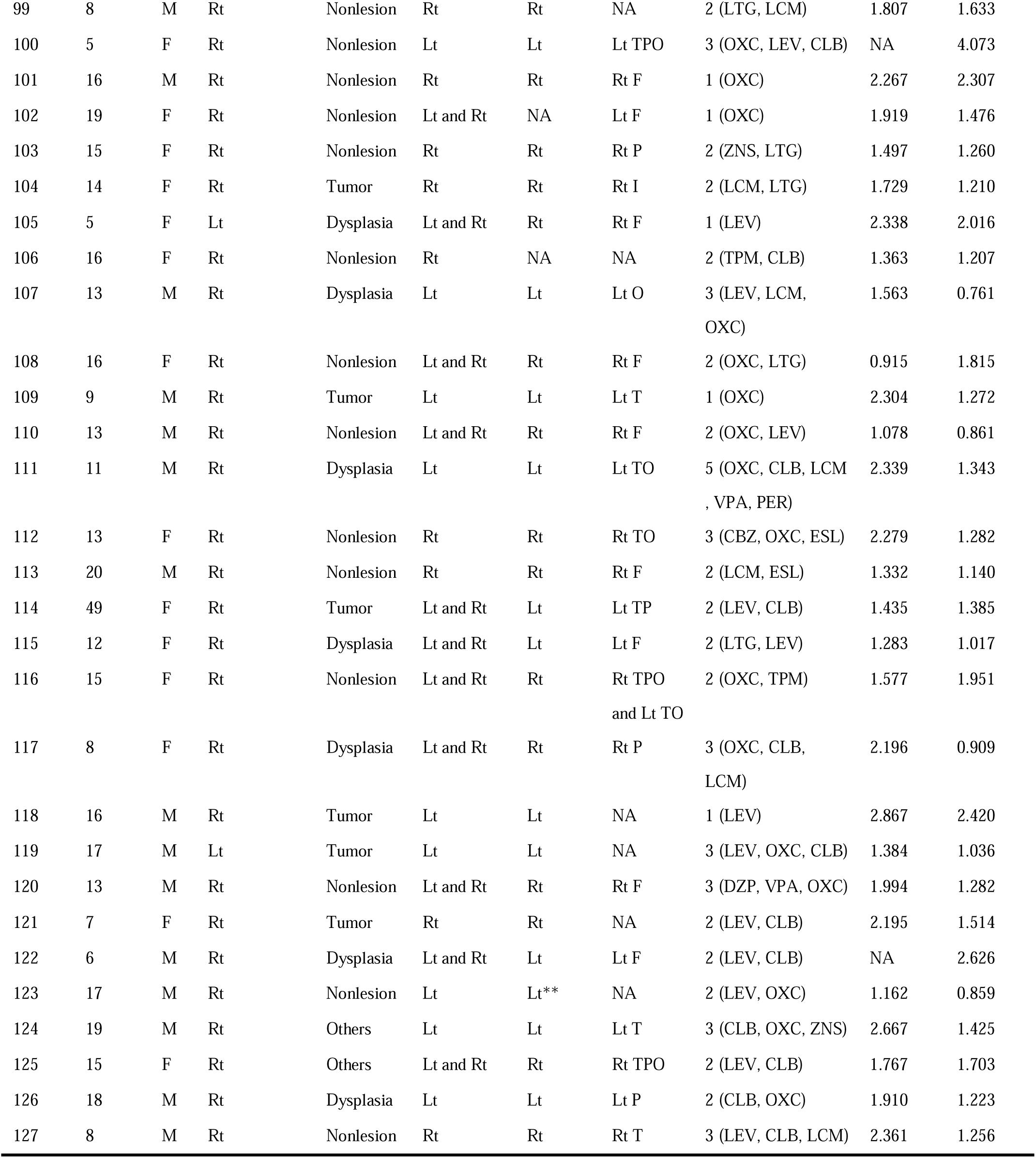
Patient profiles. Sixty-one females (F), and 66 males (M) were included in the study. Lt: left. Rt: right. SOZ: seizure onset zone. NA: not available. MRI lesions other than dysplasia and tumor included focal cortical atrophy (patient 34, 52, 125), hippocampal sclerosis (patients 44, 57, 58, 59, 90, 124), arachnoid cyst (patient 50), and arteriovenous malformations (patient 55). *: multiple subpial transections on the right Rolandic areas. **: responsive neurostimulation employed to the left temporal region. F: Frontal. T: Temporal. O: Occipital. P: Parietal. I: insula. CBZ: Carbamazepine. CLB: Clobazam. CZP: Clonazepam. DZP: Diazepam. ESL: Eslicarbazepine. LCM: Lacosamide. LEV: Levetiracetam. LTG: Lamotrigine. OXC: Oxcarbazepine. PER: Perampanel. PHT: Phenytoin. TPM: Topiramate. VPA: Valproic acid. ZNS: Zonisamide.

**eTable 2.** Number of artifact-free, nonepileptic electrode sites. Numbers in parentheses indicate the number of patients who contributed data. The IG region was excluded from the group-level region-of-interest analysis because fewer than seven patients in a hemisphere provided usable data.

| Auditory naming |  |  | Picture naming |  |  |
| --- | --- | --- | --- | --- | --- |
| Regions of interest | Left | Right | Regions of interest | Left | Right |
| aSFG: anterior superior frontal gyrus | 82 (19) | 143 (35) | aSFG: anterior superior frontal gyrus | 91 (21) | 136 (32) |
| pSFG: posterior superior frontal gyrus | 79 (23) | 99 (35) | pSFG: posterior superior frontal gyrus | 85 (25) | 88 (30) |
| rMFG: rostral middle frontal gyrus | 254 (47) | 339 (51) | rMFG: rostral middle frontal gyrus | 259 (43) | 296 (46) |
| cMFG: caudal middle frontal gyrus | 230 (52) | 259 (54) | cMFG: caudal middle frontal gyrus | 226 (48) | 229 (48) |
| pOp: pars opercularis | 158 (47) | 107 (45) | pOp: pars opercularis | 152 (43) | 96 (37) |
| pTr: pars triangularis | 110 (38) | 180 (44) | pTr: pars triangularis | 108 (41) | 156 (38) |
| pOrb: pars orbitalis | 83 (32) | 83 (26) | pOrb: pars orbitalis | 82 (31) | 72 (24) |
| LOrbF: lateral orbitofrontal gyrus | 120 (38) | 99 (33) | LOrbF: lateral orbitofrontal gyrus | 116 (36) | 97 (30) |
| MOrbF: medial orbitofrontal gyrus | 27 (19) | 40 (24) | MOrbF: medial orbitofrontal gyrus | 28 (20) | 39 (24) |
| aCG: anterior cingulate gyrus | 13 (8) | 18 (9) | aCG: anterior cingulate gyrus | 13 (8) | 17 (9) |
| IG: insular gyrus | 4 (2) | 4 (2) | IG: insular gyrus | 6 (3) | 6 (3) |
| aSTG: anterior superior temporal gyrus | 162 (47) | 224 (50) | aSTG: anterior superior temporal gyrus | 172 (46) | 185 (42) |
| pSTG: posterior superior temporal gyrus | 271 (54) | 207 (49) | pSTG: posterior superior temporal gyrus | 254 (50) | 158 (40) |
| aMTG: anterior middle temporal gyrus | 91 (32) | 89 (32) | aMTG: anterior middle temporal gyrus | 101 (32) | 85 (32) |
| pMTG: posterior middle temporal gyrus | 258 (50) | 131 (39) | pMTG: posterior middle temporal gyrus | 231 (46) | 88 (31) |
| aITG: anterior inferior temporal gyrus | 65 (26) | 99 (35) | aITG: anterior inferior temporal gyrus | 65 (25) | 81 (30) |
| pITG: posterior inferior temporal gyrus | 168 (52) | 105 (41) | pITG: posterior inferior temporal gyrus | 174 (49) | 91 (37) |
| aFG: anterior fusiform gyrus | 44 (24) | 76 (36) | aFG: anterior fusiform gyrus | 47 (24) | 63 (29) |
| pFG: posterior fusiform gyrus | 198 (47) | 161 (47) | pFG: posterior fusiform gyrus | 201 (47) | 153 (42) |
| EG: entorhinal gyrus | 48 (23) | 69 (28) | EG: entorhinal gyrus | 49 (24) | 60 (23) |
| PHG: parahippocampal gyrus | 38 (19) | 27 (20) | PHG: parahippocampal gyrus | 37 (18) | 24 (17) |
| sPreCG: superior precentral gyrus | 113 (40) | 127 (42) | sPreCG: superior precentral gyrus | 115 (42) | 105 (38) |
| iPreCG: inferior precentral gyrus | 362 (59) | 404 (57) | iPreCG: inferior precentral gyrus | 367 (57) | 359 (49) |
| sPoCG: superior postcentral gyrus | 119 (41) | 91 (41) | sPoCG: superior postcentral gyrus | 113 (38) | 88 (35) |
| iPoCG: inferior postcentral gyrus | 328 (59) | 321 (55) | iPoCG: inferior postcentral gyrus | 335 (56) | 282 (49) |
| PCL: paracentral lobule | 32 (15) | 65 (24) | PCL: paracentral lobule | 42 (18) | 54 (21) |
| SPG: superior parietal gyrus | 69 (19) | 57 (19) | SPG: superior parietal gyrus | 65 (18) | 33 (14) |
| IPG: inferior parietal gyrus | 96 (26) | 162 (38) | IPG: inferior parietal gyrus | 91 (26) | 129 (34) |
| SMG: supramarginal gyrus | 353 (56) | 339 (51) | SMG: supramarginal gyrus | 343 (53) | 290 (46) |
| LOG: lateral occipital gyrus | 229 (41) | 217 (42) | LOG: lateral occipital gyrus | 244 (41) | 205 (38) |
| LG: lingual gyrus | 122 (34) | 171 (46) | LG: lingual gyrus | 132 (36) | 160 (42) |
| CG: cuneus gyrus | 23 (12) | 39 (21) | CG: cuneus gyrus | 26 (15) | 35 (18) |
| Pcun: precuneus gyrus | 52 (23) | 75 (29) | Pcun: precuneus gyrus | 49 (24) | 66 (25) |
| pCG: posterior cingulate gyrus | 67 (22) | 83 (27) | pCG: posterior cingulate gyrus | 67 (23) | 73 (25) |
| Total | 4468 (73) | 4710 (72) | Total | 4486 (70) | 4099 (63) |

## Legends for Supplementary Videos: eVideo Legends

**eVideo 1. Neural information flows and functional coactivation/co-deactivation during auditory naming.** Left: Excitatory (red/yellow) and inhibitory (blue/green) flows through white matter tracts. Right: Functional coactivation (red/yellow) and co-deactivation (blue/green).

**eVideo 2. Neural information flows and functional coactivation/co-deactivation during visual naming.** Left: Excitatory (red/yellow) and inhibitory (blue/green) flows through white matter tracts. Right: Functional coactivation (red/yellow) and co-deactivation (blue/green).

**eVideo 3. Neural information flows and cortical high-gamma modulations during auditory naming.** Left: Excitatory (red/yellow) and inhibitory (blue/green) flows through white matter tracts. Right: Cortical high-gamma modulations.

**eVideo 4. Neural information flows and cortical high-gamma modulations during visual naming.** Left: Excitatory (red/yellow) and inhibitory (blue/green) flows through white matter tracts. Right: Cortical high-gamma modulations.

